# Enhancer architecture and chromatin accessibility constrain phenotypic space during development

**DOI:** 10.1101/2022.06.02.494376

**Authors:** Rafael Galupa, Gilberto Alvarez-Canales, Noa Ottilie Borst, Timothy Fuqua, Lautaro Gandara, Natalia Misunou, Kerstin Richter, Mariana R. P. Alves, Esther Karumbi, Melinda Liu Perkins, Tin Kocijan, Christine A. Rushlow, Justin Crocker

## Abstract

Developmental enhancers are DNA sequences that when bound to transcription factors dictate specific patterns of gene expression during development. It has been proposed that the evolution of such cis-regulatory elements is a major source of adaptive evolution; however, the regulatory and evolutionary potential of such elements remains little understood, masked by selective constraints, drift and contingency. Here, using mutation libraries in *Drosophila melanogaster* embryos, we observed that most mutations in classical developmental enhancers led to changes in gene expression levels but rarely resulted in novel expression outside of the native cell- and tissue-types. In contrast, random sequences often acted as developmental enhancers, driving expression across a range of levels and cell-types, in patterns consistent with transcription factor motifs therein; random sequences including motifs for transcription factors with pioneer activity acted as enhancers even more frequently and resulting in higher levels of expression. Together, our findings suggest that the adaptive phenotypic landscapes of developmental enhancers are constrained by both enhancer architecture and chromatin accessibility. We propose that the evolution of existing enhancers is limited in its capacity to generate novel phenotypes, whereas the activity of *de novo* elements is a primary source of phenotypic novelty.

**QUOTE:** “Chance and chance alone has a message for us.” Milan Kundera, *The Unbearable Lightness of Being*

## MAIN TEXT

Morphological changes generally result from changes in the spatiotemporal regulation of gene expression during development, and thus a major theory in evolutionary developmental biology proposes anatomical evolution to be based on the genetic and molecular mechanisms underlying the evolution of spatial gene regulation (Carroll, 2008). In line with this, the evolution of cis-regulatory elements, such as developmental enhancers (Jindal and Farley, 2021), has been proposed to be a major component of phenotypical evolution across animals (Carroll, 2008; Koshikawa, 2015; Majic and Payne, 2020; Monteiro and Gupta, 2016; Nghe et al., 2020; Stern and Orgogozo, 2008). The so-called “cis-regulatory hypothesis” proposes that mutations in enhancers are a common and continuous source of morphological variation, and a means to escape the pleotropic effects of mutations to protein coding regions (Carroll, 2008; Stern and Orgogozo, 2008). For instance, the evolution of wing pigmentation “spots” in *Drosophila* involved the gain of binding sites for different transcription factors in an enhancer controlling a pigmentation gene (Gompel et al., 2005), whereas the loss of pelvic structures in stickleback fish occurred via mutations that abrogate the activity of an enhancer controlling the homeobox gene *Pitx1* (Chan et al., 2010). Molecular mechanisms of cis-regulatory evolution have also been proposed to include duplications of existing enhancers, *de novo* emergence from existing non-regulatory DNA and co-option or exaptation of transposable elements or enhancers with unrelated activities (Emera et al., 2016; Erwin and Davidson, 2009; Fong and Capra, 2022; Indjeian et al., 2016; Koshikawa et al., 2015; Kvon et al., 2021; Long et al., 2016; Lynch et al., 2011; Rebeiz et al., 2011).

Despite elegant case studies, the extent to which these mechanisms contribute to the regulatory evolution of developmental enhancers remains an open question (Arnold et al., 2014; Smith et al., 2013). It is still unknown which changes in enhancer function are evolutionarily accessible, or how the distribution of transcription factor binding sites might constrain the evolutionary potential of enhancers (Fuqua et al., 2020). As such, there is a lack of clarity on the molecular genetic pathways for evolutionary change in animal development based on what is functionally possible *versus* what is probable and permissible from the standpoint of mutational events and natural selection (Carroll, 2008).

Here, we explored how molecular evolution of existing enhancers versus *de novo* sequences contributes to producing novel patterns of gene expression across *Drosophila melanogaster* embryos. We generated and characterized a panel of unbiased mutation libraries for both classical developmental enhancers and randomly generated sequences; this approach allows to distinguish constraints that emerge from the prior function or evolutionary histories of existing enhancers from constraints that arise from properties of the sequence or locus unrelated to selection processes.

### Constrained capacity for enhancer-driven expression outside of native expression patterns

We first set out to investigate whether and how mutations across developmental enhancers could lead to ectopic, novel expression patterns. We have previously generated a mutation library for the *E3N* enhancer, which regulates the expression of *shavenbaby* (**Fig. 1A-B**) (Fuqua et al., 2020). This mutation library included 749 variants and most mutations led to changes in transcriptional outputs (e.g., levels, location) (Fuqua et al., 2020). This library represents a ∼6 times larger sequence space than the natural variation found for *D. melanogaster E3N* from samples across the world (**Fig. 1C-F, Fig. S1**). To investigate novel expression patterns, we selected a subset of lines harboring 1-10 point mutations for further characterization; these lines come from different regions of the sequence space covered by the total library (**Fig. 1F**; **Table S1**) and showed a spectrum of effects in terms of expression levels (**Fig. 1G**). We found that 22% of the lines showed expression outside of the usual *E3N-*driven ventral stripes, in regions such as prospective anal pads, wing and haltere imaginal discs and other structures (**Fig. 1H-K**). However, these regions are ectoderm-derived and correspond to regions where the target gene of *E3N* (*svb*) is expressed (Frankel et al., 2010; Preger-Ben Noon et al., 2018).

**Figure 1:**
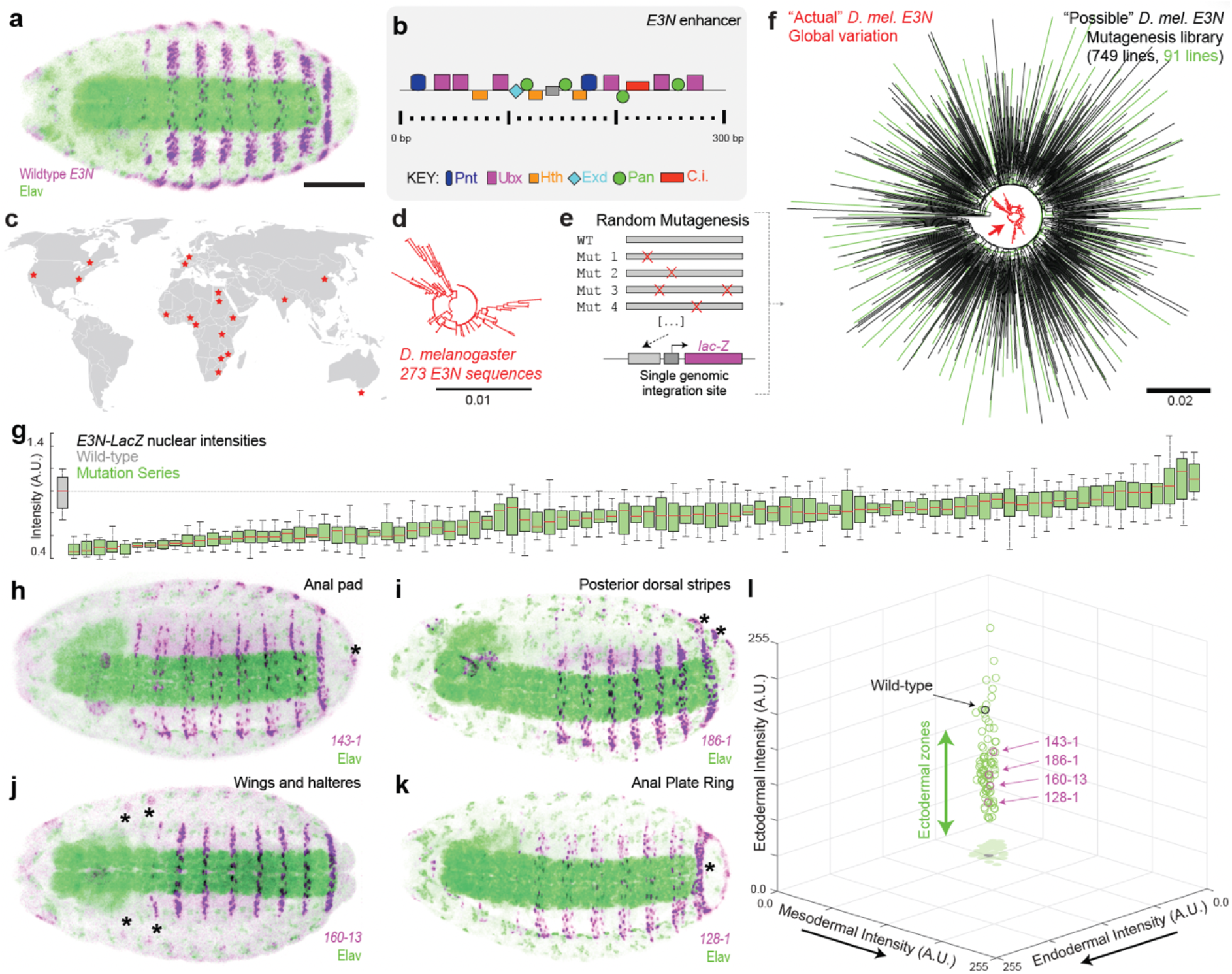
Mutant variants of the *E3N* enhancer have a limited capacity for expression outside native tissues- and cell-types. (**A**) Pattern of expression driven by wildtype *E3N* at stage 15 (beta-galactosidase protein staining). Scale bar 100 μm. (**B**) Mapped binding site architecture for *E3N*. (**C**) Collection locations of sequenced *Drosophila melanogaster* strains (Lack et al., 2016). (**D**) Phylogenetic tree of *E3N* sequences across *D. melanogaster* strains. (**E**) Schematic of enhancer variants and reporter gene construct used for integration into the *D. melanogaster* genome. (**F**) Phylogenetic tree of *E3N* sequences across *D. melanogaster* strains (red) and of *E3N* sequences from our mutational library (black and green; in green, 91 lines selected for further characterization). (**G**) Nuclear intensities of the A2 segment across 91 lines, normalized to wildtype *E3N* (n=10 embryos per line). A.U., arbitrary units of fluorescence intensity. (**H-K**) Examples of mutant variants leading to reporter expression outside the wildtype *E3N* pattern. In panels (h) and (i), expression associated to esophagus is likely an artifact of the construct used, as observed in other lines unrelated to *E3N*. (**L**) 3D plot showing fluorescence intensities for 91 lines across three regions of the embryo with different germ-layer origins (see Fig. S2). Each dot corresponds to the average value for one variant enhancer line.

To evaluate ectopic expression across regions derived from different germ layers, we quantified reporter-expression intensity in the selected lines (**Fig. S2**) and detected no expression in regions derived from germ layers other than the ectoderm (**Fig. 1L**), whereas variable levels of expression along the “ectoderm” axis could be seen (**Fig. 1L**). These results suggest that evolving new patterns of expression upon point mutations of a developmental enhancer is possible but developmentally biased to specific lineages.

### The emergence of ectopic expression patterns upon mutagenesis of developmental enhancers is rare

To explore whether the transcriptional constraints we observed for *E3N* mutagenesis are a general property of developmental enhancers, and given that *E3N* regulates a terminal selector gene in later development (Allan and Thor, 2015), we chose to explore additional “classical” enhancers involved in early development. These include *eveS2*, important for anterior-posterior specification (**Fig. 2A-B**), (Small et al., 1991, 1992; Stanojevic et al., 1991), and *rhoNEE* and *twiPE*, both involved in dorsoventral patterning (**Fig. 2E-G**), in the neurogenic ectoderm and mesoderm, respectively (Bier et al., 1990; Ip et al., 1992; Jiang et al., 1991; Markstein et al., 2004; Pan et al., 1991; Thisse et al., 1991). For each of these enhancers, we generated mutant libraries using the same setup as for the *E3N* library (Fuqua et al., 2020): each variant was cloned upstream of a heterologous *hsp70* promoter driving *lacZ* reporter expression and integrated into the *Drosophila* genome at a specific landing site, amenable to expression across different tissues and stages (**Fig. S1**). Using a PCR error-rate of ∼0.5% per molecule, we isolated enhancer variants containing approximately 1-5 mutations in 12-36 independent fly lines per enhancer (Table S1).

**Figure 2:**
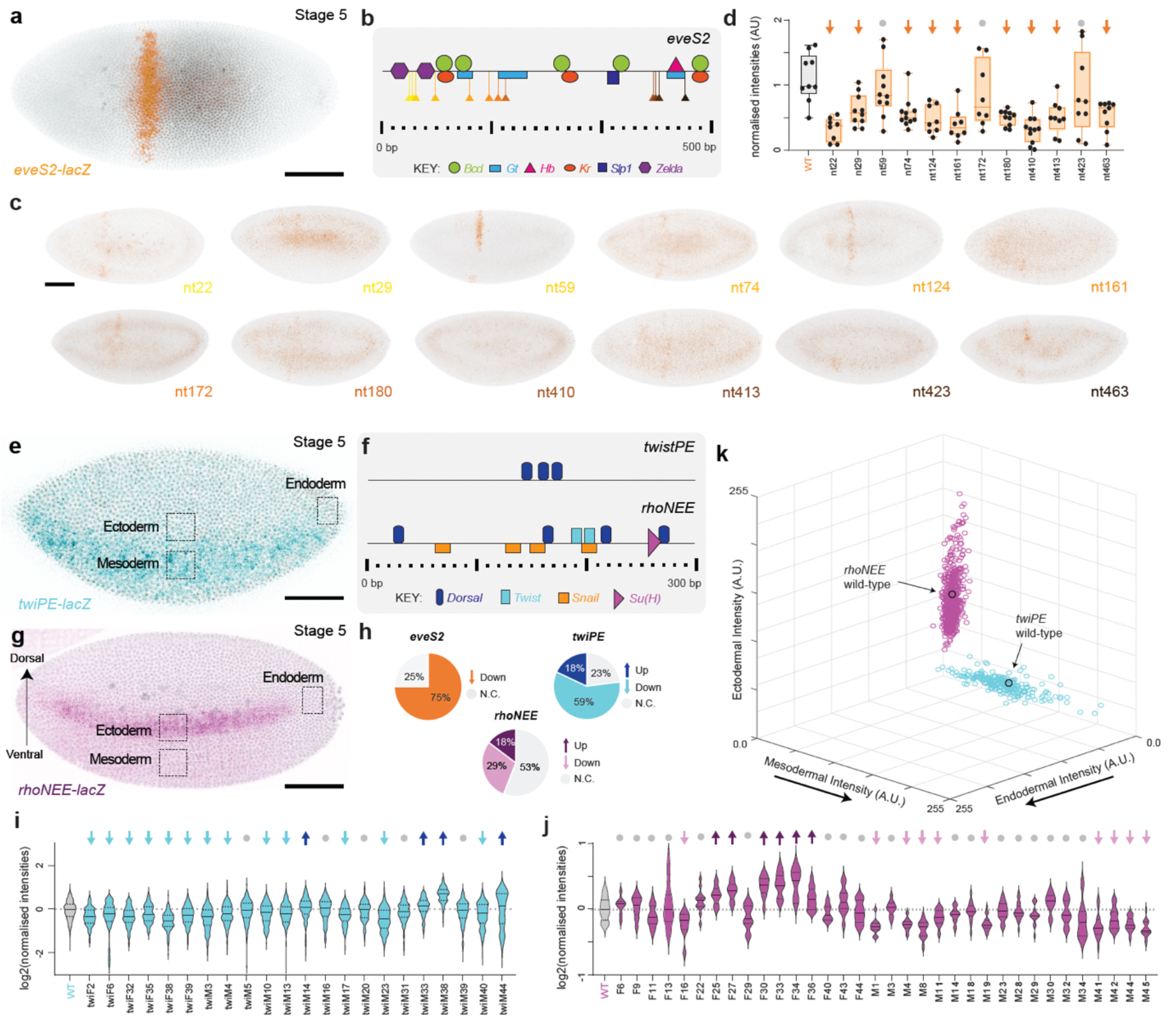
Mutagenesis across early developmental enhancers alters gene expression only within native patterns of expression. (**A**) Pattern of expression driven by wildtype *eveS2* at stage 5 (lacZ mRNA staining). Scale bar 100 μm. (**B**) Known binding site architecture for *eveS2*. Location of point mutations is indicated. (**C**) Examples of stained embryos from different *eveS2* single-nucleotide mutant variants. The name of each line corresponds to the location of the point mutation (compare with **B**). (**D**) Fluorescence intensities of the region where the wildtype *eveS2* shows a stripe across 12 single-nucleotide *eveS2* variants (n=8-11 embryos per line). Lines marked with an arrow are statistically significantly different from wildtype (p<0.05; two-tailed t-test). A.U., arbitrary units of fluorescence intensity. (**E**) Pattern of expression driven by wildtype *twiPE* at stage 5 (lacZ mRNA staining). (**F**) Known binding site architecture for *twiPE* and *rhoNEE*. (**G**) Pattern of expression driven by wildtype *rhoNEE* at stage 5 (lacZ mRNA staining). (**H**) Summary of changes in expression levels for the *eveS2, twiPE* and *rhoNEE* lines. (**I-J**) Nuclear intensities across *twiPE* (**I**) and rhoNEE (**J**) variants (n=6-27 embryos per line). Lines marked with an arrow (up or down) are statistically significant from wildtype (p<0.05; two-tailed t-test). (**K**) 3D plot showing fluorescence intensities for *twiPE* (blue) and *rhoNEE* (purple) lines across three regions of the embryo illustrated in (**I**) and (**J**). Each dot corresponds to one embryo; three embryos per line were quantified.

We examined reporter activity across all lines in the early embryo (stage 5) and found similar trends for all of them. On the one hand, mutations often led to significant changes in expression levels, and on the other hand, changes in expression were restricted to the native pattern – no ectopic expression was observed. For *eveS2* (**Fig. 2A**), each variant contained a single mutation only, almost none overlapping a known binding site (**Fig. 2B-C**). Yet, 75% led to significantly reduced expression compared to control (**Fig. 2D, 2H**), suggesting that it is relatively easy to “break” the minimal eveS2 enhancer, consistent with unsuccessful attempts to build this enhancer *de novo* (Crocker and Ilsley, 2017; Vincent et al., 2016). In no case did we observe expression outside of the eve stripe 2 region. Similar results were found for *rhoNEE* and *twiPE*: 47% and 77% of enhancer variants, respectively, showed statistically significant changes in nuclear intensities compared to control (**Fig. 2H**); for *rhoNEE*, 18% showed higher expression and 29% showed lower expression (**Fig. 2J**); for *twiPE*, these values were 18% and 59% respectively (**Fig. 2I**). These effects did not seem to correlate with the number of mutations per enhancer (**Fig. S3**) nor with the length of the enhancer (compare Fig. 2b and 2f with 2h). Again, despite clear changes in levels for most mutant variants, we noted that expression outside of the typical area of expression for each enhancer was never observed – quantification of expression in control and mutant lines across regions of the embryo that will give rise to ectoderm (lateral region of the embryo), endoderm (posterior region of the embryo) and mesoderm (ventral region of the embryo; regions highlighted in **Fig. 2E, 2G**) revealed that mutant lines showed changed levels of expression but always within the “ectoderm” and “mesoderm” regions only, for *rhoNEE* and *twiPE* enhancers respectively (**Fig. 2K**). In summary, most mutations led to changes in expression levels within native zones of expression; thus, the results suggest that the “molecular evolution” by point mutations of developmental enhancers is not likely to result in novel expression patterns.

Considering that such pleiotropic effects could be revealed throughout development (Preger-Ben Noon et al., 2018), we analyzed expression in embryos at later stages (stage 9 and 14) for the *rhoNEE* (**Fig. 3A; Fig. S4**) and *twiPE* libraries (**Fig. 3E; Fig. S5**) but we observed no ectopic expression in the mutant lines compared to the control (**Fig. 3B-D, 3F-G**). We also generated an additional mutational library for *tinB*, an enhancer that controls a mesoderm-specific gene throughout a broad developmental window (**Fig. 3H-I;** Table S1) (Yin et al., 1997; Zaffran et al., 2006). Similar to what we found for early enhancers, 47% of enhancer variants showed significant changes in enhancer activity (**Fig. 3J**; 20% showed increased expression, 27% showed decreased expression), yet no ectopic expression was observed (**Fig. 3K**).

**Figure 3:**
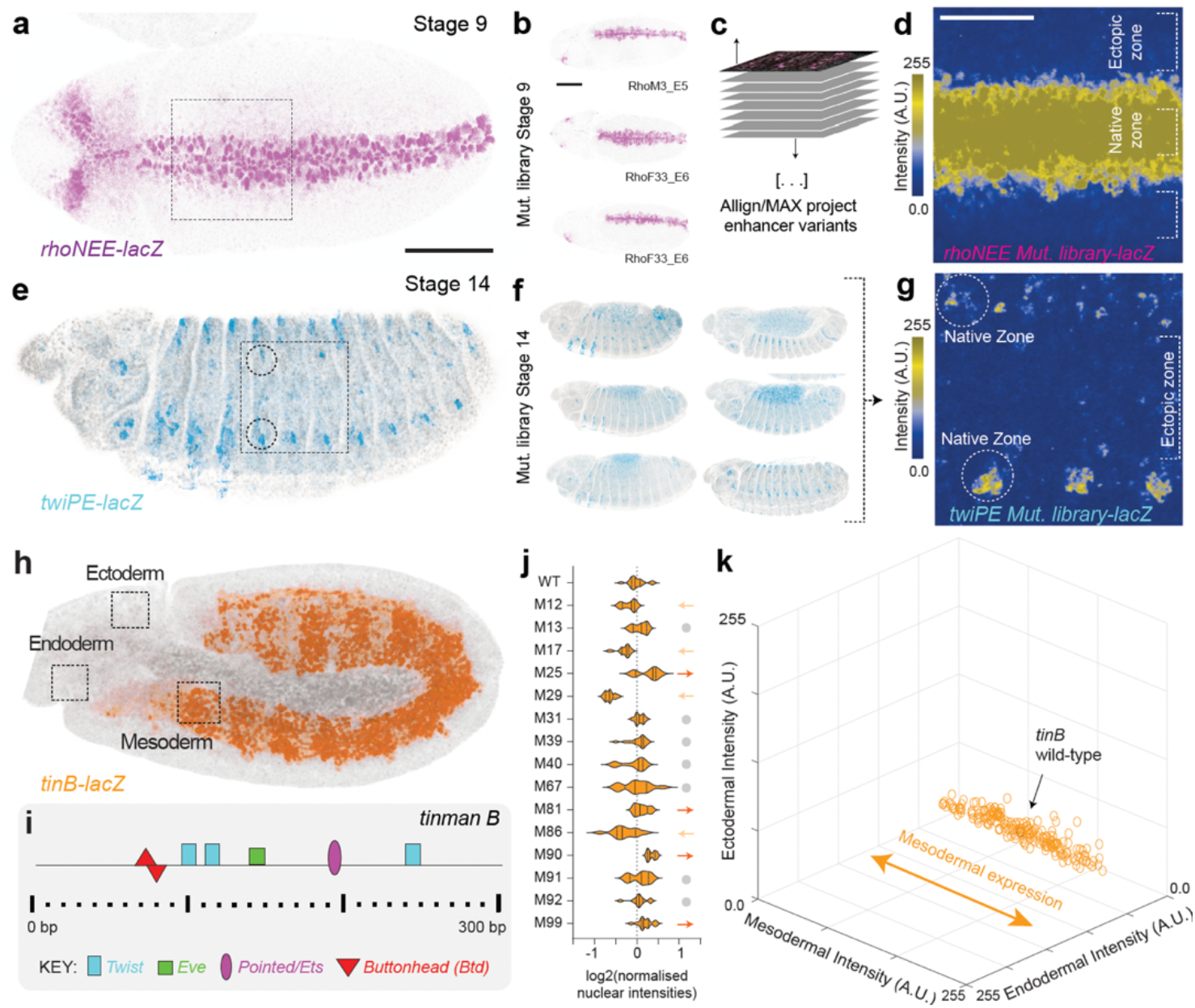
Mutagenesis across late developmental enhancers alters gene expression only within native patterns of expression. (**A**) Pattern of expression driven by wildtype *rhoNEE* at stage 9 (beta-galactosidase protein staining). Scale bar 100 μm. (**B**) Examples of stained embryos from different *rhoNEE* mutant variants. Scale bar 100 μm. (**C**) Schematic of alignment and overlaying of individual Z-projections of maximum intensity for *rhoNEE* mutant variants. (**D**) Heatmap of aggregated Z-projections. Scale bar 50 μm. (**E**) Pattern of expression driven by wildtype *twiPE* at stage 14 (beta-galactosidase protein staining). (**F**) Examples of stained embryos from different *twiPE* mutant variants. (**G**) Heatmap of aggregated Z-projections upon alignment of individual Z-projections of maximum intensity for *twiPE* mutant variants. (**H**) Pattern of expression driven by wildtype *tinB* at stage 10 (beta-galactosidase protein staining). (**I**) Known binding site architecture for *tinB*. (**J**) Nuclear intensities across *tinB* variants (n=10-18 embryos per line). Lines marked with an arrow (up or down) are statistically significant from wildtype (p<0.05; two-tailed t-test). (**K**) 3D plot showing fluorescence intensities for *tinB* lines across three regions of the embryo as illustrated in (**H**). Each dot corresponds to one embryo; at least ten embryos per line were quantified.

Finally, we tested whether ectopic expression could be “forced” upon recruitment of a ubiquitously expressed synthetic transcription factor. The *rhoNEE* enhancer has been previously engineered to contain binding sites for a transcription activator-like effector (TALE) DNA-binding protein (Crocker et al., 2016). We crossed fly lines harboring *rhoNEE* enhancers with one, two or three TALE binding sites with a line containing a TALE protein fused to the strong activation domain VP64 (Beerli et al., 1998) and expressed via a nos::Gal4 driver, and quantified expression across different regions of the early embryo (**Fig. S7**). The higher the number of binding sites for the synthetic transcription factor, the higher the expression within the usual regions of *rhoNEE* expression. However, it was not until there were two or more binding sites (16bp long) that appreciable expression was generated outside of the native zones of expression (**Fig. S7**). Together, these results reveal that the *rhoNEE* enhancer is not ‘intrinsically’ refractory to expression outside of its usual pattern of expression, but rather requires a considerably larger recruitment of activators to the locus. The fact that we do not observe ectopic expression in the enhancer libraries analyzed suggests that regulatory constraints are imposed on developmental enhancers.

### Random sequences lead to extensive expression across developmental time and space

We interrogated the extent to which *de novo* sequences, devoid of evolutionary constraints, could act as enhancers and drive expression across the embryo and across development. We synthesized random sequences (∼180bp), inserted them upstream of *hsp70* promoter driving *lacZ* (similarly to the enhancer libraries) and integrated them into the fly genome at the same genomic location (**Fig. 4A, Fig. S8**). These sequences included a motif (UAS) for the yeast Gal4 transcription factor (Kakidani and Ptashne, 1988; Webster et al., 1988), which is not present in the fly and thus this motif should be “neutral”; this design was chosen so that these sequences have a comparable architecture to libraries containing other motifs (see later). We isolated 56 fly lines harboring unique sequences (Table S1), for which we stained embryos at different stages to determine reporter gene’s expression pattern(s). Surprisingly, 86% of sequences led to changes in reporter expression at least in some cells and/or at some developmental stage, compared to expression of the reporter with no sequence cloned upstream (**Fig. 4B-D**; **Fig. S9**). The other surprising observation was that despite such pervasive expression, we never observed expression in the early embryo (**Fig. 4C**). Given the variable consensus sites found in multicellular systems, such libraries are expected to have a range of motifs with variable information content (de Boer et al., 2019; Wunderlich and Mirny, 2009) (**Fig. 4E**). To explore the expression patterns observed, we conducted motif searches across all random sequences for *Drosophila* developmental transcription factors (**Fig. 4E**; Methods). Motifs found included Ultrabithorax (Ubx), GATA, Grainyhead (Grh) and Bicoid (Bcd) motifs (**Fig. 4F-I**). Interestingly, 100% or 80% of the random DNA elements containing, respectively, a GATA or Grh motif showed expression (**Fig. 4G-H**), consistent with their previously reported predictive power (de Almeida et al., 2022; Kvon et al., 2014) and with the expression patterns of the respective transcription factors. In contrast, only 14% of elements with a Ubx motif showed expression (**Fig. 4F**) and none of the elements containing a Bcd motif showed expression (**Fig. 4I**), consistent with the absence of expression in the early embryo for all random sequences. We calculated whether our random sequences were biased for motifs of late-development transcription factors (TFs), but this did not explain the absence of early expression (average per sequence: ∼3.9 hits per early-specific motif *versus* ∼3.4 hits per late-specific motif; see Methods).

**Figure 4:**
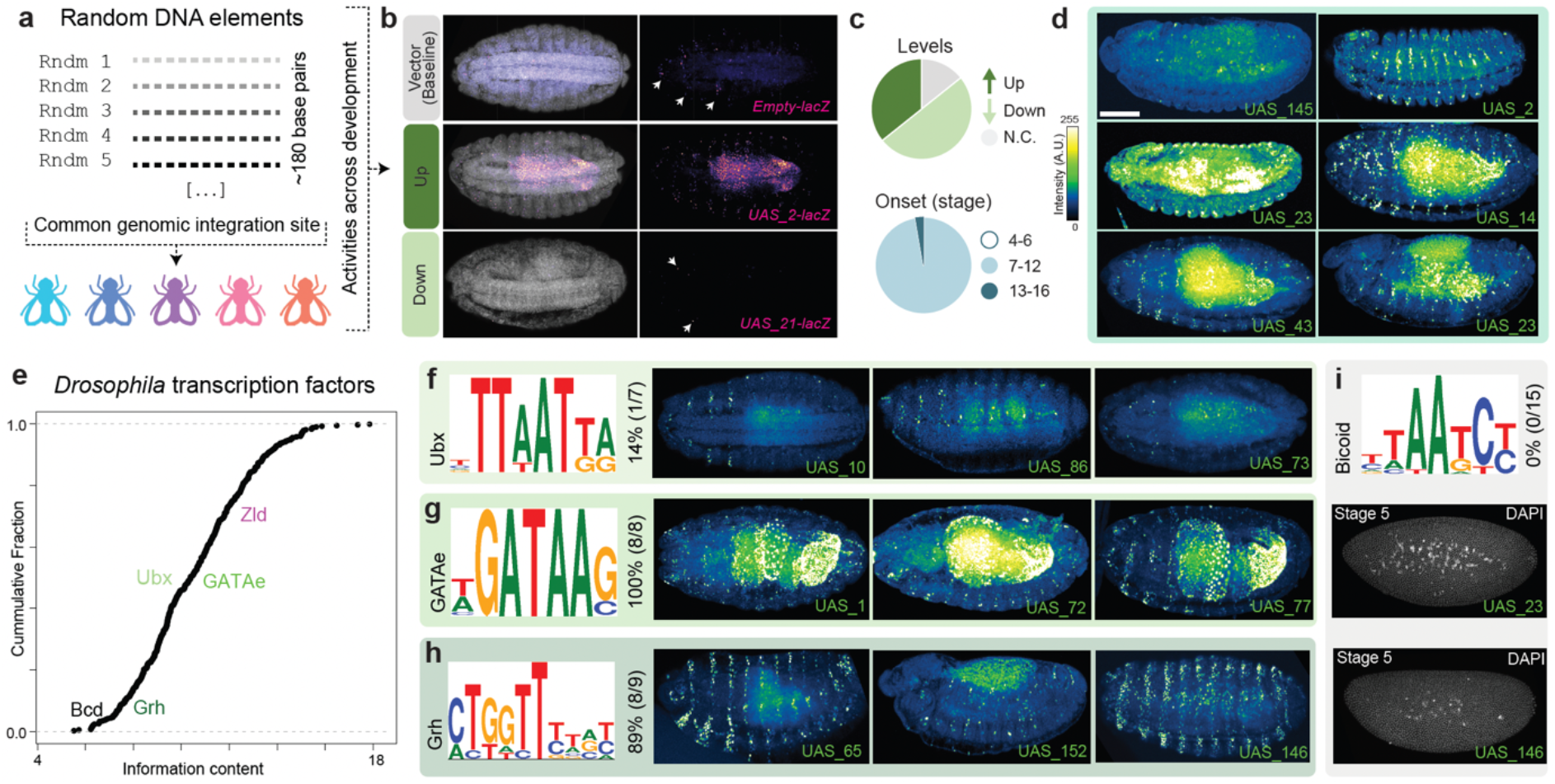
Random DNA sequences often drive reporter expression during development. (**A**) Schematic of the UAS-library. (**B**) Expression patterns at stage 15 were compared to the reporter with no sequence cloned upstream (top) and classified as “up” (middle) or “down” (bottom), depending on whether expression was increased or decreased, respectively. (**C**) Summary of changes in expression levels at stage 15 (top) based on panel (**B**), and of developmental period in which expression is first observed (bottom). (**D**) Examples of stained embryos from different random DNA sequences. (**E**) Cumulative distribution function of the expected frequency of *Drosophila* TF motifs in random DNA. (**F**) Ubx motif, percentage of lines showing expression among random DNA lines with a Ubx motif and examples of corresponding embryos. (**G**) GATA motif, percentage of lines showing expression among random DNA lines with a GATA motif and examples of corresponding embryos. (**H**) Grh motif, percentage of lines showing expression among random DNA lines with a Grh motif and examples of corresponding embryos. (**I**) Bicoid motif, percentage of lines showing expression among random DNA lines with a Bcd motif and examples of corresponding embryos.

### Specific motifs can potentiate emergence of enhancer activity

Completely random sequences thus seem to have a high potential of driving expression, and this can be associated to particular motifs. Given the association between chromatin accessibility and transcriptional permissiveness (Klemm et al., 2019), as well as studies suggesting that chromatin accessibility might underlie enhancer evolution (Peng et al., 2019; Xin et al., 2020), we generated “biased” random libraries in which we included a Grh motif (**Fig. 5A**; 7 lines, Table S1) or a Zelda motif (**Fig. 5E**; 41 lines, Table S1) approximately at the center of random sequences. Grainyhead and Zelda are transcription factors in the fly reported to have “pioneer activity” (Hansen et al., 2022; Zaret and Carroll, 2011) – their binding is associated with “opening” chromatin, rendering enhancers more accessible to binding by other transcription factors (Foo et al., 2014; Harrison et al., 2011; Iwafuchi-Doi, 2019; Jacobs et al., 2018; Larson et al., 2021; Nevil et al., 2020; Schulz et al., 2015; Sun et al., 2015). Though Zelda is usually associated with early fly development, it is expressed throughout development (**Fig. S10**) and its late embryonic knockout has phenotypical consequences (**Fig. S11**). Consistent with the idea of “pioneer activity”, an even higher proportion of random sequences from the Grh and Zld “biased” libraries drove expression compared to the UAS library (**Fig. 5B-C, 5F-G; Fig. S12**). Not only a higher number of lines was associated with expression for the “biased” libraries, but also expression levels were higher when compared to the UAS-library, regardless of the region of the embryo (**Fig. 5D, 5H-I**). To further test the potential of these motifs, we added one or two Zelda motifs to the developmental enhancers we tested initially (*eveS2, rhoNEE, twiPE, tinB*) and found a significant increase in reporter expression levels for all enhancers within their native patterns of expression (**Fig. S13**). For the *eveS2* lines, we additionally observed novel, ectopic expression (**Fig. S13**), suggesting that the Zelda motifs might “unlock” cryptic sites contained in *eveS2*. We tested whether *eveS2* contained more predicted motifs than the other enhancers, but we did not find any significant differences in the number of hits (0.07 for *eveS2* versus 0.10, 0.12 and 0.05 for *rhoNEE, tinB* and *twiPE*, respectively; normalized per enhancer length).

**Figure 5:**
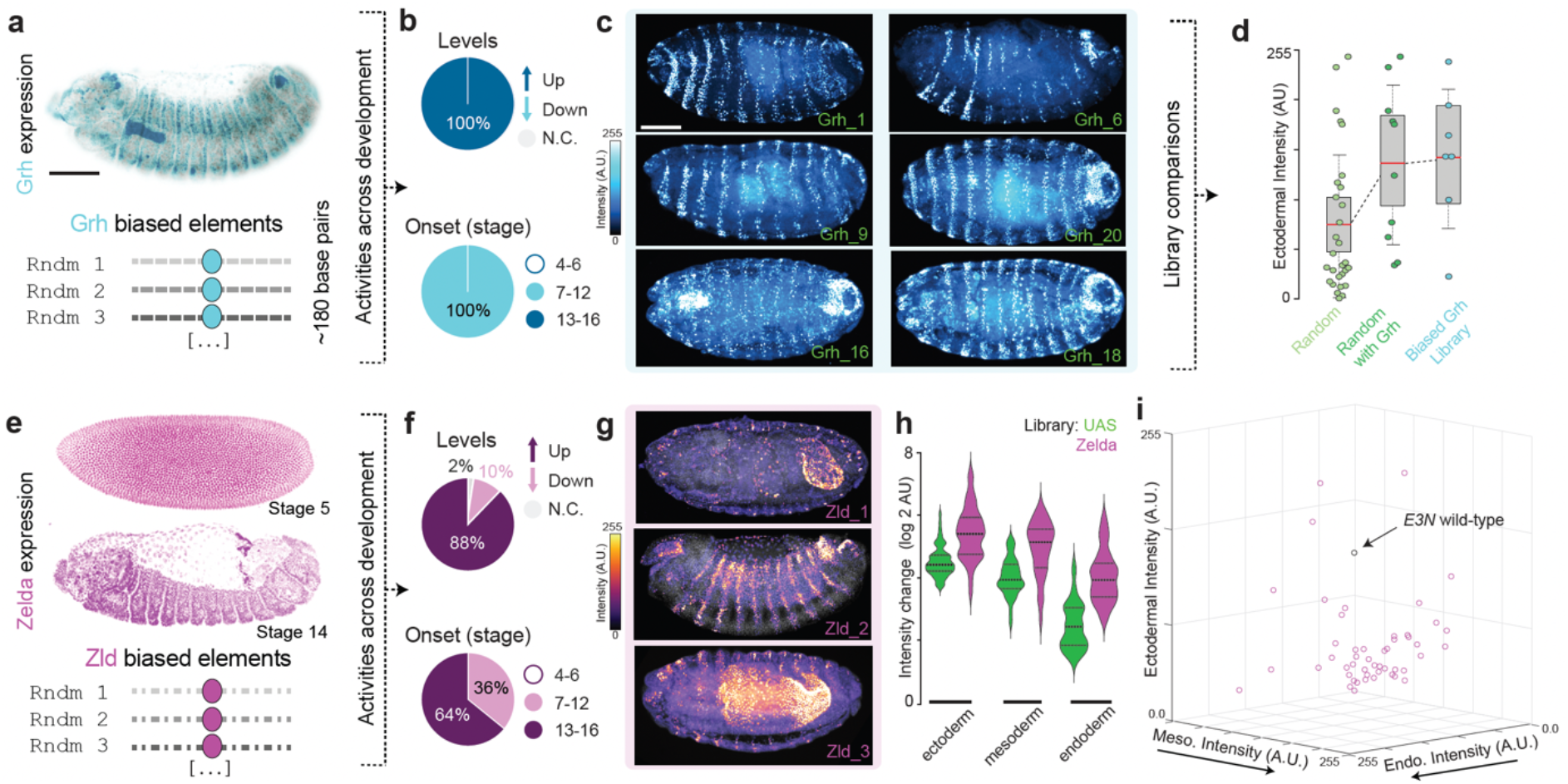
Specific DNA motifs enhance likelihood of reporter expression during development. (**A**) Staining for Grh transcription factor (top) and schematic of the Grh-library (bottom). (**B**) Summary of changes in expression levels (top) compared to the reporter with no sequence cloned upstream (Fig. S9) and of developmental period in which expression is first observed (bottom). (**C**) Examples of stained embryos from different Grh-biased sequences. (**D**) Quantification of fluorescent intensities in ectoderm-associated region for all random DNA sequences, for random DNA sequences with Grh motifs (subset of all random DNA sequences) and for Grh-biased sequences. (**E**) Staining for Zld transcription factor (top) and schematic of the Zld-library (bottom). (**F**) Summary of changes in expression levels at stage 15 (top) compared to the reporter with no sequence cloned upstream (Fig. S9) and of developmental period in which expression is first observed (bottom). (**G**) Examples of stained embryos from different Zld-biased sequences. (**H**) Quantification of fluorescent intensities for Zld-biased lines across three regions of the embryo (see Fig. S2). (**I**) 3D plot showing fluorescence intensities for Zld-biased lines, based on (**H**). Each dot corresponds to one line. For reference, fluorescence intensity for the wildtype *E3N* sequence is shown (from Fig. 1L).

To explore the possibility that the occurrence of specific motifs throughout the genome might contribute to the emergence of (*de novo*) enhancers, we selected genomic sequences containing high-affinity Ubx/Hth motifs (ATGATTTATGAC) (Slattery et al., 2011) present in *D. melanogaster* but not in other *Drosophila* species (**Fig. S14**). Such motifs have been demonstrated to augment chromatin accessibility (Loker et al., 2021) and are broadly used across development, providing a counterpoint to our synthetic libraries. Strikingly, when we tested their enhancer potential with the *lacZ* reporter assay, all sequences showed enhancer activity (**Fig. S14**). Mutating the Ubx/Hth motif in each of those sequences led to a dramatic reduction in expression for six out of seven of those sequences (**Fig. S14**), indicating that these motifs clearly have the capacity to drive expression across development. These results support the idea that specific sequence motifs might prime genomic sequences to act and/or evolve as enhancers.

## DISCUSSION

We used transgenesis-based mutagenesis and *de novo* gene synthesis during fly embryogenesis to investigate evolutionary pathways for enhancer activity. We used fly development to explore how novel patterns of gene expression might appear from either “molecular evolution” of developmental enhancers or random sequences. Notably, while reporter gene assays and minimal enhancers may not reflect the full regulatory activities of native loci (Halfon, 2019; Lindhorst and Halfon, 2022; López-Rivera et al., 2020), such an approach allows us to evaluate a broad range of “possible” enhancer variation in a controlled experimental setup, without associated fitness costs and allowing a broader exploration of evolution and development without the complexities and historical contingencies found in nature. Furthermore, using such an assay in a developmental model system, which generates an embryo in 24 hours, we can assay regulatory activities across ∼100,000 cells of different lineage origins (Song et al., 2019).

Using this approach, we found that most mutations in enhancers led to changes in levels of reporter gene expression, but almost entirely within their native zones of expression (**Figs. 1-3**), similar to previous studies using transgenic mutagenesis of the *Shh* enhancer in murine embryos (Kvon et al., 2020), or the *E3N* enhancer (Fuqua et al., 2020) and the wing spot^196^ enhancer (Le Poul et al., 2020) in fly embryos. Consistent with our results, known phenotypic evolution through nucleotide mutations of standing regulatory elements seems to appear either through changes in the levels or timings of expression within native zones or the loss of regulatory activities. For example, the evolution of pigmentation spots in fly wings occurred via a specific spatial increase in the melanic protein Yellow, which is uniformly expressed at low levels throughout the developing wings of fruit flies (Gompel et al., 2005); see (Frankel et al., 2011; Rebeiz et al., 2009) for other examples of evolution within native patterns of expression. Evolution of other traits such as thoracic ribs in vertebrates (Guerreiro et al., 2013), limbs in snakes (Kvon et al., 2016), pelvic structures in sticklebacks (Chan et al., 2010) and seed shattering in rice (Konishi et al., 2006) are all associated with loss of enhancer activity due to internal enhancer mutations. Additionally, mutations have been found to occur less often in functionally constrained regions of the genome, suggesting that mutation bias may reduce the occurrence of deleterious mutations in regulatory regions (Monroe et al., 2022).

Consistent with these results, phenotypic novelties underlain by enhancer-associated ectopic gains of expression are reportedly due to transposon mobilisation (Bourque et al., 2008; Emera and Wagner, 2012; Feschotte, 2008; Oliver and Greene, 2009), rearrangements in chromosome topology (Galupa and Heard, 2017; Gilbertson et al., 2022; Lupiáñez et al., 2016) or *de novo* evolution of enhancers from DNA sequences with unrelated or nonregulatory activities (Arnold et al., 2014; Birnbaum et al., 2012; Eichenlaub and Ettwiller, 2011; Emera et al., 2016; Li et al., 2022; Prabhakar et al., 2008; Rebeiz et al., 2011). Previous studies have explored the potential of random DNA sequences to lead to reporter gene expression, either as enhancers or promoters, especially in cell lines of prokaryotic or eukaryotic origin (de Almeida et al., 2022; Vaishnav et al., 2022; Yona et al., 2018). These have shown that there is a short (or sometimes null) mutational distance between random sequences and active cis-regulatory elements (Yona et al., 2018), which may improve evolvability. In our study, we tested random sequences in a developmental context and found that most showed enhancer activity across several types of tissues and developmental stages (**Fig. 4**). These results are consistent with a study that tested enhancer activity of all 6-mers in developing zebrafish embryos and found a diverse range of expression for ∼38% of the sequences at two developmental stages (Smith et al., 2013). We observed expression driven by random sequences even in the absence of motifs within their sequence for transcription factors with pioneering activity (**Fig. 4**). Yet, when such motifs were included, nearly all sequences acted as “strong” enhancers (leading to high levels of expression) (**Fig. 5**), consistent with the “evolutionary barrier” to the formation of a novel enhancer being lower in regions that already contain motifs for DNA binding factors, which can “act cooperatively with newly emerging sites” (Long et al., 2016).

It is interesting to note that, despite the high potential of random sequences to be expressed during development and across cell types, we never observed expression prior to gastrulation; this was not evaluated in the zebrafish study or in other studies. This may be due to the rapid rates of early fruit fly development, in which gene expression patterns are highly dynamic, and cell-fate specifications occur within minutes (Surkova et al., 2018). As such, there may be extensive regulatory demands placed on transcriptional enhancers, reflected in the clusters of high-affinity binding sites common across early embryonic developmental enhancers (Crocker et al., 2015) as well as their extensive conservation in function (Hare et al., 2008) and location (Cande et al., 2009). In the future, it will be interesting to explore how regulatory demands that change across development – such as nuclear differentiation, network cross-talk, and metabolic changes – are reflected in regulatory architectures and their evolvability.

The observation that most random sequences led to expression suggests that the potential of any sequence within the genome to drive expression is enormous and thus “an important playground for creating new regulatory variability and evolutionary innovation” (Eichenlaub and Ettwiller, 2011). This was further supported by the regulatory potential of the genomic sequences we tested, containing Ubx/Hth motifs. Perhaps the challenge from an evolutionary perspective has not been what allows expression, but what prevents expression; thus, mechanisms that repress “spurious” expression might have evolved across genomes. This is in line with propositions that nucleosomal DNA in eukaryotes has evolved to repress transcription (Muers, 2013; Wade and Grainger, 2018), along with transcriptional repressors and other mechanisms such as DNA methylation, as a response (at least partially) to “the unbearable ease of expression” present in prokaryotes (Gophna, 2018). The action of such repressive mechanisms could also explain why mutagenesis of developmental enhancers, which are subject to evolutionary selection, does not easily lead to expression outside their native patterns of expression. In sum, our findings raise exciting questions about the evolution of enhancers and the emergence of novel patterns of expression that may underlie new phenotypes, suggesting an underappreciated role for *de novo* evolution of enhancers by happenstance. Genetic theories of morphological evolution will benefit from comparing controlled, multi-dimensional laboratory experiments with standing variation (Laland et al., 2015); such an integrative approach could provide the frameworks that will enable us to make both transcriptional and evolutionary predictions.

## METHODS

### Fly strains and constructs

All mutant and random enhancer sequences were synthesized and cloned (GenScript) into pLacZattB plasmid at HindIII/XbaI site. *E3N*- and *eveS2*-related lines were injected into attP2 line, all other constructs were injected into VK33 line; injections done by Genetivision. Transgenic lines were homozygosed and genotyped; sequences are listed in Table S1.

### Embryos collection and fixation

Flies were loaded into egg collection chambers, left to acclimatize for 3-4 days and then embryos were collected for either four or sixteen hours, for early and late stages, respectively. Embryos were dechorionated in 5% bleach for 2min, abundantly rinsed with water and washed in a saline solution (0.1 M NaCl and 0.04% Triton X-100), before transfer to scintillation vials containing fixative solution (700 μl 16% PFA, 1.7 ml PBS/EGTA, 3.0 ml 100% heptane). Embryos were fixed for 25 min, shaking at 250 rpm. The lower phase was then removed, 4.6 mL 100% methanol added and vials vortexed at maximum speed for 1min. The interphase and upper phase were removed and the embryos were washed thrice in fresh methanol. Embryos were stored at -20 ºC until processed.

### Reporter gene expression analysis

#### In situ hybridization (probes)

probes for *lacZ* (reporter) and *snail* (internal control) were generated from PCR products using the in vitro transcription (IVT) kit from Roche (#11175025910) and following manufacturer’s instructions. A list of primer sequences for each PCR product can be found in Table S1. For each gene, distinct PCR products were pooled before IVT reaction. Probes were diluted in hybridization buffer (Hyb; 50% formamide, 4X SSC, 100 μg/mL salmon DNA, 50 μg/mL heparin, 0.1% Tween-20) at 50ng/μL. Prior to hybridization, a probe solution was prepared (per sample, 50 ng of each probe in 100 μL), denatured at 80 ºC for 5min, then immediately put on ice for 5min, and finally incubated at 56 ºC for 10min before added to the embryos.

#### In situ hybridization (procedure)

embryos stored in methanol were washed in methanol/ethanol (50:50), three-times in 100% ethanol and then permeabilized in xylenes (90% in ethanol) for 1h, after which embryos were washed six times in ethanol and three times in methanol. Embryos were then washed three times in PBT (PBS + 0.1% Tween-20) before post-fixation for 25min in fixative solution (225 μl 16% PFA, 500 μl PBT). Embryos were then washed several times in PBT for 40 min, followed by a wash in PBT/Hyb (50:50) at room temperature and a 30min-wash in pre-warmed Hyb at 56 ºC. Embryos were then incubated with probe solution at 56ºC overnight. The next day, embryos were washed in Hyb (three quick washes followed by three 30-min washes), then in Hyb/PBT (50:50), then in PBT several times for one hour before incubated for 30 min in blocking solution (Roche #11921673001; diluted 1:5 in PBT). Embryos were then incubated in blocking + primary antibodies diluted 1:500 (anti-DIG, Roche #11333089001; anti-FITC, ThermoFisher #A889) at 4 ºC overnight. The next day, embryos were washed in PBT (three quick washes followed by four 15-min washes), and then incubated at room temperature in blocking solution + secondary antibodies diluted 1:500 (AlexaFluor 488 and 555, ThermoFisher #A21206 and #A21436, respectively). After 2 hours, embryos were washed in PBT (three quick washes followed by four 15-min washes), mounted on Prolong Gold with DAPI (ThermoFisher, P36935) and left to curate overnight before imaging.

#### Immunofluorescence

embryos stored in methanol were washed in PBT (three quick washes followed by four 15-min washes), then in blocking solution for 30 min (Roche #11921673001; diluted 1:5 in PBT), before incubated overnight at 4 ºC in blocking solution + primary antibody diluted 1:500 (mouse anti-betagalactosidase, Promega #Z378). The next day, embryos were washed in PBT (three quick washes followed by four 15-min washes), and then incubated at room temperature in blocking solution + secondary antibody (donkey anti-mouse AlexaFluor 555, ThermoFisher #A31570). After 2 hours, embryos were washed in PBT (three quick washes followed by four 15-min washes), mounted on Prolong Gold with DAPI (ThermoFisher, P36935) and left to curate overnight before imaging.

#### Microscopy and data analysis

embryos were imaged using a confocal microscope Zeiss LSM 880 confocal. Images were processed using a combination of automated scripts with manual curation. For 3D plots showing signal intensity across three regions of the embryo (Fig. 1l, 2k, 3g, 5i), images were analyzed in ImageJ: a circular ROI of constant size was used to measure average intensity across the different regions (selected as shown in figures); number of lines/embryos analysed for each case are indicated in figure legends. For analyzing *E3N* mutant lines (Fuqua et al., 2021), individual nuclei were identified using the automated threshold algorithm on ImageJ and a watershed to split large ROIs; average intensities for each nucleus were measured. For analyzing *eveS2* mutant lines, we used ImageJ to perform Z-projections of max intensity, and a MATLAB (version R2018b; The MathWorks, Inc.) automated image analysis pipeline (named *Script-GAC*) was developed to capture expression signal along the AP axis on stage 5 embryos. For automated rotation, an ellipse was fitted on a masked embryo, and embryos were rotated based on the maximum Feret diameter. For quantification, a section with 30% of the height of the embryo was taken at a middle position and along the AP axis of each embryo. From this image section, the intensities from all the rows in the image matrix were averaged for each pixel position along the AP axis. The integration and analysis from each of these resultant AP embryo expression profiles were done in R (R Core Team, 2021). These expression profiles were smoothed with a Gaussian filter and then a linear interpolation was performed in order to have fixed samples number for the AP axis. Background removal and normalization were done based on the 10% and 50% quantile intensities, respectively, from the last 20% of the egg length. All embryos expression profiles per each genetic line were bootstrapped in order to see their reporter expression distribution along the AP axis. The bootstrapping was done using a confidence interval of 95% with 1000 replicates. For analyzing *twiPE* mutant lines, we used ImageJ to perform background subtraction from Z-projections of max intensity, rotate embryos to a vertical position and select a ROI at a defined position based on the intersection between 50% of the embryo long axis and the border of the *snail* RNA signal. We then used MorphoLibJ plugin in ImageJ to mask nuclei (volume higher than 3) and extracted intensities. For analyzing *rhoNEE* mutant lines, we used a custom code written in MATLAB (version R2018b; The MathWorks, Inc.), named *Script-MLP*; briefly, individual nuclei were segmented from the DAPI channel using a subroutine from the LivemRNA software package (Garcia et al., 2013). Stripes were then automatically identified by the following procedure: (1) bin nuclei by anterior-posterior (AP) coordinate; (2) within each bin, calculate a smoothed fluorescence profile along the dorsoventral (DV) coordinate based on the average fluorescence of each nucleus and its DV position; (3) identify peaks in the fluorescence profile for each bin; (4) align peaks across bins. Within each bin, nuclei falling within the AP coordinates for the half maximum height of a peak (on either side) were automatically considered to belong to the corresponding stripe. Manual curation was applied to fix any errors in stripe identification. Each stripe was then fitted lengthwise (AP axis) with a piecewise linear function through the middle, where for each line segment the stripe width was calculated perpendicular to the segment as the largest distance between the centers of nuclei “belonging” to the segment (i.e., nuclei with AP position falling between the AP coordinates of the two ends of the segment). Overall stripe width was calculated as the average of the widths of constituent segments. For analyzing *tinB* mutant lines, Z-projections of max intensity were generated using ImageJ and then embryos rotated and cropped to the minimum size in which the entire embryo still fitted the image. Composite images were then concatenated together and a montage was made using a scale factor of 1.0. Next, nuclear intensities were measured for each embryo in the montage. Channels were split, and in the DAPI channel the montage was smoothened twice. A threshold was manually set and applied, after which we used the “analyze particles” function based on a selection range of 100 to infinity. This threshold range was overlaid with the reporter channel, and nuclear intensities per embryo were retrieved using the ROI Manager.

### Motif prediction analysis of random sequences

Position weight matrices (PWMs) for *Drosophila melanogaster* and their logos were obtained from FlyFactorSurvey (Zhu et al., 2011). PWMs for specific stages of fly development were retrieved from (Li and Wunderlich, 2017). Motif search analysis was done using FIMO (Grant et al., 2011) and setting a threshold p-value of 0.001. The top 30% highest PWM-scores were selected to explore putative candidates for TFs binding sites.

### Information content

Information content for each of the TF motifs can be estimated using the Kullback-Leibler distance:

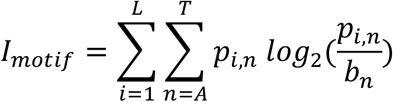

Where *p*_*i,n*_ is the probability of observing the nucleotide “n” at position “i” and *b*_*n*_ is the background frequency of nucleotide “n”. These values can be an indicative of how frequent a motif hit is expected by chance where 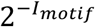 is an approximation of the probability for this event (Schneider et al., 1986). The empirical cumulative distribution plot for the information content scores was done in R.

## Supporting information

Script-MLP

Script-GAC

## DATA AND CODE AVAILABILITY

All fly lines and resources will be made available from corresponding author upon reasonable request. Automated scripts used can be found attached to this paper.

## ACKNOWLEDGEMENTS

We thank GenScript for gifting us the random DNA libraries, Hsiao-Yun Liu for making the protein for the Zelda antibody prep, Garth Ilsley for discussions about pattern quantifications, Claire Standley for title suggestions and Denis Krndija for critical feedback on the manuscript. We are grateful to other members of the Crocker lab for helpful suggestions and discussions during the course of the project, in particular Xueying Li. We also thank the ALMF imaging platform and Alessandra Reversi for some of the fly injections. R.G. and L.G. are supported by fellowships from the European Molecular Biology Laboratory Interdisciplinary Postdoc Programme (EIPOD) under Marie Skłodowska-Curie Actions COFUND (664726 and 847543, respectively). Research in the Crocker lab is supported by the European Molecular Biology Laboratory (EMBL).

## AUTHOR CONTRIBUTIONS

Conceptualization: RG, TF, JC. Investigation: RG, GAC, MRPA, NB, TF, LG, NM, KR, EK, TK, CAR, JC. Methodology: RG, GAC, NB, TF, LG, KR, JC. Formal analysis: RG, GAC, NB, TF, LG, NM, JC. Data curation: RG, NM, JC. Visualization: RG, TF, KR, NM, JC. Software: GAC, MLP, JC. Resources: CAR. Supervision: RG, TF, JC. Project administration: RG, JC. Funding acquisition: JC. Writing, original draft: RG, JC. Writing, review & editing: RG, GAC, NB, TF, LG, NM, MRPA, MLP, JC.

