## Supplementary material for "Enhancer architecture and chromatin accessibility constrain phenotypic space during development": Script-MLP: LICENSE.md

## License
<a rel="license" href="http://creativecommons.org/licenses/by-nc-sa/4.0/"><img alt="Creative Commons License" style="border-width:0" src="https://i.creativecommons.org/l/by-nc-sa/4.0/80x15.png" /></a><br /><span xmlns:dct="http://purl.org/dc/terms/" property="dct:title">TIFFStack</span> by <a xmlns:cc="http://creativecommons.org/ns#" href="http://dylan-muir.com" property="cc:attributionName" rel="cc:attributionURL">Dylan Muir</a> is licensed under a <a rel="license" href="http://creativecommons.org/licenses/by-nc-sa/4.0/">Creative Commons Attribution-NonCommercial-ShareAlike 4.0 International License</a>.<br />Based on a work at <a xmlns:dct="http://purl.org/dc/terms/" href="http://github.com/DylanMuir/TIFFStack" rel="dct:source">http://github.com/DylanMuir/TIFFStack</a>.

This work was published in [Frontiers in Neuroinformatics]: DR Muir and
BM Kampa. 2015. *[FocusStack and StimServer: A new open source MATLAB
toolchain for visual stimulation and analysis of two-photon calcium
neuronal imaging data](http://dx.doi.org/10.3389/fninf.2014.00085)*, **Frontiers in Neuroinformatics** 8 *85*. DOI: [dx.doi.org/10.3389/fninf.2014.00085](http://dx.doi.org/10.3389/fninf.2014.00085).
Please cite our publication in lieu of thanks, if you use this code.

  [Frontiers in Neuroinformatics]: http://www.frontiersin.org/neuroinformatics
