## Supplementary material for "Enhancer architecture and chromatin accessibility constrain phenotypic space during development": Script-MLP: mapped_tensor_repsum.c

```
/* mapped_tensor_repsum.c - MEX FUNCTION Replicate a vector, add elements of another vector
 *
 * Usage: [vfSummedRep] = mapped_tensor_repsum(vfSourceA, vfSourceB)
 *
 * 'vnSummedRep' will be a vector of length (numel(vfSourceA) * numel(vfSourceB)).
 * It will contain replicated copies of vector 'vfSourceA', where each copy
 * has been added with elements from 'vfSourceB' in turn.
 *
 * Author: Dylan Muir <>
 * Created: 17th July, 2012
 */

/* - Fix char16_t definition bug in OS X 10.9 */
#ifndef char16_t
	#include <stdint.h>
	typedef uint16_t char16_t;
#endif
#include "mex.h"

void mexFunction(int nlhs, mxArray *plhs[], int nrhs, const mxArray *prhs[]) {
   /* - Local variables */
   double   *vfSourceA, *vfSourceB, *vfDest;
   mwSize   uNumelA, uNumelB,
            uIndexA, uIndexB, uIndexDest;
   
   /* - Get arguments and sizes */
   uNumelA = mxGetM(prhs[0]) * mxGetN(prhs[0]);
   uNumelB = mxGetM(prhs[1]) * mxGetN(prhs[1]);
   
   vfSourceA = mxGetPr(prhs[0]);
   vfSourceB = mxGetPr(prhs[1]);
   
   /* - Allocate destination vector */
   if ((plhs[0] = mxCreateNumericMatrix(uNumelA * uNumelB, 1, mxDOUBLE_CLASS, mxREAL)) == NULL) {
      mexErrMsgIdAndTxt("MappedTensor:mapped_tensor_repsum:Memory", "Could not allocate memory.");
   }
   vfDest = mxGetPr(plhs[0]);
   
   
   /* -- Replicate vector A, add elements of vector B in turn */
      
   uIndexDest = 0;
   for (uIndexB = 0; uIndexB < uNumelB; uIndexB++) {
      for (uIndexA = 0; uIndexA < uNumelA; uIndexA++, uIndexDest++) {
         vfDest[uIndexDest] = vfSourceA[uIndexA] + vfSourceB[uIndexB];
      }
   }   
}

/* --- END of mapped_tensor_repsum.c --- */
```
