## Supplementary material for "Enhancer architecture and chromatin accessibility constrain phenotypic space during development": Script-MLP: tiffread31_header.m

```
function [TIF, HEADER, INFO] = tiffread31_header(file_name)

DEF_nPreallocateBlock = 1000;
DEF_HEADER = struct( 'SamplesPerPixel', [], 'index', [], 'ifd_pos', [], ...
                     'width', [], 'height', [], 'bits', [], 'StripOffsets', [], ...
                     'StripNumber', [], 'RowsPerStrip', [], 'StripByteCounts', [], ...
                     'cmap', [], 'colors', []);
DEF_INFO = struct('SamplesPerPixel', [], 'ByteOrder', [], 'Width', [], ...
                  'Height', [], 'BitsPerSample', [], ...
                  'RowsPerStrip', [], 'PlanarConfiguration', [], ...
                  'MaxSampleValue', [], 'MinSampleValue', []);
                     

%% --- Initialise variables and open TIFF file

TIF = struct('ByteOrder', 'ieee-le');  % byte order string
TIF.SamplesPerPixel = 1;
HEADER = DEF_HEADER;
HEADER.SamplesPerPixel = 1;
INFO = DEF_INFO;
INFO.SamplesPerPixel = 1;
TIF.PlanarConfiguration = 1;

% - Obtain the full file path
[status, file_attrib] = fileattrib(file_name);

if status == 0
   error('tiffread3:filenotfound', ['File "',file_name,'" not found.']);
end

file_name = file_attrib.Name;

% - Open file for reading
TIF.file = fopen(file_name,'r','l');

% - Obtain the short file name
[~, name, ext] = fileparts(file_name);
TIF.image_name = [name, ext];
TIF.file_name = file_name;


%% --- Read byte order: II = little endian, MM = big endian

bos = fread(TIF.file, 2, '*char');
if ( strcmp(bos', 'II') )
   TIF.ByteOrder = 'ieee-le';       % Little-endian format
   INFO.ByteOrder = 'little-endian';

elseif ( strcmp(bos','MM') )   
   TIF.ByteOrder = 'ieee-be';       % Big-endian format
   INFO.ByteOrder = 'big-endian';

else
   error('This is not a TIFF file (no MM or II).');
end


%% --- Read in a number which identifies TIFF format

TIF.tiff_id = fread(TIF.file,1,'uint16', TIF.ByteOrder);

if (TIF.tiff_id ~= 42) && (TIF.tiff_id ~= 43)
   error('This is not a TIFF file (missing 42 or 43).');
end

% - By default, read 4-byte pointers
TIF.strIFDNumEntriesSize = 'uint16';
TIF.strIFDClassSize = 'uint32';
TIF.nIFDTagBytes = 12;
TIF.nIFDClassBytes = 2;
TIF.strTagSizeClass = 'uint32';
TIF.nInlineBytes = 4;

% - Handle a BigTIFF file
if (TIF.tiff_id == 43)
   TIF.offset_size = fread(TIF.file, 1, 'uint16', TIF.ByteOrder);
   test_val = fread(TIF.file, 1, 'uint16', TIF.ByteOrder);
   
   % - Test check value
   if (test_val ~= 0)
      error('This is not a valid BigTIFF file (invalid test value).');
   end
   
   % - Get IFD pointer data class
   switch TIF.offset_size
      case 8
         TIF.strIFDNumEntriesSize = 'uint64';
         TIF.strIFDClassSize = 'uint64';
         TIF.nIFDClassBytes = 8;
         
      case 16
         TIF.strIFDNumEntriesSize = 'uint64';
         TIF.strIFDClassSize = '2*uint64';
         TIF.nIFDClassBytes = 16;
         
      otherwise
         error('Unknown IFD pointer size for BigTIFF file.');
   end
   
   TIF.nIFDTagBytes = 20;
   TIF.strTagSizeClass = 'uint64';
   TIF.nInlineBytes = 8;
end


%% --- Maps to accelerate types conversions in readIFDentry function

nbBytesMap = [ ...
   1, ... byte
   1, ... ascii string
   2, ... word
   4, ... dword/uword
   8, ... rational
   1, ... signed byte
   1, ... undefined
   2, ... signed short
   4, ... signed long
   8, ... long rational
   4, ... ???
   8, ... ???
   4, ... TIFF_IFD
   8, ... Long8
   8, ... SLong8
   8, ... Unsigned IFD offet 8 bytes
   ];

matlabTypeMap = { ...
   'uint8',   ... byte
   '*char',   ... ascii string
   'uint16',  ... word
   'uint32',  ... dword/uword
   'uint32',  ... rational
   'int8',    ... signed byte
   'uchar',   ... undefined
   'int16',   ... signed short
   'int32',   ... signed long
   'int32',   ... long rational
   'float32', ... ???
   'float64', ... ???
   'uint32',  ... TIFF_IFD
   'uint64',  ... Long8
   'int64',   ... SLong8
   'uint64',  ... Unsigned IFD offet 8 bytes
   };

%% --- Read the image file directories (IFDs)

% - Subset of tags to be parsed
parsed_tags = [256, 257, 258, 259, 262, 273, 277, 278, 279, 284, 317, 320, 339];

% - Read file offset of next IFD
ifd_pos = fread(TIF.file, 1, TIF.strIFDClassSize, TIF.ByteOrder);
img_indx = 0;

% - Attempt to preallocate
nPreallocateBlock = DEF_nPreallocateBlock;
HEADER(nPreallocateBlock) = DEF_HEADER;
INFO(nPreallocateBlock) = DEF_INFO;

% - Loop until the last IFD
while (ifd_pos ~= 0)

   img_indx = img_indx + 1;
   
   % - Extend arrays
   if (img_indx > nPreallocateBlock)
      nPreallocateBlock = nPreallocateBlock + DEF_nPreallocateBlock;
      HEADER(nPreallocateBlock) = DEF_HEADER;
      INFO(nPreallocateBlock) = DEF_INFO;
   end

   % - Store the IFD (frame) index and the IFD file offset
   HEADER(img_indx).index = img_indx; 
   HEADER(img_indx).ifd_pos = ifd_pos;

   % - Move in the file to the next IFD
   file_seek(ifd_pos);

   % - Read in the number of IFD entries
   num_entries = fread(TIF.file,1, TIF.strIFDNumEntriesSize, TIF.ByteOrder);

   % - Store current position
   entry_pos = ifd_pos + TIF.nIFDClassBytes;

   % - Read the next IFD file offset
   file_seek(ifd_pos + TIF.nIFDTagBytes * num_entries + TIF.nIFDClassBytes);
   ifd_pos = fread(TIF.file, 1, TIF.strIFDClassSize, TIF.ByteOrder);

   % - Read all the IFD entries
   for (inx = 1:num_entries)

      % - Move to next IFD entry in the file
      file_seek(entry_pos + TIF.nIFDTagBytes * (inx - 1));

      % - Read entry
      [entry_tag, entry_val, entry_cnt] = readIFDentry(TIF.strTagSizeClass, TIF.nInlineBytes);

      % - Not all valid tiff tags have been included, but tags can easily
      % be added to this code. See the official list of tags:
      % http://partners.adobe.com/asn/developer/pdfs/tn/TIFF6.pdf
      switch (entry_tag)

         % - Image width = number of columns
         case 256
            HEADER(img_indx).width = entry_val;
            INFO(img_indx).Width = entry_val;

         % - Image height = number of rows
         case 257
            HEADER(img_indx).height = entry_val;
            TIF.ImageLength = entry_val;
            INFO(img_indx).Height = entry_val;

         % - Bits per sample
         case 258
            TIF.BitsPerSample = entry_val;
            TIF.BytesPerSample = TIF.BitsPerSample / 8;
            HEADER(img_indx).bits = TIF.BitsPerSample(1);
            INFO(img_indx).BitsPerSample = entry_val;
         
         % - Compression type
         case 259         
            if entry_val ~= 1
               error('TIFFStack:Compression', ...
                   ['Compression format ', num2str(entry_val),' not supported.']);
            end

         % - Photometric interpretation
         case 262         
            TIF.PhotometricInterpretation = entry_val;
            if TIF.PhotometricInterpretation == 3
               warning('tiffread:LookUp', 'Ignoring TIFF look-up table.');
            end

         % - Strip offset
         case 273         
            HEADER(img_indx).StripOffsets = entry_val;
            HEADER(img_indx).StripNumber = entry_cnt;

         % - Samples per pixel
         case 277
            TIF.SamplesPerPixel = entry_val;
            INFO(img_indx).SamplesPerPixel = entry_val;

         % - Rows per strip
         case 278
            INFO(img_indx).RowsPerStrip = entry_val;
            HEADER(img_indx).RowsPerStrip = entry_val;

         % - Strip byte counts - number of bytes in each strip after any compression
         case 279         
            HEADER(img_indx).StripByteCounts= entry_val;

         % - Planar configuration describe the order of RGB
         case 284
            TIF.PlanarConfiguration = entry_val;

         % - Predictor for compression
         case 317
            if entry_val ~= 1
                error('Unsuported predictor value.')
            end

         % - Color map
         case 320
            HEADER(img_indx).cmap = entry_val;
            HEADER(img_indx).colors = entry_cnt / 3;

         % - Sample format
         case 339
            TIF.SampleFormat = entry_val;
      end
   end

   % - Total number of bytes per image plane
   if TIF.PlanarConfiguration == 1
      TIF.BytesPerPlane = TIF.SamplesPerPixel * HEADER(img_indx).width * HEADER(img_indx).height * TIF.BytesPerSample;
   else
      TIF.BytesPerPlane = HEADER(img_indx).width * HEADER(img_indx).height * TIF.BytesPerSample;
   end
end

% - Determine the data storage format
SampleFormat = 1;
if isfield(TIF, 'SampleFormat')
   SampleFormat = TIF.SampleFormat(1);
end

switch (SampleFormat)
   case 1   % - Unsigned integer
      TIF.classname = sprintf('uint%i', TIF.BitsPerSample(1));
      [INFO.MaxSampleValue] = deal(2.^TIF.BitsPerSample - 1);
      [INFO.MinSampleValue] = deal(0);

   case 2   % - Signed integer
      TIF.classname = sprintf('int%i', TIF.BitsPerSample(1));
      [INFO.MaxSampleValue] = deal(2.^(TIF.BitsPerSample-1) - 1);
      [INFO.MinSampleValue] = deal(-2.^(TIF.BitsPerSample-1));
   
   case 3   % - Real values
      if (TIF.BitsPerSample(1) == 32)
         % - Single-precision real value
         TIF.classname = 'single';
         [INFO.MaxSampleValue] = realmax('single');
         [INFO.MinSampleValue] = -realmax('single');

      else
         % - Double-precision real value
         TIF.classname = 'double';
         [INFO.MaxSampleValue] = realmax('double');
         [INFO.MinSampleValue] = -realmax('double');
      end
      
   otherwise
      % - Unknown / unsupported storage format
      error('TIFFStack:Format', '*** TIFFStack: Error: Unsuported TIFF sample format [%i].', SampleFormat);
end

% - Trim pre-allocated blocks
HEADER(img_indx+1:end) = [];
INFO(img_indx+1:end) = [];

% - Try to consolidate the TIFF strips if possible, to minimise disk reads
consolidate_strips(TIF.BytesPerPlane);

% --- END of tiffread31_header FUNCTION ---

%% --- Utility functions

   function file_seek(fpos)
      status = fseek(TIF.file, fpos, -1);
      if status == -1
         error('Invalid file offset (invalid fseek)');
      end
   end

   function consolidate_strips(BytesPerPlane)

      for (nFrame = 1:numel(HEADER))

         % - Try to consolidate the strips into a single one to speed-up reading
         nBytes = HEADER(nFrame).StripByteCounts(1);

         if (nBytes < BytesPerPlane)

            idx = 1;
            % - Accumulate continguous strips that contribute to the plane
            while (HEADER(nFrame).StripOffsets(1) + nBytes == HEADER(nFrame).StripOffsets(idx+1))
               idx = idx + 1;
               nBytes = nBytes + HEADER(nFrame).StripByteCounts(idx);
               if ( nBytes >= BytesPerPlane ); break; end
            end

            % - Consolidate the strips
            if (nBytes <= BytesPerPlane(1)) && (idx > 1)
               HEADER(nFrame).StripByteCounts = [nBytes; HEADER(nFrame).StripByteCounts(idx+1:HEADER(nFrame).StripNumber)];
               HEADER(nFrame).StripOffsets = HEADER(nFrame).StripOffsets([1 , idx+1:HEADER(nFrame).StripNumber]);
               HEADER(nFrame).StripNumber = 1 + HEADER(nFrame).StripNumber - idx;
            end
         end
      end
   end


%%  ---- Function to read an IFD entry

   function [entry_tag, entry_val, entry_cnt] = readIFDentry(strTagSizeClass, nInlineBytes)

      % - Read the IFD entry descriptor
      buffer = fread(TIF.file, 2, 'uint16', TIF.ByteOrder);
      entry_tag = buffer(1);
      tiffType = buffer(2);

      % - Skip tags that are not used afterwards
      if (~any(entry_tag == parsed_tags))  % Faster than `ismember` for this usage
          entry_val = [];
          entry_cnt = [];
          return;
      end

      % - Read the IFD entry
      entry_cnt = fread(TIF.file, 1, strTagSizeClass, TIF.ByteOrder);

      % - Map this IFD entry type to matlab storage classes
      try
         nbBytes = nbBytesMap(tiffType);
         matlabType = matlabTypeMap{tiffType};
         
      catch ME
         if strcmp(ME.identifier, 'MATLAB:badsubscript')
            error('TIFFSTack:Unsupported', 'TIFFStack/tiffread31_header/readIFDentry: tiff tag type %i not supported.', tiffType);
         else
            rethrow(ME);
         end
      end

      % - Check for whether the data for this tag is inline or not
      if (nbBytes * entry_cnt > nInlineBytes)
         % - The next field contains an offset
         fpos = fread(TIF.file, 1, 'uint32', TIF.ByteOrder);
         file_seek(fpos);
      end

      % - Handle TIFF 'rational' type
      if (tiffType == 5)
         val = fread(TIF.file, 2 * entry_cnt, matlabType, TIF.ByteOrder);
         entry_val = val(1:2:end) ./ val(2:2:end);

      else
         % - Read the data for this entry
         entry_val = fread(TIF.file, entry_cnt, matlabType, TIF.ByteOrder)';
      end

   end

end
```
