## Supplementary material for "Enhancer architecture and chromatin accessibility constrain phenotypic space during development": Script-MLP: tiffread31_readimage.m

```
function [data, stack] = tiffread31_readimage(TIF, HEADER, vnFrames)

opt.ReadUnknownTags = true;
opt.ConsolidateStrips = true;
opt.SimilarImages = false;
opt.DistributeMetaData = true;

% - Preallocate data block
stack = [];

for (nFrame = numel(vnFrames):-1:1) % Go backwards to pre-allocate
   
   nFrameID = vnFrames(nFrame);
   
   % Read pixel data
   
   if isfield(TIF, 'MM_stack')
      
      sel = ( vnFrames <= TIF.MM_stackCnt );
      vnFrames = vnFrames(sel);
      
      %this loop reads metamorph stacks:
      for ii = 1:numel(vnFrames)
         
         TIF.StripCnt = 1;
         
         %read the image data
         cFrame = read_pixels(TIF.BytesPerPlane * (ii-1), HEADER(vnFrames(ii)));
         data(:, :, nFrame, :) = cat(4, cFrame{:});
         
         [ IMG.MM_stack, IMG.MM_wavelength, IMG.MM_private2 ] = splitMetamorph(ii);
         
         IMG.index = ii;
         stack = cat(2, stack, IMG);
         
      end
      
      break;
      
   else
      
      if opt.SimilarImages && ~isempty(stack)
         if HEADER(nFrameID).width ~= stack(1).width || HEADER(nFrameID).height ~= stack(1).height
            % setting read_img=0 will skip dissimilar images:
            % comment-out the line below to allow dissimilar stacks
            fprintf('Tiffread skipped %ix%i image\n', HEADER(nFrameID).width, HEADER(nFrameID).height);
            continue;
         end
      end
      
      TIF.StripCnt = 1;
      
      %read image data
      cFrame = read_pixels(zeros(size(TIF.BytesPerPlane)), HEADER(nFrameID));
      data(:, :, nFrame, :) = cat(4, cFrame{:});
      
%       try
%          stack = cat(2, stack, IMG);
%       catch
%          try
%             stack = rmfield(stack, 'meta_data');
%             stack = rmfield(stack, 'meta_data_byte_counts');
%             stack = cat(2, stack, IMG);
%             fprintf('Tiffread had to discard some meta_data\n');
%          catch
%             fprintf('Tiffread could not load dissimilar image %i\n', img_indx);
%          end
%       end
      
   end
end


% remove the cell structure if there is always only one channel

flat = 1;
for ii = 1:length(stack)
   if length(stack(ii).data) ~= 1
      flat = 0;
      break;
   end
end

if flat
   for ii = length(stack):-1:1
      stack(ii).data = stack(ii).data{1};
   end
end


% distribute Andor IMG to all planes.

if opt.DistributeMetaData  &&  exist('ANDOR', 'var')
   
   fieldNames = fieldnames(ANDOR);
   for ii = length(stack):-1:1
      for jj = 1:size(fieldNames,1)
         stack(ii).(fieldNames{jj})=ANDOR.(fieldNames{jj});
      end
      stack(ii).planeNumber = ii;
   end
   % set nFrames if it doesn't exist
   if ~ isfield(stack,'nFrames')
      nFrames = length(stack);
      stack = setfield(stack, {1}, 'nFrames', nFrames);
   end
   
end

% distribute the MetaMorph info

if opt.DistributeMetaData
   
   if isfield(TIF, 'MM_stack') && isfield(IMG, 'info') && ~isempty(IMG.info)
      
      IMG.info = regexprep(IMG.info, '\r\n|\0', '\n');
      lines = textscan(IMG.info, '%s', 'Delimiter', '\n');
      mmi = parseMetamorphInfo(lines{1}, TIF.MM_stackCnt);
      for ii = length(stack):-1:1
         stack(ii).MM = mmi{stack(ii).index};
      end
      
   end
   
   % duplicate the LSM info
   if exist('LSM_info', 'var')
      for ii = length(stack):-1:1
         stack(ii).lsm = TIF.LSM_info;
      end
   end
   
end

return

%% Sub Function

   function file_seek(fpos)
      status = fseek(TIF.file, fpos, -1);
      if status == -1
         error('Invalid file offset (invalid fseek)');
      end
   end


   function pixels = read_pixels(offsets, FRAME)
      n_pix = FRAME.width * FRAME.height;
      bytes = read_plane(offsets(1), 1, TIF.SamplesPerPixel*n_pix, FRAME);
      pixels = cell(TIF.SamplesPerPixel, 1);
      if TIF.PlanarConfiguration == 2
         for c = 1:TIF.SamplesPerPixel
            pixels{c} = reshape(bytes((1+n_pix*(c-1)):(n_pix*c)), FRAME.width, FRAME.height)';
         end
      else
         spp = TIF.SamplesPerPixel;
         for c = 1:TIF.SamplesPerPixel
            pixels{c} = reshape(bytes(c:spp:c+spp*n_pix-spp), FRAME.width, FRAME.height)';
         end
      end
   end


   function bytes = read_plane(offset, plane_nb, bytes_nb, FRAME)
      
      %return an empty array if the sample format has zero bits
      if ( TIF.BitsPerSample(plane_nb) == 0 )
         bytes=[];
         return;
      end
      
      % fprintf('reading plane %i size %i %i\n', plane_nb, width, height);
            
      % Preallocate memory:
      try
         bytes = zeros(bytes_nb, 1, TIF.classname);
      catch
         %compatibility with older matlab versions:
         eval(['plane = ', TIF.classname, '(zeros(bytes_nb, 1));']);
      end
      
      % consolidate all strips:
      
      
      pos = 0;
      while 1
         
         strip = read_next_strip(offset, plane_nb, FRAME);
         
         if isempty(strip)
            break
         end
         
         bytes(1+pos:pos+length(strip)) = strip;
         pos = pos + length(strip);
         
      end
      
      if pos < bytes_nb
         warning('tiffread: %s','found fewer bytes than needed');
      end
   end

   function strip = read_next_strip(offset, plane_nb, FRAME)
      
      if ( TIF.StripCnt > FRAME.StripNumber )
         strip = [];
         return;
      end
      
      %fprintf('reading strip at position %i\n',TIF.StripOffsets(TIF.StripCnt) + offset);
      StripLength = FRAME.StripByteCounts(TIF.StripCnt) ./ TIF.BytesPerSample(plane_nb);
      
      %fprintf( 'reading strip %i\n', TIF.StripCnt);
      file_seek(FRAME.StripOffsets(TIF.StripCnt) + offset);
      
      strip = fread(TIF.file, StripLength, TIF.classname, TIF.ByteOrder);
      
      if any( length(strip) ~= StripLength )
         error('End of file reached unexpectedly.');
      end
      
      TIF.StripCnt = TIF.StripCnt + 1;
      
   end

%% =============distribute the metamorph infos to each frame:

   function [MMstack, MMwavelength, MMprivate2] = splitMetamorph(imgCnt)
      
      MMstack = [];
      MMwavelength = [];
      MMprivate2 = [];
      
      if TIF.MM_stackCnt == 1
         return;
      end
      
      left  = imgCnt - 1;
      
      if isfield( TIF, 'MM_stack' )
         S = length(TIF.MM_stack) / TIF.MM_stackCnt;
         MMstack = TIF.MM_stack(S*left+1:S*left+S);
      end
      
      if isfield( TIF, 'MM_wavelength' )
         S = length(TIF.MM_wavelength) / TIF.MM_stackCnt;
         MMwavelength = TIF.MM_wavelength(S*left+1:S*left+S);
      end
      
      if isfield( TIF, 'MM_private2' )
         S = length(TIF.MM_private2) / TIF.MM_stackCnt;
         MMprivate2 = TIF.MM_private2(S*left+1:S*left+S);
      end
      
   end


%% %%  Parse the Metamorph camera info tag into respective fields
% EVBR 2/7/2005, FJN Dec. 2007

   function mmi = parseMetamorphInfoLine(line, mmi)
      
      [tok, val] = strtok(line, ':');
      
      tok = regexprep(tok, ' ', '');
      val = strtrim(val(2:length(val)));
      
      if isempty(val)
         return;
      end
      
      %fprintf( '"%s" : "%s"\n', tok, val);
      
      if strcmp(tok, 'Exposure')
         [v, c, e, pos] = sscanf(val, '%i'); %#ok<ASGLU>
         unit = val(pos:length(val));
         %return the exposure in milli-seconds
         switch( unit )
            case 'ms'
               mmi.Exposure = v;
            case 's'
               mmi.Exposure = v * 1000;
            otherwise
               warning('tiffread:MetaMorphExposure', 'Exposure unit not recognized');
               mmi.Exposure = v;
         end
      else
         switch tok
            case 'Binning'
               % Binning: 1 x 1 -> [1 1]
               mmi.Binning = sscanf(val, '%d x %d')';
            case 'Region'
               mmi.Region = sscanf(val, '%d x %d, offset at (%d, %d)')';
            otherwise
               try
                  if strcmp(val, 'Off')
                     mmi.(tok) = 0;
                     %eval(['mm.',tok,'=0;']);
                  elseif strcmp(val, 'On')
                     mmi.(tok) = 1;
                     %eval(['mm.',tok,'=1;']);
                  elseif isstrprop(val,'digit')
                     mmi.(tok) = val;
                     %eval(['mm.',tok,'=str2num(val)'';']);
                  else
                     mmi.(tok) = val;
                     %eval(['mm.',tok,'=val;']);
                  end
               catch
                  warning('tiffread:MetaMorph', ['Invalid token "' tok '"']);
               end
         end
      end
   end

   function res = parseMetamorphInfo(lines, cnt)
      chk = length(lines) / cnt;
      res = cell(cnt, 1);
      length(lines)
      for n = 1:cnt
         mmi = [];
         for k = 1:chk
            mmi = parseMetamorphInfoLine(lines{chk*(n-1)+k}, mmi);
         end
         res{n} = mmi;
      end
   end

end
```
