## Supplementary material for "Enhancer architecture and chromatin accessibility constrain phenotypic space during development": Script-MLP: tiffread31_readtags.m

```
function [sTags] = tiffread31_readtags(TIF, HEADER, vnFrame)

%% set defaults values:

opt.ReadUnknownTags = true;
opt.ConsolidateStrips = true;
opt.SimilarImages = false;
opt.DistributeMetaData = true;

%% Maps to accelerate types conversions in readIFDentry function

nbBytesMap = [ ...
   1, ... byte
   1, ... ascii string
   2, ... word
   4, ... dword/uword
   8, ... rational
   1, ... signed byte
   1, ... undefined
   2, ... signed short
   4, ... signed long
   8, ... long rational
   4, ... ???
   8, ... ???
   4, ... TIFF_IFD
   8, ... Long8
   8, ... SLong8
   8, ... Unsigned IFD offet 8 bytes
   ];

matlabTypeMap = { ...
   'uint8',   ... byte
   '*char',   ... ascii string
   'uint16',  ... word
   'uint32',  ... dword/uword
   'uint32',  ... rational
   'int8',    ... signed byte
   'uchar',   ... undefined
   'int16',   ... signed short
   'int32',   ... signed long
   'int32',   ... long rational
   'float32', ... ???
   'float64', ... ???
   'uint32',  ... TIFF_IFD
   'uint64',  ... Long8
   'int64',   ... SLong8
   'uint64',  ... Unsigned IFD offet 8 bytes
   };

%% ---- read the image file directories (IFDs)

for (img_indx = 1:numel(vnFrame))
   
   % move in the file to the next IFD
   file_seek(HEADER(vnFrame(img_indx)).ifd_pos);
   sTags(img_indx).IFDPosition = HEADER(vnFrame(img_indx)).ifd_pos;
   
   %read in the number of IFD entries
   num_entries = fread(TIF.file,1,TIF.strIFDNumEntriesSize, TIF.ByteOrder);
   %fprintf('num_entries = %i\n', num_entries);
   
   % store current position:
   entry_pos = HEADER(vnFrame(img_indx)).ifd_pos+TIF.nIFDClassBytes;
   
   % read the next IFD address:
   file_seek(HEADER(vnFrame(img_indx)).ifd_pos+TIF.nIFDTagBytes*num_entries+TIF.nIFDClassBytes);
   
   %fprintf('reading IFD %i at position %i\n', img_indx, ifd_pos);
   
   % read all the IFD entries
   for inx = 1:num_entries
      
      % move to next IFD entry in the file
      file_seek(entry_pos+TIF.nIFDTagBytes*(inx-1));
      
      % read entry
      [entry_tag, entry_val, entry_cnt] = readIFDentry(TIF.strTagSizeClass, TIF.nInlineBytes);
      
      %fprintf('found entry with tag %i\n', entry_tag);
      
      % not all valid tiff tags have been included, but tags can easily be added to this code
      % See the official list of tags:
      % http://partners.adobe.com/asn/developer/pdfs/tn/TIFF6.pdf
      switch entry_tag
         case 256         % image width = number of column
            sTags(img_indx).Width = entry_val;
         case 257         % image height = number of row
            sTags(img_indx).Height = entry_val;
         case 258
            sTags(img_indx).BitsPerSample = entry_val;
            %fprintf('BitsPerSample %i %i %i\n', entry_val);
         case 259         % compression
            sTags(img_indx).Compression = entry_val;
         case 262         % photometric interpretation
            sTags(img_indx).PhotometricInterpretation = entry_val;
         case 269
            sTags(img_indx).DocumentName = entry_val;
         case 270         % general comments:
            sTags(img_indx).ImageDescription = entry_val;
         case 271
            sTags(img_indx).make = entry_val;
         case 273         % strip offset
            sTags(img_indx).StripOffsets = entry_val;
            sTags(img_indx).StripNumber = entry_cnt;
            %fprintf('StripNumber = %i, size(StripOffsets) = %i %i\n', TIF.StripNumber, size(TIF.StripOffsets));
         case 274
            % orientation is read, but the matrix is not rotated
            if ( 1 < entry_val ) && ( entry_val < 9 )
               sTags(img_indx).OrientationID = entry_val;
               keys = {'TopLeft', 'TopRight', 'BottomRight', 'BottomLeft', 'LeftTop', 'RightTop', 'RightBottom', 'LeftBottom'};
               sTags(img_indx).Orientation = keys{entry_val};
            end
         case 277
            sTags(img_indx).SamplesPerPixel = entry_val;
            %fprintf('Color image: sample_per_pixel=%i\n',  TIF.SamplesPerPixel);
         case 278
            sTags(img_indx).RowsPerStrip   = entry_val;
         case 279         % strip byte counts - number of bytes in each strip after any compressio
            sTags(img_indx).StripByteCounts= entry_val;
         case 282
            sTags(img_indx).XResolution   = entry_val;
         case 283
            sTags(img_indx).YResolution   = entry_val;
         case 284         % planar configuration describe the order of RGB
            sTags(img_indx).PlanarConfiguration = entry_val;
         case 296
            sTags(img_indx).ResolutionUnit= entry_val;
         case 305
            sTags(img_indx).Software = entry_val;
         case 306
            sTags(img_indx).FileModDate       = entry_val;
         case 315
            sTags(img_indx).Artist = entry_val;
         case 317        %predictor for compression
            sTags(img_indx).CompressionPredictorVal = entry_val;
         case 320         % color map
            sTags(img_indx).Colormap           = entry_val;
            sTags(img_indx).NumColors         = entry_cnt/3;
         case 339
            sTags(img_indx).SampleFormat   = entry_val;
            
            % ANDOR tags
         case 4864
            if ~exist('ANDOR', 'var')
               %the existence of the variable indicates that we are
               %handling a Andor generated file
               sTags(img_indx).ANDOR = struct([]);
            end
         case 4869       %ANDOR tag: temperature in Celsius when stabilized
            if ~(entry_val == -999)
               sTags(img_indx).ANDOR(1).temperature = entry_val;
            end
         case 4876       %exposure time in seconds
            sTags(img_indx).ANDOR(1).exposureTime   = entry_val;
         case 4878
            sTags(img_indx).ANDOR(1).kineticCycleTime = entry_val;
         case 4879       %number of accumulations
            sTags(img_indx).ANDOR(1).nAccumulations = entry_val;
         case 4881
            sTags(img_indx).ANDOR(1).acquisitionCycleTime = entry_val;
         case 4882       %Readout time in seconds, 1/readoutrate
            sTags(img_indx).ANDOR(1).readoutTime = entry_val;
         case 4884
            if (entry_val == 9)
               sTags(img_indx).ANDOR(1).isPhotonCounting = 1;
            else
               sTags(img_indx).ANDOR(1).isPhotonCounting = 0;
            end
         case 4885         %EM DAC level
            sTags(img_indx).ANDOR(1).emDacLevel = entry_val;
         case 4890
            sTags(img_indx).ANDOR(1).nFrames = entry_val;
         case 4896
            sTags(img_indx).ANDOR(1).isFlippedHorizontally = entry_val;
         case 4897
            sTags(img_indx).ANDOR(1).isFlippedVertically = entry_val;
         case 4898
            sTags(img_indx).ANDOR(1).isRotatedClockwise = entry_val;
         case 4899
            sTags(img_indx).ANDOR(1).isRotatedAnticlockwise = entry_val;
         case 4904
            sTags(img_indx).ANDOR(1).verticalClockVoltageAmplitude = entry_val;
         case 4905
            sTags(img_indx).ANDOR(1).verticalShiftSpeed = entry_val;
         case 4907
            sTags(img_indx).ANDOR(1).preAmpSetting = entry_val;
         case 4908         %Camera Serial Number
            sTags(img_indx).ANDOR(1).serialNumber = entry_val;
         case 4911       %Actual camera temperature when not equal to -999
            if ~(entry_val == -999)
               sTags(img_indx).ANDOR(1).unstabilizedTemperature = entry_val;
            end
         case 4912
            sTags(img_indx).ANDOR(1).isBaselineClamped = entry_val;
         case 4913
            sTags(img_indx).ANDOR(1).nPrescans = entry_val;
         case 4914
            sTags(img_indx).ANDOR(1).model = entry_val;
         case 4915
            sTags(img_indx).ANDOR(1).chipXSize = entry_val;
         case 4916
            sTags(img_indx).ANDOR(1).chipYSize  = entry_val;
         case 4944
            sTags(img_indx).ANDOR(1).baselineOffset = entry_val;
            
         case 33550       % GeoTIFF
            sTags(img_indx).ModelPixelScaleTag = entry_val;
         case 33628       % Metamorph specific data
            sTags(img_indx).MM_private1 = entry_val;
         case 33629       % this tag identify the image as a Metamorph stack!
            sTags(img_indx).MM_stack = entry_val;
            sTags(img_indx).MM_stackCnt = entry_cnt;
         case 33630       % Metamorph stack data: wavelength
            sTags(img_indx).MM_wavelength = entry_val;
         case 33631       % Metamorph stack data: gain/background?
            sTags(img_indx).MM_private2 = entry_val;
            
         case 33922       % GeoTIFF
            sTags(img_indx).ModelTiePointTag = entry_val;
         case 34412       % Zeiss LSM data
            sTags(img_indx).LSM_info = entry_val;
         case 34735       % GeoTIFF
            sTags(img_indx).GeoKeyDirTag = entry_val;
         case 34737       % GeoTIFF
            sTags(img_indx).GeoASCII = entry_val;
         case 42113       % GeoTIFF
            sTags(img_indx).GDAL_NODATA = entry_val;
            
         case 50838       % Adobe
            sTags(img_indx).meta_data_byte_counts = entry_val;
         case 50839       % Adobe
            sTags(img_indx).meta_data = entry_val;
            
         otherwise
            if opt.ReadUnknownTags
               sTags(img_indx).(['tag', num2str(entry_tag)])=entry_val;
               %eval(['IMG(img_indx+1).Tag',num2str(entry_tag),'=',entry_val,';']);
            else
               fprintf( 'Unknown TIFF entry with tag %i (cnt %i)\n', entry_tag, entry_cnt);
            end
      end

      % - Calculate the bounding box  if we have the required information
      if isfield(sTags, 'ModelPixelScaleTag') && isfield(sTags, 'ModelTiePointTag') && isfield(sTags, 'Height')&& isfield(sTags, 'Width'),
         sTags(img_indx).North=sTags(img_indx).ModelTiePointTag(5)-sTags(img_indx).ModelPixelScaleTag(2)*sTags(img_indx).ModelTiePointTag(2);
         sTags(img_indx).South=sTags(img_indx).North-sTags(img_indx).height*sTags(img_indx).ModelPixelScaleTag(2);
         sTags(img_indx).West=sTags(img_indx).ModelTiePointTag(4)+sTags(img_indx).ModelPixelScaleTag(1)*sTags(img_indx).ModelTiePointTag(1);
         sTags(img_indx).East=sTags(img_indx).West+sTags(img_indx).width*sTags(img_indx).ModelPixelScaleTag(1);
      end
   end
   
   
   % total number of bytes per image:
   if TIF.PlanarConfiguration == 1
      sTags(img_indx).BytesPerPlane = TIF.SamplesPerPixel * sTags(img_indx).Width * sTags(img_indx).Height * TIF.BytesPerSample;
   else
      sTags(img_indx).BytesPerPlane = sTags(img_indx).Width * sTags(img_indx).Height * TIF.BytesPerSample;
   end
end

% determine the type of data stored in the pixels:
SampleFormat = 1;
if isfield(TIF, 'SampleFormat')
   SampleFormat = TIF.SampleFormat(1);
end

switch( SampleFormat )
   case 1
      [sTags.MaxSampleValue] = deal(2.^TIF.BitsPerSample - 1);
      [sTags.MinSampleValue] = deal(0);

   case 2
      [sTags.MaxSampleValue] = deal(2.^(TIF.BitsPerSample-1) - 1);
      [sTags.MinSampleValue] = deal(-2.^(TIF.BitsPerSample-1));
   
   case 3
      if (TIF.BitsPerSample(1) == 32 )
         [sTags.MaxSampleValue] = realmax('single');
         [sTags.MinSampleValue] = -realmax('single');
      else
         [sTags.MaxSampleValue] = realmax('double');
         [sTags.MinSampleValue] = -realmax('double');
      end
      
   otherwise
      error('TIFFStack:Format', '*** TIFFStack: Error: Unsuported TIFF sample format [%i].', SampleFormat);
end

% - Try to consolidate the TIFF strips if possible

if opt.ConsolidateStrips
   consolidate_strips(TIF.BytesPerPlane);
end


%% Sub Functions

   function file_seek(fpos)
      status = fseek(TIF.file, fpos, -1);
      if status == -1
         error('Invalid file offset (invalid fseek)');
      end
   end

   function consolidate_strips(BytesPerPlane)
      
      for (nFrame = 1:numel(sTags))
         
         %Try to consolidate the strips into a single one to speed-up reading:
         nBytes = sTags(nFrame).StripByteCounts(1);
         
         if nBytes < BytesPerPlane
            
            idx = 1;
            % accumulate continguous strips that contribute to the plane
            while sTags(nFrame).StripOffsets(1) + nBytes == sTags(nFrame).StripOffsets(idx+1)
               idx = idx + 1;
               nBytes = nBytes + sTags(nFrame).StripByteCounts(idx);
               if ( nBytes >= BytesPerPlane ); break; end
            end
            
            %Consolidate the Strips
            if ( nBytes <= BytesPerPlane(1) ) && ( idx > 1 )
               %fprintf('Consolidating %i stripes out of %i', ConsolidateCnt, TIF.StripNumber);
               sTags(nFrame).StripByteCounts = [nBytes; sTags(nFrame).StripByteCounts(idx+1:sTags(nFrame).StripNumber ) ];
               sTags(nFrame).StripOffsets = sTags(nFrame).StripOffsets( [1 , idx+1:sTags(nFrame).StripNumber] );
               sTags(nFrame).StripNumber  = 1 + sTags(nFrame).StripNumber - idx;
            end
         end
      end
   end


%% ==================sub-functions that reads an IFD entry:===================

   function [entry_tag, entry_val, entry_cnt] = readIFDentry(strTagSizeClass, nInlineBytes)
      
      buffer = fread(TIF.file, 2, 'uint16', TIF.ByteOrder);
      entry_tag = buffer(1);
      tiffType = buffer(2);
      entry_cnt = fread(TIF.file, 1, strTagSizeClass, TIF.ByteOrder);
      
      try
         nbBytes = nbBytesMap(tiffType);
         matlabType = matlabTypeMap{tiffType};
      catch ME
         if strcmp(ME.identifier, 'MATLAB:badsubscript')
            error('tiff type %i not supported', tiffType);
         else
            rethrow(ME);
         end
      end
      
      if nbBytes * entry_cnt > nInlineBytes
         % next field contains an offset
         fpos = fread(TIF.file, 1, strTagSizeClass, TIF.ByteOrder);
         file_seek(fpos);
      end
      
      if entry_tag == 33629   % metamorph stack plane specifications
         entry_val = fread(TIF.file, 6 * entry_cnt, matlabType, TIF.ByteOrder);
         
      elseif entry_tag == 34412  % TIF_CZ_LSMINFO
         entry_val = readLSMinfo();
         
      elseif tiffType == 5  % TIFF 'rational' type
         val = fread(TIF.file, 2 * entry_cnt, matlabType, TIF.ByteOrder);
         entry_val = val(1:2:end) ./ val(2:2:end);
         
      else
         entry_val = fread(TIF.file, entry_cnt, matlabType, TIF.ByteOrder)';
      end
      
   end


%% ==============partial-parse of LSM info:

   function R = readLSMinfo()
      
      % Read part of the LSM info table version 2
      % this provides only very partial information, since the offset indicate that
      % additional data is stored in the file
      
      R.MagicNumber          = sprintf('0x%09X',fread(TIF.file, 1, 'uint32', TIF.ByteOrder));
      S.StructureSize        = fread(TIF.file, 1, 'uint32', TIF.ByteOrder);
      R.DimensionX           = fread(TIF.file, 1, 'uint32', TIF.ByteOrder);
      R.DimensionY           = fread(TIF.file, 1, 'uint32', TIF.ByteOrder);
      R.DimensionZ           = fread(TIF.file, 1, 'uint32', TIF.ByteOrder);
      R.DimensionChannels    = fread(TIF.file, 1, 'uint32', TIF.ByteOrder);
      R.DimensionTime        = fread(TIF.file, 1, 'uint32', TIF.ByteOrder);
      R.IntensityDataType    = fread(TIF.file, 1, 'uint32', TIF.ByteOrder);
      R.ThumbnailX           = fread(TIF.file, 1, 'uint32', TIF.ByteOrder);
      R.ThumbnailY           = fread(TIF.file, 1, 'uint32', TIF.ByteOrder);
      R.VoxelSizeX           = fread(TIF.file, 1, 'float64', TIF.ByteOrder);
      R.VoxelSizeY           = fread(TIF.file, 1, 'float64', TIF.ByteOrder);
      R.VoxelSizeZ           = fread(TIF.file, 1, 'float64', TIF.ByteOrder);
      R.OriginX              = fread(TIF.file, 1, 'float64', TIF.ByteOrder);
      R.OriginY              = fread(TIF.file, 1, 'float64', TIF.ByteOrder);
      R.OriginZ              = fread(TIF.file, 1, 'float64', TIF.ByteOrder);
      R.ScanType             = fread(TIF.file, 1, 'uint16', TIF.ByteOrder);
      R.SpectralScan         = fread(TIF.file, 1, 'uint16', TIF.ByteOrder);
      R.DataType             = fread(TIF.file, 1, 'uint32', TIF.ByteOrder);
      S.OffsetVectorOverlay  = fread(TIF.file, 1, 'uint32', TIF.ByteOrder);
      S.OffsetInputLut       = fread(TIF.file, 1, 'uint32', TIF.ByteOrder);
      S.OffsetOutputLut      = fread(TIF.file, 1, 'uint32', TIF.ByteOrder);
      S.OffsetChannelColors  = fread(TIF.file, 1, 'uint32', TIF.ByteOrder);
      R.TimeInterval         = fread(TIF.file, 1, 'float64', TIF.ByteOrder);
      S.OffsetChannelDataTypes = fread(TIF.file, 1, 'uint32', TIF.ByteOrder);
      S.OffsetScanInformatio = fread(TIF.file, 1, 'uint32', TIF.ByteOrder);
      S.OffsetKsData         = fread(TIF.file, 1, 'uint32', TIF.ByteOrder);
      S.OffsetTimeStamps     = fread(TIF.file, 1, 'uint32', TIF.ByteOrder);
      S.OffsetEventList      = fread(TIF.file, 1, 'uint32', TIF.ByteOrder);
      S.OffsetRoi            = fread(TIF.file, 1, 'uint32', TIF.ByteOrder);
      S.OffsetBleachRoi      = fread(TIF.file, 1, 'uint32', TIF.ByteOrder);
      S.OffsetNextRecording  = fread(TIF.file, 1, 'uint32', TIF.ByteOrder);
      
      % There is more information stored in this table, which is skipped here
      
      % read real acquisition times:
      if ( S.OffsetTimeStamps > 0 )
         
         status =  fseek(TIF.file, S.OffsetTimeStamps, -1);
         if status == -1
            warning('tiffread:TimeStamps', 'Could not locate LSM TimeStamps');
            return;
         end
         
         StructureSize          = fread(TIF.file, 1, 'int32', TIF.ByteOrder); %#ok<NASGU>
         NumberTimeStamps       = fread(TIF.file, 1, 'int32', TIF.ByteOrder);
         for i=1:NumberTimeStamps
            R.TimeStamp(i)      = fread(TIF.file, 1, 'float64', TIF.ByteOrder);
         end
         
         %calculate elapsed time from first acquisition:
         R.TimeOffset = R.TimeStamp - R.TimeStamp(1);
         
      end
      
      % anything else assigned to S is discarded
      
   end

end
```
