## Supplementary material for "Enhancer architecture and chromatin accessibility constrain phenotypic space during development": Script-MLP: TS_UnitTest.m

```
% TS_UnitTest - FUNCITON Unit test for TIFFStack
%
% Usage: TS_UnitTest
%
% This function performs a series of unit tests for referencing a TIFFStack
% object, in comparison with a matlab tensor. If all tests are passed, a
% simple message reporting success will be displayed. Any error indicates a
% failure in a unit test, which will be reported in detail. Any error
% should be considered a serious bug.

% Author: Dylan Muir <>
% Created: 17th Novemeber, 2015

function TS_UnitTest(strFilename)
   % - Make sure TIFFStack is on the path
   strTSDir = fileparts(which('TIFFStack'));
   
   if (isempty(strTSDir))
      error('TIFFStack:Path', '*** Error: ''TIFFStack'' is not available on the matlab path.');
   end
   
   strBaseDir = fileparts(strTSDir);
   addpath(strBaseDir);

   % - If we are being called from within 'private', then make a temp copy and run unit tests from there
   if (isequal(mfilename, 'TS_UnitTest'))
      % - Get a temporary directory and name
      strUTTestDir = fullfile(tempdir, tempname);
      mkdir(strUTTestDir);
      [~, strUTFilename] = fileparts(tempname);
      
      % - Copy the unit test file to the temporary directory
      copyfile(fullfile(strTSDir, 'private', 'TS_UnitTest.m'), fullfile(strUTTestDir, [strUTFilename '.m']));
      addpath(strUTTestDir);
      
      % - Call the unit test function
      fhUnitTest = str2func(strUTFilename);
      fhUnitTest();
      
      % - Clear up and return
      delete(fullfile(strUTTestDir, [strUTFilename '.m']));
      w = warning('off', 'MATLAB:RMDIR:RemovedFromPath');
      rmdir(strUTTestDir);
      warning(w);
      return;
   end
   
   % - Open a test TIFF file
   if (exist('strFilename', 'var') && ~isempty(strFilename))
      % - Run on user-supplied file
      TSUT_RunTestsOnFile(strFilename);
      
   else
      % - Find unit test file
      strFilename = fullfile(strTSDir, 'private', 'TS_UnitTestImage.tif');
      TSUT_RunTestsOnFile(strFilename);

      % - Find unit test bigtiff file
      strFilename = fullfile(strTSDir, 'private', 'TS_UnitTestImage_BigTIFF.tif');
      TSUT_RunTestsOnFile(strFilename);

      % - Find unit test ImageJ Big stack file
      % Currently not integrated into unit test
      % strFilename = fullfile(strTSDir, 'private', 'TS_UnitTestImage_ImageJ_BigStack_8frames.tif');
      % TSUT_RunTestsOnFile(strFilename);
   end
end

function TSUT_RunTestsOnFile(strFilename)
   
   fprintf('--- TS_UnitTest: Running on file [%s]...\n', strFilename);

   % - Try to make a TIFFStack object
   tsStack = TIFFStack(strFilename);
   
   % - Load test image using 'imread'
   sInfo = imfinfo(strFilename);
   nNumFrames = numel(sInfo);
   
   for (nFrame = nNumFrames:-1:1)
      tfFrame = permute(imread(strFilename, 'Index', nFrame, 'Info', sInfo), [1 2 4 3]);
      tfStack(:, :, nFrame, :) = tfFrame;
   end
   
   % - Check that test stack is big enough
   vnStackSize = size(tfStack);
   if (numel(vnStackSize) < 3)
      error('TIFFStack:Arguments', '*** TS_UnitTest: Stack provided for testing [%s] must have at least 3 dimensions.', strFilename);
   end
   
   if (any(vnStackSize(1:3) < [3 3 3]))
      error('TIFFStack:Arguments', '*** TS_UnitTest: Stack provided for testing [%s] must be at least [3x3x3].', strFilename);
   end      
   
   
   %% - Test straightforward referencing
   TSUT_TestReferencing(tsStack, tfStack, 'Raw stack');
   
   %% - Test permuted stack
   tsStack = permute(tsStack, [3 1 2 4]);
   tfStack = permute(tfStack, [3 1 2 4]);
   TSUT_TestReferencing(tsStack, tfStack, 'Simple permutation');
   tsStack = ipermute(tsStack, [3 1 2 4]);
   tfStack = ipermute(tfStack, [3 1 2 4]);   
      
   %% - Test inverted stack
   sTSInfo = getImageInfo(tsStack);
   fMaxSampleValue = sTSInfo(1).MaxSampleValue;
   fMinSampleValue = sTSInfo(1).MinSampleValue;
   
   tsStack.bInvert = true;
   tfStack = fMaxSampleValue - (tfStack - fMinSampleValue);
   TSUT_TestReferencing(tsStack, tfStack, 'Inverted stack');
   tsStack.bInvert = false;

   % - Re-extract stack
   for (nFrame = nNumFrames:-1:1)
      tfFrame = permute(imread(strFilename, 'Index', nFrame, 'Info', sInfo), [1 2 4 3]);
      tfStack(:, :, nFrame, :) = tfFrame;
   end
   
   %% - Test non-accelerated reading funcitons
   w = warning('off', 'TIFFStack:SlowAccess');
   tsStack = TIFFStack(strFilename, [], [], true);
   TSUT_TestReferencing(tsStack, tfStack, 'Non-accelerated TIFF reading');
   warning(w);

   %% - Test deinterleaved stack
   vnStackSize = size(tfStack);
   nNumFrames = size(tfStack, 3);

   % - Test that de-interleaving succeeded
   tsStack = TIFFStack(strFilename, [], [1 1 nNumFrames 1]);
   vnTestSize = [vnStackSize([1 2]) 1 1 nNumFrames 1 vnStackSize(4:end)];
   if (vnTestSize(end) == 1)
      vnTestSize = vnTestSize(1:(find(vnTestSize == 1, 1, 'last') - 1));
   end
   assert(isequal(size(tsStack), vnTestSize), ...
          'TIFFStack:UnitTestFailed', 'De-interleaving the stack was unsuccessful.');

   tsStack = TIFFStack(strFilename, [], [1 1 nNumFrames]);   
   vnTestSize = [vnStackSize([1 2]) 1 1 nNumFrames vnStackSize(4:end)];
   if (vnTestSize(end) == 1)
      vnTestSize = vnTestSize(1:(find(vnTestSize == 1, 1, 'last') - 1));
   end
   assert(isequal(size(tsStack), vnTestSize), ...
          'TIFFStack:UnitTestFailed', 'De-interleaving the stack was unsuccessful.');

   tsStack = TIFFStack(strFilename, [], [1 1]);
   vnTestSize = [vnStackSize([1 2]) 1 1 nNumFrames vnStackSize(4:end)];
   if (vnTestSize(end) == 1)
      vnTestSize = vnTestSize(1:(find(vnTestSize == 1, 1, 'last') - 1));
   end
   assert(isequal(size(tsStack), vnTestSize), ...
          'TIFFStack:UnitTestFailed', 'De-interleaving the stack was unsuccessful.');

   TSUT_assertFail('MATLAB:TIFFStack:expectedInteger', 'TIFFStack(strFilename, [], .5);');
       
   tfStack = reshape(tfStack, [size(tfStack, 1) size(tfStack, 2) 1 1 nNumFrames size(tfStack, 4)]);
   TSUT_TestReferencing(tsStack, tfStack, 'Deinterleaved stack');

   %% - Test saving / loading a stack
   strMatFile = tempname;
   save(strMatFile, 'tsStack');
   s = load(strMatFile);
   TSUT_TestReferencing(tsStack, s.tsStack, 'Serialised/Deserialised stack');
   
   %% - Test error conditions
   TSUT_assertFail('TIFFStack:Concatenation', '[tsStack tsStack];');
   TSUT_assertFail('TIFFStack:Concatenation', '[tsStack; tsStack];');
   TSUT_assertFail('TIFFStack:Concatenation', 'cat(1, tsStack, tsStack);');
   TSUT_assertFail('TIFFStack:InvalidReferencing', 'tsStack{1}');
   TSUT_assertFail('TIFFStack:InvalidReferencing', 'tsStack.diagnostic();');
   TSUT_assertFail('TIFFStack:DimensionMustBePositiveInteger', 'size(tsStack, 0);');
   TSUT_assertFail('TIFFStack:InvalidArgument', 'tsStack.bInvert = 2;');
   
   %% - Success if we reach here with no errors
   disp('--- TS_UnitTest: Unit tests for ''TIFFStack'' passed.');
end

function TSUT_TestReferencing(tsStack, tfStack, strTestName)
   % - Test stack sizes
   assert(isequal(size(tsStack), size(tfStack)), ...
          'TIFFStack:UnitTestFailed', 'The result of calling ''size'' was not equal between the two stacks.');
   assert(isequal(size(tsStack, 1), size(tfStack, 1)), ...
          'TIFFStack:UnitTestFailed', 'The result of calling ''size'' was not equal between the two stacks.');
   assert(isequal(size(tsStack, 2), size(tfStack, 2)), ...
          'TIFFStack:UnitTestFailed', 'The result of calling ''size'' was not equal between the two stacks.');
   assert(isequal(size(tsStack, 3), size(tfStack, 3)), ...
          'TIFFStack:UnitTestFailed', 'The result of calling ''size'' was not equal between the two stacks.');
   assert(isequal(size(tsStack, 4), size(tfStack, 4)), ...
          'TIFFStack:UnitTestFailed', 'The result of calling ''size'' was not equal between the two stacks.');

   % - Test overloaded functions
   TSUT_TestOverloads(tsStack, tfStack, strTestName);
       
   % - Test referencing entire stack
   TSUT_compareRef(':');
   TSUT_compareRef(':', ':');
   TSUT_compareRef(':', ':', ':');
   TSUT_compareRef(':', ':', ':', ':');
   TSUT_compareRef(':', ':', ':', ':', ':');
   TSUT_compareRef(':', ':', ':', ':', ':', ':');
   
   % - Test referencing subpixels
   TSUT_compareRef(1:2, ':');
   TSUT_compareRef(1:2, 1:2, ':');
   TSUT_compareRef(1:2, 9);
   TSUT_compareRef([1 3], 9);
   TSUT_compareRef(':', ':', 3);
   
   % - Test empty referencing
   TSUT_compareRef([]);
   TSUT_compareRef([], []);
   TSUT_compareRef([], [], []);
   TSUT_compareRef([], [], [], []);
   TSUT_compareRef([], [], [], [], []);
   TSUT_compareRef(1, 1, 1, [], 1);
   
   % - Test referencing using 'end'
   TSUT_compareEndRefs();
   
   % - Test referencing with unit indices
   TSUT_compareRef(1);
   TSUT_compareRef(1, 1);
   TSUT_compareRef(1, 1, 1);
   TSUT_compareRef(1, 1, 1, 1);
   TSUT_compareRef(1, 1, 1, 1, 1);
   TSUT_compareRef(1, 1, 1, 1, 1, 1);
   
   % - Test logical indexing
   TSUT_compareRef([true false true], [true true false], [false false true]);

   % - Test limit cases that should cause errors
   vnStackSize = size(tfStack);
   TSUT_testInvalidRef('TIFFStack:badsubscript', 0);
   TSUT_testInvalidRef('TIFFStack:badsubscript', 0.1);
   TSUT_testInvalidRef('TIFFStack:badsubscript', {1});
   TSUT_testInvalidRef('TIFFStack:badsubscript', complex(1, 0));
   TSUT_testInvalidRef('TIFFStack:badsubscript', 'a');
   TSUT_testInvalidRef('TIFFStack:badsubscript', prod(vnStackSize)+1);
   TSUT_testInvalidRef('TIFFStack:badsubscript', 1, prod(vnStackSize(2:end))+1);
   TSUT_testInvalidRef('TIFFStack:badsubscript', 1, vnStackSize(2)+1, 1);
   TSUT_testInvalidRef('TIFFStack:badsubscript', 1, 1, vnStackSize(3)+1, 1);
   TSUT_testInvalidRef('TIFFStack:badsubscript', {1});
   TSUT_testInvalidRef('TIFFStack:badsubscript', 1, 1, 1, 1, 1, 1, 1, 2);
   
   % - Function to compare results of referencing stack
   function TSUT_compareRef(varargin)
      try
         % - Reference stacks, check for equality
         s.type = '()';
         s.subs = varargin;
         tfTS = subsref(tsStack, s);
         tfTF = tfStack(varargin{:});
         assert(isequal(tfTS, tfTF), 'TIFFStack:UnitTestFailed:ResultsNotEqual', ...
            'The referencing result was not equal between TIFFStack and standard tensor, with subs %s.', TSUT_subs2str(varargin));
         
      catch mErr
         mUTErr = MException('TIFFStack:UnitTestFailed', 'Failure referencing stack with subs %s, during test [%s]', ...
            TSUT_subs2str(varargin), strTestName);
         mErr = mErr.addCause(mUTErr);
         rethrow(mErr);
      end
   end

   % - Function to test referencing using 'end'
   function TSUT_compareEndRefs
      try
         % - Reference stacks, check for equality
         ae(tsStack(end), ...
            tfStack(end));
         ae(tsStack(1:end, 1), ...
            tfStack(1:end, 1));
         ae(tsStack(1, 1:end), ...
            tfStack(1, 1:end));
         ae(tsStack(1, 1:end-1), ...
            tfStack(1, 1:end-1));
         ae(tsStack(1, 1, 1:end), ...
            tfStack(1, 1, 1:end));
         ae(tsStack(1, 1, 1, 1:end), ...
            tfStack(1, 1, 1, 1:end));
         ae(tsStack(1, 1, 1, 1, 1:end), ...
            tfStack(1, 1, 1, 1, 1:end));
         ae(tsStack(1, 1, 1, 1, 1, 1:end), ...
            tfStack(1, 1, 1, 1, 1, 1:end));
         
      catch mErr
         mUTErr = MException('TIFFStack:UnitTestFailed', 'Failure referencing stack with using ''end'' referencing, during test [%s]', ...
            strTestName);
         mErr = mErr.addCause(mUTErr);
         rethrow(mErr);
      end
      
      function ae(a, b)
         assert(isequal(a, b), 'TIFFStack:UnitTestFailed:ResultsNotEqual', ...
            'The referencing result was not equal between TIFFStack and standard tensor, using ''end'' referencing');
      end
   end

   % - Function to test errors with invalid references
   function TSUT_testInvalidRef(strErrorID, varargin)
      try
         % - Reference stacks, check for equality
         s.type = '()';
         s.subs = varargin;
         tfTS = subsref(tsStack, s); %#ok<NASGU>
      
      catch mErr
         % - Check whether the correct error was reported
         if ~isequal(mErr.identifier, strErrorID)
            mUTErr = MException('TIFFStack:UnitTestFailed', 'Incorrect error raised during error test with subs %s, during test [%s]. Desired error [%s], raised error [%s].', ...
               TSUT_subs2str(varargin), strTestName, strErrorID, mErr.identifier);
            mErr = mErr.addCause(mUTErr);
            rethrow(mErr);
         end
         return;
      end
      
      % - We should never get here, so raise an error
      error('TIFFStack:UnitTestFailed:ErrorNotThrown', ...
            'An error should have occurred but did not, with subs %s, during test [%s]. Desired error [%s].', ...
            TSUT_subs2str(varargin), strTestName, strErrorID);
   end
end

function TSUT_TestOverloads(tsStack, tfStack, strTestName)

   TSUT_TO_TestOverload(@sum);
   TSUT_TO_TestOverload(@nansum);
   TSUT_TO_TestOverload(@mean);
   TSUT_TO_TestOverload(@nanmean);

   function TSUT_TO_TestOverload(fhFun)
      assert(isequal(fhFun(tsStack), fhFun(tfStack)), ...
         'TIFFStack:UnitTestFailed', 'The result of called [%s] was not equal between the two stacks, during test [%s]', ...
         func2str(fhFun), strTestName);
      assert(isequal(fhFun(tsStack, 3), fhFun(tfStack, 3)), ...
         'TIFFStack:UnitTestFailed', 'The result of called [%s(x, 3)] was not equal between the two stacks, during test [%s]', ...
         func2str(fhFun), strTestName);

      % - Test 'flag' argument for sum and mean
      if (~isequal(fhFun, @nansum) && ~isequal(fhFun, @nanmean))
         assert(isequal(fhFun(tsStack, 'native'), fhFun(tfStack, 'native')), ...
            'TIFFStack:UnitTestFailed', 'The result of called [%s(x, ''native'')] was not equal between the two stacks, during test [%s]', ...
            func2str(fhFun), strTestName);
         assert(isequal(fhFun(tsStack, 3, 'native'), fhFun(tfStack, 3, 'native')), ...
            'TIFFStack:UnitTestFailed', 'The result of called [%s(x, 3, ''native''] was not equal between the two stacks, during test [%s]', ...
            func2str(fhFun), strTestName);
      end
   end

end


% - Function to test errors with invalid references
function TSUT_assertFail(strErrorID, strCommand)
try
   % - Evalulate command in called workspace
   evalin('caller', strCommand);
   
catch mErr
   % - Check whether the correct error was reported
   if ~isequal(mErr.identifier, strErrorID)
      mUTErr = MException('TIFFStack:UnitTestFailed', 'Incorrect error raised during error test with command [%s]. Desired error [%s], raised error [%s].', ...
         strCommand, strErrorID, mErr.identifier);
      mErr = mErr.addCause(mUTErr);
      rethrow(mErr);
   end
   return;
end

% - We should never get here, so raise an error
error('TIFFStack:UnitTestFailed:ErrorNotThrown', ...
   'An error should have occurred but did not, with subs %s, during test [%s]. Desired error [%s].', ...
   TSUT_subs2str(varargin), strTestName, strErrorID);
end


function strSubs = TSUT_subs2str(varargin)
   cSubsStr = cellfun(@num2str, varargin{:}, 'UniformOutput', false);
   strSubs = ['(' ...
              sprintf('[%s], ', cSubsStr{:}) ...
              sprintf('\b\b') ...
              ')'];
end
```
