## Supplementary material for "Enhancer architecture and chromatin accessibility constrain phenotypic space during development": Script-MLP: README.md

# TIFFStack
Load TIFF files into matlab fast, with lazy loading

This class allows you to access a TIFF file as a matlab tensor, while
only reading the data that you need from disk. A `TIFFStack` object
appears like a four-dimensional tensor, with dimensions for rows,
columns, frames and channels (multiple samples per pixel). These objects
can be passed transparently into other functions that expect matlab
tensors. If you need to process only a portion, or only one channel of a
TIFF stack, then this class will save you allocating the enormous
amounts of memory required to load the entire file. `TIFFStack` is also
much faster than using `imread` to read each frame of the TIFF file
separately.

`TIFFStack` attempts to use `libTiff`, which is directly supported
in recent Matlab versions. This provides dramatic speed-ups, and is a good deal faster than using `imread` or
the Matlab `Tiff` class. If `libTiff` is not available, then Matlab-only
code is used to read image data. `permute`, `ipermute`, `transpose` and
`ctranspose` are also transparently supported.

## Download and install

Clone `TIFFStack` into a directory called
*@TIFFStack*. The ampersand symbol (@) is important, since it indicates
to [Matlab] that `TIFFStack` is an object-oriented module. Add the parent
directory &mdash; not the *@TIFFStack* directory &mdash; to the [Matlab] path.

## Usage

    tsStack = TIFFStack(strFilename <, bInvert>)

A `TIFFStack` object behaves like a read-only memory mapped TIFF file.
The entire image stack is treated as a Matlab tensor. Each frame of the
file must have the same dimensions. Reading the image data is optimised
to the extent possible; the header information is only read once.

This class attempts to use the Matlab libTiff interface, if available.
If not, it uses a modified version of `tiffread` \[1, 2\] to read data.
Code is included (but disabled) to use the matlab `imread` function, but
this function returns invalid data for some TIFF formats.

## Construction of a `TIFFStack` object

    >> tsStack = TIFFStack('test.tiff'); % Construct a TIFF stack associated with a file

    >> tsStack = TIFFStack('test.tiff', true); % Indicate that the image data should be inverted 
       tsStack = 
         TIFFStack handle 
         Properties: 
            bInvert: 1 
            strFilename: [1x9 char] 
            sImageInfo: [5x1 struct] 
            strDataClass: 'uint16'

## Accessing data

    >> tsStack(:, :, 3); % Retrieve the 3rd frame of the stack, all planes 
    >> tsStack(:, :, 1, 3); % Retrieve the 3rd plane of the 1st frame 
    >> size(tsStack) % Find the size of the stack (rows, cols, frames, planes per pixel)

    ans = 
       128 128 5 1

    >> tsStack(4); % Linear indexing is supported 
    >> tsStack.bInvert = true; % Turn on data inversion

## Stacks with interleaved frame, channel and slice dimensions

Some TIFF generation software stores multiple samples per pixel as
interleaved frames in a TIFF file. Other complex stacks may include
multiple different images per frame of time (e.g. multiple cameras or
different imaged locations per frame). `TIFFStack` allows these files to be
de-interleaved, such that each conceptual data dimension has its own
referencing dimension within Matlab.

This functionality uses the optional `vnInterleavedFrameDims` argument.
This is a vector of dimensions that were interleaved into the single
frame dimension in the stack.

For example, a stack contains 2 channels of data per pixel, and 3 imaged
locations per frame, all interleaved into the TIFF frame dimension. The
stack contains 10 conceptual frames, and each frame contains 5x5 pixels.

The stack is therefore conceptually of dimensions [5 5 2 3 10 1], but
appears on disk with dimensions [5 5 60 1]. (The final dimension
corresponds to the samples-per-pixel dimension of the TIFF file).

```
>> tsStack = TIFFStack('file.tif', [], [2 3 10]);
>> size(tsStack)

ans =
     5    5    2    3   10
```


Permutation and indexing now works seamlessly on this stack, with each
conceptual dimension de-interleaved.

If desired, the final number of frames can be left off
`vnInterleavedFrameDims`; for example

```
>> tsStack = TIFFStack('file.tif', [], [2 3]);
>> size(tsStack)

ans =
     5    5    2    3   10
```

*Note*: You must be careful that you specify the dimensions in the
appropriate order, exactly as interleaved in the stack. Also, if the stack
contains multiple samples per pixel in native TIFF format, the
samples-per-pixel dimension will always be pushed to the final dimension.

## Publications

This work was published in [Frontiers in Neuroinformatics]: DR Muir and
BM Kampa. 2015. *[FocusStack and StimServer: A new open source MATLAB
toolchain for visual stimulation and analysis of two-photon calcium
neuronal imaging data](http://dx.doi.org/10.3389/fninf.2014.00085)*, **Frontiers in Neuroinformatics** 8 *85*. DOI: [dx.doi.org/10.3389/fninf.2014.00085](http://dx.doi.org/10.3389/fninf.2014.00085).
Please cite our publication in lieu of thanks, if you use this code.

## References

\[1\] Francois Nedelec, Thomas Surrey and A.C. Maggs. Physical Review
Letters 86: 3192-3195; 2001. DOI: [10.1103/PhysRevLett.86.3192]

\[2\] <http://www.cytosim.org>

## Acknowledgements

This work optionally uses `tiffread` from [Francois Nedelec] to access the data in
the TIFF file. [Matlab] includes the ability to read TIFF files in
`imread`, including niceties such as only reading a region of interest
from each frame, but imread is incredibly slow and amazingly buggy (as
of July 2011). TIFFStack uses `tiffread` in an optimised fashion, by
reading and caching the header information (the image file directories —
IFDs). Each frame can then be read directly without re-opening the file
and re-reading the IFDs.

## License
<a rel="license" href="http://creativecommons.org/licenses/by-nc-sa/4.0/"><img alt="Creative Commons License" style="border-width:0" src="https://i.creativecommons.org/l/by-nc-sa/4.0/80x15.png" /></a><br /><span xmlns:dct="http://purl.org/dc/terms/" property="dct:title">TIFFStack</span> by <a xmlns:cc="http://creativecommons.org/ns#" href="http://dylan-muir.com" property="cc:attributionName" rel="cc:attributionURL">Dylan Muir</a> is licensed under a <a rel="license" href="http://creativecommons.org/licenses/by-nc-sa/4.0/">Creative Commons Attribution-NonCommercial-ShareAlike 4.0 International License</a>.<br />Based on a work at <a xmlns:dct="http://purl.org/dc/terms/" href="http://github.com/DylanMuir/TIFFStack" rel="dct:source">http://github.com/DylanMuir/TIFFStack</a>.

  [Frontiers in Neuroinformatics]: http://www.frontiersin.org/neuroinformatics
  [FFSS]: http://journal.frontiersin.org/Journal/10.3389/fninf.2014.00085
  [10.1103/PhysRevLett.86.3192]: //dx.doi.org/10.1103/PhysRevLett.86.3192
  [Francois Nedelec]: http://www.cytosim.org
  [Matlab]: http://mathworks.com/
  [Download `TIFFStack`]: /resources/code/TIFFStack.zip
  [Matlab]: http://mathworks.com/
