## Supplementary material for "Enhancer architecture and chromatin accessibility constrain phenotypic space during development": Script-MLP: TIFFStack.m

```
% <strong>TIFFStack</strong> - Manipulate a TIFF file like a tensor
% 
% Usage: tsStack = <<strong>TIFFStack</strong>(strFilename <, bInvert, vnInterleavedFrameDims>)
% 
% A TIFFStack object behaves like a read-only memory mapped TIFF file.  The
% entire image stack is treated as a matlab tensor.  Each frame of the file must
% have the same dimensions.  Reading the image data is optimised to the extent
% possible; the header information is only read once.
%
% If this software is useful to your academic work, please cite our
% publication in lieu of thanks:
%
% D R Muir and B M Kampa, 2015. "FocusStack and StimServer: a new open
%    source MATLAB toolchain for visual stimulation and analysis of two-photon
%    calcium neuronal imaging data". Frontiers in Neuroinformatics 8 (85).
%    DOI: <a href="http://dx.doi.org/10.3389/fninf.2014.00085">10.3389/fninf.2014.00085</a>
% 
% This class attempts to use the version of tifflib built-in to recent
% versions of Matlab, if available.  Otherwise this class uses a modified
% version of tiffread [2, 3] to read data.
%
% permute, ipermute and transpose are now transparantly supported. Note
% that to read a pixel, the entire frame containing that pixel is read. So
% reading a Z-slice of the stack will read in the entire stack.
%
% Some TIFF file writing software introduces custom or poorly-formed tags.
% This causes tifflib to produce lots of warnings. These warnings can be
% ignored by setting:
%
% >> w = warning('off', 'MATLAB:imagesci:tiffmexutils:libtiffWarning');
% >> warning('off', 'MATLAB:imagesci:tifftagsread:expectedTagDataFormat');
%
% and later restored with:
%
% >> warning(w);
%
% -------------------------------------------------------------------------
% 
% <strong>Construction</strong>
% >> tsStack = TIFFStack('test.tiff');       % Construct a TIFF stack associated with a file
% 
% >> tsStack = TIFFStack('test.tiff', true); % Indicate that the image data should be inverted
% 
% tsStack = 
% 
%   TIFFStack handle
% 
%   Properties:
%          bInvert: 0
%      strFilename: [1x9 char]
%       sImageInfo: [5x1 struct]
%     strDataClass: 'uint16'
% 
% Usage:
% 
% >> tsStack(:, :, 3);     % Retrieve the 3rd frame of the stack, all planes
% 
% >> tsStack(:, :, 1, 3);  % Retrieve the 3rd plane of the 1st frame
% 
% >> size(tsStack)         % Find the size of the stack (rows, cols, frames, planes per pixel)
% 
% ans =
% 
%    128   128     5     1
% 
% >> tsStack(4);           % Linear indexing is supported
% 
% >> tsStack.bInvert = true;  % Turn on data inversion
%
% >> getFilename(tsStack)  % Retrieve the 'strFilename' property
% >> getInvert(tsStack)    % Retrive the 'bInvert' property
% >> getImageInfo(tsStack) % Retrieve the 'sImageInfo' property
% >> getDataClass(tsStack) % Retrive the 'strDataClass' property
% 
% -------------------------------------------------------------------------
%
% <strong>De-interleaving frame dimensions in complex stacks</strong>
% Some TIFF generation software stores multiple samples per pixel as
% interleaved frames in a TIFF file. Other complex stacks may include
% multiple different images per frame of time (e.g. multiple cameras or
% different imaged locations per frame). TIFFStack allows these files to be
% de-interleaved, such that each conceptual data dimension has its own
% referencing dimension within matlab.
%
% This functionality uses the optional 'vnInterleavedFrameDims' argument.
% This is a vector of dimensions that were interleaved into the single
% frame dimension in the stack.
%
% For example, a stack contains 2 channels of data per pixel, and 3 imaged
% locations per frame, all interleaved into the TIFF frame dimension. The
% stack contains 10 conceptual frames, and each frame contains 5x5 pixels.
%
% The stack is therefore conceptually of dimensions [5 5 2 3 10 1], but
% appears on disk with dimensions [5 5 60 1]. (The final dimension
% corresponds to the samples-per-pixel dimension of the TIFF file).
%
% >> tsStack = TIFFStack('file.tif', [], [2 3 10]);
% >> size(tsStack)
%
% ans =
%
%     5    5    2    3   10
%
% Permutation and indexing now works seamlessly on this stack, with each
% conceptual dimension de-interleaved.
%
% If desired, the final number of frames can be left off
% 'vnInterleavedFrameDims'; for example:
%
% >> tsStack = TIFFStack('file.tif', [], [2 3]);
% >> size(tsStack)
%
% ans =
%
%     5    5    2    3   10
%
% Note: You must be careful that you specify the dimensions in the
% appropriate order, as interleaved in the stack. Also, if the stack
% contains multiple samples per pixel in native TIFF format, the
% samples-per-pixel dimension will always be pushed to the final dimension.
%
% -------------------------------------------------------------------------
%
% <strong>ImageJ stacks</strong>
% ImageJ HyperStacks are automatically deinterleaved, if encountered. By
% default, the stacks will be presented as [Y X T Z C]. They can of course
% be permuted. If desired, the interleaving can be overridden by providing
% an explicit 'vnInterleavedFrameDims'.
%
% ImageJ writes "fake" big stacks as raw binary data, with a TIF file shim
% as a header. These appear to Matlab as a TIF file containing a single
% frame. TIFFStack loads these files using MappedTensor, when available. If
% MappedTensor is not available, a warning will be issued.
%
% References:
% [1] D R Muir and B M Kampa, 2015. "FocusStack and StimServer: a new open
%        source MATLAB toolchain for visual stimulation and analysis of two-photon
%        calcium neuronal imaging data". Frontiers in Neuroinformatics 8 (85).
%        DOI: <a href="http://dx.doi.org/10.3389/fninf.2014.00085">10.3389/fninf.2014.00085</a>
%
% [2] Francois Nedelec, Thomas Surrey and A.C. Maggs. Physical Review Letters
%        86: 3192-3195; 2001. DOI: <a href="http://dx.doi.org/10.1103/PhysRevLett.86.3192">10.1103/PhysRevLett.86.3192</a>
% 
% [3] <a href="http://www.cytosim.org">http://www.cytosim.org</a>

% Author: Dylan Muir <>
% Created: 28th June, 2011

%% Class definition

classdef TIFFStack < handle
   properties
      bInvert;             % - A boolean flag that determines whether or not the image data will be inverted
   end
   
   properties (SetAccess = private)
      strFilename = [];    % - The name of the TIFF file on disk
      sImageInfo;          % - The TIFF header information
      strDataClass;        % - The matlab class in which data will be returned
   end
   
   properties (SetAccess = private, GetAccess = private)
      bForceTiffread       % - Force the use of tiffread, rather than trying to use TiffLib
      vnDataSize;          % - Cached size of the TIFF stack
      vnApparentSize;      % - Apparent size of the TIFF stack
      TIF;                 % \
      TIF_tr31;            % |- Cached header info for tiffread31 speedups
      HEADER;              % /
      bUseTiffLib;         % - Flag indicating whether TiffLib is being used
      bMTStack;            % - Flag indicating MappedTensor is being used
      fhReadFun;           % - When using Tiff class, function for reading data
      fhSetDirFun;         % - When using Tiff class, function for setting the directory
      vnDimensionOrder;    % - Internal dimensions order to support permutation
      fhRepSum;            % - Function handle to (hopefully) accellerated repsum function
      fhCastFun;           % - The matlab function that casts data to the required return class
   end
   
   methods
      % TIFFStack - CONSTRUCTOR
      function oStack = TIFFStack(strFilename, bInvert, vnInterleavedFrameDims, bForceTiffread)
         % - Check usage
         if (~exist('strFilename', 'var') || ~ischar(strFilename))
            help TIFFStack;
            error('TIFFStack:Usage', ...
                  '*** TIFFStack: Incorrect usage.');
         end
         
         % - Should we force TIFFStack to use tiffread, rather than libTiff?
         if (~exist('bForceTiffread', 'var') || isempty(bForceTiffread))
            bForceTiffread = false;
         end
         oStack.bForceTiffread = bForceTiffread;
         
         % - Can we use the accelerated TIFF library?
         if (exist('tifflib') ~= 3) %#ok<EXIST>
            % - Try to copy the library
            strTiffLibLoc = which('/private/tifflib');
            strTIFFStackLoc = fileparts(which('TIFFStack'));
            copyfile(strTiffLibLoc, fullfile(strTIFFStackLoc, 'private'), 'f');
         end
         
         oStack.bUseTiffLib = (exist('tifflib') == 3) & ~bForceTiffread; %#ok<EXIST>
         
         if (~oStack.bUseTiffLib)
            warning('TIFFStack:SlowAccess', ...
                    '--- TIFFStack: Using slower non-TiffLib access.');
         end
         
         % - Get accelerated repsum function, if possible
         oStack.fhRepSum = GetMexFunctionHandles;
         
         % - Check for inversion flag
         if (~exist('bInvert', 'var') || isempty(bInvert))
            bInvert = false;
         end
         oStack.bInvert = bInvert;
         
         % - Check for frame dimensions
         if (~exist('vnInterleavedFrameDims', 'var'))
            vnInterleavedFrameDims = [];
         else
            validateattributes(vnInterleavedFrameDims, {'single', 'double'}, {'integer', 'real', 'positive'}, ...
               'TIFFStack', 'vnInterleavedFrameDims');
         end

         % - See if filename exists
         if (~exist(strFilename, 'file'))
            error('TIFFStack:InvalidFile', ...
                  '*** TIFFStack: File [%s] does not exist.', strFilename);
         end
         
         % - Assign absolute file path to stack
         strFilename = get_full_file_path(strFilename);
         oStack.strFilename = strFilename;
         
         % - Get image information
         try
            % - Read and store image information (using tiffread for speed and compatibility)
            [oStack.TIF, oStack.HEADER, sInfo] = tiffread31_header(strFilename);
            oStack.TIF_tr31 = oStack.TIF;
            
            % - Detect a ImageJ fake BigTIFF stack
            [bIsImageJBigStack, bIsImageJHyperStack, vnStackDims, vnInterleavedIJFrameDims] = IsImageJBigStack(tiffread31_readtags(oStack.TIF_tr31, oStack.HEADER, 1), numel(oStack.HEADER));
            
            % - Handle ImageJ big stacks with MappedTensor
            if (bIsImageJBigStack)
               [oStack.TIF, oStack.bMTStack, oStack.strDataClass] = OpenImageJBigStack(oStack, vnStackDims);
               
               % - Could we use a MappedTensor?
               if (~oStack.bMTStack)
                  % - No, so just access the first frame
                  bIsImageJBigStack = false; %#ok<NASGU>
                  bIsImageJHyperStack = false;
                  
                  warning('TIFFStack:ImageJBigStackUnsupported', ...
                          '--- TIFFStack: Warning: This is an ImageJ "fake" TIF file. MappedTensor must be available to read this file.');
               end
            end
            
            % - Detect a very long stack
            if ((numel(sInfo) > 2^16) && oStack.bUseTiffLib)
               warning('TIFFStack:LongStack', ...
                       '--- TIFFSTack: Warning: This stack has more than 2^16 frames, so Matlab/tifflib cannot read it natively.\n       Using slower non-Tifflib access.');
               oStack.bUseTiffLib = false;
            end
            
            % - Deinterleave hyperstacks automatically
            bImageJDeinterleaving = bIsImageJHyperStack && isempty(vnInterleavedFrameDims);
            if bImageJDeinterleaving
               vnInterleavedFrameDims = vnInterleavedIJFrameDims;
            end

            % - Initialise object, depending on underlying access method
            if (oStack.bMTStack)
               % - Fix up stack size
               sInfo = repmat(sInfo(1), vnStackDims(3), 1);
               
            elseif (oStack.bUseTiffLib)
               % - Create a Tiff object
               oStack.TIF = tifflib('open', strFilename, 'r');
               
               % - Check data format
               if(TiffgetTag(oStack.TIF, 'Photometric') == Tiff.Photometric.YCbCr)
                  error('TIFFStack:UnsupportedFormat', ...
                        '*** TIFFStack: YCbCr images are not supported.');
               end
               
               % - Use Tiff to get the data class for this tiff
               nDataClass = TiffgetTag(oStack.TIF, 'SampleFormat');
               switch (nDataClass)
                  case Tiff.SampleFormat.UInt
                     switch (sInfo(1).BitsPerSample(1))
                        case 1
                           oStack.strDataClass = 'logical';

                        case 8
                           oStack.strDataClass = 'uint8';
                           
                        case 16
                           oStack.strDataClass = 'uint16';
                           
                        case 32
                           oStack.strDataClass = 'uint32';
                           
                        case 64
                           oStack.strDataClass = 'uint64';
                           
                        otherwise
                           error('TIFFStack:UnsupportedFormat', ...
                                 '*** TIFFStack: The sample format of this TIFF stack is not supported.');
                     end
                     
                  case Tiff.SampleFormat.Int
                     switch (sInfo(1).BitsPerSample(1))
                        case 1
                           oStack.strDataClass = 'logical';
                           
                        case 8
                           oStack.strDataClass = 'int8';
                           
                        case 16
                           oStack.strDataClass = 'int16';
                           
                        case 32
                           oStack.strDataClass = 'int32';
                           
                        case 64
                           oStack.strDataClass = 'int64';                           
                           
                        otherwise
                           error('TIFFStack:UnsupportedFormat', ...
                              '*** TIFFStack: The sample format of this TIFF stack is not supported.');
                     end
                     
                  case Tiff.SampleFormat.IEEEFP
                     switch (sInfo(1).BitsPerSample(1))
                        case {1, 8, 16, 32}
                           oStack.strDataClass = 'single';

                        case 64
                           oStack.strDataClass = 'double';
                           
                        otherwise
                           error('TIFFStack:UnsupportedFormat', ...
                              '*** TIFFStack: The sample format of this TIFF stack is not supported.');
                     end
                     
                  otherwise
                     error('TIFFStack:UnsupportedFormat', ...
                           '*** TIFFStack: The sample format of this TIFF stack is not supported.');
               end
                              
               % -- Assign accelerated reading function
               strReadFun = 'TS_read_Tiff';
               
               % - Tiled or striped
               if (tifflib('isTiled', oStack.TIF))
                  strReadFun = [strReadFun '_tiled'];
               else
                  strReadFun = [strReadFun '_striped'];
               end
               
               % - Chunky or planar
               if (isequal(TiffgetTag(oStack.TIF, 'PlanarConfiguration'), Tiff.PlanarConfiguration.Chunky))
                  strReadFun = [strReadFun '_chunky'];
               elseif (isequal(TiffgetTag(oStack.TIF, 'PlanarConfiguration'), Tiff.PlanarConfiguration.Separate))
                  strReadFun = [strReadFun '_planar'];
               else
                  error('TIFFStack:UnsupportedFormat', ...
                        '*** TIFFStack: The planar configuration of this TIFF stack is not supported.');
               end
               
               strSetDirFun = 'TS_set_directory';
               
               % - Check for zero-based referencing
               try
                   tifflib('computeStrip', oStack.TIF, 0);
               catch
                  strReadFun = [strReadFun '_pre2014'];
                  strSetDirFun = [strSetDirFun '_pre2014'];
               end
               
               % - Convert into function handles
               oStack.fhReadFun = str2func(strReadFun);
               oStack.fhSetDirFun = str2func(strSetDirFun);
               
               % - Fix up rows per strip (inconsistency between Windows and
               % OS X Tifflib
               nRowsPerStrip = TiffgetTag(oStack.TIF, 'RowsPerStrip');
               if ~isfield(sInfo, 'RowsPerStrip') || (nRowsPerStrip ~= sInfo(1).RowsPerStrip)
                   [sInfo.RowsPerStrip] = deal(nRowsPerStrip);
               end
               
               % - Attempt to read tile width and length
               try
                  [sInfo.TileWidth] = deal(TiffgetTag(oStack.TIF, 'TileWidth'));
               catch
                  [sInfo.TileWidth] = deal(1);
               end
               
               try
                  [sInfo.TileLength] = deal(TiffgetTag(oStack.TIF, 'TileLength'));
               catch
                  [sInfo.TileLength] = deal(1);
               end
               
               % - Read max and min sample values
               [sInfo.MaxSampleValue] = deal(TiffgetTag(oStack.TIF, 'MaxSampleValue'));
               [sInfo.MinSampleValue] = deal(TiffgetTag(oStack.TIF, 'MinSampleValue'));
               
            else
               % - Read TIFF header for tiffread31
               % [oStack.TIF, oStack.HEADER] = tiffread31_header(strFilename);
               oStack.TIF.file = fopen(strFilename, 'r', 'l');

               % - Use tiffread31 to get the data class for this tiff
               fPixel = tiffread31_readimage(oStack.TIF, oStack.HEADER, 1);
               fPixel = fPixel(1, 1, :);
               oStack.strDataClass = class(fPixel);
            end
            
            % - Use imread to get the data class for this tiff
            % fPixel = imread(strFilename, 'TIFF', 1, 'PixelRegion', {[1 1], [1 1]});
            % oStack.strDataClass = class(fPixel);
            
            % -- Assign casting function
            oStack.fhCastFun = str2func(oStack.strDataClass);
            
            % - Record stack size
            if (oStack.bMTStack)
               oStack.vnDataSize = vnStackDims;
               oStack.vnApparentSize = oStack.vnDataSize;
               
               % - Initialise dimension order
               oStack.vnDimensionOrder = 1:numel(oStack.vnApparentSize);
            
               % - Permute first two dimensions
               oStack = permute(oStack, [2 1 3:numel(vnStackDims)]);
               
            else
               oStack.vnDataSize = [sInfo(1).Height sInfo(1).Width numel(sInfo) sInfo(1).SamplesPerPixel];

               % - Initialise dimension order
               oStack.vnDimensionOrder = 1:numel(oStack.vnDataSize);
            end
            
            % - Initialize apparent stack size, de-interleaving along the frame dimension
            if isempty(vnInterleavedFrameDims)
               % - No de-interleaving
               oStack.vnApparentSize = oStack.vnDataSize;
            
            elseif (prod(vnInterleavedFrameDims) ~= oStack.vnDataSize(3))
               % - Be lenient by allowing frames dimension to be left out of arguments
               if (mod(oStack.vnDataSize(3), prod(vnInterleavedFrameDims)) == 0)
                  % - Work out number of apparent frames
                  nNumApparentFrames = oStack.vnDataSize(3) ./ prod(vnInterleavedFrameDims);
                  oStack.vnApparentSize = [oStack.vnDataSize(1:2) vnInterleavedFrameDims(:)' nNumApparentFrames oStack.vnDataSize(4)];
                  oStack.vnDimensionOrder = 1:numel(oStack.vnApparentSize);
                  
               else
                  % - Incorrect total number of deinterleaved frames
                  error('TIFFStack:WrongFrameDims', ...
                     '*** TIFFStack: When de-interleaving a stack, the total number of frames must not change.');
               end
               
            else
               % - Record apparent stack dimensions
               oStack.vnApparentSize = [oStack.vnDataSize(1:2) vnInterleavedFrameDims(:)' oStack.vnDataSize(4)];
               oStack.vnDimensionOrder = 1:numel(oStack.vnApparentSize);
            end
            
            % - Fix up dimensions order for ImageJ HyperStack
            if (bImageJDeinterleaving)
               oStack = permute(oStack, [1 2 5 4 3]);
            end

            % - Record image information
            oStack.sImageInfo = sInfo;
            
         catch mErr
            base_ME = MException('TIFFStack:InvalidFile', ...
                  '*** TIFFStack: Could not open file [%s].', strFilename);
            new_ME = addCause(base_ME, mErr);
            throw(new_ME);
         end
      end
      
      % delete - DESTRUCTOR
      function delete(oStack)
         if (oStack.bUseTiffLib)
            % - Close the TIFF file, if opened by TiffLib
            if (~isempty(oStack.TIF))
               tifflib('close', oStack.TIF);
            end
         end
         
         % - Close the TIFF file, if opened by tiffread31_header
         if (isfield(oStack.TIF_tr31, 'file') && ~isempty(fopen(oStack.TIF_tr31.file)))
            fclose(oStack.TIF_tr31.file);
         end
      end

      % diagnostic - METHOD Display some diagnostics about a stack
      function diagnostic(oStack)
         disp(oStack);
         fprintf('<strong>Private properties:</strong>\n');
         fprintf('   vnDataSize: ['); fprintf('%d ', oStack.vnDataSize); fprintf(']\n');
         fprintf('   vnApparentSize: ['); fprintf('%d ', oStack.vnApparentSize); fprintf(']\n');
         fprintf('   vnDimensionOrder: ['); fprintf('%d ', oStack.vnDimensionOrder); fprintf(']\n');
         fprintf('   bUseTiffLib: %d\n', oStack.bUseTiffLib);
         fprintf('   bMTStack: %d\n', oStack.bMTStack);
         fprintf('   fhReadFun: %s\n', func2str(oStack.fhReadFun));
         fprintf('   fhSetDirFun: %s\n', func2str(oStack.fhSetDirFun));
         fprintf('   fhRepSum: %s\n', func2str(oStack.fhRepSum));
         fprintf('   fhCastFun: %s\n', func2str(oStack.fhCastFun));
      end
      
      function [TIF, HEADER] = diagnostic_HEADER(oStack)
         TIF = oStack.TIF;
         HEADER = oStack.HEADER;
      end
         

%% --- Overloaded subsref

      function [varargout] = subsref(oStack, S)
         switch S(1).type
            case '()'
               % - Test for valid subscripts
               cellfun(@isvalidsubscript, S.subs);
               
               % - Record stack size
               nNumRefDims = numel(S.subs);
               
               vnReferencedTensorSize = size(oStack);
               nNumNZStackDims = numel(vnReferencedTensorSize);
               nNumTotalStackDims = max(numel(oStack.vnDimensionOrder), nNumNZStackDims);
               
               vnFullTensorSize = vnReferencedTensorSize;
               vnFullTensorSize(nNumNZStackDims+1:nNumTotalStackDims) = 1;
               vnFullTensorSize(vnFullTensorSize == 0) = 1;
               
               bLinearIndexing = false;
               
               % - Convert logical indexing to indices
               for (nDim = 1:numel(S.subs))
                  if (islogical(S.subs{nDim}))
                     S.subs{nDim} = find(S.subs{nDim});
                  end
               end

               % - Check dimensionality and trailing dimensions
               if (nNumRefDims == 1)
                  % - Catch "read whole stack" case
                  if (iscolon(S.subs{1}))
                     S.subs = num2cell(repmat(':', 1, nNumNZStackDims));
                     vnRetDataSize = [prod(vnReferencedTensorSize), 1];

                  else
                     % - Get equivalent subscripted indexes and permute
                     vnTensorSize = size(oStack);
                     if any(S.subs{1}(:) > prod(vnTensorSize))
                        error('TIFFStack:badsubscript', ...
                           '*** TIFFStack: Index exceeds stack dimensions.');
                     else
                        [cIndices{1:nNumTotalStackDims}] = ind2sub(vnTensorSize, S.subs{1});
                     end
                     
                     % - Permute dimensions
                     vnInvOrder(oStack.vnDimensionOrder(1:nNumTotalStackDims)) = 1:nNumTotalStackDims;
                     S.subs = cIndices(vnInvOrder(vnInvOrder ~= 0));
                     vnRetDataSize = size(S.subs{1});

                     bLinearIndexing = true;
                  end
                  
                  
               elseif (nNumRefDims < nNumNZStackDims)
                  % - Wrap up trailing dimensions, matlab style, using linear indexing
                  vnReferencedTensorSize(nNumRefDims) = prod(vnReferencedTensorSize(nNumRefDims:end));
                  vnReferencedTensorSize = vnReferencedTensorSize(1:nNumRefDims);
                  
                  % - Catch "read whole stack" case
                  if (all(cellfun(@iscolon, S.subs)))
                     [S.subs{nNumRefDims+1:nNumTotalStackDims}] = deal(':');
                     vnRetDataSize = vnReferencedTensorSize;

                  else
                     % - Convert to linear indexing
                     bLinearIndexing = true;
                     [S.subs{1}, vnRetDataSize] = GetLinearIndicesForRefs(S.subs, vnReferencedTensorSize, oStack.fhRepSum);
                     S.subs = S.subs(1);
                     [S.subs{1:nNumTotalStackDims}] = ind2sub(vnFullTensorSize, S.subs{1});

                     % - Inverse permute index order
                     vnInvOrder(oStack.vnDimensionOrder(1:nNumTotalStackDims)) = 1:nNumTotalStackDims;
                     S.subs = S.subs(vnInvOrder(vnInvOrder ~= 0));
                  end
                                    
               elseif (nNumRefDims == nNumNZStackDims)
                  % - Check for colon references
                  vbIsColon = cellfun(@iscolon, S.subs);
                  vnRetDataSize = cellfun(@numel, S.subs);                  
                  vnRetDataSize(vbIsColon) = vnReferencedTensorSize(vbIsColon);
                  
                  % - Permute index order
                  S.subs(nNumNZStackDims+1:nNumTotalStackDims) = {1};
                  vnInvOrder(oStack.vnDimensionOrder(1:nNumTotalStackDims)) = 1:nNumTotalStackDims;
                  S.subs = S.subs(vnInvOrder(vnInvOrder ~= 0));
                  
               else % (nNumRefDims > nNumNZStackDims)
                  % - Check for non-colon references
                  vbIsColon = cellfun(@iscolon, S.subs);
                  
                  % - Check for non-unitary references
                  vbIsUnitary = cellfun(@(c)(isequal(c, 1)), S.subs);
                  
                  % - Check for non-empty references
                  vbIsEmpty = cellfun(@isempty, S.subs);
                  
                  % - Check only trailing dimensions
                  vbTrailing = [false(1, nNumNZStackDims) true(1, nNumRefDims-nNumNZStackDims)];
                  
                  % - Check trailing dimensions for inappropriate indices
                  if (any(vbTrailing & (~vbIsColon & ~vbIsUnitary & ~vbIsEmpty)))
                     % - This is an error
                     error('TIFFStack:badsubscript', ...
                        '*** TIFFStack: Index exceeds stack dimensions.');
                  end
                  
                  % - Catch empty refs
                  if (~any(vbIsEmpty))
                     % - Only keep relevant dimensions
                     S.subs = S.subs(1:nNumNZStackDims);
                  end
                  
                  % - Determine returned data size
                  vnReferencedTensorSize(nNumNZStackDims+1:nNumRefDims) = 1;
                  vnReferencedTensorSize(vnReferencedTensorSize == 0) = 1;
                  vbIsColon = cellfun(@iscolon, S.subs);
                  vnRetDataSize = cellfun(@numel, S.subs);
                  vnRetDataSize(vbIsColon) = vnReferencedTensorSize(vbIsColon);
                  
                  % - Permute index order
                  S.subs(nNumNZStackDims+1:nNumTotalStackDims) = {1};
                  vnInvOrder(oStack.vnDimensionOrder(1:nNumTotalStackDims)) = 1:nNumTotalStackDims;
                  S.subs = S.subs(vnInvOrder(vnInvOrder ~= 0));
               end
               
               % - Catch empty refs
               if (prod(vnRetDataSize) == 0)
                  [varargout{1:nargout}] = zeros(vnRetDataSize);
                  return;
               end
               
               % - Re-interleave frame indices for deinterleaved stacks
               if (numel(oStack.vnApparentSize) > 4)
                  % - Record output data size in deinterleaved space
                  if (~bLinearIndexing)
                     vnOutputSize = cellfun(@numel, S.subs);
                     vbIsColon = cellfun(@iscolon, S.subs);
                     vnOutputSize(vbIsColon) = oStack.vnApparentSize(vbIsColon);
                  end
                  
                  % - Get frame 
                  cFrameSubs = S.subs(3:end-1);
                  if all(cellfun(@iscolon, cFrameSubs))
                     S.subs = {S.subs{1} S.subs{2}  ':' S.subs{end}};
                  else
                     if (bLinearIndexing)
                        vnFrameIndices = sub2ind(oStack.vnApparentSize(3:end-1), cFrameSubs{:});
                     else
                        tnFrameIndices = reshape(1:oStack.vnDataSize(3), ...
                           oStack.vnApparentSize(3:end-1));
                        vnFrameIndices = tnFrameIndices(cFrameSubs{:});
                     end
                     
                     % - Construct referencing subscripts for raw stack
                     S.subs = [S.subs(1:2) reshape(vnFrameIndices, [], 1) S.subs(end)];
                  end
               end

               % - Access stack (MappedTensor or tifflib or tiffread)
               if (oStack.bMTStack)
                  tfData = TS_read_data_MappedTensor(oStack, S.subs, bLinearIndexing);
               elseif (oStack.bUseTiffLib)
                  tfData = TS_read_data_Tiff(oStack, S.subs, bLinearIndexing);
               else
                  tfData = TS_read_data_tiffread(oStack, S.subs, bLinearIndexing);
               end
               
               % - Permute dimensions, if linear indexing has not been used
               if (~bLinearIndexing)
                  % - Reshape resulting data in case of deinterleaved stacks
                  if (numel(oStack.vnApparentSize) > 4)
                     tfData = reshape(tfData, vnOutputSize);
                  end

                  tfData = permute(tfData, oStack.vnDimensionOrder);
               end
               
               % - Reshape returned data to concatenate trailing dimensions (just as matlab does)
               if (~isequal(size(tfData), vnRetDataSize))
                  tfData = reshape(tfData, vnRetDataSize);
               end
               
               [varargout{1:nargout}] = tfData;
               
            otherwise
               error('TIFFStack:InvalidReferencing', ...
                     '*** TIFFStack: Only ''()'' referencing is supported by TIFFStacks.');
         end
      end
      
%% --- Getter methods

      function strFilename = getFilename(oStack)
         strFilename = oStack.strFilename;
      end

      function sImageInfo = getImageInfo(oStack)
         sImageInfo = oStack.sImageInfo;
      end
      
      function vsTags = getImageTags(oStack, vnFrames)
         % getImageTags - METHOD Read TIFF tags for individual frames
         %
         % Usage: vsTags = getImageTags(oStack, vnFrames)
         %
         % 'oStack' is a TIFFStack object. 'vnFrames' is a vector of frame
         % indices into the stack.
         %
         % 'vsTags' will be a struct array, with each element of the array
         % containing all tags for the corresponding frame index in
         % 'vnFrames'.
         
         if (nargin < 2)
            help TIFFStack/getImageTags;
            error('TIFFStack:Usage', 'TIFFStack/getImageTags: ''vnFrames'' is a required argument.');
         end
         
         % - Extract tags for these frames
         vsTags = tiffread31_readtags(oStack.TIF_tr31, oStack.HEADER, vnFrames);
      end
      
      function bInvert = getInvert(oStack)
         bInvert = oStack.bInvert;
      end
      
      function strDataClass = getDataClass(oStack)
         strDataClass = oStack.strDataClass;
      end
      
%% --- Overloaded numel, size, ndims, permute, ipermute, ctranspose, transpose, cat, horzcat, vertcat
      function [n] = numel(oStack, varargin)
         n = prod(size(oStack)); %#ok<PSIZE>
      end

      % size - METHOD Overloaded size function
      function [varargout] = size(oStack, vnDimensions)
         % - Get original tensor size, and extend dimensions if necessary
         vnApparentSize = oStack.vnApparentSize; %#ok<PROPLC,PROP>
         vnApparentSize(end+1:numel(oStack.vnDimensionOrder)) = 1; %#ok<PROPLC,PROP>
         
         % - Return the size of the tensor data element, permuted
         vnSize = vnApparentSize(oStack.vnDimensionOrder); %#ok<PROPLC,PROP>
         
         % - Trim trailing unitary dimensions
         vbIsUnitary = vnSize == 1;
         if (vbIsUnitary(end))
            nLastNonUnitary = find(~vbIsUnitary, 1, 'last');
            if (nLastNonUnitary < numel(vnSize))
               vnSize = vnSize(1:nLastNonUnitary);
            end
         end
         
         % - Return specific dimension(s)
         if (exist('vnDimensions', 'var'))
            if (~isnumeric(vnDimensions) || any(vnDimensions < 1))
               error('TIFFStack:DimensionMustBePositiveInteger', ...
                  '*** TIFFStack: Dimensions argument must be a positive integer.');
            end
            
            vbExtraDimensions = vnDimensions > numel(vnSize);
            
            % - Return the specified dimension(s)
            vnSizeOut(~vbExtraDimensions) = vnSize(vnDimensions(~vbExtraDimensions));
            vnSizeOut(vbExtraDimensions) = 1;
         else
            vnSizeOut = vnSize;
         end
         
         % - Handle differing number of size dimensions and number of output
         % arguments
         nNumArgout = max(1, nargout);
         
         if (nNumArgout == 1)
            % - Single return argument -- return entire size vector
            varargout{1} = vnSizeOut;
            
         elseif (nNumArgout <= numel(vnSizeOut))
            % - Several return arguments -- return single size vector elements,
            % with the remaining elements grouped in the last value
            varargout(1:nNumArgout-1) = num2cell(vnSizeOut(1:nNumArgout-1));
            varargout{nNumArgout} = prod(vnSizeOut(nNumArgout:end));
            
         else %(nNumArgout > numel(vnSize))
            % - Output all size elements
            varargout(1:numel(vnSizeOut)) = num2cell(vnSizeOut);
            
            % - Deal out trailing dimensions as '1'
            varargout(numel(vnSizeOut)+1:nNumArgout) = {1};
         end
      end
      
      % ndims - METHOD Overloaded ndims function
      function [nNumDims] = ndims(oStack)
         nNumDims = numel(size(oStack));
      end
      
      % permute - METHOD Overloaded permute function
      function [oStack] = permute(oStack, vnNewOrder)
         oStack.vnDimensionOrder(1:numel(vnNewOrder)) = oStack.vnDimensionOrder(vnNewOrder);
      end
      
      % ipermute - METHOD Overloaded ipermute function
      function [oStack] = ipermute(oStack, vnOldOrder)
         vnNewOrder(vnOldOrder) = 1:numel(vnOldOrder);
         oStack = permute(oStack, vnNewOrder);
      end
      
      % ctranspose - METHOD Overloaded ctranspose function
      function [oStack] = cstranspose(oStack)
         oStack = transpose(oStack);
      end
      
      % transpose - METHOD Overloaded transpose function
      function [oStack] = transpose(oStack)
         oStack = permute(oStack, [2 1]);
      end

      % cat - METHOD Overloaded cat, horzcat, vertcat functions
      function [varargout] = cat(varargin) %#ok<STOUT>
         error('TIFFStack:Concatenation', ...
            '*** TIFFStack: Concatenation is not supported by TIFFStack.');
      end
      
      function [varargout] = horzcat(varargin) %#ok<STOUT>
         error('TIFFStack:Concatenation', ...
            '*** TIFFStack: Concatenation is not supported by TIFFStack.');
      end
      
      function [varargout] = vertcat(varargin) %#ok<STOUT>
         error('TIFFStack:Concatenation', ...
            '*** TIFFStack: Concatenation is not supported by TIFFStack.');
      end      
      
%% --- Overloaded end

      function nLength = end(oStack, nEndDim, nTotalRefDims)
         vnSizes = size(oStack);
         if (nEndDim < nTotalRefDims)
            nLength = vnSizes(nEndDim);
         else
            nLength = prod(vnSizes(nEndDim:end));
         end
      end

%% --- Overloaded sum, nansum, mean, nanmean

      % sum - METHOD Overloaded sum function
      function [tfResult, tnNumNonNaNs] = sum(oStack, nDim, flag, bIgnoreNaNs)
         % - If this function is called, it must be a sum over the entire stack
         if nargin==2 && ischar(nDim)
            flag = nDim;
         elseif nargin < 3
            flag = 'default';
         end
         
         if nargin == 1 || (nargin == 2 && ischar(nDim))
            nDim = find(size(oStack)~=1,1);
            if isempty(nDim), nDim = 1; end
         end
         
         if (~exist('bIgnoreNaNs', 'var'))
            bIgnoreNaNs = true;
         end
         
         % - Set up referencing
         sSubs.type = '()';
         sSubs.subs = repmat({':'}, 1, ndims(oStack));
         
         % - Loop to perform sum in double precision
         for (nIndex = 1:size(oStack, nDim))
            sSubs.subs{nDim} = nIndex;
            tfSlice = double(subsref(oStack, sSubs));

            % - Ignore NaNs, if requested
            if (bIgnoreNaNs)
               if (~exist('tnNumNonNaNs', 'var'))
                  tnNumNonNaNs = zeros(size(tfSlice));
               end
               
               tnNumNonNaNs(~isnan(tfSlice)) = tnNumNonNaNs(~isnan(tfSlice)) + 1;
               
               % - Set NaNs to zero
               tfSlice(isnan(tfSlice)) = 0;
            end
            
            % - Perform sum
            if (~exist('tfResult', 'var'))
               tfResult = tfSlice;
            else
               tfResult = tfResult + tfSlice;
            end
         end
         
         % - Cast result to native class, if necessary
         if (strcmp(flag, 'native'))
            tfResult = oStack.fhCastFun(tfResult);
         end
      end
      
      % nansum - METHOD Overloaded nansum function
      function tfResult = nansum(oStack, nDim)
         if nargin == 1
            nDim = find(size(oStack)~=1,1);
            if isempty(nDim), nDim = 1; end
         end
         
         % - Call "sum" with "bIgnoreNaNs" set to true
         tfResult = sum(oStack, nDim, 'default', true);
      end
         
      % mean - METHOD Overloaded mean function
      function y = mean(x,dim,flag, bIgnoreNaNs)
         
         if (~exist('bIgnoreNaNs', 'var'))
            bIgnoreNaNs = true;
         end
         
         if nargin==2 && ischar(dim)
            flag = dim;
         elseif nargin < 3
            flag = 'default';
         end
         
         if nargin == 1 || (nargin == 2 && ischar(dim))
            dim = find(size(x)~=1,1);
            if isempty(dim), dim = 1; end
         end
         
         % - Compute sum in double
         [y, tnNumNonNaNs] = sum(x, dim, 'double', bIgnoreNaNs);
         y = y ./ tnNumNonNaNs;
         
         % - Re-cast result, if necessary
         if (strcmp(flag, 'native'))
            y = x.fhCastFun(y);
         end
      end
      
      % nanmean - METHOD Overloaded nanmean
      function y = nanmean(x,dim)
         if nargin == 1
            dim = find(size(x)~=1,1);
            if isempty(dim), dim = 1; end
         end
         
         % - Call mean
         y = mean(x,dim,'default', true);
      end
      
%% --- Overloaded sort, prctile

      % sort - METHOD Overloaded sort function
      function tfSorted = sort(oStack, varargin)
         % - Warn about loss of function 
         warning('TIFFStack:LostTIFFStack', '--- TIFFStack/sort: Warning: Data returned by ''sort'' is no longer a ''TIFFStack'' object.');
         
         % - Set up referencing
         sSubs.type = '()';
         sSubs.subs = repmat({':'}, 1, ndims(oStack));
         
         % - Just load stack and pass parameters to sort
         tfSorted = sort(subsref(oStack, sSubs), varargin{:});
      end
      
      % prctile - METHOD Overloaded prctile function
      function tfPrctile = prctile(oStack, varargin)
         % - Warn about loss of function 
         warning('TIFFStack:LostTIFFStack', '--- TIFFStack/prctile: Warning: Data returned by ''prctile'' is no longer a ''TIFFStack'' object.');
         
         % - Set up referencing
         sSubs.type = '()';
         sSubs.subs = repmat({':'}, 1, ndims(oStack));
         
         % - Just load stack and pass parameters to sort
         tfPrctile = prctile(subsref(oStack, sSubs), varargin{:});         
      end

%% --- Overloaded cast methods

      % cast - METHOD Overloaded cast function
      function tfStack = cast(oStack, strClass, oTemplate)
         if (nargin == 3) && (strcmp(strClass, 'like'))
            strClass = class(oTemplate);
         end
         
         % - Construct a casting function
         fhCastFun = str2func(strClass); %#ok<PROPLC,PROP>
         
         % - Set up referencing
         sSubs.type = '()';
         sSubs.subs = repmat({':'}, 1, ndims(oStack));

         % - Load the entire stack
         tfStack = subsref(oStack, sSubs);
         
         % - Cast the result
         tfStack = fhCastFun(tfStack); %#ok<PROPLC,PROP>
      end

      function tfStack = double(oStack)
         tfStack = cast(oStack, 'double');
      end
      
      function tfStack = single(oStack)
         tfStack = cast(oStack, 'single');
      end
      
      function tfStack = int8(oStack)
         tfStack = cast(oStack, 'int8');
      end
         
      function tfStack = int16(oStack)
         tfStack = cast(oStack, 'int16');
      end
         
      function tfStack = int32(oStack)
         tfStack = cast(oStack, 'int16');
      end

      function tfStack = int64(oStack)
         tfStack = cast(oStack, 'int16');
      end
      
      function tfStack = uint8(oStack)
         tfStack = cast(oStack, 'int8');
      end
         
      function tfStack = uint16(oStack)
         tfStack = cast(oStack, 'int16');
      end
         
      function tfStack = uint32(oStack)
         tfStack = cast(oStack, 'int16');
      end

      function tfStack = uint64(oStack)
         tfStack = cast(oStack, 'int16');
      end
      
%% --- Overloaded test methods

      function bIsNumeric = isnumeric(~)
         bIsNumeric = true;
      end
      
      function bIsFloat = isfloat(oStack)
         switch (oStack.strDataClass)
            case {'single', 'double'}
               bIsFloat = true;
               
            otherwise
               bIsFloat = false;
         end
      end
      
      function bIsInteger = isinteger(oStack)
         switch (oStack.strDataClass)
            case {'int8', 'int16', 'int32', 'int64', 'uint8', 'uint16', 'uint32', 'uint64'}
               bIsInteger = true;
               
            otherwise
               bIsInteger = false;
         end
      end
            
%% --- Property accessors

      % set.bInvert - SETTER method for 'bInvert'
      function set.bInvert(oStack, bInvert)
         % - Check contents
         if (~islogical(bInvert) || ~isscalar(bInvert))
            error('TIFFStack:InvalidArgument', ...
                  '*** TIFFStack/set.bInvert: ''bInvert'' must be a logical scalar.');
         else
            % - Assign bInvert value
            oStack.bInvert = bInvert;
         end
      end            
      
%% --- Overloaded save method
      % saveobj - Save method
      function sSerialised = saveobj(tsStack)
         % - Serialise object and remove transient properties
         w = warning('off', 'MATLAB:structOnObject');
         sSerialised = struct(tsStack);
         sSerialised = rmfield(sSerialised, {'TIF', 'HEADER', 'fhReadFun', 'fhSetDirFun', 'fhRepSum', 'fhCastFun'});
         warning(w);
      end
      
   end

%% -- Overloaded load method

   methods (Static)
      
      % loadobj - Load method
      function oStack = loadobj(sSavedVar)
         % - Create a new TIFFStack
         oStack = TIFFStack(sSavedVar.strFilename, sSavedVar.bInvert, [], ...
                            sSavedVar.bForceTiffread);

         % - Adjust dimensions to look like saved stack
         oStack.vnApparentSize = sSavedVar.vnApparentSize;
         oStack.vnDimensionOrder = sSavedVar.vnDimensionOrder;
      end
   end
end

%% --- Helper functions ---

% TS_read_data_tiffread - FUNCTION Read the requested pixels from the TIFF file (using tiffread31)
%
% Usage: [tfData] = TS_read_data_imread(oStack, cIndices)
%
% 'oStack' is a TIFFStack.  'cIndices' are the indices passed in from subsref.
% Colon indexing will be converted to full range indexing.  cIndices is a cell
% array with the format {rows, cols, frames, slices}.  Slices are RGB or CMYK
% or so on.

function [tfData] = TS_read_data_tiffread(oStack, cIndices, bLinearIndexing)
   % - Fix up final index dimensions, if necessary
   cIndices(end+1:4) = {1};
   
   % - Convert colon indexing
   vbIsColon = cellfun(@iscolon, cIndices);
   
   for (nColonDim = find(vbIsColon))
      cIndices{nColonDim} = 1:oStack.vnDataSize(nColonDim);
   end

   % - Fix up subsample detection for unitary dimensions
   vbIsOne = cellfun(@(c)isequal(c, 1), cIndices);
   vbIsColon(~vbIsColon) = vbIsOne(~vbIsColon) & (oStack.vnDataSize(~vbIsColon) == 1);
   
   % - Check ranges
   vnMinRange = cellfun(@(c)(min(c(:))), cIndices(~vbIsColon));
   vnMaxRange = cellfun(@(c)(max(c(:))), cIndices(~vbIsColon));
   
   if (any(vnMinRange < 1) || any(vnMaxRange > oStack.vnDataSize(~vbIsColon)))
      error('TIFFStack:badsubscript', ...
         '*** TIFFStack: Index exceeds stack dimensions.');
   end
   
   % - Find unique frames to read
   [vnFrameIndices, ~, vnOrigFrameIndices] = unique(cIndices{3});
   
   % - Read data block
   try
      tfDataBlock = tiffread31_readimage(oStack.TIF, oStack.HEADER, vnFrameIndices);
      
   catch mErr
      % - Record error state
      base_ME = MException('TIFFStack:ReadError', ...
         '*** TIFFStack: Could not read data from image file.');
      new_ME = addCause(base_ME, mErr);
      throw(new_ME);
   end
      
   % - Handle linear or subscript indexing
   if (~bLinearIndexing)
      % - Select pixels from frames, if necessary
      if any(~vbIsColon([1 2 4]))
         tfData = tfDataBlock(cIndices{1}, cIndices{2}, vnOrigFrameIndices, cIndices{4});
      else
         tfData = tfDataBlock;
      end

   else
      % - Convert frame indices to frame-linear
      vnFrameLinearIndices = sub2ind(oStack.vnDataSize([1 2 4]), cIndices{1}, cIndices{2}, cIndices{4});
      
      % - Allocate return vector
      tfData = zeros(numel(cIndices{1}), 1, oStack.strDataClass);

      % - Loop over images in stack and extract required frames
      for (nFrameIndex = 1:numel(vnFrameIndices))
         vbThesePixels = cIndices{3} == vnFrameIndices(nFrameIndex);
         mfThisFrame = tfDataBlock(:, :, nFrameIndex, :);
         tfData(vbThesePixels) = mfThisFrame(vnFrameLinearIndices(vbThesePixels));
      end
   end      
      
   % - Invert data if requested
   if (oStack.bInvert)
      tfData = oStack.fhCastFun(oStack.sImageInfo(1).MaxSampleValue) - (tfData - oStack.fhCastFun(oStack.sImageInfo(1).MinSampleValue));
   end
end


% TS_read_data_Tiff - FUNCTION Read the requested pixels from the TIFF file (using tifflib)
%
% Usage: [tfData] = TS_read_data_Tiff(oStack, cIndices)
%
% 'oStack' is a TIFFStack.  'cIndices' are the indices passed in from subsref.
% Colon indexing will be converted to full range indexing.  cIndices is a cell
% array with the format {rows, cols, frames, slices}.  Slices are RGB or CMYK
% or so on.

function [tfData] = TS_read_data_Tiff(oStack, cIndices, bLinearIndexing)
   % - Convert colon indexing
   vbIsColon = cellfun(@iscolon, cIndices);
   
   for (nColonDim = find(vbIsColon))
      cIndices{nColonDim} = 1:oStack.vnDataSize(nColonDim);
   end

   % - Fix up subsample detection for unitary dimensions
   vbIsOne = cellfun(@(c)isequal(c, 1), cIndices(~vbIsColon));
   vbIsColon(~vbIsColon) = vbIsOne & (oStack.vnDataSize(~vbIsColon) == 1);
   
   % - Check ranges
   vnMinRange = cellfun(@(c)(min(c(:))), cIndices(~vbIsColon));
   vnMaxRange = cellfun(@(c)(max(c(:))), cIndices(~vbIsColon));
   
   if (any(vnMinRange < 1) || any(vnMaxRange > oStack.vnDataSize(~vbIsColon)))
      error('TIFFStack:badsubscript', ...
         '*** TIFFStack: Index exceeds stack dimensions.');
   end
   
   % - Get referencing parameters for TIF object
   w = oStack.vnDataSize(2);
   h = oStack.vnDataSize(1);
   vfrps = cellfun(@(rps)nanemptymin(rps, h), {oStack.sImageInfo(cIndices{3}).RowsPerStrip});
   vftw = cellfun(@(tw)nanemptymin(tw, w), {oStack.sImageInfo(cIndices{3}).TileWidth});
   vfth = cellfun(@(tw)nanemptymin(tw, h), {oStack.sImageInfo(cIndices{3}).TileLength});
   spp = oStack.sImageInfo(1).SamplesPerPixel;
   
   tlStack = oStack.TIF;

   % - Handle linear or subscript indexing
   if (~bLinearIndexing)
      % - Allocate single frame buffer
      vnBlockSize = oStack.vnDataSize(1:2);
      vnBlockSize(3) = numel(cIndices{3});
      vnBlockSize(4) = oStack.vnDataSize(4);
      tfImage = zeros([vnBlockSize(1:2) 1 vnBlockSize(4)], oStack.strDataClass);

      % - Allocate tensor for returning data
      vnOutputSize = cellfun(@(c)numel(c), cIndices);
      tfData = zeros(vnOutputSize, oStack.strDataClass);
      
      % - Do we need to resample the data block?
      vbTest = [true true false true];
      bResample = any(vbTest(~vbIsColon));

      try
         % - Loop over images in stack
         for (nImage = 1:numel(cIndices{3}))
            % - Skip to this image in stack
            oStack.fhSetDirFun(tlStack, cIndices{3}(nImage));
            
            % - Read data from this image, overwriting frame buffer
            [~, tfImage] = oStack.fhReadFun(tfImage, tlStack, spp, w, h, vfrps(nImage), vftw(nImage), vfth(nImage), []);
            
            % - Resample frame, if required
            if (bResample)
               tfData(:, :, nImage, :) = tfImage(cIndices{1}, cIndices{2}, cIndices{4});
            else
               tfData(:, :, nImage, :) = tfImage;
            end
         end
         
      catch mErr
         % - Record error state
         base_ME = MException('TIFFStack:ReadError', ...
            '*** TIFFStack: Could not read data from image file.');
         new_ME = addCause(base_ME, mErr);
         throw(new_ME);
      end
           
   else
      % -- Linear indexing
      
      % - Allocate return vector
      tfData = zeros(size(cIndices{1}), oStack.strDataClass);
      
      % - Allocate single-frame buffer
      vnBlockSize = oStack.vnDataSize(1:2);
      vnBlockSize(3) = numel(cIndices{3});
      vnBlockSize(4) = oStack.vnDataSize(4);
      tfImage = zeros([vnBlockSize(1:2) 1 vnBlockSize(4)], oStack.strDataClass);
      
      % - Convert frame indices to frame-linear
      vnFrameLinearIndices = sub2ind(vnBlockSize([1 2 4]), cIndices{1}, cIndices{2}, cIndices{4});
      
      % - Loop over images in stack and extract required frames
      try
         vnUniqueImages = unique(cIndices{3}(:))';
         for (nImage = 1:numel(vnUniqueImages))
            % - Find corresponding pixels
            vbThesePixels = cIndices{3} == vnUniqueImages(nImage);
            
            % - Skip to this image in stack
            oStack.fhSetDirFun(tlStack, vnUniqueImages(nImage));

            % - Read the subsampled pixels from the stack
            [tfData(vbThesePixels), tfImage] = oStack.fhReadFun(tfImage, tlStack, spp, w, h, vfrps(nImage), vftw(nImage), vfth(nImage), vnFrameLinearIndices(vbThesePixels));
         end
         
      catch mErr
         % - Record error state
         base_ME = MException('TIFFStack:ReadError', ...
            '*** TIFFStack: Could not read data from image file.');
         new_ME = addCause(base_ME, mErr);
         throw(new_ME);
      end
   end
   
   % - Invert data if requested
   if (oStack.bInvert)
      tfData = oStack.sImageInfo(1).MaxSampleValue(1) - (tfData - oStack.sImageInfo(1).MinSampleValue(1));
   end
end


% TS_read_data_MappedTensor - FUNCTION Read the requested pixels from an ImageJ fake binary TIFF file (using MappedTensor)
%
% Usage: [tfData] = TS_read_data_MappedTensor(oStack, cIndices)
%
% 'oStack' is a TIFFStack.  'cIndices' are the indices passed in from subsref.
% Colon indexing will be converted to full range indexing.  cIndices is a cell
% array with the format {rows, cols, frames, slices}.  Slices are RGB or CMYK
% or so on.

function [tfData] = TS_read_data_MappedTensor(oStack, cIndices, bLinearIndexing)

   % - Fix up subsample detection for unitary dimensions
   vbIsColon = cellfun(@iscolon, cIndices);
   vbIsOne = cellfun(@(c)isequal(c, 1), cIndices(~vbIsColon));
   vbIsColon(~vbIsColon) = vbIsOne & (oStack.vnDataSize(~vbIsColon) == 1);
   
   % - Check ranges
   vnMinRange = cellfun(@(c)(min(c(:))), cIndices(~vbIsColon));
   vnMaxRange = cellfun(@(c)(max(c(:))), cIndices(~vbIsColon));
   
   if (any(vnMinRange < 1) || any(vnMaxRange > oStack.vnDataSize(~vbIsColon)))
      error('TIFFStack:badsubscript', ...
         '*** TIFFStack: Index exceeds stack dimensions.');
   end
   
   % - Handle linear or subscript indexing
   if (~bLinearIndexing)
      
      try
         % - Just read using MappedTensor
         tfData = oStack.TIF(cIndices{1:end});
         
      catch mErr
         % - Record error state
         base_ME = MException('TIFFStack:ReadError', ...
            '*** TIFFStack: Could not read data from image file.');
         new_ME = addCause(base_ME, mErr);
         throw(new_ME);
      end
           
   else
      % -- Linear indexing
      
      % - Convert frame indices back to stack-linear
      vnStackLinearIndices = sub2ind(oStack.vnDataSize, cIndices{:});
      
      % - Loop over images in stack and extract required frames
      try
         % - Just read using MappedTensor
         tfData = oStack.TIF(vnStackLinearIndices);
         
      catch mErr
         % - Record error state
         base_ME = MException('TIFFStack:ReadError', ...
            '*** TIFFStack: Could not read data from image file.');
         new_ME = addCause(base_ME, mErr);
         throw(new_ME);
      end
   end
   
   % - Invert data if requested
   if (oStack.bInvert)
      tfData = oStack.sImageInfo(1).MaxSampleValue(1) - (tfData - oStack.sImageInfo(1).MinSampleValue(1));
   end
end


% GetLinearIndicesForRefs - FUNCTION Convert a set of multi-dimensional indices directly into linear indices
function [vnLinearIndices, vnDimRefSizes] = GetLinearIndicesForRefs(cRefs, vnLims, hRepSumFunc)

   % - Find colon references
   vbIsColon = cellfun(@iscolon, cRefs);
   
   if (all(vbIsColon))
      vnLinearIndices = 1:prod(vnLims);
      vnDimRefSizes = vnLims;
      return;
   end
   
   nFirstNonColon = find(~vbIsColon, 1, 'first');
   vbTrailingRefs = true(size(vbIsColon));
   vbTrailingRefs(1:nFirstNonColon-1) = false;
   vnDimRefSizes = cellfun(@numel, cRefs);
   vnDimRefSizes(vbIsColon) = vnLims(vbIsColon);
   
   % - Calculate dimension offsets
   vnDimOffsets = [1 cumprod(vnLims)];
   vnDimOffsets = vnDimOffsets(1:end-1);

   % - Remove trailing "1"s
   vbOnes = cellfun(@(c)isequal(c, 1), cRefs);
   nLastNonOne = find(~vbOnes, 1, 'last');
   vbTrailingRefs((nLastNonOne+1):end) = false;

   % - Check reference limits
   if (any(cellfun(@(r,l)any(r>l), cRefs(~vbIsColon), num2cell(vnLims(~vbIsColon)))))
      error('TIFFStack:badsubscript', 'Index exceeds matrix dimensions.');
   end
   
   % - Work out how many linear indices there will be in total
   nNumIndices = prod(vnDimRefSizes);
   vnLinearIndices = zeros(nNumIndices, 1);
   
   % - Build a referencing window encompassing the leading colon refs (or first ref)
   if (nFirstNonColon > 1)
      vnLinearIndices(1:prod(vnLims(1:(nFirstNonColon-1)))) = 1:prod(vnLims(1:(nFirstNonColon-1)));
   else
      vnLinearIndices(1:vnDimRefSizes(1)) = cRefs{1};
      vbTrailingRefs(1) = false;
   end
   
   % - Replicate windows to make up linear indices
   for (nDimension = find(vbTrailingRefs & ~vbOnes))
      % - How long is the current window?
      nCurrWindowLength = prod(vnDimRefSizes(1:(nDimension-1)));
      nThisWindowLength = nCurrWindowLength * vnDimRefSizes(nDimension);
      
      % - Is this dimension a colon reference?
      if (vbIsColon(nDimension))
         vnLinearIndices(1:nThisWindowLength) = hRepSumFunc(vnLinearIndices(1:nCurrWindowLength), ((1:vnLims(nDimension))-1) * vnDimOffsets(nDimension));

      else
         vnLinearIndices(1:nThisWindowLength) = hRepSumFunc(vnLinearIndices(1:nCurrWindowLength), (cRefs{nDimension}-1) * vnDimOffsets(nDimension));
      end
   end
end

% mapped_tensor_repsum_nomex - FUNCTION Slow version of replicate and sum
function [vfDest] = mapped_tensor_repsum_nomex(vfSourceA, vfSourceB)
   [mfA, mfB] = meshgrid(vfSourceB, vfSourceA);
   vfDest = mfA(:) + mfB(:);
end

% isvalidsubscript - FUNCTION Test whether a vector contains valid entries
% for subscript referencing
function isvalidsubscript(oRefs)
   try
      % - Test for colon
      if (iscolon(oRefs))
         return;
      end
      
      if (islogical(oRefs))
         % - Test for logical indexing
         validateattributes(oRefs, {'logical'}, {'binary'});
         
      else
         % - Test for normal indexing
         validateattributes(oRefs, {'single', 'double'}, {'integer', 'real', 'positive'});
      end
      
   catch
      error('TIFFStack:badsubscript', ...
            '*** TIFFStack: Subscript indices must either be real positive integers or logicals.');
   end
end

%% get_full_file_path - FUNCTION Calculate the absolute path to a given (possibly relative) filename
%
% Usage: strFullPath = get_full_file_path(strFile)
%
% 'strFile' is a filename, which may include relative path elements.  The file
% does not have to exist.
%
% 'strFullPath' will be the absolute path to the file indicated in 'strFile'.

function strFullPath = get_full_file_path(strFile)

   try
      fid = fopen(strFile);
      strFile = fopen(fid);
      fclose(fid);
      
      [strDir, strName, strExt] = fileparts(strFile);
      
      if (isempty(strDir))
         strDir = '.';
         strFullDirPath = cd(cd(strDir));
         strFullPath = fullfile(strFullDirPath, [strName strExt]);
      else
         strFullPath = strFile;
      end
      
   catch mErr
      % - Close file id, if necessary
      if (ismember(fid, fopen('all')))
         fclose(fid);
      end
      
      % - Record error state
      base_ME = MException('TIFFStack:ReadError', ...
         '*** TIFFStack: Could not open file [%s].', strFile);
      new_ME = addCause(base_ME, mErr);
      throw(new_ME);
   end
end

% iscolon - FUNCTION Test whether a reference is equal to ':'
function bIsColon = iscolon(ref)
   bIsColon = ischar(ref) && isequal(ref, ':');
end

% nanemptymin - FUNCTION Test for a min or nan or empty
function fVal = nanemptymin(fVal, fTestVal)
   fVal(isempty(fVal)) = fTestVal;
   fVal(isnan(fVal)) = fTestVal;
end

%% Accelerated Libtiff reading functions

% TS_read_Tiff_striped_separate - FUNCTION Read an image using tifflib, for
% striped separate TIFF files
function [vfOutputPixels, tfImageBuffer] = TS_read_Tiff_striped_separate(tfImageBuffer, tlStack, spp, ~, h, rps, ~, ~, vnFrameLinearIndices) %#ok<DEFNU>
   for r = 1:rps:h
      row_inds = r:min(h,r+rps-1);
      for k = 1:spp
         stripNum = tifflib('computeStrip', tlStack, r-1, k-1);
         tfImageBuffer(row_inds,:,k) = tifflib('readEncodedStrip', tlStack, stripNum-1);
      end
   end
   
   % - Perform sub-referencing, if required
   if (~isempty(vnFrameLinearIndices))
      vfOutputPixels = tfImageBuffer(vnFrameLinearIndices);
   else
      vfOutputPixels = tfImageBuffer;
   end
end

% TS_read_Tiff_striped_chunky - FUNCTION Read an image using tifflib, for
% striped chunk TIFF files
function [vfOutputPixels, tfImageBuffer] = TS_read_Tiff_striped_chunky(tfImageBuffer, tlStack, ~, ~, h, rps, ~, ~, vnFrameLinearIndices) %#ok<DEFNU>
   for r = 1:rps:h
      row_inds = r:min(h,r+rps-1);
      stripNum = tifflib('computeStrip', tlStack, r-1);
      tfImageBuffer(row_inds,:,:) = tifflib('readEncodedStrip', tlStack, stripNum-1);
   end
   
   % - Perform sub-referencing, if required
   if (~isempty(vnFrameLinearIndices))
      vfOutputPixels = tfImageBuffer(vnFrameLinearIndices);
   else
      vfOutputPixels = tfImageBuffer;
   end
end

% TS_read_Tiff_tiled_separate - FUNCTION Read an image using tifflib, for
% tiled separate TIFF files
function [vfOutputPixels, tfImageBuffer] = TS_read_Tiff_tiled_separate(tfImageBuffer, tlStack, spp, w, h, ~, tWidth, tHeight, vnFrameLinearIndices) %#ok<DEFNU>
   for r = 1:tHeight:h
      row_inds = r:min(h,r+tHeight-1);
      for c = 1:tWidth:w
         col_inds = c:min(w,c+tWidth-1);
         for k = 1:spp
            tileNumber = tifflib('computeTile', tlStack, [r c]-1, k);
            tfImageBuffer(row_inds,col_inds,k) = tifflib('readEncodedTile', tlStack, tileNumber-1);
         end
      end
   end
   
   % - Perform sub-referencing, if required
   if (~isempty(vnFrameLinearIndices))
      vfOutputPixels = tfImageBuffer(vnFrameLinearIndices);
   else
      vfOutputPixels = tfImageBuffer;
   end
end

% TS_read_Tiff_tiled_chunky - FUNCTION Read an image using tifflib, for
% tiled chunky TIFF files
function [vfOutputPixels, tfImageBuffer] = TS_read_Tiff_tiled_chunky(tfImageBuffer, tlStack, ~, w, h, ~, tWidth, tHeight, vnFrameLinearIndices) %#ok<DEFNU>
   for r = 1:tHeight:h
      row_inds = r:min(h,r+tHeight-1);
      for c = 1:tWidth:w
         col_inds = c:min(w,c+tWidth-1);
         tileNumber = tifflib('computeTile', tlStack, [r c]-1);
         tfImageBuffer(row_inds,col_inds,:) = tifflib('readEncodedTile', tlStack, tileNumber-1);
      end
   end
   
   % - Perform sub-referencing, if required
   if (~isempty(vnFrameLinearIndices))
      vfOutputPixels = tfImageBuffer(vnFrameLinearIndices);
   else
      vfOutputPixels = tfImageBuffer;
   end
end

% TS_set_direcotry - FUNCTION Set the current TIFF directory
function TS_set_directory(tlStack, nDirectory) %#ok<DEFNU>
   tifflib('setDirectory', tlStack, nDirectory-1);
end

%% Pre-2014 matlab Tifflib reading functions

% TS_read_Tiff_striped_separate - FUNCTION Read an image using tifflib, for
% striped separate TIFF files
function [vfOutputPixels, tfImageBuffer] = TS_read_Tiff_striped_separate_pre2014(tfImageBuffer, tlStack, spp, ~, h, rps, ~, ~, vnFrameLinearIndices) %#ok<DEFNU>
   for r = 1:rps:h
      row_inds = r:min(h,r+rps-1);
      for k = 1:spp
         stripNum = tifflib('computeStrip', tlStack, r, k);
         tfImageBuffer(row_inds,:,k) = tifflib('readEncodedStrip', tlStack, stripNum);
      end
   end
   
   % - Perform sub-referencing, if required
   if (~isempty(vnFrameLinearIndices))
      vfOutputPixels = tfImageBuffer(vnFrameLinearIndices);
   else
      vfOutputPixels = tfImageBuffer;
   end
end

% TS_read_Tiff_striped_chunky - FUNCTION Read an image using tifflib, for
% striped chunk TIFF files
function [vfOutputPixels, tfImageBuffer] = TS_read_Tiff_striped_chunky_pre2014(tfImageBuffer, tlStack, ~, ~, h, rps, ~, ~, vnFrameLinearIndices) %#ok<DEFNU>
   for r = 1:rps:h
      row_inds = r:min(h,r+rps-1);
      stripNum = tifflib('computeStrip', tlStack, r);
      tfImageBuffer(row_inds,:,:) = tifflib('readEncodedStrip', tlStack, stripNum);
   end
   
   % - Perform sub-referencing, if required
   if (~isempty(vnFrameLinearIndices))
      vfOutputPixels = tfImageBuffer(vnFrameLinearIndices);
   else
      vfOutputPixels = tfImageBuffer;
   end
end

% TS_read_Tiff_tiled_separate - FUNCTION Read an image using tifflib, for
% tiled separate TIFF files
function [vfOutputPixels, tfImageBuffer] = TS_read_Tiff_tiled_separate_pre2014(tfImageBuffer, tlStack, spp, w, h, ~, tWidth, tHeight, vnFrameLinearIndices) %#ok<DEFNU>
   for r = 1:tHeight:h
      row_inds = r:min(h,r+tHeight-1);
      for c = 1:tWidth:w
         col_inds = c:min(w,c+tWidth-1);
         for k = 1:spp
            tileNumber = tifflib('computeTile', tlStack, [r c], k);
            tfImageBuffer(row_inds,col_inds,k) = tifflib('readEncodedTile', tlStack, tileNumber);
         end
      end
   end
   
   % - Perform sub-referencing, if required
   if (~isempty(vnFrameLinearIndices))
      vfOutputPixels = tfImageBuffer(vnFrameLinearIndices);
   else
      vfOutputPixels = tfImageBuffer;
   end
end

% TS_read_Tiff_tiled_chunky - FUNCTION Read an image using tifflib, for
% tiled chunky TIFF files
function [vfOutputPixels, tfImageBuffer] = TS_read_Tiff_tiled_chunky_pre2014(tfImageBuffer, tlStack, ~, w, h, ~, tWidth, tHeight, vnFrameLinearIndices) %#ok<DEFNU>
   for r = 1:tHeight:h
      row_inds = r:min(h,r+tHeight-1);
      for c = 1:tWidth:w
         col_inds = c:min(w,c+tWidth-1);
         tileNumber = tifflib('computeTile', tlStack, [r c]);
         tfImageBuffer(row_inds,col_inds,:) = tifflib('readEncodedTile', tlStack, tileNumber);
      end
   end
   
   % - Perform sub-referencing, if required
   if (~isempty(vnFrameLinearIndices))
      vfOutputPixels = tfImageBuffer(vnFrameLinearIndices);
   else
      vfOutputPixels = tfImageBuffer;
   end
end

% TS_set_direcotry - FUNCTION Set the current TIFF directory
function TS_set_directory_pre2014(tlStack, nDirectory) %#ok<DEFNU>
   tifflib('setDirectory', tlStack, nDirectory);
end


%% Accelerated TiffgetTag function

function tagValue = TiffgetTag(oTiff,tagId)
% getTag  Retrieve tag from image.
%   tagValue = getTag(tagId) retrieves the value of the tag tagId
%   from the current directory.  tagId may be specified either via
%   the Tiff.TagID property or as a char string.
%
%   This method corresponds to the TIFFGetField function in the
%   LibTIFF C API.  To use this method, you must be familiar with
%   LibTIFF version 3.7.1 as well as the TIFF specification and
%   technical notes.  This documentation may be referenced at
%   <http://www.remotesensing.org/libtiff/document.html>.
%
%   Example:
%
%   t = Tiff('example.tif','r');
%   % Specify tag by tag number.
%   width = t.getTag(Tiff.TagID.ImageWidth);
%
%   % Specify tag by tag name.
%   width = t.getTag('ImageWidth');
%
%   See also setTag
%
%

   switch(class(tagId))
      case 'char'
         % The user gave a char id for the tag.
         tagValue = tifflib('getField',oTiff,Tiff.TagID.(tagId));
         
      otherwise
         % Assume numeric.
         tagValue = tifflib('getField',oTiff,tagId);
   end
end


%% -- MEX-handling functions

function [hRepSumFunc] = GetMexFunctionHandles
   % - Try to use cache for function handle
   persistent GMFH_hRepSumFunc;
   
   if ~isempty(GMFH_hRepSumFunc)
      hRepSumFunc = GMFH_hRepSumFunc;
      return;
   end

   % - Does the compiled MEX function exist?
   if (exist('mapped_tensor_repsum') ~= 3) %#ok<EXIST>
      % - Move to the MappedTensor private directory
      strMTDir = fileparts(which('TIFFStack'));
      strCWD = cd(fullfile(strMTDir, 'private'));
         
      try %#ok<TRYNC>
         % - Try to compile the MEX functions
         disp('--- TIFFStack: Compiling MEX functions.');
         mex('mapped_tensor_repsum.c', '-largeArrayDims', '-O');
      end
      
      % - Move back to previous working directory
      cd(strCWD);
   end
   
   % - Did we succeed?
   if (exist('mapped_tensor_repsum') == 3) %#ok<EXIST>
      hRepSumFunc = @mapped_tensor_repsum;
      
   else
      % - Just use the slow matlab version
      warning('TIFFStack:MEXCompilation', ...
         '--- TIFFStack: Could not compile MEX functions.  Using slow matlab versions.');
      
      hRepSumFunc = @mapped_tensor_repsum_nomex;
   end
   
   % - Record function handle in the cache
   GMFH_hRepSumFunc = hRepSumFunc;
end

%% -- ImageJ helper functions

function [bIsImageJBigStack, bIsImageJHyperStack, vnStackDims, vnInterleavedFrameDims] = IsImageJBigStack(sInfo, nApparentSize)

   % - Set up default return arguments
   bIsImageJBigStack = false;
   bIsImageJHyperStack = false;
   vnStackDims = [];
   vnInterleavedFrameDims = [];
   
   % - Check for ImageDescription field
   if (~isfield(sInfo, 'ImageDescription'))
      return;
   end
   
   % - Get image description
   strImageDesc = sInfo(1).ImageDescription;
   
   % - Look for ImageJ version information
   strImageJVer = sscanf(strImageDesc(strfind(strImageDesc, 'ImageJ='):end), 'ImageJ=%s');
   
   % - Look for stack size information
   if (~isempty(strImageJVer))
      nNumImages = sscanf(strImageDesc(strfind(strImageDesc, 'images='):end), 'images=%d');
      
      % - Does ImageJ report a greater number of images than sInfo?
      if (~isempty(nNumImages) && (nApparentSize ~= nNumImages))
         bIsImageJBigStack = true;
      end
      
      % - Is this a hyperstack?
      strHyperStack = sscanf(strImageDesc(strfind(strImageDesc, 'hyperstack='):end), 'hyperstack=%s');
      
      if (strcmpi(strHyperStack, 'true'))
         bIsImageJHyperStack = true;
         
         % - Extract information about the stack size for a hyperstack
         nNumChannels = sscanf(strImageDesc(strfind(strImageDesc, 'channels='):end), 'channels=%d');
         nNumSlices = sscanf(strImageDesc(strfind(strImageDesc, 'slices='):end), 'slices=%d');
         nNumFrames = sscanf(strImageDesc(strfind(strImageDesc, 'frames='):end), 'frames=%d');
         
         if (isempty(nNumChannels))
            nNumChannels = 1;
         end
         
         if (isempty(nNumSlices))
            nNumSlices = 1;
         end
         
         if (isempty(nNumFrames))
            nNumFrames = 1;
         end
         
         % - Check total stack size
         if (nNumFrames*nNumSlices*nNumChannels ~= nApparentSize)
            warning('TIFFStack:ImageJStackSize', ...
               'The reported size of this stack does not match the number of IFDs.\nI will attempt to continue.');
            
            nNumFrames = floor(nApparentSize / nNumSlices / nNumChannels);
         end
         
         % - Deinterleave stack
         vnStackDims = [sInfo(1).Width sInfo(1).Height nNumFrames*nNumSlices*nNumChannels 1];
         vnInterleavedFrameDims = [nNumChannels nNumSlices nNumFrames];
         
      else
         % - Extract information about the stack size for a fake big stack
         vnStackDims = [sInfo(1).Width sInfo(1).Height nNumImages sInfo(1).SamplesPerPixel];
         vnInterleavedFrameDims = [];
      end
   end
end

function [mtHandle, bUseMappedTensor, strDataClass] = OpenImageJBigStack(oStack, vnStackDims)
   bUseMappedTensor = (exist('MappedTensor', 'class') == 8);
   mtHandle = [];
   strDataClass = [];
   
   if (bUseMappedTensor)
      % - Use tiffread to get file information
      [TIF, HEADER] = tiffread31_header(oStack.strFilename);
      
      % - Use MappedTensor to open the file
      strDataClass = TIF.classname;
      mtHandle = MappedTensor(oStack.strFilename, vnStackDims, ...
         'MachineFormat', TIF.ByteOrder, 'Class', strDataClass, 'HeaderBytes', HEADER.StripOffsets);
   end
end

% --- END of TIFFStack.m ---
```
