## Supplementary material for "Enhancer architecture and chromatin accessibility constrain phenotypic space during development": Script-MLP: bfCheckJavaMemory.m

```
function [] = bfCheckJavaMemory(varargin)
% bfCheckJavaMemory warn if too little memory is allocated to Java
%
% SYNOPSIS  bfCheckJavaMemory()
%
% Input
%
%   minMemory - (Optional) The minimum suggested memory setting in MB.
%   Default: 512
%
% Output
%
%    A warning message is printed if too little memory is allocated.

% OME Bio-Formats package for reading and converting biological file formats.
%
% Copyright (C) 2014 - 2017 Open Microscopy Environment:
%   - Board of Regents of the University of Wisconsin-Madison
%   - Glencoe Software, Inc.
%   - University of Dundee
%
% This program is free software: you can redistribute it and/or modify
% it under the terms of the GNU General Public License as
% published by the Free Software Foundation, either version 2 of the
% License, or (at your option) any later version.
%
% This program is distributed in the hope that it will be useful,
% but WITHOUT ANY WARRANTY; without even the implied warranty of
% MERCHANTABILITY or FITNESS FOR A PARTICULAR PURPOSE.  See the
% GNU General Public License for more details.
%
% You should have received a copy of the GNU General Public License along
% with this program; if not, write to the Free Software Foundation, Inc.,
% 51 Franklin Street, Fifth Floor, Boston, MA  02110-1301, USA.

runtime = javaMethod('getRuntime', 'java.lang.Runtime');
maxMemory = runtime.maxMemory() / (1024 * 1024);

ip = inputParser;
ip.addOptional('minMemory', 512, @isscalar);
ip.parse(varargin{:});
minMemory = ip.Results.minMemory;

warningID = 'BF:lowJavaMemory';

if maxMemory < minMemory - 64
    warning_msg = [...
        '*** Insufficient memory detected. ***\n'...
        '*** %dm found ***\n'...
        '*** %dm or greater is recommended ***\n'...
        '*** See http://www.mathworks.com/matlabcentral/answers/92813 ***\n'...
        '*** for instructions on increasing memory allocation. ***\n'];
    warning(warningID, warning_msg, round(maxMemory), minMemory);
end
```
