## Supplementary material for "Enhancer architecture and chromatin accessibility constrain phenotypic space during development": Script-MLP: bfCheckJavaPath.m

```
function [status, version] = bfCheckJavaPath(varargin)
% bfCheckJavaPath check Bio-Formats is included in the Java class path
%
% SYNOPSIS  bfCheckJavaPath()
%           status = bfCheckJavaPath(autoloadBioFormats)
%           [status, version] = bfCheckJavaPath()
%
% Input
%
%    autoloadBioFormats - Optional. A boolean specifying the action
%    to take if Bio-Formats is not in the javaclasspath.  If true,
%    tries to find a Bio-Formats jar file and adds it to the java
%    class path.
%    Default - true
%
% Output
%
%    status - Boolean. True if the Bio-Formats classes are in the Java
%    class path.
%
%
%    version - String specifying the current version of Bio-Formats if
%    Bio-Formats is in the Java class path. Empty string otherwise.

% OME Bio-Formats package for reading and converting biological file formats.
%
% Copyright (C) 2012 - 2017 Open Microscopy Environment:
%   - Board of Regents of the University of Wisconsin-Madison
%   - Glencoe Software, Inc.
%   - University of Dundee
%
% This program is free software: you can redistribute it and/or modify
% it under the terms of the GNU General Public License as
% published by the Free Software Foundation, either version 2 of the
% License, or (at your option) any later version.
%
% This program is distributed in the hope that it will be useful,
% but WITHOUT ANY WARRANTY; without even the implied warranty of
% MERCHANTABILITY or FITNESS FOR A PARTICULAR PURPOSE.  See the
% GNU General Public License for more details.
%
% You should have received a copy of the GNU General Public License along
% with this program; if not, write to the Free Software Foundation, Inc.,
% 51 Franklin Street, Fifth Floor, Boston, MA  02110-1301, USA.

% Input check
ip = inputParser;
ip.addOptional('autoloadBioFormats', true, @isscalar);
ip.parse(varargin{:});

[status, version] = has_working_bioformats();

if ~status && ip.Results.autoloadBioFormats,
    jarPath = fullfile(fileparts(mfilename('fullpath')), ...
                       'bioformats_package.jar');
    if (exist(jarPath) == 2)
        javaaddpath(jarPath);
        [status, version] = has_working_bioformats();
        if (~status)
            javarmpath(jarPath);
        end
    end
end

% Return true if bioformats java interface is working, false otherwise.
% Not working will probably mean that the classes are not in
% the javaclasspath.
function [status, version] = has_working_bioformats()

status = true;
version = '';
try
    % If the following fails for any reason, then bioformats is not working.
    % Getting the version number and creating a reader is the bare minimum.
    reader = javaObject('loci.formats.in.FakeReader');
    if is_octave()
        version = char(java_get('loci.formats.FormatTools', 'VERSION'));
    else
        version = char(loci.formats.FormatTools.VERSION);
    end
catch
    status = false;
end
```
