## Supplementary material for "Enhancer architecture and chromatin accessibility constrain phenotypic space during development": Script-MLP: bfGetFileExtensions.m

```
function fileExt = bfGetFileExtensions
% bfGetFileExtensions list all extensions supported by Bio-Formats
%
% Synopsis: fileExt = bfGetExtensions()
%
% Input
%      none
%
% Output
%      fileExt:  a cell array of dimensions n x2 where the first colum
%      gives the extension and the second the name of the corresponding
%      format.
%      This cell array is formatted to be used with uigetfile funciton.

% OME Bio-Formats package for reading and converting biological file formats.
%
% Copyright (C) 2012 - 2017 Open Microscopy Environment:
%   - Board of Regents of the University of Wisconsin-Madison
%   - Glencoe Software, Inc.
%   - University of Dundee
%
% This program is free software: you can redistribute it and/or modify
% it under the terms of the GNU General Public License as
% published by the Free Software Foundation, either version 2 of the
% License, or (at your option) any later version.
%
% This program is distributed in the hope that it will be useful,
% but WITHOUT ANY WARRANTY; without even the implied warranty of
% MERCHANTABILITY or FITNESS FOR A PARTICULAR PURPOSE.  See the
% GNU General Public License for more details.
%
% You should have received a copy of the GNU General Public License along
% with this program; if not, write to the Free Software Foundation, Inc.,
% 51 Franklin Street, Fifth Floor, Boston, MA  02110-1301, USA.

% Get all readers and create cell array with suffixes and names
readers = javaMethod('getReaders', javaObject('loci.formats.ImageReader'));
fileExt = cell(numel(readers), 2);
for i = 1:numel(readers)
    suffixes = readers(i).getSuffixes();
    if is_octave()
        %% FIXME when https://savannah.gnu.org/bugs/?42700 gets fixed
        ExtSuf = cell(numel(suffixes), 1);
        for j = 1:numel(suffixes)
            ExtSuf{j} = char(suffixes(j));
        end
        fileExt{i, 1} = ExtSuf;
    else
        fileExt{i, 1} = arrayfun(@char, suffixes, 'Unif', false);
    end
    fileExt{i, 2} = char(readers(i).getFormat());
end

% Concatenate all unique formats
allExt = unique(vertcat(fileExt{:, 1}));
allExt = allExt(~cellfun(@isempty, allExt));
fileExt = vertcat({allExt, 'All formats'}, fileExt);

% Format file extensions
for i = 1:size(fileExt, 1)
    fileExt{i, 1} = sprintf('*.%s;', fileExt{i, 1}{:});
    fileExt{i, 1}(end) = [];
end
```
