## Supplementary material for "Enhancer architecture and chromatin accessibility constrain phenotypic space during development": Script-MLP: bfGetPlane.m

```
function I = bfGetPlane(r, varargin)
% BFGETPLANE Retrieve the plane data from a reader using Bio-Formats
%
%   I = bfGetPlane(r, iPlane) returns a specified plane from the input
%   format reader. The index specifying the plane to retrieve should be
%   contained between 1 and the number of planes for the series. Given a
%   set of (z, c, t) plane coordinates, the plane index (0-based) can be
%   retrieved using r.getIndex(z, c, t).
%
%   I = bfGetPlane(r, iPlane, x, y, width, height) only returns the tile
%   which origin is specified by (x, y) and dimensions are specified by
%   (width, height).
%
% Examples
%
%    I = bfGetPlane(r, 1) % First plane of the series
%    I = bfGetPlane(r, r.getImageCount()) % Last plane of the series
%    I = bfGetPlane(r, r.getIndex(0, 0, 0) + 1) % First plane of the series
%    I = bfGetPlane(r, 1, 1, 1, 20, 20) % 20x20 tile originated at (0, 0)
%
% See also: BFGETREADER

% OME Bio-Formats package for reading and converting biological file formats.
%
% Copyright (C) 2012 - 2017 Open Microscopy Environment:
%   - Board of Regents of the University of Wisconsin-Madison
%   - Glencoe Software, Inc.
%   - University of Dundee
%
% This program is free software: you can redistribute it and/or modify
% it under the terms of the GNU General Public License as
% published by the Free Software Foundation, either version 2 of the
% License, or (at your option) any later version.
%
% This program is distributed in the hope that it will be useful,
% but WITHOUT ANY WARRANTY; without even the implied warranty of
% MERCHANTABILITY or FITNESS FOR A PARTICULAR PURPOSE.  See the
% GNU General Public License for more details.
%
% You should have received a copy of the GNU General Public License along
% with this program; if not, write to the Free Software Foundation, Inc.,
% 51 Franklin Street, Fifth Floor, Boston, MA  02110-1301, USA.

% Input check
ip = inputParser;
isValidReader = @(x) isa(x, 'loci.formats.IFormatReader') && ...
    ~isempty(x.getCurrentFile());
ip.addRequired('r', isValidReader);
ip.parse(r);

% Plane check
isValidPlane = @(x) isscalar(x) && ismember(x, 1 : r.getImageCount());
% Optional tile arguments check
isValidX = @(x) isscalar(x) && ismember(x, 1 : r.getSizeX());
isValidY = @(x) isscalar(x) && ismember(x, 1 : r.getSizeY());
ip.addRequired('iPlane', isValidPlane);
ip.addOptional('x', 1, isValidX);
ip.addOptional('y', 1, isValidY);
ip.addOptional('width', r.getSizeX(), isValidX);
ip.addOptional('height', r.getSizeY(), isValidY);
ip.parse(r, varargin{:});

% Additional check for tile size
assert(ip.Results.x - 1 + ip.Results.width <= r.getSizeX(),...
     'MATLAB:InputParser:ArgumentFailedValidation',...
     'Invalid tile size');
assert(ip.Results.y - 1 + ip.Results.height <= r.getSizeY(),...
     'MATLAB:InputParser:ArgumentFailedValidation',...
     'Invalid tile size');

% Get pixel type
pixelType = r.getPixelType();
bpp = javaMethod('getBytesPerPixel', 'loci.formats.FormatTools', pixelType);
fp = javaMethod('isFloatingPoint', 'loci.formats.FormatTools', pixelType);
sgn = javaMethod('isSigned', 'loci.formats.FormatTools', pixelType);
little = r.isLittleEndian();

plane = r.openBytes(...
    ip.Results.iPlane - 1, ip.Results.x - 1, ip.Results.y - 1, ...
    ip.Results.width, ip.Results.height);

% convert byte array to MATLAB image
if sgn
    % can get the data directly to a matrix
    I = javaMethod('makeDataArray2D', 'loci.common.DataTools', plane, ...
        bpp, fp, little, ip.Results.height);
else
    % get the data as a vector, either because makeDataArray2D
    % is not available, or we need a vector for typecast
    I = javaMethod('makeDataArray', 'loci.common.DataTools', plane, ...
        bpp, fp, little);
end

% Java does not have explicitly unsigned data types;
% hence, we must inform MATLAB when the data is unsigned
if ~sgn
    % NB: arr will always be a vector here
    switch class(I)
        case 'int8'
            I = typecast(I, 'uint8');
        case 'int16'
            I = typecast(I, 'uint16');
        case 'int32'
            I = typecast(I, 'uint32');
        case 'int64'
            I = typecast(I, 'uint64');
    end
end

if isvector(I)
    % convert results from vector to matrix
    shape = [ip.Results.width ip.Results.height];
    I = reshape(I, shape)';
end
```
