## Supplementary material for "Enhancer architecture and chromatin accessibility constrain phenotypic space during development": Script-MLP: bfGetReader.m

```
function r = bfGetReader(varargin)
% BFGETREADER return a reader for a microscopy image using Bio-Formats
%
%   r = bfGetReader() creates an empty Bio-Formats reader extending
%   loci.formats.ReaderWrapper.
%
%   r = bfGetReader(id) where id is a path to an existing file creates and
%   initializes a reader for the input file.
%
% Examples
%
%    r = bfGetReader()
%    I = bfGetReader(path_to_file)
%
%
% See also: BFGETPLANE

% OME Bio-Formats package for reading and converting biological file formats.
%
% Copyright (C) 2012 - 2017 Open Microscopy Environment:
%   - Board of Regents of the University of Wisconsin-Madison
%   - Glencoe Software, Inc.
%   - University of Dundee
%
% This program is free software: you can redistribute it and/or modify
% it under the terms of the GNU General Public License as
% published by the Free Software Foundation, either version 2 of the
% License, or (at your option) any later version.
%
% This program is distributed in the hope that it will be useful,
% but WITHOUT ANY WARRANTY; without even the implied warranty of
% MERCHANTABILITY or FITNESS FOR A PARTICULAR PURPOSE.  See the
% GNU General Public License for more details.
%
% You should have received a copy of the GNU General Public License along
% with this program; if not, write to the Free Software Foundation, Inc.,
% 51 Franklin Street, Fifth Floor, Boston, MA  02110-1301, USA.

% Input check
ip = inputParser;
ip.addOptional('id', '', @ischar);
ip.addOptional('stitchFiles', false, @isscalar);
ip.parse(varargin{:});
id = ip.Results.id;

% verify that enough memory is allocated
bfCheckJavaMemory();

% load the Bio-Formats library into the MATLAB environment
status = bfCheckJavaPath();
assert(status, ['Missing Bio-Formats library. Either add bioformats_package.jar '...
    'to the static Java path or add it to the Matlab path.']);

% Check if input is a fake string
isFake = strcmp(id(max(1, end - 4):end), '.fake');

if ~isempty(id) && ~isFake
    % Check file existence using fileattrib
    [status, f] = fileattrib(id);
    assert(status && f.directory == 0, 'bfGetReader:FileNotFound',...
        'No such file: %s', id);
    id = f.Name;
end

% set LuraWave license code, if available
if exist('lurawaveLicense', 'var')
    path = fullfile(fileparts(mfilename('fullpath')), 'lwf_jsdk2.6.jar');
    javaaddpath(path);
    javaMethod('setProperty', 'java.lang.System', ...
               'lurawave.license', lurawaveLicense);
end

% Create a loci.formats.ReaderWrapper object
r = javaObject('loci.formats.ChannelSeparator', ...
               javaObject('loci.formats.ChannelFiller'));
if ip.Results.stitchFiles
    r = javaObject('loci.formats.FileStitcher', r);
end

% Initialize the metadata store
OMEXMLService = javaObject('loci.formats.services.OMEXMLServiceImpl');
r.setMetadataStore(OMEXMLService.createOMEXMLMetadata());

% Initialize the reader
if ~isempty(id), evalc('r.setId(id)'); end
```
