## Supplementary material for "Enhancer architecture and chromatin accessibility constrain phenotypic space during development": Script-MLP: bfInitLogging.m

```
function bfInitLogging(varargin)
% BFINITLOGGING initializes the Bio-Formats logging system
%
%   bfInitLogging() initializes the logging system at WARN level by default.
%
%   bfInitLogging(level) sets the logging level to use when initializing
%   the logging system
%
% Examples
%
%    bfInitLogging();
%    bfInitLogging('DEBUG');

% OME Bio-Formats package for reading and converting biological file formats.
%
% Copyright (C) 2016 - 2017 Open Microscopy Environment:
%   - Board of Regents of the University of Wisconsin-Madison
%   - Glencoe Software, Inc.
%   - University of Dundee
%
% This program is free software: you can redistribute it and/or modify
% it under the terms of the GNU General Public License as
% published by the Free Software Foundation, either version 2 of the
% License, or (at your option) any later version.
%
% This program is distributed in the hope that it will be useful,
% but WITHOUT ANY WARRANTY; without even the implied warranty of
% MERCHANTABILITY or FITNESS FOR A PARTICULAR PURPOSE.  See the
% GNU General Public License for more details.
%
% You should have received a copy of the GNU General Public License along
% with this program; if not, write to the Free Software Foundation, Inc.,
% 51 Franklin Street, Fifth Floor, Boston, MA  02110-1301, USA.

% Check Bio-Formats is set in the Java class path
bfCheckJavaPath();

% Input check
levels = {'ALL', 'DEBUG', 'ERROR', 'FATAL', 'INFO', 'OFF', 'TRACE', 'WARN'};
ip = inputParser;
ip.addOptional('level', 'WARN', @(x) ismember(x, levels));
ip.parse(varargin{:});

% Set logging level
javaMethod('enableLogging', 'loci.common.DebugTools', ip.Results.level);
```
