## Supplementary material for "Enhancer architecture and chromatin accessibility constrain phenotypic space during development": Script-MLP: bfOpen3DVolume.m

```
function volume = bfOpen3DVolume(filename)
% bfOpen3DVolume loads a stack of images using Bio-Formats and transforms them
% into a 3D volume
%
% SYNPOSIS  bfOpen3DVolume
%           V = bfOpen3DVolume(filename)
%
% Input
%
%   filename - Optional.  A path to the file to be opened.  If not specified,
%   then a file chooser window will appear.
%
% Output
%
%   volume - 3D array containing all images in the file.

% OME Bio-Formats package for reading and converting biological file formats.
%
% Copyright (C) 2012 - 2017 Open Microscopy Environment:
%   - Board of Regents of the University of Wisconsin-Madison
%   - Glencoe Software, Inc.
%   - University of Dundee
%
% This program is free software: you can redistribute it and/or modify
% it under the terms of the GNU General Public License as
% published by the Free Software Foundation, either version 2 of the
% License, or (at your option) any later version.
%
% This program is distributed in the hope that it will be useful,
% but WITHOUT ANY WARRANTY; without even the implied warranty of
% MERCHANTABILITY or FITNESS FOR A PARTICULAR PURPOSE.  See the
% GNU General Public License for more details.
%
% You should have received a copy of the GNU General Public License along
% with this program; if not, write to the Free Software Foundation, Inc.,
% 51 Franklin Street, Fifth Floor, Boston, MA  02110-1301, USA.

% load the Bio-Formats library into the MATLAB environment
status = bfCheckJavaPath();
assert(status, ['Missing Bio-Formats library. Either add bioformats_package.jar '...
    'to the static Java path or add it to the Matlab path.']);

% Prompt for a file if not input
if nargin == 0 || exist(filename, 'file') == 0
  [file, path] = uigetfile(bfGetFileExtensions, 'Choose a file to open');
  filename = [path file];
  if isequal(path, 0) || isequal(file, 0), return; end
end

volume = bfopen(filename);
vaux{1} = cat(3, volume{1}{:, 1});
vaux{2} = filename;
volume{1} = vaux;
end
```
