## Supplementary material for "Enhancer architecture and chromatin accessibility constrain phenotypic space during development": Script-MLP: bfopen.m

```
function [result] = bfopen(id, varargin)
% Open microscopy images using Bio-Formats.
%
% SYNOPSIS r = bfopen(id)
%          r = bfopen(id, x, y, w, h)
%
% Input
%    r - the reader object (e.g. the output bfGetReader)
%
%    x - (Optional) A scalar giving the x-origin of the tile.
%    Default: 1
%
%    y - (Optional) A scalar giving the y-origin of the tile.
%    Default: 1
%
%    w - (Optional) A scalar giving the width of the tile.
%    Set to the width of the plane by default.
%
%    h - (Optional) A scalar giving the height of the tile.
%    Set to the height of the plane by default.
%
% Output
%
%    result - a cell array of cell arrays of (matrix, label) pairs,
%    with each matrix representing a single image plane, and each inner
%    list of matrices representing an image series.
%
% Portions of this code were adapted from:
% http://www.mathworks.com/support/solutions/en/data/1-2WPAYR/
%
% This method is ~1.5x-2.5x slower than Bio-Formats's command line
% showinf tool (MATLAB 7.0.4.365 R14 SP2 vs. java 1.6.0_20),
% due to overhead from copying arrays.
%
% Thanks to all who offered suggestions and improvements:
%     * Ville Rantanen
%     * Brett Shoelson
%     * Martin Offterdinger
%     * Tony Collins
%     * Cris Luengo
%     * Arnon Lieber
%     * Jimmy Fong
%
% NB: Internet Explorer sometimes erroneously renames the Bio-Formats library
%     to bioformats_package.zip. If this happens, rename it back to
%     bioformats_package.jar.
%
% For many examples of how to use the bfopen function, please see:
%     https://docs.openmicroscopy.org/latest/bio-formats/developers/matlab-dev.html

% OME Bio-Formats package for reading and converting biological file formats.
%
% Copyright (C) 2007 - 2017 Open Microscopy Environment:
%   - Board of Regents of the University of Wisconsin-Madison
%   - Glencoe Software, Inc.
%   - University of Dundee
%
% This program is free software: you can redistribute it and/or modify
% it under the terms of the GNU General Public License as
% published by the Free Software Foundation, either version 2 of the
% License, or (at your option) any later version.
%
% This program is distributed in the hope that it will be useful,
% but WITHOUT ANY WARRANTY; without even the implied warranty of
% MERCHANTABILITY or FITNESS FOR A PARTICULAR PURPOSE.  See the
% GNU General Public License for more details.
%
% You should have received a copy of the GNU General Public License along
% with this program; if not, write to the Free Software Foundation, Inc.,
% 51 Franklin Street, Fifth Floor, Boston, MA  02110-1301, USA.

% -- Configuration - customize this section to your liking --

% Toggle the autoloadBioFormats flag to control automatic loading
% of the Bio-Formats library using the javaaddpath command.
%
% For static loading, you can add the library to MATLAB's class path:
%     1. Type "edit classpath.txt" at the MATLAB prompt.
%     2. Go to the end of the file, and add the path to your JAR file
%        (e.g., C:/Program Files/MATLAB/work/bioformats_package.jar).
%     3. Save the file and restart MATLAB.
%
% There are advantages to using the static approach over javaaddpath:
%     1. If you use bfopen within a loop, it saves on overhead
%        to avoid calling the javaaddpath command repeatedly.
%     2. Calling 'javaaddpath' may erase certain global parameters.
autoloadBioFormats = 1;

% Toggle the stitchFiles flag to control grouping of similarly
% named files into a single dataset based on file numbering.
stitchFiles = 0;

% To work with compressed Evotec Flex, fill in your LuraWave license code.
%lurawaveLicense = 'xxxxxx-xxxxxxx';

% -- Main function - no need to edit anything past this point --

% load the Bio-Formats library into the MATLAB environment
status = bfCheckJavaPath(autoloadBioFormats);
assert(status, ['Missing Bio-Formats library. Either add bioformats_package.jar '...
    'to the static Java path or add it to the Matlab path.']);

% Prompt for a file if not input
if nargin == 0 || exist(id, 'file') == 0
  [file, path] = uigetfile(bfGetFileExtensions, 'Choose a file to open');
  id = [path file];
  if isequal(path, 0) || isequal(file, 0), return; end
end

% Initialize logging
bfInitLogging();

% Get the channel filler
r = bfGetReader(id, stitchFiles);

% Test plane size
if nargin >=4
    planeSize = javaMethod('getPlaneSize', 'loci.formats.FormatTools', ...
                           r, varargin{3}, varargin{4});
else
    planeSize = javaMethod('getPlaneSize', 'loci.formats.FormatTools', r);
end

if planeSize/(1024)^3 >= 2
    error(['Image plane too large. Only 2GB of data can be extracted '...
        'at one time. You can workaround the problem by opening '...
        'the plane in tiles.']);
end

numSeries = r.getSeriesCount();
result = cell(numSeries, 2);

globalMetadata = r.getGlobalMetadata();

for s = 1:numSeries
    fprintf(['Reading series #%d', '...', '\n'], s);
    r.setSeries(s - 1);
    pixelType = r.getPixelType();
    bpp = javaMethod('getBytesPerPixel', 'loci.formats.FormatTools', ...
                     pixelType);
    bppMax = power(2, bpp * 8);
    numImages = r.getImageCount();
    imageList = cell(numImages, 2);
    colorMaps = cell(numImages);
    for i = 1:numImages
        arr = bfGetPlane(r, i, varargin{:});

        % retrieve color map data
        if bpp == 1
            colorMaps{s, i} = r.get8BitLookupTable()';
        else
            colorMaps{s, i} = r.get16BitLookupTable()';
        end

        warning_state = warning ('off');
        if ~isempty(colorMaps{s, i})
            newMap = single(colorMaps{s, i});
            newMap(newMap < 0) = newMap(newMap < 0) + bppMax;
            colorMaps{s, i} = newMap / (bppMax - 1);
        end
        warning (warning_state);


        % build an informative title for our figure
        label = id;
        if numSeries > 1
            seriesName = char(r.getMetadataStore().getImageName(s - 1));
            if ~isempty(seriesName)
                label = [label, '; ', seriesName];
            else
                qs = int2str(s);
                label = [label, '; series ', qs, '/', int2str(numSeries)];
            end
        end
        if numImages > 1
            qi = int2str(i);
            label = [label, '; plane ', qi, '/', int2str(numImages)];
            if r.isOrderCertain()
                lz = 'Z';
                lc = 'C';
                lt = 'T';
            else
                lz = 'Z?';
                lc = 'C?';
                lt = 'T?';
            end
            zct = r.getZCTCoords(i - 1);
            sizeZ = r.getSizeZ();
            if sizeZ > 1
                qz = int2str(zct(1) + 1);
                label = [label, '; ', lz, '=', qz, '/', int2str(sizeZ)];
            end
            sizeC = r.getSizeC();
            if sizeC > 1
                qc = int2str(zct(2) + 1);
                label = [label, '; ', lc, '=', qc, '/', int2str(sizeC)];
            end
            sizeT = r.getSizeT();
            if sizeT > 1
                qt = int2str(zct(3) + 1);
                label = [label, '; ', lt, '=', qt, '/', int2str(sizeT)];
            end
        end

        % save image plane and label into the list
        imageList{i, 1} = arr;
        imageList{i, 2} = label;
    end

    % save images and metadata into our master series list
    result{s, 1} = imageList;

    % extract metadata table for this series
    seriesMetadata = r.getSeriesMetadata();
    javaMethod('merge', 'loci.formats.MetadataTools', ...
               globalMetadata, seriesMetadata, 'Global ');
    result{s, 2} = seriesMetadata;
    result{s, 3} = colorMaps;
    result{s, 4} = r.getMetadataStore();
end
r.close();
```
