## Supplementary material for "Enhancer architecture and chromatin accessibility constrain phenotypic space during development": Script-MLP: bfsave.m

```
function bfsave(varargin)
% BFSAVE Save a 5D matrix into an OME-TIFF using Bio-Formats library
%
%    bfsave(I, outputPath) writes the input 5D matrix into a new file
%    specified by outputPath.
%
%    bfsave(I, outputPath, dimensionOrder) specifies the dimension order of
%    the input matrix. Default valuse is XYZCT.
%
%    bfsave(I, outputPath, 'Compression', compression) specifies the
%    compression to use when writing the OME-TIFF file.
%
%    bfsave(I, outputPath, 'BigTiff', true) allows to save the file using
%    64-bit offsets
%
%    bfsave(I, outputPath, 'metadata', metadata) allows to use a custom
%    OME-XML metadata object when saving the file instead of creating a
%    minimal OME-XML metadata object from the input 5D matrix.
%
%    For more information, see https://docs.openmicroscopy.org/latest/bio-formats/developers/matlab-dev.html
%
%    Examples:
%
%        bfsave(zeros(100, 100), outputPath)
%        bfsave(zeros(100, 100, 2, 3, 4), outputPath)
%        bfsave(zeros(100, 100, 20), outputPath, 'dimensionOrder', 'XYTZC')
%        bfsave(zeros(100, 100), outputPath, 'Compression', 'LZW')
%        bfsave(zeros(100, 100), outputPath, 'BigTiff', true)
%        bfsave(zeros(100, 100), outputPath, 'metadata', metadata)
%
% See also: BFGETREADER, CREATEMINIMALOMEXMLMETADATA

% OME Bio-Formats package for reading and converting biological file formats.
%
% Copyright (C) 2012 - 2017 Open Microscopy Environment:
%   - Board of Regents of the University of Wisconsin-Madison
%   - Glencoe Software, Inc.
%   - University of Dundee
%
% This program is free software: you can redistribute it and/or modify
% it under the terms of the GNU General Public License as
% published by the Free Software Foundation, either version 2 of the
% License, or (at your option) any later version.
%
% This program is distributed in the hope that it will be useful,
% but WITHOUT ANY WARRANTY; without even the implied warranty of
% MERCHANTABILITY or FITNESS FOR A PARTICULAR PURPOSE.  See the
% GNU General Public License for more details.
%
% You should have received a copy of the GNU General Public License along
% with this program; if not, write to the Free Software Foundation, Inc.,
% 51 Franklin Street, Fifth Floor, Boston, MA  02110-1301, USA.

% verify that enough memory is allocated
bfCheckJavaMemory();

% Check for required jars in the Java path
bfCheckJavaPath();

% Input check
ip = inputParser;
ip.addRequired('I', @isnumeric);
ip.addRequired('outputPath', @ischar);
ip.addOptional('dimensionOrder', 'XYZCT', @(x) ismember(x, getDimensionOrders()));
ip.addParamValue('metadata', [], @(x) isa(x, 'loci.formats.ome.OMEXMLMetadata'));
ip.addParamValue('Compression', '',  @(x) ismember(x, getCompressionTypes()));
ip.addParamValue('BigTiff', false , @islogical);
ip.parse(varargin{:});

% Create metadata
if isempty(ip.Results.metadata)
    metadata = createMinimalOMEXMLMetadata(ip.Results.I,...
        ip.Results.dimensionOrder);
else
    metadata = ip.Results.metadata;
end

% Create ImageWriter
writer = javaObject('loci.formats.ImageWriter');
writer.setWriteSequentially(true);
writer.setMetadataRetrieve(metadata);
if ~isempty(ip.Results.Compression)
    writer.setCompression(ip.Results.Compression);
end
if ip.Results.BigTiff
    writer.getWriter(ip.Results.outputPath).setBigTiff(ip.Results.BigTiff);
end
writer.setId(ip.Results.outputPath);

% Load conversion tools for saving planes
switch class(ip.Results.I)
    case {'int8', 'uint8'}
        getBytes = @(x) x(:);
    case {'uint16','int16'}
        getBytes = @(x) javaMethod('shortsToBytes', 'loci.common.DataTools', x(:), 0);
    case {'uint32','int32'}
        getBytes = @(x) javaMethod('intsToBytes', 'loci.common.DataTools', x(:), 0);
    case {'single'}
        getBytes = @(x) javaMethod('floatsToBytes', 'loci.common.DataTools', x(:), 0);
    case 'double'
        getBytes = @(x) javaMethod('doublesToBytes', 'loci.common.DataTools', x(:), 0);
end

% Save planes to the writer
nPlanes = metadata.getPixelsSizeZ(0).getValue() *...
    metadata.getPixelsSizeC(0).getValue() *...
    metadata.getPixelsSizeT(0).getValue();
zctCoord = [size(ip.Results.I, 3) size(ip.Results.I, 4)...
    size(ip.Results.I, 5)];
for index = 1 : nPlanes
    [i, j, k] = ind2sub(zctCoord, index);
    plane = ip.Results.I(:, :, i, j, k)';
    writer.saveBytes(index-1, getBytes(plane));
end
writer.close();

end

function dimensionOrders = getDimensionOrders()
% List all values of DimensionOrder
dimensionOrderValues = javaMethod('values', 'ome.xml.model.enums.DimensionOrder');
dimensionOrders = cell(numel(dimensionOrderValues), 1);
for i = 1 :numel(dimensionOrderValues),
    dimensionOrders{i} = char(dimensionOrderValues(i).toString());
end
end

function compressionTypes = getCompressionTypes()
% List all values of Compression
writer = javaObject('loci.formats.ImageWriter');
if is_octave()
    %% FIXME when https://savannah.gnu.org/bugs/?42700 gets fixed
    types = writer.getCompressionTypes();
    nTypes = numel(types);
    compressionTypes = cell(nTypes, 1);
    for i = 1:nTypes
        compressionTypes{i} = char(types(i));
    end
else
    compressionTypes = arrayfun(@char, writer.getCompressionTypes(),...
                                'UniformOutput', false);
end
end
```
