## Supplementary material for "Enhancer architecture and chromatin accessibility constrain phenotypic space during development": Script-MLP: bfUpgradeCheck.m

```
function bfUpgradeCheck(varargin)
% Check for new version of Bio-Formats and update it if applicable
%
% SYNOPSIS: bfUpgradeCheck(autoDownload, 'STABLE')
%
% Input
%    autoDownload - Optional. A boolean specifying of the latest version
%    should be downloaded
%
%    versions -  Optional: a string sepecifying the version to fetch.
%    Should be either trunk, daily or stable (case insensitive)
%
% Output
%    none

% OME Bio-Formats package for reading and converting biological file formats.
%
% Copyright (C) 2012 - 2017 Open Microscopy Environment:
%   - Board of Regents of the University of Wisconsin-Madison
%   - Glencoe Software, Inc.
%   - University of Dundee
%
% This program is free software: you can redistribute it and/or modify
% it under the terms of the GNU General Public License as
% published by the Free Software Foundation, either version 2 of the
% License, or (at your option) any later version.
%
% This program is distributed in the hope that it will be useful,
% but WITHOUT ANY WARRANTY; without even the implied warranty of
% MERCHANTABILITY or FITNESS FOR A PARTICULAR PURPOSE.  See the
% GNU General Public License for more details.
%
% You should have received a copy of the GNU General Public License along
% with this program; if not, write to the Free Software Foundation, Inc.,
% 51 Franklin Street, Fifth Floor, Boston, MA  02110-1301, USA.
% Check input
ip = inputParser;
ip.addOptional('autoDownload', false, @isscalar);
versions = {'stable', 'daily', 'trunk'};
ip.addOptional('version', 'STABLE', @(x) any(strcmpi(x, versions)))
ip.parse(varargin{:})

% Create UpgradeChecker
upgrader = javaObject('loci.formats.UpgradeChecker');
if upgrader.alreadyChecked(), return; end

% Check for new version of Bio-Formats
if is_octave()
    caller = 'Octave';
else
    caller = 'MATLAB';
end
if ~ upgrader.newVersionAvailable(caller)
    fprintf('*** bioformats_package.jar is up-to-date ***\n');
    return;
end

fprintf('*** A new stable version of Bio-Formats is available ***\n');
% If appliable, download new version of Bioformats
if ip.Results.autoDownload
    fprintf('*** Downloading... ***');
    path = fullfile(fileparts(mfilename('fullpath')), 'bioformats_package.jar');
    buildName = [upper(ip.Results.version) '_BUILD'];
    upgrader.install(loci.formats.UpgradeChecker.(buildName), path);
    fprintf('*** Upgrade will be finished when MATLAB is restarted ***\n');
end
```
