## Supplementary material for "Enhancer architecture and chromatin accessibility constrain phenotypic space during development": Script-MLP: createMinimalOMEXMLMetadata.m

```
function metadata = createMinimalOMEXMLMetadata(I, varargin)
% CREATEMINIMALOMEXMLMETADATA Create an OME-XML metadata object from an input matrix
%
%    createMinimalOMEXMLMetadata(I) creates an OME-XML metadata object from
%    an input 5-D array. Minimal metadata information is stored such as the
%    pixels dimensions, dimension order and type. The output object is a
%    metadata object of type loci.formats.ome.OMEXMLMetadata.
%
%    createMinimalOMEXMLMetadata(I, dimensionOrder) specifies the dimension
%    order of the input matrix. Default valuse is XYZCT.
%
%    Examples:
%
%        metadata = createMinimalOMEXMLMetadata(zeros(100, 100));
%        metadata = createMinimalOMEXMLMetadata(zeros(10, 10, 2), 'XYTZC');
%
% See also: BFSAVE

% OME Bio-Formats package for reading and converting biological file formats.
%
% Copyright (C) 2012 - 2017 Open Microscopy Environment:
%   - Board of Regents of the University of Wisconsin-Madison
%   - Glencoe Software, Inc.
%   - University of Dundee
%
% This program is free software: you can redistribute it and/or modify
% it under the terms of the GNU General Public License as
% published by the Free Software Foundation, either version 2 of the
% License, or (at your option) any later version.
%
% This program is distributed in the hope that it will be useful,
% but WITHOUT ANY WARRANTY; without even the implied warranty of
% MERCHANTABILITY or FITNESS FOR A PARTICULAR PURPOSE.  See the
% GNU General Public License for more details.
%
% You should have received a copy of the GNU General Public License along
% with this program; if not, write to the Free Software Foundation, Inc.,
% 51 Franklin Street, Fifth Floor, Boston, MA  02110-1301, USA.

% Not using the inputParser for first argument as it copies data
assert(isnumeric(I), 'First argument must be numeric');

% Input check
ip = inputParser;
ip.addOptional('dimensionOrder', 'XYZCT', @(x) ismember(x, getDimensionOrders()));
ip.parse(varargin{:});

% Create metadata
toInt = @(x) javaObject('ome.xml.model.primitives.PositiveInteger', ...
                        javaObject('java.lang.Integer', x));
OMEXMLService = javaObject('loci.formats.services.OMEXMLServiceImpl');
metadata = OMEXMLService.createOMEXMLMetadata();
metadata.createRoot();
metadata.setImageID('Image:0', 0);
metadata.setPixelsID('Pixels:0', 0);
if is_octave()
    java_true = java_get('java.lang.Boolean', 'TRUE');
else
    java_true = java.lang.Boolean.TRUE;
end
metadata.setPixelsBigEndian(java_true, 0);

% Set dimension order
dimensionOrderEnumHandler = javaObject('ome.xml.model.enums.handlers.DimensionOrderEnumHandler');
dimensionOrder = dimensionOrderEnumHandler.getEnumeration(ip.Results.dimensionOrder);
metadata.setPixelsDimensionOrder(dimensionOrder, 0);

% Set pixels type
pixelTypeEnumHandler = javaObject('ome.xml.model.enums.handlers.PixelTypeEnumHandler');
if strcmp(class(I), 'single')
    pixelsType = pixelTypeEnumHandler.getEnumeration('float');
else
    pixelsType = pixelTypeEnumHandler.getEnumeration(class(I));
end
metadata.setPixelsType(pixelsType, 0);

% Read pixels size from image and set it to the metadat
sizeX = size(I, 2);
sizeY = size(I, 1);
sizeZ = size(I, find(ip.Results.dimensionOrder == 'Z'));
sizeC = size(I, find(ip.Results.dimensionOrder == 'C'));
sizeT = size(I, find(ip.Results.dimensionOrder == 'T'));
metadata.setPixelsSizeX(toInt(sizeX), 0);
metadata.setPixelsSizeY(toInt(sizeY), 0);
metadata.setPixelsSizeZ(toInt(sizeZ), 0);
metadata.setPixelsSizeC(toInt(sizeC), 0);
metadata.setPixelsSizeT(toInt(sizeT), 0);

% Set channels ID and samples per pixel
for i = 1: sizeC
    metadata.setChannelID(['Channel:0:' num2str(i-1)], 0, i-1);
    metadata.setChannelSamplesPerPixel(toInt(1), 0, i-1);
end

end

function dimensionOrders = getDimensionOrders()
% List all values of DimensionOrder
dimensionOrderValues = javaMethod('values', 'ome.xml.model.enums.DimensionOrder');
dimensionOrders = cell(numel(dimensionOrderValues), 1);
for i = 1 :numel(dimensionOrderValues),
    dimensionOrders{i} = char(dimensionOrderValues(i).toString());
end
end
```
