## Supplementary material for "Enhancer architecture and chromatin accessibility constrain phenotypic space during development": Script-MLP: calculateProjection.m

```
function projection = calculateProjection(...
    projectionType, nSlices, imageStack, varargin)

% Calculate projection for a nuclear channel
lowerSlice = 1;
upperSlice = nSlices;

if strcmpi(projectionType, 'middleprojection')
    lowerSlice = round(nSlices * .50);
    upperSlice =round(nSlices * .75);
elseif strcmpi(projectionType, 'midsumprojection')
    lowerSlice = round(nSlices * .33);
    upperSlice =round(nSlices * .66);
end

if ~isempty(varargin)
    upperSlice = varargin{1};
    lowerSlice = varargin{2};
end

imageStack = imageStack(:, :, lowerSlice:upperSlice);
    

if strcmpi(projectionType, 'medianprojection')
    projection = median(imageStack, 3);
elseif strcmpi(projectionType, 'maxprojection') || strcmpi(projectionType, 'middleprojection')
    projection = max(imageStack, [], 3);
elseif strcmpi(projectionType, 'midsumprojection')
    projection = sum(imageStack, 3);
else
    SortedHisSlices = sort(imageStack, 3, 'descend');
    if ~isempty(varargin)
        max_custom = varargin{1};
        min_custom = varargin{2};
        if length(size(SortedHisSlices)) > 2
            projection = mean(SortedHisSlices(:, :, max_custom:min_custom), 3);
        else
            projection = SortedHisSlices;
        end
    else
        ProjectionBounds = strsplit(projectionType, ':');
        max_custom = str2double(ProjectionBounds{2});
        min_custom = str2double(ProjectionBounds{3});
        projection = mean(SortedHisSlices(:, :, max_custom:min_custom), 3);
    end
end

end
```
