## Supplementary material for "Enhancer architecture and chromatin accessibility constrain phenotypic space during development": Script-MLP: findNuclei.m

```
function [ xy, varargout ] = findNuclei(FrameInfo, hisImage,...
    nucleusDiameter, embryoMask, varargin )
%FINDNUCLEI This function finds the position of nuclei within a frame which
%is used as an initial input for the tracking algorithm.
%   Detection is made by detecting the local maxima in a
%   Laplacian-of-Gaussian filtered image.
%
%%

hisImage = double(hisImage);

%this is an optional argument from tracknuclei. it's specified as global
%because it's too hard to figure out how to get it from tracknuclei ->
%findnuclei -AR 8/8/2020
global mixedPolarity


%% Initializing variables
% Load parameters
LoGratio = getDefaultParameters(FrameInfo,'LoGratio');
space_resolution = getDefaultParameters(FrameInfo,'space resolution');
localMaximumRadius = LoGratio*nucleusDiameter/space_resolution;
LoGradius = nucleusDiameter/space_resolution*LoGratio;
edgeClearance = getDefaultParameters(FrameInfo,'edge clearance')*nucleusDiameter/space_resolution;
useEllipse = false;

% Added by GM 1/9/2019
useMultithresh=false;
k=1;
while k<=length(varargin)
    if strcmpi(varargin{k},'useMultithresh')
        useMultithresh = true;
        k=k+1;
    elseif strcmpi(varargin{k},'ellipse')
        useEllipse = true;
        k = k+1;
    end
end
% Added by GM 1/9/2019

% Added by NL and GM on 11/23/2019
% Edited by GM on 1/7/2019
xDim_um = FrameInfo(1).PixelsPerLine * FrameInfo(1).PixelSize;
yDim_um = FrameInfo(1).LinesPerFrame * FrameInfo(1).PixelSize;
if yDim_um > 150 && xDim_um > 150
    pixelvalues = unique(hisImage(:));
    thresh = pixelvalues(2);
    f_sigma = round(nucleusDiameter / FrameInfo(1).PixelSize);
    bwfill = imfill(hisImage>thresh,'holes');
    I_inside = bwselect(bwfill,round(size(bwfill,2)/2), round(size(bwfill, 1)/2));
    I_inside=uint16((2^16-1)*I_inside);
    I_blurred = imfilter(I_inside,...
        fspecial('gaussian',2*f_sigma,f_sigma),'symmetric','conv');
    level = graythresh(I_blurred);
    embryoMask = im2bw(I_blurred,level);
else
    if ~exist('embryoMask','var') || isempty(embryoMask)
        embryoMask = true(size(hisImage, 1));
    end
end

%%

if nargin > 5
    if isnumeric(varargin{1})
        targetNumber = varargin{1}; % coarse estimate of the number of nuclei that should be found.
    end
end


% Create the mask used to detect local maxima.
localMaxMask = fspecial('disk',localMaximumRadius);
localMaxMask = imbinarize(mat2gray(localMaxMask),graythresh(mat2gray(localMaxMask)));
localMaxMask(round(length(localMaxMask)/2),round(length(localMaxMask)/2))  = 0;

if useEllipse
    N = 4*ceil(localMaximumRadius);
    [x,y] = meshgrid(1:N);
    r1 = 2*localMaximumRadius;
    r2 = localMaximumRadius;
    
    localMaxMask = sqrt((x - N/2).^2 / r1^2 + (y - N/2).^2 / r2^2);
    localMaxMask = double(localMaxMask < 1);
    localMaxMask(round(N/2),round(N/2)) = 0;
end


%% Main body

% Filter the image
filteredImg = fourierFilterWithSymmetricBoundaryConditions(...
    hisImage,-fspecial('log',round(10*LoGradius),LoGradius));

%if this abs is taken, both dark and bright areas will be highlighted in
%the filtered image. off by default since that's not the expected behavior.
if mixedPolarity
    filteredImg = abs(filteredImg);
end

% Find local maxima
maxima = (filteredImg > imdilate(filteredImg,localMaxMask) );% Commented out GM on 9/3/20 Seems to no longer work after updates & embryoMask; 

% Smooth the image to get more robust maxima values:
G = imfilter(filteredImg,fspecial('disk',3),'symmetric');
maximaLinearIndices = find(maxima>0);
[x_maxima,y_maxima] = ind2sub(size(maxima),maximaLinearIndices);


%Check whether anything was found in this image. If nothing is found,
%this is usually the result of an empty frame
% if isempty(xm) || isempty(ym)
%    error(['No nuclei found in frame ',num2str(frameNumber),'. Check that that frame is not blank.'])
% end

if ~useMultithresh
    try
        [v,C] = voronoin([x_maxima y_maxima]);
    catch
        %not sure when this exception occurs-
        %it adds additional options (http://www.qhull.org/html/qh-optq.htm) -AR
        %Qbb - scale last coordinate to [0,m] for Delaunay (this is the
        %default for voronoin. it doesn't need to be included)
        %Qz -add a point-at-infinity for Delaunay triangulations
        [v,C] = voronoin([x_maxima y_maxima],{'Qbb','Qz'});
    end
    
    %get the voronoi edge pixel indices(linear) that are within the image
    ind_v = v(:,1) > 0.5 &...
        v(:,2) > 0.5...
        & v(:,1) < size(maxima,1)...
        & v(:,2) < size(maxima,2);
    
    %get the corresponding subscript indices from the above linear indices
    background_ind = sub2ind(size(maxima),round(v(ind_v,1)),round(v(ind_v,2)));
    
    %append the maxima to the voronoi edges. this corresponds to an image
    %with a bunch of voronoi edges with dots somewhere inside them
    sample_ind = [maximaLinearIndices(:) ; background_ind(:)];
    
    %these are fluorescence values in the (smoothed) nuclear image at the
    %locations of sample_ind (the voronoi vertices and maxima)
    nuc1 = G(sample_ind);
    
    %autothresholding with Otsu's method
    thresh = graythresh(mat2gray(nuc1));
    indNuclei = imbinarize(mat2gray(nuc1),thresh);
else
    b_points = boundary(x_maxima,y_maxima);
    index_vec = 1:numel(x_maxima);
    indNuclei = ~ismember(index_vec,b_points);
end

%% Check the segmentation if a target number was provided.
if exist('targetNumber', 'var')
    indNuclei = useTargetNumber(targetNumber, nuc1, indNuclei);
end
%%

indNuclei = indNuclei(1:numel(maximaLinearIndices));

% Get rid of all nuclei below thresh
maximaLinearIndices = maximaLinearIndices(indNuclei);

% Convert into coordinates
[x,y] = ind2sub(size(hisImage),maximaLinearIndices);

xy = [x y];

if nargout > 1
    % Those values are the intensities of the maxima that
    %could be used to sort them.
    varargout{1} = G(maximaLinearIndices);
    
    varargout{2} = v;
    varargout{3} = C;
end

end
```
