## Supplementary material for "Enhancer architecture and chromatin accessibility constrain phenotypic space during development": Script-MLP: fit_ellipse.m

```
function ellipse_t = fit_ellipse( x,y,axis_handle )
%
% fit_ellipse - finds the best fit to an ellipse for the given set of points.
%
% Format:   ellipse_t = fit_ellipse( x,y,axis_handle )
%
% Input:    x,y         - a set of points in 2 column vectors. AT LEAST 5 points are needed !
%           axis_handle - optional. a handle to an axis, at which the estimated ellipse 
%                         will be drawn along with it's axes
%
% Output:   ellipse_t - structure that defines the best fit to an ellipse
%                       a           - sub axis (radius) of the X axis of the non-tilt ellipse
%                       b           - sub axis (radius) of the Y axis of the non-tilt ellipse
%                       phi         - orientation in radians of the ellipse (tilt)
%                       X0          - center at the X axis of the non-tilt ellipse
%                       Y0          - center at the Y axis of the non-tilt ellipse
%                       X0_in       - center at the X axis of the tilted ellipse
%                       Y0_in       - center at the Y axis of the tilted ellipse
%                       long_axis   - size of the long axis of the ellipse
%                       short_axis  - size of the short axis of the ellipse
%                       status      - status of detection of an ellipse
%
% Note:     if an ellipse was not detected (but a parabola or hyperbola), then
%           an empty structure is returned

% =====================================================================================
%                  Ellipse Fit using Least Squares criterion
% =====================================================================================
% We will try to fit the best ellipse to the given measurements. the mathematical
% representation of use will be the CONIC Equation of the Ellipse which is:
% 
%    Ellipse = a*x^2 + b*x*y + c*y^2 + d*x + e*y + f = 0
%   
% The fit-estimation method of use is the Least Squares method (without any weights)
% The estimator is extracted from the following equations:
%
%    g(x,y;A) := a*x^2 + b*x*y + c*y^2 + d*x + e*y = f
%
%    where:
%       A   - is the vector of parameters to be estimated (a,b,c,d,e)
%       x,y - is a single measurement
%
% We will define the cost function to be:
%
%   Cost(A) := (g_c(x_c,y_c;A)-f_c)'*(g_c(x_c,y_c;A)-f_c)
%            = (X*A+f_c)'*(X*A+f_c) 
%            = A'*X'*X*A + 2*f_c'*X*A + N*f^2
%
%   where:
%       g_c(x_c,y_c;A) - vector function of ALL the measurements
%                        each element of g_c() is g(x,y;A)
%       X              - a matrix of the form: [x_c.^2, x_c.*y_c, y_c.^2, x_c, y_c ]
%       f_c            - is actually defined as ones(length(f),1)*f
%
% Derivation of the Cost function with respect to the vector of parameters "A" yields:
%
%   A'*X'*X = -f_c'*X = -f*ones(1,length(f_c))*X = -f*sum(X)
%
% Which yields the estimator:
%
%       ~~~~~~~~~~~~~~~~~~~~~~~~~~~~~~~~~~~~~~~~~~~~~~~~~~~~~~~~~~~~~~~~~~~~~~~~~~~~
%       |  A_least_squares = -f*sum(X)/(X'*X) ->(normalize by -f) = sum(X)/(X'*X)  |
%       ~~~~~~~~~~~~~~~~~~~~~~~~~~~~~~~~~~~~~~~~~~~~~~~~~~~~~~~~~~~~~~~~~~~~~~~~~~~~
%
% (We will normalize the variables by (-f) since "f" is unknown and can be accounted for later on)
%  
% NOW, all that is left to do is to extract the parameters from the Conic Equation.
% We will deal the vector A into the variables: (A,B,C,D,E) and assume F = -1;
%
%    Recall the conic representation of an ellipse:
% 
%       A*x^2 + B*x*y + C*y^2 + D*x + E*y + F = 0
% 
% We will check if the ellipse has a tilt (=orientation). The orientation is present
% if the coefficient of the term "x*y" is not zero. If so, we first need to remove the
% tilt of the ellipse.
%
% If the parameter "B" is not equal to zero, then we have an orientation (tilt) to the ellipse.
% we will remove the tilt of the ellipse so as to remain with a conic representation of an 
% ellipse without a tilt, for which the math is more simple:
%
% Non tilt conic rep.:  A`*x^2 + C`*y^2 + D`*x + E`*y + F` = 0
%
% We will remove the orientation using the following substitution:
%   
%   Replace x with cx+sy and y with -sx+cy such that the conic representation is:
%   
%   A(cx+sy)^2 + B(cx+sy)(-sx+cy) + C(-sx+cy)^2 + D(cx+sy) + E(-sx+cy) + F = 0
%
%   where:      c = cos(phi)    ,   s = sin(phi)
%
%   and simplify...
%
%       x^2(A*c^2 - Bcs + Cs^2) + xy(2A*cs +(c^2-s^2)B -2Ccs) + ...
%           y^2(As^2 + Bcs + Cc^2) + x(Dc-Es) + y(Ds+Ec) + F = 0
%
%   The orientation is easily found by the condition of (B_new=0) which results in:
% 
%   2A*cs +(c^2-s^2)B -2Ccs = 0  ==> phi = 1/2 * atan( b/(c-a) )
%   
%   Now the constants   c=cos(phi)  and  s=sin(phi)  can be found, and from them
%   all the other constants A`,C`,D`,E` can be found.
%
%   A` = A*c^2 - B*c*s + C*s^2                  D` = D*c-E*s
%   B` = 2*A*c*s +(c^2-s^2)*B -2*C*c*s = 0      E` = D*s+E*c 
%   C` = A*s^2 + B*c*s + C*c^2
%
% Next, we want the representation of the non-tilted ellipse to be as:
%
%       Ellipse = ( (X-X0)/a )^2 + ( (Y-Y0)/b )^2 = 1
%
%       where:  (X0,Y0) is the center of the ellipse
%               a,b     are the ellipse "radiuses" (or sub-axis)
%
% Using a square completion method we will define:
%       
%       F`` = -F` + (D`^2)/(4*A`) + (E`^2)/(4*C`)
%
%       Such that:    a`*(X-X0)^2 = A`(X^2 + X*D`/A` + (D`/(2*A`))^2 )
%                     c`*(Y-Y0)^2 = C`(Y^2 + Y*E`/C` + (E`/(2*C`))^2 )
%
%       which yields the transformations:
%       
%           X0  =   -D`/(2*A`)
%           Y0  =   -E`/(2*C`)
%           a   =   sqrt( abs( F``/A` ) )
%           b   =   sqrt( abs( F``/C` ) )
%
% And finally we can define the remaining parameters:
%
%   long_axis   = 2 * max( a,b )
%   short_axis  = 2 * min( a,b )
%   Orientation = phi
%
%

% initialize
orientation_tolerance = 1e-3;

% empty warning stack
warning( '' );

% prepare vectors, must be column vectors
x = x(:);
y = y(:);

% remove bias of the ellipse - to make matrix inversion more accurate. (will be added later on).
mean_x = mean(x);
mean_y = mean(y);
x = x-mean_x;
y = y-mean_y;

% the estimation for the conic equation of the ellipse
X = [x.^2, x.*y, y.^2, x, y ];
a = sum(X)/(X'*X);

% check for warnings
if ~isempty( lastwarn )
    disp( 'stopped because of a warning regarding matrix inversion' );
    ellipse_t = [];
    return
end

% extract parameters from the conic equation
[a,b,c,d,e] = deal( a(1),a(2),a(3),a(4),a(5) );

% remove the orientation from the ellipse
if ( min(abs(b/a),abs(b/c)) > orientation_tolerance )
    
    orientation_rad = 1/2 * atan( b/(c-a) );
    cos_phi = cos( orientation_rad );
    sin_phi = sin( orientation_rad );
    [a,b,c,d,e] = deal(...
        a*cos_phi^2 - b*cos_phi*sin_phi + c*sin_phi^2,...
        0,...
        a*sin_phi^2 + b*cos_phi*sin_phi + c*cos_phi^2,...
        d*cos_phi - e*sin_phi,...
        d*sin_phi + e*cos_phi );
    [mean_x,mean_y] = deal( ...
        cos_phi*mean_x - sin_phi*mean_y,...
        sin_phi*mean_x + cos_phi*mean_y );
else
    orientation_rad = 0;
    cos_phi = cos( orientation_rad );
    sin_phi = sin( orientation_rad );
end

% check if conic equation represents an ellipse
test = a*c;
switch (1)
case (test>0),  status = '';
case (test==0), status = 'Parabola found';  warning( 'fit_ellipse: Did not locate an ellipse' );
case (test<0),  status = 'Hyperbola found'; warning( 'fit_ellipse: Did not locate an ellipse' );
end

% if we found an ellipse return it's data
if (test>0)
    
    % make sure coefficients are positive as required
    if (a<0), [a,c,d,e] = deal( -a,-c,-d,-e ); end
    
    % final ellipse parameters
    X0          = mean_x - d/2/a;
    Y0          = mean_y - e/2/c;
    F           = 1 + (d^2)/(4*a) + (e^2)/(4*c);
    [a,b]       = deal( sqrt( F/a ),sqrt( F/c ) );    
    long_axis   = 2*max(a,b);
    short_axis  = 2*min(a,b);

    % rotate the axes backwards to find the center point of the original TILTED ellipse
    R           = [ cos_phi sin_phi; -sin_phi cos_phi ];
    P_in        = R * [X0;Y0];
    X0_in       = P_in(1);
    Y0_in       = P_in(2);
    
    % pack ellipse into a structure
    ellipse_t = struct( ...
        'a',a,...
        'b',b,...
        'phi',orientation_rad,...
        'X0',X0,...
        'Y0',Y0,...
        'X0_in',X0_in,...
        'Y0_in',Y0_in,...
        'long_axis',long_axis,...
        'short_axis',short_axis,...
        'status','' );
else
    % report an empty structure
    ellipse_t = struct( ...
        'a',[],...
        'b',[],...
        'phi',[],...
        'X0',[],...
        'Y0',[],...
        'X0_in',[],...
        'Y0_in',[],...
        'long_axis',[],...
        'short_axis',[],...
        'status',status );
end

% check if we need to plot an ellipse with it's axes.
if (nargin>2) & ~isempty( axis_handle ) & (test>0)
    
    % rotation matrix to rotate the axes with respect to an angle phi
    R = [ cos_phi sin_phi; -sin_phi cos_phi ];
    
    % the axes
    ver_line        = [ [X0 X0]; Y0+b*[-1 1] ];
    horz_line       = [ X0+a*[-1 1]; [Y0 Y0] ];
    new_ver_line    = R*ver_line;
    new_horz_line   = R*horz_line;
    
    % the ellipse
    theta_r         = linspace(0,2*pi);
    ellipse_x_r     = X0 + a*cos( theta_r );
    ellipse_y_r     = Y0 + b*sin( theta_r );
    rotated_ellipse = R * [ellipse_x_r;ellipse_y_r];
    
    % draw
    hold_state = get( axis_handle,'NextPlot' );
    set( axis_handle,'NextPlot','add' );
    plot( new_ver_line(1,:),new_ver_line(2,:),'r' );
    plot( new_horz_line(1,:),new_horz_line(2,:),'r' );
    plot( rotated_ellipse(1,:),rotated_ellipse(2,:),'r' );
    set( axis_handle,'NextPlot',hold_state );
end
```
