## Supplementary material for "Enhancer architecture and chromatin accessibility constrain phenotypic space during development": Script-MLP: fourierFilterWithSymmetricBoundaryConditions.m

```
function [ y ] = fourierFilterWithSymmetricBoundaryConditions( x, h )

% Compute y = h*x with symmetric boundary extension
xSym = [x,fliplr(x)];       % Symmetrize horizontally
xSym = [xSym;flipud(xSym)]; % Symmetrize vertically
[k1,k2] = size(h);
hpad = zeros(size(xSym));
hpad([end+1-floor(k1/2):end,1:ceil(k1/2)], ...
    [end+1-floor(k2/2):end,1:ceil(k2/2)]) = h;
y = real(ifft2(fft2(hpad).*fft2(xSym)));
y = y(1:size(y,1)/2,1:size(y,2)/2);


end
```
