## Supplementary material for "Enhancer architecture and chromatin accessibility constrain phenotypic space during development": Script-MLP: getDefaultParameters.m

```
function [ parameterValue ] = getDefaultParameters(FrameInfo, parameterName, varargin)
%GETDEFAULTPARAMETERS All default settings are grouped in here, so that
%they can dynamically be redefined in all functions.
%  
% ORGANIZATION
% Each function has a field in the 'parameters' structure. It contains a
% cell with one column per parameter. Each column is a cell, with the first
% field corresponding to the default value and all the following fields to
% the possible names that can be used to get the given value. 
% NAME CONFLICTS
% If within one function several parameters have identical names, the value
% of the one appearing first in the cell is returned.

%% GLOBAL PARAMETERS

% % time_resolution = 37;
% % space_resolution = .22;
% time_resolution = 9.56; %AR 9/11/2018 
% space_resolution = .0708;%AR 9/11/2018 
time_resolution = median(diff([FrameInfo.Time]));     %Median separation between frames (in seconds)
space_resolution = FrameInfo(1).PixelSize; % (um)

parameters.global = {...
    {time_resolution,'time resolution', 'timeResolution', 'time', 't'}...   % imaging period in seconds.
    {space_resolution, 'space resolution', 'spaceResolution', 'space', 's'}...% pixel size in micrometers. 
    {7.26, 'diameter nc7', 'd7'}... % diameter of nuclei during nc7 in micrometers.
    {7.26, 'diameter nc8', 'd8'}... % diameter of nuclei during nc8 in micrometers.
    {7.26, 'diameter nc9', 'd9'}... % diameter of nuclei during nc9 in micrometers.
    {7.26, 'diameter nc10', 'd10'}... % diameter of nuclei during nc10 in micrometers.
    {6.16, 'diameter nc11', 'd11'}... % diameter of nuclei during nc11 in micrometers.
    {5.72, 'diameter nc12', 'd12'}... % diameter of nuclei during nc12 in micrometers.
    {4.84, 'diameter nc13', 'd13'}... % diameter of nuclei during nc13 in micrometers.
    {3.96, 'diameter nc14', 'd14'}... % diameter of nuclei during nc14 in micrometers.
    {35/88, 'LoGratio'}... % Ratio between the radius of the LoG filter and the nucleus diameter (when both in the same unit).
    {0.81, 'max Interphase Displacement', 'maxInterphaseDisplacement'}... % in nucleus diameters/minute. Used to determine the maximal distance that a nucleus can move from one frame to the next after correcting for a bulk shift.
    {0.75, 'edge clearance'}... % in nucleus diameter. Space within which nuclei are discarded if their center falls in.
    {80, 'margin mitosis'}... % in seconds. Maximum time before or after a mitosis is detected when nuclei could still be dividing.
    {37, 'increased precision before mitosis'}... % in seconds. Before (and after) every division, the number of points that are used to estimate nuclei movements is increased to cope with the more complex movements that occur. This slows down the processing and thus is limited to a few frames.
    {111, 'increased precision after mitosis'}... % in seconds. After (and before) every division, the number of points that are used to estimate nuclei movements is increased to cope with the more complex movements that occur. This slows down the processing and thus is limited to a few frames.
    };

%% FUNCTIONS' PARAMETERS

% findMitosis
parameters.findMitosis = {...
    {space_resolution, 'smoothing factor', 'smoothing'}... % std of the spatial gaussian filter in micrometers. Was originally 1 pixel.
	{259, 'time window', 'timewindow'}... % in seconds. Was originally 7 frames.
    };

% choseFramesToStartTracking
parameters.choseFramesToStartTracking = {...
    {740, 'delta t', 'time'}... % in seconds. Time into nc14 when nuclei are in optimal shape to start the tracking.
    };

% findNuclei
parameters.findNuclei = {...
    {1.1, 'nuclei distance nc10', 'nucleidistancenc10', 'ncdistance10'}... % no units. This value times the corresponding nucleus radius is the minimal distance between two different nuclei.
    {1.1, 'nuclei distance nc11', 'nucleidistancenc11', 'ncdistance11'}... % no units. This value times the corresponding nucleus radius is the minimal distance between two different nuclei.
    {1.1, 'nuclei distance nc12', 'nucleidistancenc12', 'ncdistance12'}... % no units. This value times the corresponding nucleus radius is the minimal distance between two different nuclei.
    {1.1, 'nuclei distance nc13', 'nucleidistancenc13', 'ncdistance13'}... % no units. This value times the corresponding nucleus radius is the minimal distance between two different nuclei.
    {1.1, 'nuclei distance nc14', 'nucleidistancenc14', 'ncdistance14'}... % no units. This value times the corresponding nucleus radius is the minimal distance between two different nuclei.
    };

% trackToTheNextFrame
parameters.trackToTheNextFrame = {...
    {4, 'max shift correction'}... % no units. This value times the corresponding nucleus diameter is used as a scale for possible shifts from one frame to the next.
    };

%% CODE

% Default is set as a global parameter
if nargin == 2
    field = 'global';
else
    field = varargin{1};
end

fields = fieldnames(parameters);

% Reorganize all parameters into an array for further search of parameters.
for j = 1:numel(fields)
    Nparameters = length(parameters.(fields{j}));
    par.(fields{j}) = cell(Nparameters,2);
    
    for jj = 1:Nparameters
        par.(fields{j}){jj,1} = parameters.(fields{j}){jj}(2:end); % All possible names
        par.(fields{j}){jj,2} = parameters.(fields{j}){jj}{1}; % Value
    end
end

% Find the name called.
fieldInd = find(strcmpi(field,fields));

% Deal with an invalid name.
if isempty(fieldInd)
    error(['Trying to get parameter values for unknown function ' field '.'])
end

% Search for the corresponding parameter.
parameterFound = false;
for j = 1:size(par.(fields{fieldInd}),1)
    possibleNames = par.(fields{fieldInd}){j,1};
    if any(strcmpi(parameterName,possibleNames))
        parameterValue = par.(fields{fieldInd}){j,2};
        parameterFound = true;
        break;
    end
end
if ~parameterFound
    error(['Unable to find the default value for parameter called ' parameterName '.'])
end

%% On the fly parameters
% Overwrite the value if one of the parameters was provided by the user 
% when calling mainTracking.

% time resolution
possibleNames = par.global{1,1};
global provided_time_resolution
if any(strcmpi(parameterName,possibleNames)) && exist('provided_time_resolution','var') && ~isempty(provided_time_resolution)
    parameterValue = provided_time_resolution;
end

% space resolution
possibleNames = par.global{2,1};
global provided_space_resolution
if any(strcmpi(parameterName,possibleNames)) && exist('provided_space_resolution','var') && ~isempty(provided_space_resolution)
    parameterValue = provided_space_resolution;
end

% LoGratio
possibleNames = par.global{11,1};
global provided_LoGratio
if any(strcmpi(parameterName,possibleNames)) && exist('provided_LoGratio','var') && ~isempty(provided_LoGratio)
    parameterValue = provided_LoGratio;
end


end
```
