## Supplementary material for "Enhancer architecture and chromatin accessibility constrain phenotypic space during development": Script-MLP: iIndex.m

```
function output=iIndex(i,NDigits);

%Creates a string with the number i and a total of NDigits

if i<(10^NDigits)
    if i==0
        for j=1:NDigits
            output(j)='0';
        end
    else
        for j=1:NDigits
            if ((10^(j-1))<=i)&&(i<(10^j))
                output(1:NDigits-j)='0';
                output(end+1:end+j)=num2str(i);
            end
        end
    end
else
    output=num2str(i);
end

        
```
