## Supplementary material for "Enhancer architecture and chromatin accessibility constrain phenotypic space during development": Script-MLP: peakfinder.m

```
function varargout = peakfinder(x0, sel, thresh, extrema, includeEndpoints, interpolate)
%PEAKFINDER Noise tolerant fast peak finding algorithm
%   INPUTS:
%       x0 - A real vector from the maxima will be found (required)
%       sel - The amount above surrounding data for a peak to be,
%           identified (default = (max(x0)-min(x0))/4). Larger values mean
%           the algorithm is more selective in finding peaks.
%       thresh - A threshold value which peaks must be larger than to be
%           maxima or smaller than to be minima.
%       extrema - 1 if maxima are desired, -1 if minima are desired
%           (default = maxima, 1)
%       includeEndpoints - If true the endpoints will be included as
%           possible extrema otherwise they will not be included
%           (default = true)
%       interpolate - If true quadratic interpolation will be performed
%           around each extrema to estimate the magnitude and the
%           position of the peak in terms of fractional indicies. Note that
%           unlike the rest of this function interpolation assumes the
%           input is equally spaced. To recover the x_values of the input
%           rather than the fractional indicies you can do:
%           peakX = x0 + (peakLoc - 1) * dx
%           where x0 is the first x value and dx is the spacing of the
%           vector. Output peakMag to recover interpolated magnitudes.
%           See example 2 for more information.
%           (default = false)
%
%   OUTPUTS:
%       peakLoc - The indicies of the identified peaks in x0
%       peakMag - The magnitude of the identified peaks
%
%   [peakLoc] = peakfinder(x0) returns the indicies of local maxima that
%       are at least 1/4 the range of the data above surrounding data.
%
%   [peakLoc] = peakfinder(x0,sel) returns the indicies of local maxima
%       that are at least sel above surrounding data.
%
%   [peakLoc] = peakfinder(x0,sel,thresh) returns the indicies of local
%       maxima that are at least sel above surrounding data and larger
%       (smaller) than thresh if you are finding maxima (minima).
%
%   [peakLoc] = peakfinder(x0,sel,thresh,extrema) returns the maxima of the
%       data if extrema > 0 and the minima of the data if extrema < 0
%
%   [peakLoc] = peakfinder(x0,sel,thresh,extrema, includeEndpoints)
%       returns the endpoints as possible extrema if includeEndpoints is
%       considered true in a boolean sense
%
%   [peakLoc, peakMag] = peakfinder(x0,sel,thresh,extrema,interpolate)
%       returns the results of results of quadratic interpolate around each
%       extrema if interpolate is considered to be true in a boolean sense
%
%   [peakLoc, peakMag] = peakfinder(x0,...) returns the indicies of the
%       local maxima as well as the magnitudes of those maxima
%
%   If called with no output the identified maxima will be plotted along
%       with the input data.
%
%   Note: If repeated values are found the first is identified as the peak
%
% Example 1:
% t = 0:.0001:10;
% x = 12*sin(10*2*pi*t)-3*sin(.1*2*pi*t)+randn(1,numel(t));
% x(1250:1255) = max(x);
% peakfinder(x)
%
% Example 2:
% ds = 100;  % Downsample factor
% dt = .001; % Time step
% ds_dt = ds*dt; % Time delta after downsampling
% t0 = 1;
% t = t0:dt:5 + t0;
% x = 0.2-sin(0.01*2*pi*t)+3*cos(7/13*2*pi*t+.1)-2*cos((1+pi/10)*2*pi*t+0.2)-0.2*t;
% x(end) = min(x);
% x_ds = x(1:ds:end); % Downsample to test interpolation
% [minLoc, minMag] = peakfinder(x_ds, .8, 0, -1, false, true);
% minT = t0 + (minLoc - 1) * ds_dt; % Take into account 1 based indexing
% p = plot(t,x,'-',t(1:ds:end),x_ds,'o',minT,minMag,'rv');
% set(p(2:end), 'linewidth', 2); % Show the markers more clearly
% legend('Actual Data', 'Input Data', 'Estimated Peaks');
% Copyright Nathanael C. Yoder 2015

% Perform error checking and set defaults if not passed in
narginchk(1, 6);
nargoutchk(0, 2);

s = size(x0);
flipData =  s(1) < s(2);
len0 = numel(x0);
if len0 ~= s(1) && len0 ~= s(2)
    error('PEAKFINDER:Input','The input data must be a vector')
elseif isempty(x0)
    varargout = {[],[]};
    return;
end
if ~isreal(x0)
    warning('PEAKFINDER:NotReal','Absolute value of data will be used')
    x0 = abs(x0);
end

if nargin < 2 || isempty(sel)
    sel = (max(x0)-min(x0))/4;
elseif ~isnumeric(sel) || ~isreal(sel)
    sel = (max(x0)-min(x0))/4;
    warning('PEAKFINDER:InvalidSel',...
        'The selectivity must be a real scalar.  A selectivity of %.4g will be used',sel)
elseif numel(sel) > 1
    warning('PEAKFINDER:InvalidSel',...
        'The selectivity must be a scalar.  The first selectivity value in the vector will be used.')
    sel = sel(1);
end

if nargin < 3 || isempty(thresh)
    thresh = [];
elseif ~isnumeric(thresh) || ~isreal(thresh)
    thresh = [];
    warning('PEAKFINDER:InvalidThreshold',...
        'The threshold must be a real scalar. No threshold will be used.')
elseif numel(thresh) > 1
    thresh = thresh(1);
    warning('PEAKFINDER:InvalidThreshold',...
        'The threshold must be a scalar.  The first threshold value in the vector will be used.')
end

if nargin < 4 || isempty(extrema)
    extrema = 1;
else
    extrema = sign(extrema(1)); % Should only be 1 or -1 but make sure
    if extrema == 0
        error('PEAKFINDER:ZeroMaxima','Either 1 (for maxima) or -1 (for minima) must be input for extrema');
    end
end

if nargin < 5 || isempty(includeEndpoints)
    includeEndpoints = true;
end

if nargin < 6 || isempty(interpolate)
    interpolate = false;
end

x0 = extrema*x0(:); % Make it so we are finding maxima regardless
thresh = thresh*extrema; % Adjust threshold according to extrema.
dx0 = diff(x0); % Find derivative
dx0(dx0 == 0) = -eps; % This is so we find the first of repeated values
ind = find(dx0(1:end-1).*dx0(2:end) < 0)+1; % Find where the derivative changes sign

% Include endpoints in potential peaks and valleys as desired
if includeEndpoints
    x = [x0(1);x0(ind);x0(end)];
    ind = [1;ind;len0];
    minMag = min(x);
    leftMin = minMag;
else
    x = x0(ind);
    minMag = min(x);
    leftMin = min(x(1), x0(1));
end

% x only has the peaks, valleys, and possibly endpoints
len = numel(x);

if len > 2 % Function with peaks and valleys
    % Set initial parameters for loop
    tempMag = minMag;
    foundPeak = false;

    if includeEndpoints
        % Deal with first point a little differently since tacked it on
        % Calculate the sign of the derivative since we tacked the first
        %  point on it does not neccessarily alternate like the rest.
        signDx = sign(diff(x(1:3)));
        if signDx(1) <= 0 % The first point is larger or equal to the second
            if signDx(1) == signDx(2) % Want alternating signs
                x(2) = [];
                ind(2) = [];
                len = len-1;
            end
        else % First point is smaller than the second
            if signDx(1) == signDx(2) % Want alternating signs
                x(1) = [];
                ind(1) = [];
                len = len-1;
            end
        end
    end

    % Skip the first point if it is smaller so we always start on a
    %   maxima
    if x(1) >= x(2)
        ii = 0;
    else
        ii = 1;
    end

    % Preallocate max number of maxima
    maxPeaks = ceil(len/2);
    peakLoc = zeros(maxPeaks,1);
    peakMag = zeros(maxPeaks,1);
    cInd = 1;
    % Loop through extrema which should be peaks and then valleys
    while ii < len
        ii = ii+1; % This is a peak
        % Reset peak finding if we had a peak and the next peak is bigger
        %   than the last or the left min was small enough to reset.
        if foundPeak
            tempMag = minMag;
            foundPeak = false;
        end

        % Found new peak that was lager than temp mag and selectivity larger
        %   than the minimum to its left.
        if x(ii) > tempMag && x(ii) > leftMin + sel
            tempLoc = ii;
            tempMag = x(ii);
        end

        % Make sure we don't iterate past the length of our vector
        if ii == len
            break; % We assign the last point differently out of the loop
        end

        ii = ii+1; % Move onto the valley
        % Come down at least sel from peak
        if ~foundPeak && tempMag > sel + x(ii)
            foundPeak = true; % We have found a peak
            leftMin = x(ii);
            peakLoc(cInd) = tempLoc; % Add peak to index
            peakMag(cInd) = tempMag;
            cInd = cInd+1;
        elseif x(ii) < leftMin % New left minima
            leftMin = x(ii);
        end
    end

    % Check end point
    if includeEndpoints
        if x(end) > tempMag && x(end) > leftMin + sel
            peakLoc(cInd) = len;
            peakMag(cInd) = x(end);
            cInd = cInd + 1;
        elseif ~foundPeak && tempMag > minMag % Check if we still need to add the last point
            peakLoc(cInd) = tempLoc;
            peakMag(cInd) = tempMag;
            cInd = cInd + 1;
        end
    elseif ~foundPeak
        if x(end) > tempMag && x(end) > leftMin + sel
            peakLoc(cInd) = len;
            peakMag(cInd) = x(end);
            cInd = cInd + 1;
        elseif tempMag > min(x0(end), x(end)) + sel
            peakLoc(cInd) = tempLoc;
            peakMag(cInd) = tempMag;
            cInd = cInd + 1;
        end
    end

    % Create output
    if cInd > 1
        peakInds = ind(peakLoc(1:cInd-1));
        peakMags = peakMag(1:cInd-1);
    else
        peakInds = [];
        peakMags = [];
    end
else % This is a monotone function where an endpoint is the only peak
    [peakMags,xInd] = max(x);
    if includeEndpoints && peakMags > minMag + sel
        peakInds = ind(xInd);
    else
        peakMags = [];
        peakInds = [];
    end
end

% Apply threshold value.  Since always finding maxima it will always be
%   larger than the thresh.
if ~isempty(thresh)
    m = peakMags>thresh;
    peakInds = peakInds(m);
    peakMags = peakMags(m);
end

if interpolate && ~isempty(peakMags)
    middleMask = (peakInds > 1) & (peakInds < len0);
    noEnds = peakInds(middleMask);

    magDiff = x0(noEnds + 1) - x0(noEnds - 1);
    magSum = x0(noEnds - 1) + x0(noEnds + 1)  - 2 * x0(noEnds);
    magRatio = magDiff ./ magSum;

    peakInds(middleMask) = peakInds(middleMask) - magRatio/2;
    peakMags(middleMask) = peakMags(middleMask) - magRatio .* magDiff/8;
end

% Rotate data if needed
if flipData
    peakMags = peakMags.';
    peakInds = peakInds.';
end

% Change sign of data if was finding minima
if extrema < 0
    peakMags = -peakMags;
    x0 = -x0;
end

% Plot if no output desired
if nargout == 0
    if isempty(peakInds)
        disp('No significant peaks found')
    else
        figure;
        plot(1:len0,x0,'.-',peakInds,peakMags,'ro','linewidth',2);
    end
else
    varargout = {peakInds,peakMags};
end
```
