## Supplementary material for "Enhancer architecture and chromatin accessibility constrain phenotypic space during development": Script-MLP: saveNuclearProjection.m

```
function saveNuclearProjection(hisMat, file)

nFrames = size(hisMat, 3);

if max(hisMat(:)) < 255
    hisMat = uint8(hisMat);
else
    hisMat = uint16(hisMat);
end

for f = 1:nFrames
    
    image = hisMat(:, :, f);
    if f == 1
        imwrite(image,file);
    else
        imwrite(image,file,'WriteMode', 'append');
    end
     
end

end
```
