## Supplementary material for "Enhancer architecture and chromatin accessibility constrain phenotypic space during development": Script-MLP: findFlatFieldInformation.m

```
% Find the FF information
function [FFPaths, FFToUse, LSMFF] = findFlatFieldInformation(Folder)
  % The FF can be in the folder with the data or in the folder
  % corresponding to the day.
  D1 = dir([Folder, filesep, 'FF*.lsm']);
  D1 = [D1, dir([Folder, filesep, 'FF*.czi'])];
  D2 = dir([Folder, filesep, '..', filesep, 'FF*.lsm']);
  D2 = [D2, dir([Folder, filesep, '..', filesep, 'FF*.czi'])];

  FFPaths = {};
  for i = 1:length(D1)
    FFPaths{end + 1} = [Folder, filesep, D1(i).name];
  end

  for i = 1:length(D2)
    FFPaths{end + 1} = [Folder, filesep, '..', filesep, D2(i).name];
  end

  %Go through the FF files and see which one matches the pixel size
  %and image pixel number
  FFToUse = [];
  LSMFF = [];
  for i = 1:length(FFPaths)
    LSMFF = bfopen(FFPaths{i});
    LSMFFMeta = LSMFF{:, 4};
  
    if (LSMFFMeta.getPixelsPhysicalSizeX(0) == LSMFFMeta.getPixelsPhysicalSizeX(0)) & ...
      (LSMFFMeta.getPixelsSizeY(0) == LSMFFMeta.getPixelsSizeY(0)) &...
      (LSMFFMeta.getPixelsSizeX(0) == LSMFFMeta.getPixelsSizeX(0))
      FFToUse = [FFToUse, i];
    end
  end
end
```
