## Supplementary material for "Enhancer architecture and chromatin accessibility constrain phenotypic space during development": Script-MLP: getNDigits.m

```
function NDigits = getNDigits(NFrames, LSMIndex)
  % First, get the starting time. This is not accessible in the
  % OME format, so we need to pull it out from the original
  % Metadata

  if NFrames(LSMIndex) < 10
    NDigits = 1;
  elseif NFrames(LSMIndex) < 100
    NDigits = 2;
  elseif NFrames(LSMIndex) < 1000
    NDigits = 3;
  else
    error('This program cannot support more than 1000 frames. This can be easily fixed')
  end
end
```
