## Supplementary material for "Enhancer architecture and chromatin accessibility constrain phenotypic space during development": Script-MLP: getZeissFrameInfo.m

```
function [FrameInfo,AllLSMImages,NSlices, NPlanes, NFrames,Frame_Times,NChannels] = getZeissFrameInfo(RawDataFiles,NSeries,FrameInfo)

  AllLSMImages = cell(NSeries,1);
  Frame_Times = []; % Store the frame information
  for LSMIndex = 1:NSeries        
    Folder = RawDataFiles(LSMIndex).folder;
    %Load the file using BioFormats
    LSMImages = bfopen([Folder, filesep, RawDataFiles(LSMIndex).name]);

    % Extract the metadata for each series
    LSMMeta = LSMImages{:, 4}; % OME Metadata
    LSMMeta2 = LSMImages{:, 2}; % Original Metadata
    AllLSMImages{LSMIndex} = LSMImages{1};

    % Figure out the number of slices in each series
    NSlices(LSMIndex) = str2double(LSMMeta.getPixelsSizeZ(0));
    % Number of channels
    if LSMIndex == 1 %NL: assuming all series have same # of channels for consistency with LIF mode
      NChannels = LSMMeta.getChannelCount(0);
    end
    % Total number of planes acquired
    NPlanes(LSMIndex) = LSMMeta.getPlaneCount(0);
    % Finally, use this information to determine the number of frames in
    % each series
    NFrames(LSMIndex) = NPlanes(LSMIndex) / NSlices(LSMIndex) / NChannels;

    % Check that the acquisition wasn't stopped before the end of a
    % cycle. If that is the case, some images in the last frame will
    % be blank and we need to remove them.
    if sum(sum(LSMImages{1}{end,1})) == 0
      % Reduce the number of frames by one
      NFrames(LSMIndex) = NFrames(LSMIndex) - 1;
      % Reduce the number of planes by NChannels*NSlices
      NPlanes(LSMIndex) = NPlanes(LSMIndex) - NChannels * NSlices(LSMIndex);
    end

    NDigits = getNDigits(NFrames, LSMIndex);

    try
        StartingTime(LSMIndex) = obtainZeissStartingTime(Folder, LSMIndex, LSMMeta2, NDigits);
    catch
        StartingTime(LSMIndex) = obtainZeissStartingTime(Folder, LSMIndex, LSMMeta2, NDigits+1);
    end       
    
    % add times
    [ValueField, Frame_Times] = obtainZeissFrameTimes(LSMMeta, NSlices, LSMIndex, NPlanes, NChannels, StartingTime, Frame_Times);
    
    % update frame info
    [~, FrameInfo] = updateZeissFrameInfo(LSMIndex, NFrames, NSlices, FrameInfo, LSMMeta, Frame_Times, ValueField);
  end

  
```
