## Supplementary material for "Enhancer architecture and chromatin accessibility constrain phenotypic space during development": Script-MLP: getZeissFrameInfoFISH.m

```
function [FrameInfo,LSMImages,NSlices, NPlanes, ...
    NFrames,Frame_Times,NChannels] = getZeissFrameInfoFISH(RawDataFile,FrameInfo)

Frame_Times = []; % Store the frame information

%Load the file using BioFormats
LSMImages = bfopen(RawDataFile);

% Extract the metadata for each series
LSMMeta = LSMImages{:, 4}; % OME Metadata

% Figure out the number of slices in each series
NSlices = str2double(LSMMeta.getPixelsSizeZ(0));
% Number of channels
if 1 == 1 %NL: assuming all series have same # of channels for consistency with LIF mode
    NChannels = LSMMeta.getChannelCount(0);
end
% Total number of planes acquired
NPlanes = LSMMeta.getPlaneCount(0);
% Finally, use this information to determine the number of frames in
% each series
NFrames = NPlanes / NSlices / NChannels;

StartingTime = 0;

% add times
[ValueField, Frame_Times] = obtainZeissFrameTimes(LSMMeta, NSlices, 1, NPlanes, NChannels, StartingTime, Frame_Times);

% update frame info
[~, FrameInfo] = updateZeissFrameInfo(1, NFrames, NSlices, FrameInfo, LSMMeta, Frame_Times, ValueField);
end
```
