## Supplementary material for "Enhancer architecture and chromatin accessibility constrain phenotypic space during development": Script-MLP: obtainZeissChannels.m

```
%Figure out the different channels
function [coatChannel, histoneChannel, fiducialChannel] = obtainZeissChannels(Channel1, Channel2, ExperimentType)
  histoneChannel = 0;
  Channel1{1} = lower(Channel1{1});
  Channel2{1} = lower(Channel2{1});
  if (~strcmpi(ExperimentType, 'inputoutput'))
    if ~isempty(strfind(Channel1{1}, lower('MCP'))) || ~isempty(strfind(Channel1{1}, lower('PCP'))) || ~isempty(strfind(Channel1{1}, lower('PP7')))
      coatChannel = 1;
    elseif strfind(Channel1{1}, lower('His'))
      histoneChannel = 1;
    else
      error('LSM Mode error: Channel name not recognized. Check MovieDatabase')
    end

    if ~isempty(strfind(Channel2{1}, lower('MCP'))) || ~isempty(strfind(Channel2{1}, lower('PCP'))) || ~isempty(strfind(Channel2{1}, lower('PP7')))
      coatChannel = 2;
    elseif strfind(Channel2{1}, lower('His'))
      histoneChannel = 2;
    else
      error('LSM Mode error: Channel name not recognized. Check MovieDatabase')
    end
  else 
    if ~isempty(strfind(Channel1{1}, lower('MCP')))
      coatChannel = 1;
      histoneChannel = 1;
    elseif ~isempty(strfind(Channel2{1}, lower('MCP')))
      coatChannel = 2;
      histoneChannel = 2;
    end
  end

  fiducialChannel = histoneChannel;
end
```
