## Supplementary material for "Enhancer architecture and chromatin accessibility constrain phenotypic space during development": Script-MLP: obtainZeissFrameTimes.m

```
function [ValueField, Frame_Times] = obtainZeissFrameTimes(LSMMeta, NSlices, LSMIndex, NPlanes, NChannels, StartingTime, Frame_Times)
  % There seems to be some ambiguity with how information is being
  % pulled out of the metadata. This might be due to BioFormats
  % versioning. We'll check which case we have to provide
  % compatibility for both types of outcome. Whether we get the
  % "value" field or not is determined by the ValueField flag.
  ValueField = 1;
  try
    LSMMeta.getPlaneDeltaT(0,0).value
  catch
    ValueField = 0;
  end

  % Now get the time for each frame. We start the timer at the first time point
  % of the first data series.
  for j = 0:(NSlices(LSMIndex) * NChannels):(NPlanes(LSMIndex) - 1)
    if ~isempty(LSMMeta.getPlaneDeltaT(0, j))
      if ValueField                    
        Frame_Times = [Frame_Times, str2num(LSMMeta.getPlaneDeltaT(0, j).value) + StartingTime(LSMIndex) - StartingTime(1)];
      else
        Frame_Times = [Frame_Times, str2num(LSMMeta.getPlaneDeltaT(0, j)) + StartingTime(LSMIndex) - StartingTime(1)];
      end
    else  % If there's only one frame the DeltaT will be empty
      Frame_Times = [Frame_Times, StartingTime(LSMIndex) - StartingTime(1)];
    end
  end
end
```
