## Supplementary material for "Enhancer architecture and chromatin accessibility constrain phenotypic space during development": Script-MLP: obtainZeissStartingTime.m

```
function StartingTime = obtainZeissStartingTime(Folder, LSMIndex, LSMMeta2, NDigits)
  % Get the starting time of this acquisition
  % This is different if I have an LSM or CZI file
  LSMDir = dir([Folder, filesep, '*.lsm']);     %Zeiss confocal, old LSM format
  CZIDir = dir([Folder, filesep, '*.czi']);     %Zeiss confocal, new CZI format
  
  if ~isempty(LSMDir)
    StartingTime = LSMMeta2.get(['TimeStamp #', iIndex(1, NDigits)]); %SEANCHANGE
  elseif ~isempty(CZIDir)
%     TimeStampString = LSMMeta2.get('Global Information|Image|T|StartTime #1');
%     TimeStampStrings{LSMIndex} = TimeStampString;
%     %Get the number of days since 1/1/0000
%     TimeStamp = datetime(TimeStampString(1:19),'InputFormat','yyyy-MM-dd''T''HH:mm:ss');
%     %Get the milliseconds, note that we're not using them!
%     MilliSeconds = TimeStamp(21:end-1);
%     %Convert the time to seconds. We're ignoring the seconds.
%     StartingTime = datenum(TimeStamp)*24*60*60; %SEANCHANGE from StartingTime(LSMIndex)
    StartingTime = 0;
  else
    error('Wrong file format. The code should not have gotten this far.')
  end
end
```
