## Supplementary material for "Enhancer architecture and chromatin accessibility constrain phenotypic space during development": Script-MLP: processFlatFieldInformation.m

```
function processFlatFieldInformation(Prefix, OutputFolder, FFPaths, FFToUse, LSMFF)
  if length(FFToUse) > 1
    warning('Too many flat field images match the pixel and image size size');
    FFToUse = uigetfile('Select which flat field image to use');
  elseif length(FFToUse) == 0
    warning('No flat field image found');
    % harrypotel: Is this clear needed?
    clear LIFFF;
  else
    LSMFF = bfopen(FFPaths{FFToUse});

    % Find the channel with the highest counts
    for i = 1:size(LSMFF{1}, 1)
      MaxValue(i) = max(max(LSMFF{1}{i, 1}));
    end

    [Dummy, ChannelToUse] = max(MaxValue);
    imwrite(LSMFF{1}{ChannelToUse, 1}, [OutputFolder, filesep, Prefix,'_FF.tif']);
  end
end
```
