## Supplementary material for "Enhancer architecture and chromatin accessibility constrain phenotypic space during development": Script-MLP: processLSMData.m

```
% NL: Updated this drawing heavily from processLIFExportMode
% Last updated: 2020-09-03

function FrameInfo = processLSMData(Folder, RawDataFiles, FrameInfo,...
    Channels, ProjectionType, Prefix, OutputFolder,nuclearGUI,...
    skipExtraction)
    
    disp('Exporting movie file...');
    
    cleanupObj = onCleanup(@myCleanupFun);

    moviePrecision = 'uint16';
    hisPrecision = 'uint16';
    
    %Load the reference histogram for the fake histone channel
    load('ReferenceHist.mat', 'ReferenceHist');    
    
    % initialize FrameInfo
    FrameInfo = [];
        
    % get basic info
    NSeries = length(RawDataFiles);      
    
    if ~skipExtraction
      % This chunk makes FrameInfo                     
      [FrameInfo,AllLSMImages,NSlices, ~, NFrames,~,NChannels] ...
        = getZeissFrameInfo(RawDataFiles,NSeries,FrameInfo);
      
      % save FrameInfo
      liveExperiment = LiveExperiment(Prefix);
      save([liveExperiment.resultsFolder,filesep,'FrameInfo.mat'], 'FrameInfo')
      
      % this function exports tif z stacks
      exportTifStacks(AllLSMImages, 'LSM', NChannels, NFrames, NSlices, Prefix, ...
          moviePrecision, hisPrecision, nuclearGUI, ProjectionType, Channels, ReferenceHist)           
      
      % Look for flat field images
      [FFPaths, FFToUse, LSMFF] = findFlatFieldInformation(Folder);
      % Proceed accordingly
      processFlatFieldInformation(Prefix, OutputFolder, FFPaths, FFToUse, LSMFF);
      
      if nuclearGUI

        chooseAnaphaseFrames(...
            Prefix, 'ProjectionType', ProjectionType,...
            'ReferenceHist', ReferenceHist);

      end
    end
end
```
