## Supplementary material for "Enhancer architecture and chromatin accessibility constrain phenotypic space during development": Script-MLP: processZeissChannels.m

```
function processZeissChannels(channelIndex, Prefix, FrameRange, framesIndex, OutputFolder, BlankImage, NSlices, NChannels, LSMIndex, LSMImages)
  % First do the coat protein channel
  % Save the blank images at the beginning and end of the stack
  NameSuffix = ['_ch', iIndex(channelIndex, 2)];

  NewName = [Prefix, '_', iIndex(FrameRange(framesIndex), 3), '_z', iIndex(1, 2), NameSuffix, '.tif'];
  imwrite(BlankImage, [OutputFolder, filesep, NewName]);
  NewName = [Prefix, '_', iIndex(FrameRange(framesIndex), 3), '_z', iIndex(min(NSlices) + 2, 2), NameSuffix, '.tif'];
  imwrite(BlankImage, [OutputFolder, filesep, NewName]);
  
  %Copy the rest of the images
  slicesCounter = 1; %Counter for slices
  
  for slicesIndex = ((framesIndex - 1) * NSlices(LSMIndex) * NChannels(LSMIndex) + 1 + (channelIndex - 1)):...
    NChannels:(framesIndex * NSlices(LSMIndex)) * NChannels
    if slicesCounter <= min(NSlices)
      NewName = [Prefix, '_', iIndex(FrameRange(framesIndex), 3), '_z', iIndex(slicesCounter + 1, 2), NameSuffix, '.tif'];
      imwrite(LSMImages{slicesIndex, 1}, [OutputFolder, filesep, NewName]);
      slicesCounter = slicesCounter + 1;
    end
  end
end
```
