## Supplementary material for "Enhancer architecture and chromatin accessibility constrain phenotypic space during development": Script-MLP: processZeissConfocalLSMData.m

```
function FrameInfo = processZeissConfocalLSMData(Folder, D, FrameInfo, ExperimentType, Channel1, Channel2, ProjectionType,Prefix, OutputFolder)
  % What type of experiment do we have?

    [coatChannel, ~, fiducialChannel] = obtainZeissChannels(Channel1, Channel2, ExperimentType);

    NSeries = length(D);
    Frame_Times = []; % Store the frame information

    waitbarFigure = waitbar(0, 'Extracting LSM images');
    load('ReferenceHist.mat')
    for LSMIndex = 1:NSeries
      waitbar(LSMIndex / NSeries, waitbarFigure);
      
      %Load the file using BioFormats
      LSMImages = bfopen([Folder, filesep, D(LSMIndex).name]);
      % Extract the metadata for each series
      LSMMeta = LSMImages{:, 4}; % OME Metadata
      LSMMeta2 = LSMImages{:, 2}; % Original Metadata

      % Figure out the number of slices in each series
      NSlices(LSMIndex) = str2num(LSMMeta.getPixelsSizeZ(0));
      % Number of channels
      NChannels(LSMIndex) = LSMMeta.getChannelCount(0);
      % Total number of planes acquired
      NPlanes(LSMIndex) = LSMMeta.getPlaneCount(0);
      % Finally, use this information to determine the number of frames in
      % each series
      NFrames(LSMIndex) = NPlanes(LSMIndex) / NSlices(LSMIndex) / NChannels(LSMIndex);

      % Check that the acquisition wasn't stopped before the end of a
      % cycle. If that is the case, some images in the last frame will
      % be blank and we need to remove them.
      if sum(sum(LSMImages{1}{end,1})) == 0
        % Reduce the number of frames by one
        NFrames(LSMIndex) = NFrames(LSMIndex) - 1;
        % Reduce the number of planes by NChannels*NSlices
        NPlanes(LSMIndex) = NPlanes(LSMIndex) - NChannels(LSMIndex) * NSlices(LSMIndex);
      end

      NDigits = getNDigits(NFrames, LSMIndex);

      StartingTime(LSMIndex) = obtainZeissStartingTime(Folder, LSMIndex, LSMMeta2, NDigits); %SEANCHANGE
      [ValueField, Frame_Times] = obtainZeissFrameTimes(LSMMeta, NSlices, LSMIndex, NPlanes, NChannels, StartingTime, Frame_Times);
      [FrameRange, FrameInfo] = createZeissFrameInfo(LSMIndex, NFrames, NSlices, FrameInfo, LSMMeta, Frame_Times, ValueField);

      processZeissFrames(Prefix, ExperimentType, Channel1, Channel2,ProjectionType, OutputFolder, LSMImages, LSMIndex, FrameRange, NSlices, NChannels, coatChannel, fiducialChannel, ReferenceHist)
    end
  
    close(waitbarFigure);

    [FFPaths, FFToUse, LSMFF] = findFlatFieldInformation(Folder);
    processFlatFieldInformation(Prefix, OutputFolder, FFPaths, FFToUse, LSMFF);

end
```
