## Supplementary material for "Enhancer architecture and chromatin accessibility constrain phenotypic space during development": Script-MLP: processZeissFrames.m

```
function processZeissFrames(Prefix, ExperimentType, Channel1, Channel2,ProjectionType, OutputFolder, LSMImages, LSMIndex, FrameRange, NSlices, NChannels, coatChannel, fiducialChannel, ReferenceHist)
  % Create a blank image
  BlankImage = uint16(zeros(size(LSMImages{1}{1, 1})));

  % Redefine LSMImages as a cell. I think we need this so that the
  % parfor loop doesn't freak out.
  LSMImages = LSMImages{1};

  for framesIndex = 1:length(FrameRange)
    
    if ~strcmpi(ExperimentType, 'inputoutput')
      for channelIndex = 1:NChannels
         processZeissChannels(channelIndex, Prefix, FrameRange, framesIndex, OutputFolder, BlankImage, NSlices, NChannels, LSMIndex, LSMImages);
      end
     else 
      for channelIndex = 1:NChannels
        processZeissChannels(channelIndex, Prefix, FrameRange, framesIndex, OutputFolder, BlankImage, NSlices, NChannels, LSMIndex, LSMImages);
      end
    end

    if fiducialChannel
        %Now do His-RFP
        HisSlices = zeros([size(LSMImages{1, 1}, 1), size(LSMImages{1, 1}, 2), NSlices(LSMIndex)]);
        slicesCounter = 1;

        for slicesIndex = ((framesIndex - 1) * NSlices(LSMIndex) * NChannels(LSMIndex) + 1 + (fiducialChannel - 1)):NChannels(LSMIndex):...
          (framesIndex * NSlices(LSMIndex)) * NChannels(LSMIndex)

          HisSlices(:, :, slicesCounter) = LSMImages{slicesIndex, 1};
          slicesCounter = slicesCounter + 1;
        end
    end
    
    if fiducialChannel
        Projection = median(HisSlices, 3);
    end
    if strcmpi(ExperimentType, 'inputoutput')
      %YJK : Think about the case when there is no His channel,
      %and it is inputoutput mode or 1spot mode or 2spot2color.
      %We can use (MCP-mCherry) either inverted or raw
      %images to make fake histone images.
      if (isempty(strfind(Channel1{1}, 'His'))) && (isempty(strfind(Channel2{1}, 'His')))
        if (~isempty(strfind(Channel1{1}, 'NLS')))|(~isempty(strfind(Channel2{1}, 'NLS')))
          %don't invert with NLS-MCP-mCherry
        else
          %We don't want to use all slices. Only the center ones
          StackCenter = round((min(NSlices) - 1) / 2);
          StackRange = StackCenter - 1:StackCenter + 1;
          if strcmp(ProjectionType, 'medianprojection')
              Projection = median(HisSlices(:,:,StackRange), [], 3);
          else
              Projection = max(HisSlices(:,:,StackRange), [], 3);
          end
          %invert images to make nuclei bright
          Projection = imcomplement(Projection);
        end
        Projection = histeq(mat2gray(Projection), ReferenceHist);
        Projection = Projection * 10000;
      end
    end
    
    if fiducialChannel
        imwrite(uint16(Projection), [OutputFolder, filesep, Prefix, '-His_', iIndex(FrameRange(framesIndex), 3), '.tif']);
    end
end

end
```
