## Supplementary material for "Enhancer architecture and chromatin accessibility constrain phenotypic space during development": Script-MLP: updateZeissFrameInfo.m

```
function [FrameRange, FrameInfo] = updateZeissFrameInfo(LSMIndex, NFrames,...
        NSlices, FrameInfo, LSMMeta, Frame_Times, ValueField)
  %Save the information in FrameInfo
  if LSMIndex == 1
    FrameRange = 1:NFrames(LSMIndex);
  else
    FrameRange = (1:NFrames(LSMIndex)) + length(FrameInfo);
  end
  
  for i = FrameRange
    FrameInfo(i).LinesPerFrame = str2double(LSMMeta.getPixelsSizeY(0));
    FrameInfo(i).PixelsPerLine = str2double(LSMMeta.getPixelsSizeX(0));
    FrameInfo(i).NumberSlices = min(NSlices); % JP: because of z-padding, NL: added this back in, What was the issue?
    % feature, we need to set NumberSlices after we've processed all series
    FrameInfo(i).FileMode = 'LSMExport';

    if ValueField
      FrameInfo(i).PixelSize = str2num(LSMMeta.getPixelsPhysicalSizeX(0).value);
      FrameInfo(i).ZStep = str2double(LSMMeta.getPixelsPhysicalSizeZ(0).value);
    else
      FrameInfo(i).PixelSize = str2num(LSMMeta.getPixelsPhysicalSizeX(0));
      FrameInfo(i).ZStep = str2double(LSMMeta.getPixelsPhysicalSizeZ(0));
    end

    FrameInfo(i).Time = Frame_Times(i); % In seconds
  end
end
```
