## Supplementary material for "Enhancer architecture and chromatin accessibility constrain phenotypic space during development": Script-MLP: main.m

```
% Code in lib is from Hernan Garcia's lab mRNADynamics package (some of
% which is downloaded from elsewhere, e.g., Bio-Formats).
%
% Creator: Mindy Liu Perkins
% Last edited: MLP 2021-09-28

clc; clear; close all;

% Establish file structure.
proj = 'rhoNEE';
date = '2021-05-06';
rootdir = 'C:\Users\rafaelgalupa\Desktop\MLP_script\data';
data_dir = fullfile(rootdir,proj,date);

files = dir(fullfile(data_dir, '*.czi'));
filenames = {files.name};

% Create fish_seg_pool() object, which stores results from individual
% embryos in the same batch.  If fsp was saved previously it will be
% automatically loaded.
fsp = fish_seg_pool(proj,filenames{1}(1:end-4),date,rootdir);


%% EXTRACT MICROSCOPE IMAGES
% You can skip this if images were already extracted.

% Check if fsp has already been saved.  If not, set the histone channel (or
% nuclear-marked channel to use for segmentation) and extract the
% individual images.  Files are saved in rootdir/processed_data/proj/date
if ~exist(fullfile(fsp.savedir,[fsp.prefix '.mat']),'file')
    % Set the histone (nuclear marker) channel.
    fsp.histchannel = 2;
    
    % Extract individual embryos from batch CZI file.  You can ignore the
    % warning messages about individual CZI files not existing.  This step
    % creates TIFF files for channel projections in
    % rootdir/proj/date/tifstacks
    fsp.batch_extract('filepath',fullfile(data_dir,filenames{1}), ...
        'ProjectionType','maxprojection');
    % You can change ProjectionType (which applies only to the histone
    % channel for segmentation) to 'maxprojection', 'medianprojection', or
    % 'midsumprojection'.
end

% Save fsp so if you accidentally run the whole script again it won't
% extract the images a second time.  Note if you want to try a different
% projection type you will have to comment out the "if" statement above or
% delete the saved file, which will be found in rootdir/processed_data/proj
fsp.save();
disp('Finished extracting microscope images.')


%% PERFORM EMBRYO BY EMBRYO ANALYSIS
% Set the fluorescence channel with rho.
target_channel = 3;

% Options to redo various parts of the pipeline.
redo_all = true;
redo_segmentation = true;
redo_fluorescence = true;
redo_rotation = true;
redo_stripes = true;

for ii = 24:numel(fsp.daughter_fish) % for each embryo in the batch
    cur_fs = fsp.daughter_fish(ii);
    
    if isempty(cur_fs.skip)
        cur_fs.mark_skip();
    end
    
    if ~cur_fs.skip
        % Segment nuclei and manually identify outliers.
        if redo_all || redo_segmentation || isempty(cur_fs.nucleus_xy)
            cur_fs.segment('plot',false);
            cur_fs.identify_outliers();
        end
        
        % Calculate nuclear fluorescence.
        if redo_all || redo_segmentation || ...
                redo_fluorescence || isempty(cur_fs.fluo)
            cur_fs.calc_nuclear_fluorescence(target_channel);
        end
        
        % Rotate the embryo.
        if redo_all || redo_rotation || isempty(cur_fs.rotated_xy)
            cur_fs.s = [];
            cur_fs.rotate('long_axis_up',true);
        end
        
        % Identify and manually curate stripes.
        if redo_all || redo_stripes || isempty(cur_fs.ixs_in_stripe)
            cur_fs.identify_stripes('debug_plot',false, ...
                'interp_stripes',true,'manual_box',true,'overwrite',true, ...
                'defaultnstripes',2); close;
            cur_fs.choose_stripes_to_use();
            cur_fs.manual_correct_stripes();
        end
        
        % Calculate stripe width.
        cur_fs.get_stripe_width()
    end
    
    % Save the results.
    fsp.save(); % fsp.save('y') if you don't want to be prompted to overwrite
end

disp('Finished processing embryos.')


%% SAVE RESULTS IN TXT FILES

stripe_fluo = {};
stripe_width_in_micrometer = [];
for ii = 1:numel(fsp.daughter_fish) % for each embryo in the batch
    cur_fish = fsp.daughter_fish(ii);

    if isempty(cur_fish.ixs_in_stripe)
        disp(['Embryo ' num2str(ii) ' has not been processed--skipping...'])
    else
        cur_fluo = cur_fish.fluo;
        cur_ixs_in_stripe = cur_fish.ixs_in_stripe;
        sw_table = cur_fish.get_stripe_width();
        
        for ii_stripe = 1:size(cur_fish.ixs_in_stripe,2)
            if cur_fish.stripes_to_use(ii_stripe)
                stripe_fluo{end+1} = cur_fluo(cur_ixs_in_stripe(:,ii_stripe));
                if sum(cur_fish.stripes_to_use) == 1 && ii_stripe == 2
                    stripe_width_in_micrometer(end+1) = sw_table{2,1};
                else
                    stripe_width_in_micrometer(end+1) = sw_table{2,ii_stripe};
                end
            end
        end
    end
end

fluo_table = cell2table(stripe_fluo');
writetable(fluo_table,fullfile(fsp.savedir,'pooled_stripe_fluorescence_nuclei'), ...
    'WriteVariableNames',false);
writetable(array2table(stripe_width_in_micrometer), ...
    fullfile(fsp.savedir,'pooled_stripe_width_micrometer'), ...
    'WriteVariableNames',false);

disp(['Pooled data saved in ' fsp.savedir]);
```
