## Supplementary material for "Enhancer architecture and chromatin accessibility constrain phenotypic space during development": Script-MLP: embryo_heatmap.m

```
function embryo_heatmap(vals,opt,varargin)

vals = vals(:);
vals = vals - nanmin(vals);
ellipse_colors = (vals/nanmax(vals));
ellipse_colors(isnan(ellipse_colors)) = 0;

if strcmp(opt,'ellipse')
    ellipse(varargin{:},ellipse_colors,10,[],1.5,true);
elseif strcmp(opt,'scatter')
    scatter(varargin{:},ellipse_colors,'filled')
    colormap jet
else
    error(['unknown plot option ' opt]);
end

axis equal
axis off

end
```
