## Supplementary material for "Enhancer architecture and chromatin accessibility constrain phenotypic space during development": Script-MLP: fish_seg.m

```
% Object for analyzing a single FISH embryo.
%
% Creator: Mindy Liu Perkins
% Please note this code is not packaged for general use, so there are some
% unused/"test" functions, incomplete edge case detection, and some implied
% functionalities that are not fully implemented.
%
% Uses code from Hernan Garcia's mRNADynamics package.
% Uses code from Matlab Central File Exchange:
% - fit_ellipse: Ohad Gal (2009). fit_ellipse (https://www.mathworks.com/matlabcentral/fileexchange/3215-fit_ellipse), MATLAB Central File Exchange. Retrieved November 26, 2020.
% - InterX: NS (2010). Curve intersections (https://www.mathworks.com/matlabcentral/fileexchange/22441-curve-intersections), MATLAB Central File Exchange. Retrieved April 7, 2022. 
% - peakfinder (*debugged MLP):  Nathanael Yoder (2015). peakfinder(x0, sel, thresh, extrema, includeEndpoints, interpolate) (https://www.mathworks.com/matlabcentral/fileexchange/25500-peakfinder-x0-sel-thresh-extrema-includeendpoints-interpolate), MATLAB Central File Exchange. Retrieved November 25, 2020. 
% - violin: Holger Hoffmann (2015). Violin Plot (https://www.mathworks.com/matlabcentral/fileexchange/45134-violin-plot), MATLAB Central File Exchange. Retrieved May 8, 2021.

classdef fish_seg < handle
    
    properties
        s = [];             % image rotated extra pi/2? (y/n)
        histchannel = 1;    % index of histone channel
        fluochannels;
        
        fluo;               % fluorescence per nucleus
        spotfluo_ratio;     % ratio spot/nuclear fluorescence
        spotfluo;           % spot fluorescence
        
        outlier_ix;         % indices of outliers
        calibration_ix;     % indices of cells belonging to calibration (e.g., Fkh staining)
        
        nucleus_xy;         % coordinates of nuclei in original frame
        rotated_xy;         % coordinates of nuclei in rotated frame
        
        ixs_in_stripe;      % indices of nuclei belonging to stripes
        stripes_to_use;     % logical array indicating stripes to use
        ixs_in_calib;       % indices of nuclei belonging to calibration region
        
        nucdist;            % distance of nuclei from stripe centroids
        
        skip;
    end
    
    properties (SetAccess = protected)
        date;       % date of sample
        project;    % name of project (subfolder in datadir)
        prefix;     % sample identifier (filename sans .***)
        
        rootdir = 'C:\Users\mindylp\Documents\EMBL\Microscopy\';	% root directory for project (set default here)
        datadir;    % directory of original source data
        savedir;    % directory where final object is saved
        filepath;   % filepath to CZI file
        tifstackdir;    % directory where extracted TIFF stack is saved
        
        histprojection_type = 'midsumprojection';
        
        % for compatibility with livemRNA nuclear segmentation code
        FrameInfo;          % struct with image info
        
        nuclear_diameter;
        spot_radius;
        gaussfilt_radius;
    end
    
    methods
        
        % CONSTRUCTOR
        % Optional inputs: same as this.initialize
        function this = fish_seg(varargin)
            if ~isempty(varargin)
                this = this.initialize(varargin{:});
            end
        end
        
        
        % This function specifies where relevant files are located.  If a
        % MAT file exists in the directory for processed data that is
        % specified by the inputs, that object is loaded automatically.
        % Otherwise directories for raw and processed data are set and no
        % further actions are performed.
        %
        % INPUTS: Specify where relevant files are located.
        % - project = string with project name
        % - date = date of project
        % - prefix = name of individual FISH
        % - rootdir = root directory (optional; if desired set default directly under object properties above)
        %
        % For example, to process a file stored in
        % 'C:\Users\mindylp\Documents\EMBL\Microscopy\eve_tales\2021-05-15\ngt40x179eve46TALEA_eve-sna-FITC-555_DAPI_2021_05_15__10_23_53.czi'
        % the inputs are
        % - project = 'eve_tales'
        % - date = '2021-05-15'
        % - prefix = 'ngt40x179eve46TALEA_eve-sna-FITC-555_DAPI_2021_05_15__10_23_53'
        % - rootdir = 'C:\Users\mindylp\Documents\EMBL\Microscopy\'
        %
        % This function sets the following directories for saving TIFF file
        % outputs (from processing CZI files) and the object itself (with
        % call to save()):
        % - TIFF files : rootdir\project\date\tifstacks
        % - .mat file for object : rootdir\processed_data\project\date\
        function this = initialize(this,project,date,prefix,rootdir)
            this.project = project;
            this.date = date;
            this.prefix = prefix;
            if nargin > 4
                this.rootdir = rootdir;
            end
            
            this.datadir = fullfile(this.rootdir,this.project);
            this.savedir = fullfile(this.rootdir,'processed_data',this.project,this.date);
            this.filepath = fullfile(this.datadir,this.date,[this.prefix '.czi']);
            this.tifstackdir = fullfile(this.datadir,this.date,'tifstacks');
            
            if ~exist(this.filepath,'file')
                warning([this.filepath ' does not exist'])
            end
            
            this = this.load();
            this.fix_savedir(); % for backwards compatibility
        end
        
        
        % Extract the channels into separate TIFF stacks and generate
        % projection for histone channel.  TIFF files are saved in
        % this.tifstackdir
        function extract(this,varargin)
            disp('Extracting raw data...')
            
            ProjectionType = this.histprojection_type;
            series_ix = [];
            
            v = 1;
            while v <= numel(varargin)
                switch varargin{v}
                    case 'ProjectionType'
                        assert(numel(varargin) > v);
                        v = v+1;
                        ProjectionType = varargin{v};
                    case 'Series'
                        assert(numel(varargin) > v);
                        v = v+1;
                        series_ix = varargin{v};
                    otherwise
                        error(['unrecognized input option ' varargin{v}]);
                end
                v = v+1;
            end
            
            [this.FrameInfo,LSMImages,NSlices,~,NFrames,~,NChannels] ...
                = getZeissFrameInfoFISH(this.filepath,[]);
            
            if isempty(series_ix)
                if size(LSMImages,1) > 1
                    error('CZI file contains a series--please specify a series index or use a fish_seg_pool object with batch_extract()')
                else
                    series_ix = 1;
                end
            end
            
            LSMImage = LSMImages(series_ix);
            
            assert(NFrames == 1);

            this.gen_tif_stack(LSMImage,NChannels,NSlices,ProjectionType);
            
            disp('Done.')
        end
        
        
        % Segment nuclei from the histone channel.
        function segment(this,varargin)
            nuclear_cycle = 'd14';
            show_plot = true;
            
            v = 1;
            while v <= numel(varargin)
                switch varargin{v}
                    case 'nuclear_cycle'
                        assert(numel(varargin) > v);
                        v = v+1;
                        nuclear_cycle = varargin{v};
                    case 'nuclear_diameter'
                        assert(numel(varargin) > v);
                        v = v+1;
                        this.nuclear_diameter = varargin{v};
                    case 'plot'
                        assert(numel(varargin) > v);
                        v = v+1;
                        show_plot = varargin{v};
                    otherwise
                        error(['unrecognized input option' varargin{v}]);
                end
                v = v+1;
            end
            
            if isempty(this.FrameInfo)
                error('call this.extract() before segmentation');
            end
            
            if isempty(this.nuclear_diameter)
                this.nuclear_diameter = getDefaultParameters(this.FrameInfo,nuclear_cycle);
            end
            
            disp('Segmenting...')
            
            % load image to segment
            img2seg = this.get_img('his');
            
            % segment
            this.nucleus_xy = fliplr(findNuclei(this.FrameInfo,img2seg,this.nuclear_diameter));
            this.outlier_ix = false(size(this.nucleus_xy,1),1);
            
            if show_plot
                figure
                imagesc(img2seg); colormap gray
                hold on
                scatter(this.nucleus_xy(:,1),this.nucleus_xy(:,2),'r')
                hold off
                axis equal
                axis off
            end
            
            disp('Done.')
        end
        
        
        % Loads image associated with this in situ.
        % Options: opt = 'his' for the image used for segmentation, or an
        % integer channel number for the channel to be loaded.  If the
        % latter, optionally specify to project the image using the
        % specified projection type (default midsum).
        function img = get_img(this,opt,proj,proj_type,show_plot,newfig)
            if nargin < 6
                newfig = true;
                if nargin < 5
                    show_plot = false;
                    if nargin < 4
                        proj_type = this.histprojection_type;
                        if nargin < 3
                            proj = false;
                        end
                    end
                end
            end
            
            if isempty(proj_type)
                proj_type = this.histprojection_type;
            end
            
            if isnumeric(opt)
                img = read_tiff_stack(fullfile(this.tifstackdir,[this.prefix, ...
                    '_ch0' num2str(opt) '.tif']));
                if proj
                    img = calculateProjection(proj_type,size(img,3),img);
                end
            elseif strcmpi(opt,'his')
                img = double(imread(fullfile(this.tifstackdir,[this.prefix '-His.tif'])));
            else
                error('unknown image type')
            end
            
            if show_plot
                if size(img,3) > 1
                    warning('cannot plot image with multiple stacks');
                else
                    if newfig
                        figure
                    end
                    imagesc(img); axis equal; axis off; colormap gray;
                    imlims = [0,quantile(as_vector(img),0.95)];
                    caxis(imlims);
                end
            end
        end
        
        
        % Plots a histogram for the given image ('his' or channel) with the
        % indicated projection type (defaults to this.histprojection_type).
        function img_hist(this,opt,proj_type,figtitle)
            if nargin < 4
                figtitle = 'Image histogram';
                if nargin < 3
                    proj_type = this.histprojection_type;
                end
            end
            
            img = this.get_img(opt,true,proj_type);
            
            figure
            hist(as_vector(img));
            title(figtitle);
        end
        
        
        % Solicits user input as to whether this embryo is flagged to skip
        % during other analysis.
        function mark_skip(this)
            this.get_img('his',true,this.histprojection_type,true);
            
            opt = 'x';
            while ~isempty(opt) && ~strcmpi(opt,'s')
                opt = input('Press enter to use this embryo or s to skip it. ','s');
            end
            
            if strcmpi(opt,'s')
                this.skip = true;
            else
                this.skip = false;
            end
            
            close(gcf);
        end
        
        % Identifies outliers manually (default) or based on thresholding
        % for the specified property.
        function identify_outliers(this,varargin)
            manual = true;
            imgopt = 'his';
            overwrite = false;
            
            v = 1;
            while v <= numel(varargin)
                switch varargin{v}
                    case 'threshold'
                        manual = false;
                        assert(numel(varargin) > v+1);
                        properties = varargin(v+1:2:end);
                        threshes = varargin(v+2:2:end);
                        break;
                    case 'imgopt'
                        assert(numel(varargin) > v);
                        v = v+1;
                        imgopt = varargin{v};
                    case 'overwrite'
                        assert(numel(varargin) > v);
                        v = v+1;
                        overwrite = varargin{v};
                    otherwise
                        error(['unrecognized input option ' varargin{v}]);
                end
                v = v+1;
            end
            
            if overwrite
                new_outliers = false(size(this.outlier_ix));
            else
                new_outliers = this.outlier_ix;
            end
            
            if manual
                fig = figure;
                this.scatter_nuclei(imgopt,false,true);
                new_outliers = this.manual_select(fig,this.outlier_ix, ...
                    'include_outliers','selmarkerspec',{'y'},'newmarkerspec',{'w'});
                close(fig);
            else
                for ii_prop = 1:numel(properties)
                    cur_prop = properties{ii_prop};
                    cur_thresh = threshes{ii_prop};
                    
                    if isnumeric(cur_prop)
                        cur_vals = cur_prop;
                    else
                        cur_vals = this.(cur_prop);
                    end
                    
                    if numel(cur_vals) ~= numel(this.outlier_ix)
                        error('property must have the same number of entries as cells');
                    end
                    
                    new_outliers = new_outliers | (cur_vals < cur_thresh(1)) | ...
                        (cur_vals > cur_thresh(2));
                end
            end
            
            this.outlier_ix = new_outliers;
        end
        
        
        % Scatterplots nuclei on top of specified image ('his' for
        % segmentation image or channel number for other).  Optionally show
        % outliers in yellow or all segmented objects in the same color.
        function scatter_nuclei(this,imgopt,show_outliers,show_all,newfig)
            if nargin < 5
                newfig = false;
                if nargin < 4
                    show_all = false;
                    if nargin < 3
                        show_outliers = true;
                        if nargin < 2
                            imgopt = 'his';
                        end
                    end
                end
            end
            
            if isempty(this.nucleus_xy)
                error('call this.segment() before plotting nuclei')
            end
            
            if newfig
                figure
            end
            img = this.get_img(imgopt,true);
            imagesc(img); colormap gray; hold on;
            
            if show_all
                xy2plt = this.nucleus_xy;
            else
                xy2plt = this.nucleus_xy(~this.outlier_ix,:);
            end

            scatter(xy2plt(:,1),xy2plt(:,2),'g');
            if show_outliers
                scatter(this.nucleus_xy(this.outlier_ix,1),this.nucleus_xy(this.outlier_ix,2),'y');
            end
            hold off; axis equal; axis off
        end
        
        
        % Plots outliers in red over specified image; see this.get_img for
        % imgopt options.
        function fig = plot_outliers(this,imgopt)
            img = this.get_img(imgopt,true);
            
            fig = figure;
            imagesc(img); colormap gray
            hold on
            scatter(this.nucleus_xy(:,1),this.nucleus_xy(:,2),'r');
            hold off
            axis equal
            axis off
        end
        
        
        % Calculate the average fluorescence per nucleus separately for the
        % indicated channels.
        function calc_nuclear_fluorescence(this,channels,varargin)
            calc_spot_fluo = true;
            stacks_to_use = [];
            
            v = 1;
            while v <= numel(varargin)
                switch varargin{v}
                    case 'calc_spot_fluo'
                        assert(numel(varargin) > v);
                        v = v+1;
                        calc_spot_fluo = varargin{v};
                    case 'spot_radius'
                        if ~calc_spot_fluo
                            disp('calc_spot_fluo is set to false; spot_radius has no effect')
                        end
                        assert(numel(varargin) > v);
                        v = v+1;
                        this.spot_radius = varargin{v};
                    case 'gaussfilt_radius'
                        if ~calc_spot_fluo
                            disp('calc_spot_fluo is set to false; gaussfilt_radius has no effect')
                        end
                        assert(numel(varargin) > v);
                        v = v+1;
                        this.gaussfilt_radius = varargin{v};
                    case 'stacks'
                        assert(numel(varargin) > v);
                        v = v+1;
                        stacks_to_use = varargin{v};
                    otherwise
                        error(['unrecognized input option' varargin{v}]);
                end
                v = v+1;
            end
            
            if isempty(this.nucleus_xy)
                error('call this.segment() before calculating nuclear fluorescence');
            end
            
            nchannels = numel(channels);
            ncells = size(this.nucleus_xy,1);
            
            this.fluo = nan(ncells,nchannels);
            if calc_spot_fluo
                this.spotfluo = nan(ncells,nchannels);
                this.spotfluo_ratio = nan(ncells,nchannels);
            end
            for ii_channel = 1:nchannels
                cur_channel = channels(ii_channel);
                disp(['Calculating nuclear fluorescence for channel ' num2str(cur_channel) '...'])
                
                imgfluo = this.get_img(cur_channel);
                
                % calculate nuclear fluorescence
                nuclear_radius_in_pixels = floor(this.nuclear_diameter/(2*this.FrameInfo.PixelSize));
                nuclear_radius_in_pixels = nuclear_radius_in_pixels + mod(nuclear_radius_in_pixels,2);
                nuclear_mask = fspecial('disk',nuclear_radius_in_pixels) > 0;
                
                for ii = 1:ncells
                    box_rows = this.nucleus_xy(ii,2) + nuclear_radius_in_pixels*[-1,1];
                    box_cols = this.nucleus_xy(ii,1) + nuclear_radius_in_pixels*[-1,1];
                    
                    if ~(any(box_rows < 1) || any(box_cols < 1) || ...
                            any(box_rows > size(imgfluo,1)) || any(box_cols > size(imgfluo,2)))
                        cur_nuc = imgfluo(box_rows(1):box_rows(2),box_cols(1):box_cols(2),:);
                        
                        % nuclear fluorescence calculated as mean of max projection
                        if isempty(stacks_to_use)
                            cur_nuc = max(cur_nuc,[],3);
                        else
                            cur_nuc = max(cur_nuc(stacks_to_use),[],3);
                        end
                        this.fluo(ii,ii_channel) = mean(as_vector(cur_nuc(nuclear_mask)));
                        
                        if calc_spot_fluo
                            if isempty(this.spot_radius)
                                this.spot_radius = this.nuclear_diameter / 10;
                            end
                            
                            spot_radius_px = floor(this.spot_radius/this.FrameInfo.PixelSize);
                            
                            if isempty(this.gaussfilt_radius) || this.gaussfilt_radius == 0
                                this.gaussfilt_radius = this.spot_radius/this.FrameInfo.PixelSize/15;
                            end
                            
                            cur_nuc_gaussfilt = imgaussfilt(cur_nuc,this.gaussfilt_radius);

                            % first pass at spot brightness: mean fluorescence of pixels in
                            % original image surrounding brightest in Gaussian filtered image
                            [~,spotcen] = find_2d_max(cur_nuc_gaussfilt);
                            rowix = (1:size(cur_nuc_gaussfilt,1))';
                            colix = 1:size(cur_nuc_gaussfilt,2);
                            dist_from_center = sqrt((rowix-spotcen(1)).^2 + (colix-spotcen(2)).^2);
                            valid_px = (dist_from_center < spot_radius_px);
                            this.spotfluo(ii,ii_channel) = mean(as_vector(cur_nuc(valid_px)));
                            
                            this.spotfluo_ratio(ii,ii_channel) = ...
                                this.spotfluo(ii,ii_channel)/mean(cur_nuc(~valid_px & nuclear_mask));
                        end
                    end
                end
            end
            
            this.fluochannels = channels;
        end
        
        
        % Plots embryo heatmap with given values.  Specify rotated or
        % unrotated coordinates; defaults to rotated if available.
        function heatmap(this,vals,rotated,custom_xy,newfig)
            if nargin < 5
                newfig = false;
                if nargin < 4
                    custom_xy = [];
                    if nargin < 3
                        rotated = true;
                    end
                end
            end
            
            if ~isempty(custom_xy)
                xy = custom_xy;
            else
                if isempty(this.nucleus_xy)
                    error('call this.segment()');
                end
                
                xy = this.nucleus_xy;
                if rotated
                    if ~isempty(this.rotated_xy)
                        xy = this.rotated_xy;
                    end
                end
            end
            
            if ~isnumeric(vals)
                vals = this.(vals);
            end
            
            if numel(vals) ~= size(xy,1)
                error('property to plot must have same number of entries as nuclei');
            end
            
            nuclear_radius_in_pixels = floor(this.nuclear_diameter/(2*this.FrameInfo.PixelSize));
            
            if newfig
                figure
            end
            embryo_heatmap(vals(~this.outlier_ix),'scatter',xy(~this.outlier_ix,1), ...
                xy(~this.outlier_ix,2),nuclear_radius_in_pixels);
        end
        
        
        % Rotates embryo to lie along an axis.
        function rotate(this,varargin)
            debug_plot = false;  % display results of ellipse fit
            overwrite = false;
            repeat_rot = true;   % prompt user to rotate extra 90 or 180 degrees
            long_axis_up = false;   % true to rotate long axis vertical
            
            v = 1;
            while v <= numel(varargin)
                switch varargin{v}
                    case 'debug_plot'
                        assert(numel(varargin) > v);
                        v = v+1;
                        debug_plot = varargin{v};
                    case 'overwrite'
                        assert(numel(varargin) > v);
                        v = v+1;
                        overwrite = varargin{v};
                    case 'long_axis_up'
                        assert(numel(varargin) > v);
                        v = v+1;
                        long_axis_up = varargin{v};
                    otherwise
                        error(['unrecognized input option' varargin{v}]);
                end
                v = v+1;
            end
            
            cenx = this.nucleus_xy(~this.outlier_ix,1);
            ceny = this.nucleus_xy(~this.outlier_ix,2);
            
            % fit ellipse to boundaries of point cloud
            boundary_ixs = boundary(cenx,ceny);
            
            if debug_plot
                figure
                scatter(cenx,ceny);
                ellipse_t = fit_ellipse(cenx(boundary_ixs),ceny(boundary_ixs),gca);
                pause
                close
            else
                ellipse_t = fit_ellipse(cenx(boundary_ixs),ceny(boundary_ixs));
            end
            
            theta = -ellipse_t.phi;
            
            if overwrite
                this.s = [];
            end
            
            first_pass = true;
            % rotate embryo based on ellipse orientation
            Z = this.nucleus_xy;
            while repeat_rot
                R = [cos(theta),sin(theta); -sin(theta),cos(theta)];
                
                Z = (R*Z')';%(R*this.nucleus_xy')';
                
                if first_pass
                    % adjust if embryo was rotated with A-P axis up/down
                    if (diff(range(Z)) < 0 && long_axis_up) || ...
                            (diff(range(Z)) > 0 && ~long_axis_up)
                        Z = ([0, 1; -1, 0]*Z')';
                    end
                    first_pass = false;
                end
                
                % check rotation with user
                rotf = figure;
                this.heatmap(this.fluo,true,Z)
                
                while ~strcmpi(this.s,'y') && ~strcmpi(this.s,'n') && ~strcmpi(this.s,'h')
                    this.s = input('Rotate by additional 180 (y) or 90 (h) degrees? (y/h/n) ','s');
                end
                
                close(rotf);
                
                if strcmpi(this.s,'y')
                    repeat_rot = true;
                    theta = pi;
                    this.s = 'n';
                elseif strcmpi(this.s,'h')
                    repeat_rot = true;
                    theta = pi/2;
                    this.s = [];
                else
                    repeat_rot = false;
                end
            end
            
            this.rotated_xy = Z;
        end
        
        
        % Identify costain region for normalizing feature of interest.
        % Currently only supports fkh assuming that embryo has been rotated
        % with long axis horizontal.
        function identify_calibration_region(this,channel,varargin)
            if isempty(this.fluo)
                error('call this.calc_nuclear_fluorescence() before identifying calibration region')
            end
            
            if nargin < 2 || isempty(channel)
                channel = this.fluochannels(1);
            end
            
            assert(isnumeric(channel) && channel > 0 && channel < 3);
            
            ii_channel = find((this.fluochannels == channel));
            debug_plot = false;
            debug_plot_peaks = false;
            overwrite = false;
            stain = [];
            x_cutoff = 0.8;
            stripecutofffactor = 2;
            slope_cutoff = 1;
            manual = false;
            
            v = 1;
            while v <= numel(varargin)
                switch varargin{v}
                    case 'debug_plot'
                        assert(numel(varargin) > v);
                        v = v+1;
                        debug_plot = varargin{v};
                    case 'overwrite'
                        assert(numel(varargin) > v);
                        v = v+1;
                        overwrite = varargin{v};
                    case 'stain'
                        assert(numel(varargin) > v);
                        v = v+1;
                        stain = varargin{v};
                    case 'x_cutoff'
                        assert(numel(varargin) > v);
                        v = v+1;
                        x_cutoff = varargin{v};
                    case 'manual'
                        manual = true;
                    otherwise
                        error(['unrecognized input option' varargin{v}]);
                end
                v = v+1;
            end
            
            opt = [];
            if ~isempty(this.ixs_in_calib) && ~overwrite
                while ~strcmpi(opt,'y') && ~strcmpi(opt,'n')
                    opt = input('Overwrite existing calibration region? (y/n) ','s');
                end
            end
            
            if strcmpi(opt,'n')
                return;
            end
            
            if ~manual && isempty(stain)
                disp('No calibration region given--defaulting to Fkh posterior...');
                stain = 'fkh';
            end
            
            if manual
                fig = figure;
                this.heatmap('fluo');
                this.ixs_in_calib = this.manual_select(fig,[],'rotated','polygon', ...
                    'selmarkerspec',{50,'y'},'newmarkerspec',{50,'k'});
                close(fig);
                return;
            end
            
            if strcmpi(stain,'fkh')
                ncells = size(this.rotated_xy,1);
                this.ixs_in_calib = false(ncells,1);
                
                if ~isempty(this.ixs_in_stripe)
                    [sw,stripe_cenx,stripe_ceny] = this.get_stripe_width();
                    xcutoff = stripe_cenx(:,end) + sw(end);
                    valix = ~isnan(xcutoff);
                else
                    disp(['No stripes detected; using x-axis cutoff ' num2str(x_cutoff) '...']);
                    
                    xcutoff = min(this.rotated_xy(~this.outlier_ix,1)) + ...
                        range(this.rotated_xy(~this.outlier_ix,1))*x_cutoff;
                end
                
                exclude_ix = this.outlier_ix | (this.rotated_xy(:,1) < min(xcutoff));
                
                [~,~,bins_temp] = histcounts(this.rotated_xy(~exclude_ix,2));
                nbins = max(bins_temp);
                
                bins = nan(ncells,1);    % resize bins for outliers
                bins(~exclude_ix) = bins_temp;
                
                xbound = nan(nbins,1);
                ybound = nan(nbins,1);
                
                if ~isempty(this.ixs_in_stripe)
                    yqc = linspace(min(stripe_ceny(:)),max(stripe_ceny(:)),nbins);
                    xcutoff = interp1(stripe_ceny(valix,end),xcutoff(valix),yqc,...
                        'linear','extrap');
                else
                    xcutoff = xcutoff*ones(nbins,1);
                end
                
                % identify fluorescence peaks independently in each y-coordinate bin
                for ii_bin = 1:nbins
                    cur_ix = (bins == ii_bin) & ~exclude_ix;
                    
                    if ~isempty(xcutoff(ii_bin))
                        cur_ix = cur_ix & (this.rotated_xy(:,1) > xcutoff(ii_bin));
                    end
                    
                    if sum(cur_ix) > 2
                        cur_cenx = this.rotated_xy(cur_ix,1);
                        cur_ceny = this.rotated_xy(cur_ix,2);
                        cur_fluo = this.fluo(cur_ix,ii_channel);
                        
                        [cur_cenx,sort_ix] = sort(cur_cenx);
                        cur_ceny = cur_ceny(sort_ix);
                        cur_fluo = cur_fluo(sort_ix);
                        
                        % perform smoothing on A-P profile at current D-V coordinate
                        cur_fluo = smooth(cur_cenx,cur_fluo,0.5,'loess');
                        
                        if any(~isnan(cur_fluo))
                            trough_ix = peakfinder(cur_fluo,range(cur_fluo)/8,[],-1,true);
                            trough_ix = unique(trough_ix);
                            
                            if ~isempty(trough_ix)
                                [maxfluo,max_ix] = max(cur_fluo(min(trough_ix):end));
                                
                                trough_ix(trough_ix > min(max_ix)) = [];
                                if ~isempty(trough_ix)
                                    
                                    if debug_plot
                                        subplot(nbins,1,ii_bin)
                                        plot(cur_cenx,cur_fluo); hold on;
                                        scatter(cur_cenx(trough_ix),cur_fluo(trough_ix),'ro');
                                        hold off;
                                    end
                                    
                                    trough_ix = trough_ix(1);
                                    height_thresh = ...
                                        cur_fluo(trough_ix)+(maxfluo - cur_fluo(trough_ix))/2;
                                    
                                    tf = find(cur_fluo(trough_ix+1:end) > height_thresh,1);
                                    if ~isempty(tf)
                                        border_ix = trough_ix + tf;
                                        
                                        xbound(ii_bin) = cur_cenx(border_ix);
                                        ybound(ii_bin) = cur_ceny(border_ix);
                                        
                                        % assign nuclei to calibration region
                                        this.ixs_in_calib = this.ixs_in_calib | ...
                                            cur_ix & this.rotated_xy(:,1) > xbound(ii_bin);
                                    end
                                end
                            end
                        end
                    else
                        warning(['Only ' num2str(sum(cur_ix)) ' nuclei in bin ' num2str(ii_bin) ...
                            '...ignoring bin...']);
                    end
                end
                
                [xbound,ybound] = smooth_stripes(xbound,ybound,[],[],0.8);
                
                valix = ~isnan(xbound);
                yq = linspace(nanmin(this.rotated_xy(~exclude_ix,2)), ...
                    nanmax(this.rotated_xy(~exclude_ix,2)),100);
                xq = interp1(ybound(valix),xbound(valix),yq,'linear','extrap');
                
                if debug_plot
                    figure
                    this.heatmap(this.fluo);
                    hold on; plot(xq,yq,'r','LineWidth',3); hold off;
                end
            elseif strcmpi(stain,'sna')
                exclude_ix = [];%any(this.ixs_in_stripe,2);
                
                expectednstripes = 1;
                maxnstripes = 2;
                horizontal = true;
                interp_stripes = true;
                loess_smooth_factor = 0.3;
                
                [stripe_cenx,stripe_ceny,stripe_width_left,stripe_width_right,bins,nbins] = ...
                    this.find_stripes(ii_channel,expectednstripes,maxnstripes,horizontal, ...
                    exclude_ix,loess_smooth_factor,debug_plot,debug_plot_peaks, ...
                    stripecutofffactor,slope_cutoff);
                
                this.ixs_in_calib = assign_stripes(this,stripe_cenx,stripe_ceny, ...
                    stripe_width_left,stripe_width_right,stripecutofffactor,interp_stripes, ...
                    horizontal,bins,nbins);
                
                % throw out indices that do not have at least 2 of their
                % nearest neighbors also in the calibration region
                nn = this.nearest_n_neighbors(this.ixs_in_calib,6);
                
                inds_in_calib = find(this.ixs_in_calib);
                for ii = 1:numel(inds_in_calib)
                    if sum(this.ixs_in_calib(nn(:,ii))) <= 2
                        this.ixs_in_calib(inds_in_calib(ii)) = false;
                    end
                end
            else
                error('stain options are fkh and sna')
            end
            
            f = figure;
            this.plot_calibration_region(channel);
        end
        
        
        % FUNCTION FOR TESTING ONLY.  NOT FINALIZED.  USE WITH CAUTION.
        function ycoord = sna_testing(this)
            channel = 1;
            loess_smooth_factor = 0.5;
            exclude_ix = this.outlier_ix;
            
            [N,~,bins_temp] = histcounts(this.rotated_xy(~exclude_ix,2));
            nbins = max(bins_temp);
            
            bins = nan(size(this.rotated_xy,1),1);    % resize bins for outliers
            bins(~exclude_ix) = bins_temp;
            
            subplot(1,3,2)
            this.heatmap('fluo')
            axis on
            hold on
            % identify fluorescence peaks independently in each bin
            for ii = 1:nbins
                cur_bin = ii;
                
                if N(ii) > 4
                    cur_ix = (bins == cur_bin) & ~exclude_ix;
                    
                    cur_fluo = this.fluo(cur_ix,channel);
                    
                    cur_cenx = this.rotated_xy(cur_ix,1);
                    cur_ceny = this.rotated_xy(cur_ix,2);
                    
                    [cur_cenx,sort_ix] = sort(cur_cenx);
                    cur_ceny = cur_ceny(sort_ix);
                    cur_fluo = cur_fluo(sort_ix);
                    
                    % perform smoothing on appropriate profile at current
                    % coordinate
                    cur_fluo = smooth(cur_cenx,cur_fluo,loess_smooth_factor,'loess');
                    
                    if any(~isnan(cur_fluo))
                        ceny_mean(ii,1) = nanmean(cur_ceny);
                        fluo_mean(ii,1) = nanmean(cur_fluo);
                        
                        cur_col = [1 0 0]*(ii/nbins);
                        
                        subplot(1,3,1)
                        plot(cur_cenx,cur_fluo,'color',cur_col,'linewidth',1.5); hold on;
                        plot(cur_cenx,fluo_mean(ii)*ones(size(cur_cenx)),'color',cur_col,'linestyle','--','linewidth',1.5);
                        
%                         subplot(1,3,2)
%                         plot([min(this.rotated_xy(:,1)),max(this.rotated_xy(:,1))],ceny_mean(ii)*[1 1], ...
%                             'color',cur_col,'linewidth',1.5);
                        
%                         legend_entries{ii} = num2str(nanmean(cur_ceny));
                    end
                else
                    warning(['Only ' num2str(N(ii)) ' nuclei in bin ' num2str(ii) ...
                        '...ignoring bin...']);
                end
            end
            subplot(1,3,1)
            hold off;
            xlim([min(this.rotated_xy(:,1)),max(this.rotated_xy(:,1))])
%             l = legend(legend_entries);
%             set(l,'location','eastoutside')
            
            subplot(1,3,2)
            hold off
            axis tight
            
            subplot(1,3,3)
            plot(fluo_mean,ceny_mean);
            
            fluo_mean_smooth = fluo_mean;%smooth(ceny_mean,fluo_mean,0.1,'loess');
            
            [~,peak_ix] = max(fluo_mean);
            if isempty(peak_ix)
                warning('peak_ix is empty')
            end
            hold on;
            scatter(fluo_mean_smooth(peak_ix),ceny_mean(peak_ix),'rx');
            hold off;
            axis tight;
            
            subplot(1,3,2)
            hold on
            plot([min(this.rotated_xy(:,1)),max(this.rotated_xy(:,1))], ...
                ceny_mean(peak_ix)*[1,1],'r','linewidth',1.5);
            hold off
            
            set(gcf,'position',1e3*[4.1,0.3,1.5,0.42]);
            
            
            ycoord = ceny_mean(peak_ix);
        end
        
        
        % Automatically identifies stripes (default y-axis).
        % Optimized for late-stage eve stripes.  Basic method consists of:
        % (1) binning nuclei by DV coordinate
        % (2) for each DV bin (slice), smooth AP fluorescence profile and
        %     identify peaks (assumed to be stripe centers)
        % (3) align peaks across slices and perform automated smoothing
        % Recommended to use manual correction after automated
        % identification: call this.manual_correct_stripes()
        function identify_stripes(this,varargin)
            maxnstripes = [];
            expectednstripes = [];
            stripecutofffactor = 2;
            debug_plot = false;
            debug_plot_peaks = false;
            x_cutoff = [];
            y_cutoff = [];
            interp_stripes = false;
            slope_cutoff = [];
            overwrite = [];
            manual_select = false;
            defaultnstripes = 7;
            loess_smooth_factor = 0.1;
            horizontal = false;
            
            if isempty(this.fluochannels)
                error('call this.calc_nuclear_fluorescence before identifying stripes');
            end
            
            channel = this.fluochannels(1);
            
            v = 1;
            while v <= numel(varargin)
                switch varargin{v}
                    case 'expectednstripes'
                        assert(numel(varargin) > v);
                        v = v+1;
                        expectednstripes = varargin{v};
                        assert(isnumeric(expectednstripes) && expectednstripes > 0);
                    case 'maxnstripes'
                        assert(numel(varargin) > v);
                        v = v+1;
                        maxnstripes = varargin{v};
                        assert(isnumeric(maxnstripes) && maxnstripes > 0);
                    case 'stripecutofffactor'
                        assert(numel(varargin) > v);
                        v = v+1;
                        stripecutofffactor = varargin{v};
                        assert(isnumeric(stripecutofffactor) && stripecutofffactor > 0);
                    case 'debug_plot'
                        assert(numel(varargin) > v);
                        v = v+1;
                        debug_plot = varargin{v};
                        assert(islogical(debug_plot));
                    case 'debug_plot_peaks'
                        assert(numel(varargin) > v);
                        v = v+1;
                        debug_plot_peaks = varargin{v};
                        assert(islogical(debug_plot_peaks));
                    case 'channel'
                        assert(numel(varargin) > v);
                        v = v+1;
                        channel = varargin{v};
                        assert(isnumeric(channel) && ...
                            (channel == 1 || channel == 2 || channel == 3));
                    case 'x_cutoff'
                        assert(numel(varargin) > v);
                        v = v+1;
                        x_cutoff = varargin{v};
                        assert(isnumeric(x_cutoff) && numel(x_cutoff) == 2);
                    case 'y_cutoff'
                        assert(numel(varargin) > v);
                        v = v+1;
                        y_cutoff = varargin{v};
                        assert(isnumeric(y_cutoff) && numel(y_cutoff) == 2);
                    case 'interp_stripes'
                        assert(numel(varargin) > v);
                        v = v+1;
                        interp_stripes = varargin{v};
                        assert(islogical(interp_stripes));
                    case 'slope_cutoff'
                        assert(numel(varargin) > v);
                        v = v+1;
                        slope_cutoff = varargin{v};
                        assert(isnumeric(slope_cutoff) && slope_cutoff > 0);
                    case 'overwrite'
                        assert(numel(varargin) > v);
                        v = v+1;
                        overwrite = varargin{v};
                        assert(islogical(overwrite));
                    case 'manual_box'
                        assert(numel(varargin) > v);
                        v = v+1;
                        manual_select = varargin{v};
                        assert(islogical(manual_select));
                    case 'defaultnstripes'
                        assert(numel(varargin) > v);
                        v = v+1;
                        defaultnstripes = varargin{v};
                        assert(isnumeric(defaultnstripes) && defaultnstripes > 0);
                    case 'vertical'
                        horizontal = false;
                    case 'horizontal'
                        horizontal = true;
                    case 'loess'
                        assert(numel(varargin) > v);
                        v = v+1;
                        loess_smooth_factor = varargin{v};
                        assert(isnumeric(loess_smooth_factor) && loess_smooth_factor > 0);
                    otherwise
                        error(['unrecognized input option ' varargin{v}]);
                end
                v = v+1;
            end
            
            if ~isempty(this.ixs_in_stripe) && isempty(overwrite)
                opt = [];
                while ~strcmpi(opt,'y') && ~strcmpi(opt,'n')
                    opt = input('Stripes already identified--overwrite? (y/n) ','s');
                end
                
                if strcmpi(opt,'y')
                    overwrite = true;
                else
                    overwrite = false;
                end
            end
            
            if ~overwrite
                return;
            end
            
            if isempty(this.rotated_xy)
                error('call this.rotate() before identifying stripes')
            end
            
            if isempty(x_cutoff)
                x_cutoff = [0.2,0.9];
            end
            
            if isempty(y_cutoff)
                y_cutoff = [0,1];
            end
            
            ii_channel = find(this.fluochannels == channel);
            
            exclude_ix = this.outlier_ix;
            
            if manual_select
                disp('Note: manual select overrides x- and y-cutoff.');
                
                fig = figure;
                this.heatmap('fluo');
                selected_ix = this.manual_select(fig,[],'rotated','box','newmarkerspec',{'ko'});
                close(fig);
                exclude_ix = exclude_ix | ~selected_ix;
                
                curnstripe = 0;
                while ~(curnstripe >= 1)
                    curnstripe = input(['Enter expected number of stripes in this region ' ...
                        '(or press enter to take default ' num2str(defaultnstripes) '): ']);
                    
                    if isempty(curnstripe)
                        break
                    end
                end
                
                if ~isempty(curnstripe)
                    expectednstripes = curnstripe;
                    maxnstripes = expectednstripes + 1;
                end
            else
                xlen = range(this.rotated_xy(~this.outlier_ix,1));
                xstart = min(this.rotated_xy(~this.outlier_ix,1));
                ylen = range(this.rotated_xy(~this.outlier_ix,2));
                ystart = min(this.rotated_xy(~this.outlier_ix,2));
                exclude_ix = exclude_ix | ...
                    (this.rotated_xy(:,1) < (xstart + x_cutoff(1)*xlen)) | ...
                    (this.rotated_xy(:,1) > (xstart + x_cutoff(2)*xlen)) | ...
                    (this.rotated_xy(:,2) < (ystart + y_cutoff(1)*ylen)) | ...
                    (this.rotated_xy(:,2) > (ystart + y_cutoff(2)*ylen));
            end
            
            if isempty(expectednstripes)
                disp(['No expected number of stripes given--defaulting to ' ...
                    num2str(defaultnstripes) '...']);
                expectednstripes = defaultnstripes;
            end
            
            if isempty(maxnstripes)
                maxnstripes = expectednstripes + 1;
            end
            
            stripecutofffactor = 2;
            slope_cutoff = 1;
            
            [stripe_cenx,stripe_ceny,stripe_width_left,stripe_width_right,bins,nbins] = ...
                this.find_stripes(ii_channel,expectednstripes,maxnstripes, ...
                horizontal,exclude_ix,loess_smooth_factor,debug_plot,debug_plot_peaks, ...
                stripecutofffactor,slope_cutoff);
            
            this.ixs_in_stripe = assign_stripes(this,stripe_cenx,stripe_ceny, ...
                stripe_width_left,stripe_width_right,stripecutofffactor,interp_stripes, ...
                horizontal,bins,nbins);
            
            this.stripes_to_use = true(size(this.ixs_in_stripe,2),1);
            
            f = figure;
            this.plot_stripes(channel,'show_legend',true); axis equal;
            curpos = get(f,'Position');
            set(f,'Position',[900,100,curpos(3:4)]);
        end
        
        
        % Toggle which stripes to use.  Call with no arguments to visualize
        % embryo and request user input, or with one argument containing
        % numeric or logical indices to set to valid.
        function choose_stripes_to_use(this,ixs_to_use)
            if nargin > 1
                this.stripes_to_use = false(1,size(this.ixs_in_stripe,2));
                this.stripes_to_use(ixs_to_use) = true;
                return;
            end
            
            nstripes = size(this.ixs_in_stripe,2);
            
            if nstripes > 0
                if isempty(this.stripes_to_use)
                    this.stripes_to_use = true(1,size(this.ixs_in_stripe,2));
                end
                
                f = figure;
                curpos = get(f,'Position');
                set(f,'Position',[900,100,curpos(3:4)]);
                g = this.plot_stripes([],'show_legend',true,'hide_nonstripe'); axis equal;
                
                while true
                    disp(['Current valid stripes are ' num2str(find(this.stripes_to_use(:)')) ' of ' ...
                        num2str(nstripes) ' total.']);
                    opt = input('Enter the number of the stripe to toggle or enter to end: ');
                    
                    if isempty(opt)
                        close(f);
                        return;
                    end
                    
                    if opt < 1 || opt > nstripes
                        warning(['Stripe number must be between 1 and ' num2str(nstripes) '.']);
                    else
                        this.stripes_to_use(opt) = ~this.stripes_to_use(opt);
                        
                        figure(f);
                        delete(g);
                        hold on;
                        g = this.plot_stripes([],'show_img',false,'show_legend',true,'hide_nonstripe');
                        hold off;
                    end
                end
            else
                disp('Call this.identify_stripes()');
            end
        end
        
        
        % Estimates stripe width from identified nuclei.  First bins by
        % y-coordinate and calculates centroids for each stripe segment,
        % then fits a line between each centroid and calculates the spread
        % of nuclei in the direction perpendicular to that line.  Final
        % estimate obtained as the mean of segment widths.
        function [sw_table,stripe_cenx,stripe_ceny] = get_stripe_width(this,opt)
            [~,~,bins_temp] = histcounts(this.rotated_xy(~this.outlier_ix,2));
            nbins = max(bins_temp);
            
            bins = nan(size(this.rotated_xy,1),1);    % resize bins for outliers
            bins(~this.outlier_ix) = bins_temp;
            
            nstripes = size(this.ixs_in_stripe,2);
            
            % calculate centroids within each bin
            stripe_cenx = nan(nbins,nstripes);
            stripe_ceny = nan(nbins,nstripes);
            for ii_stripe = 1:nstripes
                for jj_bin = 1:nbins
                    cur_x = this.rotated_xy(this.ixs_in_stripe(:,ii_stripe) & bins == jj_bin,1);
                    cur_y = this.rotated_xy(this.ixs_in_stripe(:,ii_stripe) & bins == jj_bin,2);
                    stripe_cenx(jj_bin,ii_stripe) = nanmean(cur_x);
                    stripe_ceny(jj_bin,ii_stripe) = nanmean(cur_y);
                end
            end
            
            sw = nan(nbins-1,nstripes);
            for ii_stripe = 1:nstripes
                for jj_stretch = 1:nbins-1
                    % get line connecting two centroids
                    xcoords = stripe_cenx(jj_stretch:jj_stretch+1,ii_stripe);
                    ycoords = stripe_ceny(jj_stretch:jj_stretch+1,ii_stripe);
                    
                    % get nuclei between centroids
                    cur_xy = this.rotated_xy(this.ixs_in_stripe(:,ii_stripe),:);
                    cur_xy = cur_xy(cur_xy(:,2) > ycoords(1) & ...
                        cur_xy(:,2) < ycoords(2),:);
                    
                    if ~isempty(cur_xy)
                        % project coordinates of contained nuclei onto line
                        % perpendicular to segment between centroids,
                        % intersecting midpoint of segment
                        mp = [mean(xcoords),mean(ycoords)];
                        theta = -tan(diff(xcoords)/diff(ycoords));
                        R = [cos(theta),sin(theta);-sin(theta),cos(theta)];
                        rot_xy = (cur_xy - [mp(1),mp(2)])*R';
                        sw(jj_stretch,ii_stripe) = range(rot_xy(:,1));
                    end
                end
            end
            
            sw = nanmean(sw,1); % in units of pixels
            sw_um = sw * this.FrameInfo.PixelSize;  % in units of um

            % in units of internuclear distance
            dists = sqrt((this.nucleus_xy(:,1) - this.nucleus_xy(:,1)').^2 + ...
                (this.nucleus_xy(:,2) - this.nucleus_xy(:,2)').^2);
            dists = dists + diag(nan(numel(this.outlier_ix),1));      
            dists(this.outlier_ix,:) = nan;
            dists(:,this.outlier_ix) = nan;
            dists = min(dists,[],1);
            internuc_dist = nanmedian(dists); % distance from midpoint to midpoint
            sw_inuc = sw/internuc_dist;
            
            % as fraction of embryo length/width
            approx_embryo_x = range(this.rotated_xy(~this.outlier_ix,1));
            approx_embryo_y = range(this.rotated_xy(~this.outlier_ix,2));
            if approx_embryo_y > approx_embryo_x
                sw_ap = sw/approx_embryo_y;
                sw_dv = sw/approx_embryo_x;
            else
                sw_ap = sw/approx_embryo_x;
                sw_dv = sw/approx_embryo_y;
            end
            
            % in units of nuclear diameter
            sw_nucd = sw_um/getDefaultParameters(this.FrameInfo,'d14');
            
            vnamefun = @(x) ['stripe' num2str(x)];
            
            sw_table = array2table([sw;sw_um;sw_ap;sw_dv;sw_inuc;sw_nucd]);
            sw_table.Properties.RowNames = {'pixels','micrometers', ...
                'fraction embryo length (A/P)','fraction embryo width (D/V)', ...
                'measured internuclear distances','nuclear diameters (nc14)'};
            sw_table.Properties.VariableNames = cellfun(vnamefun,num2cell(1:numel(sw)),'UniformOutput',false);
            sw_table = sw_table(:,this.stripes_to_use);
        end
        
        
        % Calculates minimum distance from all nuclei to midline of valid
        % stripes (approximated from centroids).
        function calc_dist_from_stripe_centroid(this,opt,channel)
            if nargin < 3
                if ~isempty(this.fluochannels)
                    channel = this.fluochannels(1);
                else
                    channel = 1;
                end
                
                if nargin < 2
                    opt = 'normalized';
                end
            end
            
            if ~strcmpi(opt,'raw') && ~strcmpi(opt,'normalized')
                error('options are raw and normalized');
            end
            
            ii_channel = find(this.fluochannels == channel);
            
            if isempty(ii_channel)
                disp('warning: channel not found...defaulting to index 1');
                ii_channel = 1;
            end
            
            [~,~,bins_temp] = histcounts(this.rotated_xy(~this.outlier_ix,2));
            nbins = max(bins_temp);
            
            bins = nan(size(this.rotated_xy,1),1);    % resize bins for outliers
            bins(~this.outlier_ix) = bins_temp;
            
            nstripes = size(this.ixs_in_stripe,2);
            
            % calculate centroids within each bin
            stripe_cenx = nan(nbins,nstripes);
            stripe_ceny = nan(nbins,nstripes);
            for ii_stripe = 1:nstripes
                if this.stripes_to_use(ii_stripe)
                    for jj_bin = 1:nbins
                        cur_x = this.rotated_xy(this.ixs_in_stripe(:,ii_stripe) & bins == jj_bin,1);
                        cur_y = this.rotated_xy(this.ixs_in_stripe(:,ii_stripe) & bins == jj_bin,2);
                        stripe_cenx(jj_bin,ii_stripe) = nanmean(cur_x);
                        stripe_ceny(jj_bin,ii_stripe) = nanmean(cur_y);
                    end
                end
            end
            
            this.nucdist = nan(size(this.rotated_xy,1),nstripes);
            for ii_stripe = 1:nstripes
                if this.stripes_to_use(ii_stripe)
                    for jj_stretch = 1:nbins-1
                        % get line connecting two centroids
                        xcoords = stripe_cenx(jj_stretch:jj_stretch+1,ii_stripe);
                        ycoords = stripe_ceny(jj_stretch:jj_stretch+1,ii_stripe);
                        
                        % estimate min distance to each line segment
                        
                        % get nuclei between centroids
                        cur_xy = this.rotated_xy;
                        cur_ix = cur_xy(:,2) > ycoords(1) & cur_xy(:,2) < ycoords(2);
                        cur_xy = cur_xy(cur_ix,:);
                        
                        if ~isempty(cur_xy)
                            % project coordinates of contained nuclei onto line
                            % perpendicular to segment between centroids,
                            % intersecting midpoint of segment
                            mp = [mean(xcoords),mean(ycoords)];
                            theta = -tan(diff(xcoords)/diff(ycoords));
                            R = [cos(theta),sin(theta);-sin(theta),cos(theta)];
                            rot_xy = (cur_xy - [mp(1),mp(2)])*R';
                            this.nucdist(cur_ix,ii_stripe) = rot_xy(:,1);
                        end
                    end
                end
            end
            
            if strcmpi(opt,'normalized')
                approx_embryo_length = range(this.rotated_xy(~this.outlier_ix,1));
                this.nucdist = this.nucdist/approx_embryo_length;
            end
        end
        
        
        % Generates violin plots for fluorescence at different distances
        % from stripe midpoints.
        function [fluo_for_violinplot,binmids] = violinplot_by_dist_from_stripe_centroid(this,opt,varargin)
            col = [0.2,0.2,1];
            channel = [];
            xmax = [];
            overlay = false;
            stripe_ids = [];
            plot_single = false;
            
            v = 1;
            while v <= numel(varargin)
                switch varargin{v}
                    case 'xmax'
                        assert(v+1 <= numel(varargin));
                        xmax = varargin{v+1};
                        v = v+1;
                    case 'overlay'
                        overlay = true;
                    case 'no_overlay'
                        overlay = false;
                    case 'color'
                        assert(v+1 <= numel(varargin));
                        col = varargin{v+1};
                        v = v+1;
                    case 'channel'
                        assert(v+1 <= numel(varargin));
                        channel = varargin{v+1};
                        v = v+1;
                    case 'stripes'
                        assert(v+1 <= numel(varargin));
                        stripe_ids = varargin{v+1};
                        v = v+1;
                    case 'single'
                        plot_single = true;
                    otherwise
                        error(['unrecognized option ' varargin{v}]);
                end
                v = v+1;
            end
            
            if nargin < 2
                opt = 'normalized';
            end
            
            if ~strcmpi(opt,'raw') && ~strcmpi(opt,'normalized')
                error('options are raw and normalized');
            end
            
            ii_channel = find(this.fluochannels == channel);
            
            if isempty(ii_channel)
                disp('warning: channel not found...defaulting to index 1');
                ii_channel = 1;
            end
            
            xlab = 'distance from stripe';
            
            if strcmpi(opt,'normalized')
                xlab = [xlab, ' (normalized)'];
                if isempty(xmax)
                    xmax = 0.2;
                end
            elseif isempty(xmax)
                xmax = inf;
            end
            
            nstripes = size(this.ixs_in_stripe,2);
            
            val_ix = ~this.outlier_ix & ~this.ixs_in_calib;
            dists_for_plot = abs(this.nucdist(val_ix,:));
            fluo_for_plot = this.fluo(val_ix);
            spotfluo_for_plot = this.spotfluo(val_ix);
            fluodiff_for_plot = spotfluo_for_plot - nanmedian(fluo_for_plot);
            
            titles = {'Average fluorescence','Spot fluorescence','Spot fluo - median avg fluo across all nuclei'};
            if ~isempty(this.ixs_in_calib)
                disp('Normalizing fluorescence values to average of nuclei in calibration region...');
                calfactor = nanmean(this.fluo(this.ixs_in_calib,ii_channel));
                fluo_for_plot = fluo_for_plot/calfactor;
                spotfluo_for_plot = spotfluo_for_plot/calfactor;
                fluodiff_for_plot = fluodiff_for_plot/calfactor;
                titles{1} = 'Average fluorescence (normalized)';
                titles{2} = 'Spot fluorescence (normalized)';
            end
            
            if ~overlay
                figure
            end
            
            if ~isempty(stripe_ids)
                dists_for_violinplot = min(dists_for_plot(:,stripe_ids),[],2);
            else
                dists_for_violinplot = min(dists_for_plot,[],2);
            end

            [~,edges,bins] = histcounts(dists_for_violinplot,10,'BinLimits',[0,xmax]);
            nbins = max(bins);
            
            fluo_for_violinplot = cell(1,nbins);
            spotfluo_for_violinplot = cell(1,nbins);
            fluodiff_for_violinplot = cell(1,nbins);
            for ii = 1:nbins
                fluo_for_violinplot{ii} = fluo_for_plot(bins == ii);
                spotfluo_for_violinplot{ii} = spotfluo_for_plot(bins == ii);
                fluodiff_for_violinplot{ii} = fluodiff_for_plot(bins == ii);
            end
            
            binmids = (edges(1:end-1)+edges(2:end))/2;
            xlabs = cellfun(@num2str,num2cell(binmids),'UniformOutput',false);
            
            if ~plot_single
                subplot(3,1,1)
            end
            violin(fluo_for_violinplot,'xlabel',xlabs,'edgecolor',col,'facealpha',0,'mc','none','medc',col);%'facecolor',col,'edgecolor','none');
            legend off;
            title(titles{1});
            ylim([0.25 1]);
            
            if ~plot_single
                subplot(3,1,2)
                violin(spotfluo_for_violinplot,'xlabel',xlabs,'edgecolor',col,'facealpha',0,'mc','none','medc',col);%'facecolor',col,'edgecolor','none');
                legend off;
                title(titles{2});
                ylim([0.25,2]);
                
                subplot(3,1,3)
                violin(fluodiff_for_violinplot,'xlabel',xlabs,'edgecolor',col,'facealpha',0,'mc','none','medc',col);%'facecolor',col,'edgecolor','none');
                legend off;
                title(titles{3});
                ylim([-0.25,0.75]);
                
                pos = get(gcf,'Position');
                set(gcf,'Position',[300,200,670,580]);
            else
                pos = get(gcf,'Position');
                set(gcf,'Position',[300,200,670,580/3]);
            end
            
            xlabel('distance from stripe midpoint')
        end
        
        
        % Plot calibration region on top of fluorescence image.
        function g = plot_calibration_region(this,channel,varargin)
            show_img = true;
            
            v = 1;
            while v <= numel(varargin)
                switch varargin{v}
                    case 'show_img'
                        assert(numel(varargin) > v);
                        v = v+1;
                        show_img = varargin{v};
                    otherwise
                        error(['unrecognized input option' varargin{v}]);
                end
                v = v+1;
            end
            
            if isempty(channel)
                channel = this.fluochannels(1);
            end
            
            if isempty(this.ixs_in_calib)
                error('call this.identify_calibration_region() before plotting');
            end
            
            if show_img
                imgfluo = this.get_img(channel);
                imagesc(max(imgfluo,[],3)); axis off; colormap gray;
                hold on;
            end
            
            g = scatter(this.nucleus_xy(this.ixs_in_calib,1), ...
                this.nucleus_xy(this.ixs_in_calib,2),'r');
            hold off;
            axis equal;
        end
        
        
        % Scatter stripes on top of fluorescence image.
        function g = plot_stripes(this,channel,varargin)
            if nargin < 2 || isempty(channel)
                channel = this.fluochannels(1);
            end
            
            show_img = true;
            show_nonstripe = true;
            stripes_to_plot = [];
            plot_leg = false;
            
            v = 1;
            while v <= numel(varargin)
                switch varargin{v}
                    case 'show_img'
                        assert(numel(varargin) > v);
                        v = v+1;
                        show_img = varargin{v};
                        assert(islogical(show_img));
                    case 'show_legend'
                        assert(numel(varargin) > v);
                        v = v+1;
                        plot_leg = varargin{v};
                        assert(islogical(plot_leg));
                    case 'hide_nonstripe'
                        show_nonstripe = false;
                    otherwise
                        error(['unrecognized input option' varargin{v}]);
                end
                v = v+1;
            end
            
            if isempty(this.ixs_in_stripe)
                error('call this.identify_stripes() before plotting');
            end
            
            nstripes = size(this.ixs_in_stripe,2);
            
            if show_img
                imgfluo = this.get_img(channel);
                imgvals = max(imgfluo,[],3);
                imagesc(imgvals); axis off; colormap gray;
                imlims = [0,quantile(as_vector(imgvals),0.95)];
                caxis(imlims);
                hold on;
            end
            
            g = [];
            color_order = get(gca,'colororder');
            for ii_stripe = 1:nstripes
                if this.stripes_to_use(ii_stripe)
                    g = [g,scatter(this.nucleus_xy(this.ixs_in_stripe(:,ii_stripe),1), ...
                        this.nucleus_xy(this.ixs_in_stripe(:,ii_stripe),2),50, ...
                        color_order(ii_stripe,:))];
                    hold on;
                end
            end
            
            if show_nonstripe
                ixs_any_stripe = any(this.ixs_in_stripe(:,this.stripes_to_use),2);
                g = [g,scatter(this.nucleus_xy(~ixs_any_stripe & ~this.outlier_ix,1), ...
                    this.nucleus_xy(~ixs_any_stripe & ~this.outlier_ix,2),50,'m')];%0.5*[1 1 1])];
            end
            hold off;
            
            if plot_leg
                stripens = 1:nstripes;
                legend_entries = cellfun(@num2str,num2cell(stripens(this.stripes_to_use)),'UniformOutput',false);
                l = legend(legend_entries);
                set(l,'location','eastoutside','autoupdate','off','autoupdate','off')
            end
        end
        
        
        % Manually correct calibration region using a GUI.
        function manual_correct_calibration(this,channel)
            if nargin < 2
                channel = this.fluochannels(1);
            end
            
            if isempty(this.ixs_in_calib)
                warning('No calibration region found: initializing...')
                this.ixs_in_calib = false(size(this.nucleus_xy,1),1);
                xlims = [1,this.FrameInfo.PixelsPerLine];
                ylims = xlims;
            else
                padding = 500;
                xlims = [max(min(this.nucleus_xy(this.ixs_in_calib,1)-padding),1), ...
                    min(max(this.nucleus_xy(this.ixs_in_calib,1)+padding),this.FrameInfo.PixelsPerLine)];
                ylims = [max(min(this.nucleus_xy(this.ixs_in_calib,2)-padding),1), ...
                    min(max(this.nucleus_xy(this.ixs_in_calib,2)+padding),this.FrameInfo.PixelsPerLine)];
            end
            
            fig = figure;
            this.scatter_nuclei(channel,false);
            this.ixs_in_calib = this.manual_select(fig,this.ixs_in_calib,'selmarkerspec',{'y'});
            close(fig);
        end
        
        
        % Manually correct stripes using a GUI.
        function manual_correct_stripes(this,channel)
            if nargin < 2
                channel = this.fluochannels(1);
            end
            
            if isempty(this.ixs_in_stripe)
                error('call this.identify_stripes() before manual correction');
            end
            
            nstripes = size(this.ixs_in_stripe,2);
            
            fig = figure;
            gg = this.plot_stripes(channel,'show_legend',true);
            cur_stripe = 0;
            while true
                while ~(cur_stripe >= 1 && cur_stripe <= nstripes)
                    cur_stripe = input(['Enter stripe number to correct (1-' num2str(nstripes) ...
                        ') or hit enter to stop: ']);
                    
                    if isempty(cur_stripe)
                        break
                    end
                end
                
                if isempty(cur_stripe)
                    break
                else
                    disp(['Click nuclei to add them to or remove them from stripe ' ...
                        num2str(cur_stripe) '.  Press enter when finished.']);
                    
                    cur_sel_ix = this.ixs_in_stripe(:,cur_stripe);
                    cur_sel_ix = this.manual_select(fig,cur_sel_ix,'newmarkerspec',{50,'k.'});
                    rem_stripes = 1:nstripes;
                    rem_stripes(cur_stripe) = [];
                    this.ixs_in_stripe(:,cur_stripe) = cur_sel_ix;
                    this.ixs_in_stripe(cur_sel_ix,rem_stripes) = false;
                    
                    delete(gg);
                    
                    figure(fig);
                    hold on;
                    gg = this.plot_stripes(channel,'show_img',false, ...
                        'show_legend',true);
                    hold off;
                    
                    cur_stripe = 0;
                end
            end
            close(fig);
        end
        
        
        % Loads saved file associated with this object, if it exists.
        function this = load(this)
            filepath_to_load = fullfile(this.savedir,[this.prefix '.mat']);
            
            if exist(filepath_to_load,'file')
                disp(['Loading ' filepath_to_load])
                load(filepath_to_load,'this');
                disp('Loaded.')
            else
                disp('No file found for this experiment.')
            end
        end
        
        
        % Save this object in savedir.
        function save(this,opt)
            if nargin < 2
                opt = [];
            end
            
            filepath_to_save = fullfile(this.savedir,this.prefix);
            overwrite = true;
            
            if exist([filepath_to_save '.mat'],'file')
                while ~strcmpi(opt,'y') && ~strcmpi(opt,'n')
                    opt = input('Overwrite existing file? (y/n) ','s');
                end
                
                if strcmpi(opt,'n')
                    overwrite = false;
                end
            elseif ~exist(this.savedir,'dir')
                mkdir(this.savedir);
            end
            
            if overwrite
                save(filepath_to_save,'this');
                disp(['Saved to ' filepath_to_save])
            end
        end
        
        
        % Return specified field calibrated to fluorescence in calibration
        % region.
        function calib_prop = calibrated(this,prop,opt,quantval)
            if nargin < 4
                quantval = 0.95;
                if nargin < 3
                    opt = 'quantile';
                end
            end
            
            if isempty(this.ixs_in_calib)
                error('call this.identify_calibration_region()')
            end
            
            calib_prop = this.(prop);
            
            switch opt
                case 'mean'
                    calfactor = nanmean(this.fluo(this.ixs_in_calib,:),1);
                case 'quantile'
                    calfactor = quantile(this.fluo(this.ixs_in_calib,:),quantval);
                otherwise
                    error(['unsupported option ' opt]);
            end
            
            calib_prop = calib_prop./calfactor;
        end
        
        
        % Generates boxplots for fluorescence, spot fluorescence, and spot
        % fluorescence ratio by stripe.  Indices for non-stripe nuclei
        % exclude the indices used for fluorescence calibration.
        function fbp = boxplots(this,channel)
            if nargin < 2
                if ~isempty(this.fluochannels)
                    channel = this.fluochannels(1);
                else
                    channel = 1;
                end
            end
            
            ii_channel = find(this.fluochannels == channel);
            
            if isempty(ii_channel)
                disp('warning: channel not found...defaulting to index 1');
                ii_channel = 1;
            end
            
            if isempty(this.ixs_in_stripe)
                error('call this.identify_stripes() before plotting');
            end
            
            ixs_any_stripe = any(this.ixs_in_stripe,2);
            ixs_no_stripe_no_calib = ~ixs_any_stripe & ~this.outlier_ix & ~this.ixs_in_calib;
            nstripes = size(this.ixs_in_stripe,2);
            
            fluo_by_stripe = repmat(this.fluo(:,ii_channel), ...
                [1,nstripes+2]) .* [this.ixs_in_stripe,ixs_any_stripe,ixs_no_stripe_no_calib];
            fluo_by_stripe(fluo_by_stripe == 0) = NaN;
            
            spotfluo_by_stripe = repmat(this.spotfluo(:,ii_channel), ...
                [1,nstripes+2]) .* [this.ixs_in_stripe,ixs_any_stripe,ixs_no_stripe_no_calib];
            spotfluo_by_stripe(spotfluo_by_stripe == 0) = NaN;
            
            spotfluo_ratio_by_stripe = repmat(this.spotfluo_ratio(:,ii_channel), ...
                [1,nstripes+2]) .* [this.ixs_in_stripe,ixs_any_stripe,ixs_no_stripe_no_calib];
            spotfluo_ratio_by_stripe(spotfluo_ratio_by_stripe == 0) = NaN;
            
            labels = cell(nstripes+2,1);
            labels(1:nstripes) = cellfun(@num2str,num2cell(1:nstripes),'UniformOutput',false);
            labels(end-1:end) = {'all','none'};
            
            titles = {'Average fluorescence','Spot fluorescence','Spot fluorescence ratio'};
            if ~isempty(this.ixs_in_calib)
                disp('Normalizing fluorescence values to average of nuclei in calibration region...');
                calfactor = nanmean(this.fluo(this.ixs_in_calib,ii_channel));
                fluo_by_stripe = fluo_by_stripe/calfactor;
                spotfluo_by_stripe = spotfluo_by_stripe/calfactor;
                titles{1} = 'Average fluorescence (normalized)';
                titles{2} = 'Spot fluorescence (normalized)';
            end
            
            fbp = figure('name',this.prefix);
            subplot(1,3,1)
            boxplot(fluo_by_stripe,'Labels',labels);
            title(titles{1})
            xlabel('stripe')
            ylim([0,1.8])
            
            subplot(1,3,2)
            boxplot(spotfluo_by_stripe,'Labels',labels);
            title(titles{2})
            xlabel('stripe')
            ylim([0,3])
            
            subplot(1,3,3)
            boxplot(spotfluo_ratio_by_stripe,'Labels',labels);
            title(titles{3})
            xlabel('stripe')
            ylim([0,4])
            
            set(gcf,'Position',[50, 450, 1500, 400]);
        end
        
        
        % Returns the indices of the n nearest neighbors of cells with
        % indices ix (a number or logical array).  Output is a logical
        % array with one column per valid index in ixs.
        function nn = nearest_n_neighbors(this,ix,n)
            dists = (this.nucleus_xy(:,1) - this.nucleus_xy(:,1)').^2 + ...
                (this.nucleus_xy(:,2) - this.nucleus_xy(:,2)').^2;
            dists = dists + diag(inf(numel(this.outlier_ix),1));
            
            dists(this.outlier_ix,:) = inf;
            dists(:,this.outlier_ix) = inf;
            
            if islogical(ix)
                ix = find(ix);
            end
            
            nn = false(numel(this.outlier_ix),numel(ix));
            for ii = 1:numel(ix)
                [~,sort_ix] = sort(dists(ix(ii),:),'ascend');
                nn_ind = sort_ix(1:n);
                nn(nn_ind,ii) = true;
            end
        end
        
    end
    
    
    methods (Access = protected)
        function gen_tif_stack(this,LSMImage,NChannels,NSlices,ProjectionType,upperSlice,lowerSlice)
            if nargin < 7
                lowerSlice = 1;
                upperSlice = NSlices;
            end
            
            if ~exist(this.tifstackdir,'dir')
                disp(['Creating folder ' this.tifstackdir '...']);
                mkdir(this.tifstackdir);
            end
            
            disp('Converting to TIFF stacks...')
            exportTifStacksFISH(this.tifstackdir,LSMImage, ...
                NChannels,NSlices,this.prefix,ProjectionType, ...
                this.histchannel,upperSlice,lowerSlice);
        end
        
        % GUI to select nuclei.
        % Inputs:
        % - fig = image on which to draw selection
        % - cur_sel_ix = starting selection
        function selected_ix = manual_select(this,fig,cur_sel_ix,varargin)
            if nargin < 2 || isempty(cur_sel_ix)
                cur_sel_ix = false(size(this.outlier_ix));
            end
            
            assert(islogical(cur_sel_ix));
            
            rotated = false;
            selmarkerspec = {'ko'};
            newmarkerspec = {50,'r.'};
            sel_type = 'x';
            forced_sel = false;
            include_outliers = false;
            
            v = 1;
            while v <= numel(varargin)
                switch varargin{v}
                    case 'rotated'
                        rotated = true;
                    case 'selmarkerspec'
                        assert(numel(varargin) > v);
                        v = v+1;
                        selmarkerspec = varargin{v};
                        assert(iscell(selmarkerspec));
                    case 'newmarkerspec'
                        assert(numel(varargin) > v);
                        v = v+1;
                        newmarkerspec = varargin{v};
                        assert(iscell(newmarkerspec));
                    case 'box'
                        sel_type = 'b';
                        forced_sel = true;
                    case 'polygon'
                        sel_type = 'p';
                        forced_sel = true;
                    case 'include_outliers'
                        include_outliers = true;
                    otherwise
                        error(['unrecognized option ' varargin{v}]);
                end
                v = v + 1;
            end
            
            if rotated
                xycoords = this.rotated_xy;
            else
                xycoords = this.nucleus_xy;
            end
            
            
            while true
                % refresh display
                if exist('gg','var')
                    delete(gg);
                end
                figure(fig);
                hold on;
                gg = scatter(xycoords(cur_sel_ix,1),xycoords(cur_sel_ix,2),selmarkerspec{:});
                hold off;
                
                manopt = sel_type;
                while ~strcmpi(manopt,'i') && ~strcmpi(manopt,'b') && ...
                        ~strcmpi(manopt,'p') && ~isempty(manopt)
                    manopt = input('Select nuclei individually (i), by box (b), by polygon (p), or press enter to finish: ','s');
                end
                
                if isempty(manopt)
                    break;
                end
                
                switch manopt
                    case 'i'
                        disp('Click to (de)select.  Press enter to exit selection mode.')
                    case 'b'
                        disp('Click opposite corners to box selection.  Press enter to exit selection mode.')
                    case 'p'
                        disp('Click points to outline a polygon around selection.  Press enter when finished.')
                    otherwise
                        disp('Unrecognized option.')
                        manopt = [];
                end
                
                while true
                    switch manopt
                        case 'i' % individual
                            xy = ginput(1);
                            
                            if isempty(xy)
                                delete(gg);
                                break;
                            end
                            [~,nearest_ix] = min(sum((xycoords - xy).^2,2));
                            cur_sel_ix(nearest_ix) = ~cur_sel_ix(nearest_ix);
                            
                            if ~include_outliers
                                cur_sel_ix(this.outlier_ix) = false;
                            end
                            
                            % refresh display
                            delete(gg);
                            hold on;
                            gg = scatter(xycoords(cur_sel_ix,1),xycoords(cur_sel_ix,2),selmarkerspec{:});
                            hold off;                            
                        case 'b' % box
                            new_sel_ix = false(size(cur_sel_ix));
                            
                            xy = ginput(2);
                            
                            if size(xy,1) ~= 2
                                break;
                            end
                            
                            % (de)select points inside box
                            bounds = [min(xy(:,1)),min(xy(:,2)); ...
                                max(xy(:,1)),max(xy(:,2))];
                            
                            ix = (xycoords(:,1) > bounds(1,1)) & ...
                                (xycoords(:,1) < bounds(2,1)) & ...
                                (xycoords(:,2) > bounds(1,2)) & ...
                                (xycoords(:,2) < bounds(2,2));
                            new_sel_ix(ix) = true;
                            
                            if ~include_outliers
                                new_sel_ix(this.outlier_ix) = false;
                            end
                        case 'p' % polygon
                            curpts = [];
                            while true
                                xy = ginput(1);
                                
                                if size(xy,2) ~= 2
                                    break;
                                end
                                
                                curpts = [curpts;xy];
                                
                                hold on;
                                if size(curpts,1) == 1
                                    pb = scatter(curpts(1),curpts(2),75,'r.');
                                else
                                    delete(pb);
                                    pb = plot(curpts(:,1),curpts(:,2),'r','linewidth',2);
                                end
                                hold off;
                            end
                            
                            if exist('pb','var')
                                delete(pb);
                            end
                            
                            if isempty(curpts)
                                break;
                            else
                                % (de)select points inside polygon
                                new_sel_ix = inpolygon(xycoords(:,1),xycoords(:,2), ...
                                    curpts(:,1),curpts(:,2));
                                
                                if ~include_outliers
                                    new_sel_ix(this.outlier_ix) = false;
                                end
                            end
                    end
                    
                    if ~strcmpi(manopt,'i')
                        % show new selection
                        hold on;
                        ns = scatter(xycoords(new_sel_ix,1),xycoords(new_sel_ix,2),newmarkerspec{:});
                        hold off;
                        
                        opt = 'x';
                        if ~forced_sel
                            while ~isempty(opt) && ~strcmpi(opt,'d') && ~strcmpi(opt,'c')
                                opt = input('Press enter to select, d to deselect, or c to clear. ','s');
                            end
                            
                            switch opt
                                case []
                                    cur_sel_ix = cur_sel_ix | new_sel_ix;
                                    delete(ns);
                                    break;
                                case 'd'
                                    cur_sel_ix = cur_sel_ix & ~new_sel_ix;
                                    delete(ns);
                                    break;
                                case 'c'
                                    delete(ns);
                            end
                        else
                            while ~isempty(opt) && ~strcmpi(opt,'c')
                                opt = input('Press enter to select or c to clear. ','s');
                            end
                            
                            switch opt
                                case []
                                    selected_ix = new_sel_ix;
                                    delete(ns);
                                    return;
                                case 'c'
                                    delete(ns);
                            end
                        end
                    end
                    
                end
            end
            delete(gg);
            selected_ix = cur_sel_ix;
        end
        
        
        % Helper function for this.identify_stripes().
        function [stripe_cenx,stripe_ceny,stripe_width_left,stripe_width_right,bins,nbins] = ...
                find_stripes(this,channel,expectednstripes,maxnstripes, ...
                horizontal,exclude_ix,loess_smooth_factor,debug_plot,debug_plot_peaks, ...
                stripecutofffactor,slope_cutoff)
            
            if isempty(exclude_ix)
                exclude_ix = false(size(this.outlier_ix));
            end
            
            exclude_ix = exclude_ix | this.outlier_ix;
            
            if horizontal
                % bin along x axis
                [N,~,bins_temp] = histcounts(this.rotated_xy(~exclude_ix,1));
            else % stripes are vertical
                % bin along y axis
                [N,~,bins_temp] = histcounts(this.rotated_xy(~exclude_ix,2));
            end
            nbins = max(bins_temp);
            
            bins = nan(size(this.rotated_xy,1),1);    % resize bins for outliers
            bins(~exclude_ix) = bins_temp;

            stripe_cenx = nan(nbins,maxnstripes);
            stripe_ceny = nan(nbins,maxnstripes);
            stripe_width_left = nan(nbins,maxnstripes);
            stripe_width_right = nan(nbins,maxnstripes);
            
            % identify fluorescence peaks independently in each bin
            for ii = 1:nbins
                cur_bin = ii;
                
                if N(ii) > 4
                    cur_ix = (bins == cur_bin) & ~exclude_ix;
                    
                    cur_fluo = this.fluo(cur_ix,channel);
                    
                    if horizontal
                        % swap coordinates
                        cur_cenx = this.rotated_xy(cur_ix,2);
                        cur_ceny = this.rotated_xy(cur_ix,1);
                    else
                        cur_cenx = this.rotated_xy(cur_ix,1);
                        cur_ceny = this.rotated_xy(cur_ix,2);
                    end
                    
                    [cur_cenx,sort_ix] = sort(cur_cenx);
                    cur_ceny = cur_ceny(sort_ix);
                    cur_fluo = cur_fluo(sort_ix);
                    
                    % perform smoothing on appropriate profile at current
                    % coordinate
                    cur_fluo = smooth(cur_cenx,cur_fluo,loess_smooth_factor,'loess');
                    
                    if any(~isnan(cur_fluo))
                        peak_ix = peakfinder(cur_fluo,range(cur_fluo)/8,[],[],false);
                        peak_ix = unique(peak_ix);
                        trough_ix = peakfinder(cur_fluo,range(cur_fluo)/8,[],-1,true);
                        trough_ix = unique(trough_ix);
                        
                        % width = distance to half max relative to side of smaller height,
                        % i.e., peak height is defined relative to the higher of the two
                        % neighboring troughs
                        left_trough_ix = (trough_ix(1:end-1));
                        right_trough_ix = (trough_ix(2:end));
                        
                        left_height = cur_fluo(peak_ix) - cur_fluo(left_trough_ix);
                        right_height = cur_fluo(peak_ix) - cur_fluo(right_trough_ix);
                        
                        min_height = min(left_height,right_height);
                        left_thresh = cur_fluo(peak_ix) - (min_height/2);
                        right_thresh = cur_fluo(peak_ix) - (min_height/2);
                        
                        right_width = nan(1,numel(peak_ix));
                        left_width = nan(1,numel(peak_ix));
                        % for computational efficiency, no interpolation
                        for jj = 1:numel(peak_ix)
                            dist_to_below_thresh_left = cur_cenx(peak_ix(jj)) - cur_cenx(cur_fluo < left_thresh(jj));
                            % only consider threshhold crossings to left of current peak
                            dist_to_below_thresh_left = dist_to_below_thresh_left(dist_to_below_thresh_left > 0);
                            left_width(jj) = min(dist_to_below_thresh_left);
                            
                            dist_to_below_thresh_right = cur_cenx(peak_ix(jj)) - cur_cenx(cur_fluo < right_thresh(jj));
                            % only consider threshold crossings to right of current peak
                            dist_to_below_thresh_right = dist_to_below_thresh_right(dist_to_below_thresh_right < 0);
                            right_width(jj) = min(abs(dist_to_below_thresh_right));
                        end
                        
                        if debug_plot_peaks
                            subplot(5,5,ii)
                            plot(cur_cenx,cur_fluo);
                            hold on
                            scatter(cur_cenx(peak_ix),cur_fluo(peak_ix));
                            scatter(cur_cenx(trough_ix),cur_fluo(trough_ix));
                            scatter(cur_cenx(peak_ix) - left_width', ...
                                cur_fluo(peak_ix) - min_height/2,'r');
                            scatter(cur_cenx(peak_ix) + right_width', ...
                                cur_fluo(peak_ix) - min_height/2,'r');
                            hold off
                            title(num2str(cur_bin))
                        end
                        
                        n_detected_stripes = numel(peak_ix);
                        
                        if n_detected_stripes > maxnstripes
                            warning([num2str(n_detected_stripes) ' stripes detected; ' ...
                                'truncating results to first ' num2str(maxnstripes)]);
                            n_detected_stripes = maxnstripes;
                            peak_ix = peak_ix(1:n_detected_stripes);
                            left_width = left_width(1:n_detected_stripes);
                            right_width = right_width(1:n_detected_stripes);
                        end
                        
                        stripe_cenx(ii,1:n_detected_stripes) = cur_cenx(peak_ix);
                        stripe_ceny(ii,1:n_detected_stripes) = cur_ceny(peak_ix);
                        
                        stripe_width_left(ii,1:n_detected_stripes) = left_width;
                        stripe_width_right(ii,1:n_detected_stripes) = right_width;
                    end
                else
                    warning(['Only ' num2str(N(ii)) ' nuclei in bin ' num2str(ii) ...
                        '...ignoring bin...']);
                end
            end
            
            % realign stripes
            [stripe_cenx,stripe_ceny,stripe_width_left,stripe_width_right] = ...
                realign_stripes_pool(expectednstripes, ...
                stripe_cenx,stripe_ceny,stripe_width_left,stripe_width_right);
            
            % prune
            [stripe_cenx,stripe_ceny,stripe_width_left,stripe_width_right] = ...
                realign_stripes_pool(expectednstripes, ...
                stripe_cenx,stripe_ceny,stripe_width_left,stripe_width_right,true);
            
            % prune stripe width outliers
            for ii = 1:maxnstripes
                stripe_width_cutoff = nanmean([stripe_width_left(:,ii);stripe_width_right(:,ii)]);
                stripe_width_left(stripe_width_left(:,ii) > ...
                    stripecutofffactor*stripe_width_cutoff,ii) = stripe_width_cutoff;
                stripe_width_right(stripe_width_right(:,ii) > ...
                    stripecutofffactor*stripe_width_cutoff,ii) = stripe_width_cutoff;
            end
            
            % fix crossover points between neighboring stripes
            [stripe_cenx,stripe_ceny,stripe_width_left,stripe_width_right] = ...
                remove_stripe_xings(stripe_cenx,stripe_ceny,stripe_width_left,stripe_width_right);
            
            % remove "outlier" points (high slope)
            [stripe_cenx,stripe_ceny,stripe_width_left,stripe_width_right] = ...
                smooth_stripes(stripe_cenx,stripe_ceny,stripe_width_left,stripe_width_right,slope_cutoff);
            
            if debug_plot
                figure
                show_stripes_xy(stripe_cenx,stripe_ceny,stripe_width_left,stripe_width_right);
            end

            stripe_cenx = stripe_cenx(:,1:expectednstripes);
            stripe_ceny = stripe_ceny(:,1:expectednstripes);
            stripe_width_left = stripe_width_left(:,1:expectednstripes);
            stripe_width_right = stripe_width_right(:,1:expectednstripes);
        end
        
        
        % Helper function for this.identify_stripes().  Returns ncells x
        % nstripes logical array indicating which nuclei belong to which
        % stripe.
        function ixs = assign_stripes(this,stripe_cenx,stripe_ceny, ...
                stripe_width_left,stripe_width_right,stripecutofffactor, ...
                interp_stripes,horizontal,bins,nbins)
            nstripes = size(stripe_cenx,2);
            
            % assign nuclei to stripes
            if interp_stripes
                yinterp = nanmean(stripe_ceny,2);
                for kk = 1:nstripes
                    valix_range = find(~isnan(stripe_cenx(:,kk)));
                    if numel(valix_range) > 2
                        valix = valix_range(1):valix_range(end);
                        [stripe_cenx(valix,kk),~, ...
                            stripe_width_left(valix,kk),stripe_width_right(valix,kk)] = ...
                            interpolate_stripes(stripe_cenx(:,kk),stripe_ceny(:,kk), ...
                            stripe_width_left(:,kk),stripe_width_right(:,kk), ...
                            yinterp(valix));
                    end
                end
                
                % prune stripe width outliers
                for ii = 1:nstripes
                    stripe_width_cutoff = nanmean([stripe_width_left(:,ii);stripe_width_right(:,ii)]);
                    stripe_width_left(stripe_width_left(:,ii) > ...
                        stripecutofffactor*stripe_width_cutoff,ii) = stripe_width_cutoff;
                    stripe_width_right(stripe_width_right(:,ii) > ...
                        stripecutofffactor*stripe_width_cutoff,ii) = stripe_width_cutoff;
                end
            end
            
            ixs = false(size(this.rotated_xy,1),nstripes);
            left_bounds = stripe_cenx(:,1:nstripes) - stripe_width_left(:,1:nstripes);
            right_bounds = stripe_cenx(:,1:nstripes) + stripe_width_right(:,1:nstripes);
            
            for ii_bin = 1:nbins
                cur_nuc_ix = (bins == ii_bin);
                for ii_stripe = 1:nstripes
                    xbounds = [left_bounds(ii_bin,ii_stripe),right_bounds(ii_bin,ii_stripe)];
                    if horizontal
                        ixs(:,ii_stripe) = ~any(ixs(:,1:ii_stripe-1),2) & ...
                            ((ixs(:,ii_stripe)) | ...
                            (cur_nuc_ix(:) & (this.rotated_xy(:,2) > xbounds(1)) & ...
                            (this.rotated_xy(:,2) < xbounds(2))));
                    else
                        ixs(:,ii_stripe) = ~any(ixs(:,1:ii_stripe-1),2) & ...
                            ((ixs(:,ii_stripe)) | ...
                            (cur_nuc_ix(:) & (this.rotated_xy(:,1) > xbounds(1)) & ...
                            (this.rotated_xy(:,1) < xbounds(2))));
                    end
                end
            end
            
            % make sure nuclei are assigned to only one stripe
            disputed_nuclei = (sum(ixs,2) > 1);
            if any(disputed_nuclei)
                error('disputed nuclei')
            end
        end
        
    end
    
    
    methods (Access = private)
        % Fix for backwards compatibility with files saved using a previous
        % version of this code.
        function fix_savedir(this)
            if ~contains(this.savedir,this.date)
                this.savedir = fullfile(this.savedir,this.date);
            end
        end
        
        
        % If ix is in a stripe, returns the number of the stripe; otherwise
        % returns empty.
        function n = is_in_stripe(this,ix)
            n = find(this.ixs_in_stripe(ix,:));
        end
    end
    
end
```
