## Supplementary material for "Enhancer architecture and chromatin accessibility constrain phenotypic space during development": Script-MLP: fish_seg_pool.m

```
% Object for analyzing several FISH embryos.
%
% Creator: Mindy Liu Perkins
% Please note this code is not packaged for general use, so there are some
% unused/"test" functions, incomplete edge case detection, and some implied
% functionalities that are not fully implemented.

classdef fish_seg_pool < fish_seg
    
    properties (SetAccess = private)
        daughter_fish;	% array of fish_seg objects belonging to this pool
        
        stripe_widths_normalized;   % normalized to approx. length of embryo
    end
    
    methods
        
        function this = fish_seg_pool(varargin)
            this.daughter_fish = fish_seg.empty(0,0);
            if ~isempty(varargin)
                this = this.initialize(varargin{:});
            end
        end
        
        function this = initialize(this,project,name,date,rootdir)
            this.project = project;
            this.prefix = name;           
            if nargin > 3
                this.date = date;
                if nargin > 4
                    this.rootdir = rootdir;
                end
            end
            
            this.datadir = fullfile(this.rootdir,this.project);
            this.savedir = fullfile(this.rootdir,'processed_data',this.project);
            this.filepath = fullfile(this.datadir,this.date,[this.prefix '.czi']);
            this.tifstackdir = fullfile(this.datadir,this.date,'tifstacks');
            
            if ~exist(this.filepath,'file')
                warning([this.filepath ' does not exist'])
            end
            
            this = this.load();
        end
        
        function init_by_filename_containing(this,dates,filenames,searchterm)
            if ~iscell(dates)
                dates = {dates};
            end
            
            nf = numel(filenames);
            
            if numel(dates) == 1
                dates = repmat(dates,[nf,1]);
            elseif numel(dates) ~= nf
                error('dates must be a singular date for all filenames or a date per filename');
            end
            
            for ii = 1:nf
                filenames{ii} = erase(filenames{ii},'.czi');
                if contains(filenames{ii},searchterm)
                    this.add_daughter(this.project,dates{ii},filenames{ii});
                end
            end
        end
        
        function batch_extract(this,varargin)
            disp('Extracting raw data...')
            
            ProjectionType = this.histprojection_type;
            cur_filepath = this.filepath;
            slice_range = [];
            
            v = 1;
            while v < numel(varargin)
                switch varargin{v}
                    case 'filepath'
                        assert(numel(varargin) > v);
                        v = v+1;
                        cur_filepath = varargin{v};
                    case 'ProjectionType'
                        assert(numel(varargin) > v);
                        v = v+1;
                        ProjectionType = varargin{v};
                        this.histprojection_type = ProjectionType;
                    case 'slice_range'
                        assert(numel(varargin) > v);
                        v = v+1;
                        slice_range = varargin{v};
                        assert(isnumeric(slice_range) && numel(slice_range) == 2);
                    otherwise
                        error(['unrecognized input option ' varargin{v}]);
                end
                v = v+1;
            end
            
            [this.FrameInfo,LSMImages,NSlices,~,NFrames,~,NChannels] ...
                = getZeissFrameInfoFISH(cur_filepath,[]);
            
            assert(NFrames == 1);
            
            NSeries = size(LSMImages,1);
            for ii_series = 1:NSeries
                this.add_daughter(this.project,this.date,[this.prefix,'-',num2str(ii_series)]);
                LSMImage = LSMImages(ii_series);
                
                this.daughter_fish(end).histprojection_type = this.histprojection_type;
                this.daughter_fish(end).histchannel = this.histchannel;
                if ~isempty(slice_range)
                    this.daughter_fish(end).gen_tif_stack(LSMImage,NChannels,NSlices,ProjectionType,slice_range(2),slice_range(1));
                else
                    this.daughter_fish(end).gen_tif_stack(LSMImage,NChannels,NSlices,ProjectionType);
                end
                this.daughter_fish(end).FrameInfo = this.FrameInfo;
            end
            disp('Done.')
        end
        
        function add_daughter(this,project,date,prefix)
            nel = numel(this.daughter_fish);
            
            if ~isempty(this.rootdir)
                this.daughter_fish(nel+1) = fish_seg(project,date,prefix,this.rootdir);
            else
                this.daughter_fish(nel+1) = fish_seg(project,date,prefix);
            end
        end
        
        % Currently does not support multiple fluorescence channels.
        function populate(this)
            valid_ix = vertcat(this.daughter_fish.skip);
            
            if numel(valid_ix) ~= numel(this.daughter_fish)
                for ii = 1:numel(this.daughter_fish)
                    if isempty(this.daughter_fish(ii).skip)
                        this.daughter_fish(ii).mark_skip();
                    end
                end
                valid_ix = vertcat(this.daughter_fish.skip);
            end
            
            this.ixs_in_stripe = vertcat(this.daughter_fish(valid_ix).ixs_in_stripe);
            this.outlier_ix = vertcat(this.daughter_fish(valid_ix).outlier_ix);
            this.ixs_in_calib = vertcat(this.daughter_fish(valid_ix).ixs_in_calib);
            
            this.spotfluo_ratio = vertcat(this.daughter_fish(valid_ix).spotfluo_ratio);
            
            this.nucdist = vertcat(this.daughter_fish(valid_ix).nucdist);
            
            if ~isempty(this.ixs_in_calib)
                this.fluo = [];
                this.spotfluo = [];
                this.stripe_widths_normalized = [];
                for ii = 1:numel(this.daughter_fish)
                    if valid_ix(ii)
                        this.fluo = [this.fluo; ...
                            this.daughter_fish(ii).calibrated('fluo')];
                        this.spotfluo = [this.spotfluo; ...
                            this.daughter_fish(ii).calibrated('spotfluo')];
                    end
                end
            else
                this.fluo = vertcat(this.daughter_fish(valid_ix).fluo);
                this.spotfluo = vertcat(this.daughter_fish(valid_ix).spotfluo);
            end
            
            for ii = 1:numel(this.daughter_fish)
                if valid_ix(ii)
                    this.stripe_widths_normalized = [this.stripe_widths_normalized; ...
                        this.daughter_fish(ii).get_stripe_width('normalized')];
                end
            end
        end
        
        
        function all_daughters(this,fnname,varargin)
            for ii = 1:numel(this.daughter_fish)
                this.daughter_fish(ii).(fnname)(varargin{:});
            end
        end

    end

end
```
