## Supplementary material for "Enhancer architecture and chromatin accessibility constrain phenotypic space during development": Script-MLP: exportTifStacksFISH.m

```
function exportTifStacksFISH(savedir, AllImages, NChannels, NSlices,...
    Prefix, ProjectionType, histchannel_ix,upperSlice,lowerSlice)

PreProcFolder = savedir;

ySize = size(AllImages{1}{1,1}, 1);
xSize = size(AllImages{1}{1,1}, 2);
BlankImage = zeros(ySize, xSize, 'uint16');

topZSlice = min(NSlices);

for channelIndex = 1:NChannels
    
    NameSuffix = ['_ch',iIndex(channelIndex,2)];
    
    NewName = [Prefix,NameSuffix, '.tif'];
    
    % write bottom slice (always a blank padding image)
    imwrite(BlankImage, [PreProcFolder, filesep, NewName]);
    
    % copy the rest of the images
    slicesCounter = 1;
    firstImageIndex = 1 + (channelIndex - 1);
    lastImageIndex = NSlices(1) * NChannels;
    if firstImageIndex == lastImageIndex
        firstImageIndex = 1;
        lastImageIndex = 1;
    end
    for imageIndex = firstImageIndex:NChannels:lastImageIndex
        if slicesCounter <= topZSlice
            % if zPadding, it will process all images (because topZSlice would be max(NSlices)
            % if no zPadding, it will process images rounding down to the series with least
            % zSlices, because topZSlice would be min(NSlices)
            
            imwrite(AllImages{1,1}{imageIndex,1},...
                [PreProcFolder, filesep, NewName], 'WriteMode', 'append');
            slicesCounter = slicesCounter + 1;
        end
    end
    
    % save as many blank images at the end of the stack are needed
    %(depending on zPadding being active or not)
    for zPaddingIndex = slicesCounter+1:topZSlice+2
        imwrite(BlankImage, [PreProcFolder, filesep, NewName], 'WriteMode', 'append');
    end
    
    if channelIndex == histchannel_ix
        filepath = [PreProcFolder,filesep,NewName];
        
        % generate projection
        disp(['Generating projection for channel ' num2str(histchannel_ix) '...']);
        hisstack = read_tiff_stack(filepath);
        
        hisMat = calculateProjection(ProjectionType,size(hisstack,3),hisstack, ...
            upperSlice,lowerSlice);
        
        saveNuclearProjection(hisMat, [PreProcFolder, filesep, Prefix, '-His.tif']);
    end
    
end

end
```
