## Supplementary material for "Enhancer architecture and chromatin accessibility constrain phenotypic space during development": Script-MLP: read_tiff_stack.m

```
function img = read_tiff_stack(filepath)

img_info = imfinfo(filepath);
nstacks = numel(img_info);
img = nan(img_info(1).Height,img_info(1).Width,nstacks);
for ii_stack = 1:nstacks
    img(:,:,ii_stack) = imread(filepath,ii_stack);
end

end
```
