## Supplementary material for "Enhancer architecture and chromatin accessibility constrain phenotypic space during development": Script-MLP: interpolate_stripes.m

```
function [xq,yq,xlq,xrq] = interpolate_stripes(stripe_cenx,stripe_ceny, ...
    stripe_width_left,stripe_width_right,interpts)

if nargin < 5 || isempty(interpts)
    interpts = 100;
end

% interpts is either a scalar (for a number of linearly spaced points) or a
% vector (for the interpolation points themselves)
if numel(interpts) == 1
    ninterpts = interpts;
else
    ninterpts = numel(interpts);
end

nstripes = size(stripe_cenx,2);

xq = nan(ninterpts,nstripes);
yq = xq;
xlq = xq;
xrq = xq;

% interpolate stripes
for kk = 1:nstripes
    valix = ~isnan(stripe_cenx(:,kk));
    if sum(valix) > 1
        if numel(interpts) ~= 1
            yq(:,kk) = interpts;
        else
            yq(:,kk) = linspace(nanmin(stripe_ceny(:,kk)),nanmax(stripe_ceny(:,kk)),ninterpts)';
        end
        xq(:,kk) = interp1(stripe_ceny(valix,kk),stripe_cenx(valix,kk),yq(:,kk),'linear','extrap');
        xlq(:,kk) = interp1(stripe_ceny(valix,kk),stripe_width_left(valix,kk),yq(:,kk),'linear','extrap');
        xrq(:,kk) = interp1(stripe_ceny(valix,kk),stripe_width_right(valix,kk),yq(:,kk),'linear','extrap');
    end
end

end
```
