## Supplementary material for "Enhancer architecture and chromatin accessibility constrain phenotypic space during development": Script-MLP: realign_stripes.m

```
function [stripe_cenx,stripe_ceny,stripe_width_left,stripe_width_right] = ...
    realign_stripes(expectednstripes, ...
    stripe_cenx,stripe_ceny,stripe_width_left,stripe_width_right,reldir,pruneflag)

if nargin < 7
    pruneflag = false;
    if nargin < 6
        reldir = -1;
    end
end

t_cutoff = 5;  % max iteration time

if isempty(expectednstripes)
    warning('no expected number of stripes provided; defaulting to 7');
    expectednstripes = 7;
end

prev_stripe_cenx = nan(size(stripe_cenx));

count = 0;
start_t = tic;
while (numel(stripe_cenx(~isnan(stripe_cenx))) ~= numel(prev_stripe_cenx(~isnan(prev_stripe_cenx)))) ...
        || (any(stripe_cenx(~isnan(stripe_cenx)) ~= prev_stripe_cenx(~isnan(stripe_cenx))))
    prev_stripe_cenx = stripe_cenx;
    [stripe_cenx,stripe_ceny,stripe_width_left,stripe_width_right] = ...
        realign_stripes_iterate(expectednstripes,reldir, ...
        stripe_cenx,stripe_ceny,stripe_width_left,stripe_width_right,pruneflag);

    if count == 0
        reldir = -reldir;
    end
    count = count + 1;
    
    if toc(start_t) > t_cutoff
        disp('reached maximum time for iteration--stopping...')
        break;
    end
end


end
```
