## Supplementary material for "Enhancer architecture and chromatin accessibility constrain phenotypic space during development": Script-MLP: realign_stripes_iterate.m

```
function [stripe_cenx,stripe_ceny,stripe_width_left,stripe_width_right] = ...
    realign_stripes_iterate(expectednstripes,reldir,...
    stripe_cenx,stripe_ceny,stripe_width_left,stripe_width_right,pruneflag)

if nargin < 7
    pruneflag = false;
end

nbins = size(stripe_cenx,1);

if reldir == -1
    condif = 1;
elseif reldir == 1
    condif = nbins;
else
    error('unrecognized option for relative direction')
end

nstripes_per_bin = sum(~isnan(stripe_cenx),2);
nstripes = min(expectednstripes,max(nstripes_per_bin));

rowixs = (1:nbins)';

for ii = 1:nbins
    cur_peak_cenx = stripe_cenx(ii,:);
    valid_ix = ~isnan(cur_peak_cenx);
    
    avg_stripe_cenx = nanmean(stripe_cenx(nstripes_per_bin == nstripes & rowixs ~= ii,:),1);
    
    if (ii == condif) || (nstripes_per_bin(ii+reldir) < nstripes_per_bin(ii))
        % align to mean of stripe
        xdist = abs(avg_stripe_cenx' - cur_peak_cenx);
    else    % align to weighted combo of neighboring non-NaN entry and stripe mean
        temp_stripe_cenx = nan([1,size(stripe_cenx,2)]);
        for jj = 1:size(stripe_cenx,2)
            if pruneflag
                nextix_prevstripe = nan(nbins,1);
                nextix_nextstripe = nan(nbins,1);
                
                if jj > 1
                    nextix_prevstripe = find(stripe_cenx(:,jj-1));
                    nextix_prevstripe(isnan(stripe_cenx(:,jj-1))) = [];
                end
                
                if jj < size(stripe_cenx,2)
                    nextix_nextstripe = find(stripe_cenx(:,jj+1));
                    nextix_nextstripe(isnan(stripe_cenx(:,jj+1))) = [];
                end
            end
            
            nextix = find(stripe_cenx(:,jj));
            nextix(isnan(stripe_cenx(:,jj))) = [];
            
            if reldir == 1
                nextix = min(nextix(nextix > ii));
                if pruneflag
                    nextix_prevstripe = min(nextix_prevstripe(nextix_prevstripe > ii));
                    nextix_nextstripe = min(nextix_nextstripe(nextix_nextstripe > ii));
                    nextix = max([nextix_prevstripe,nextix,nextix_nextstripe]);
                end
            else
                nextix = max(nextix(nextix < ii));
                if pruneflag
                    nextix_prevstripe = max(nextix_prevstripe(nextix_prevstripe < ii));
                    nextix_nextstripe = max(nextix_nextstripe(nextix_nextstripe < ii));
                    nextix = min([nextix_prevstripe,nextix,nextix_nextstripe]);
                end
            end
            
            if ~isempty(nextix)
                temp_stripe_cenx(jj) = stripe_cenx(nextix,jj);
            end
        end
        
        xdist = 2*abs(temp_stripe_cenx' - cur_peak_cenx) + ...
            abs(avg_stripe_cenx' - cur_peak_cenx);
    end
    
    [~,min_dist_ix] = min(xdist,[],1);
    min_dist_ix(isnan(cur_peak_cenx)) = NaN;    % remove distance to self
    
    % if repeats found in min_dist_ix, align to smaller of the two
    unique_min_dist_ix = unique(min_dist_ix);
    if numel(unique_min_dist_ix) < numel(~isnan(min_dist_ix))
        neach = sum(unique_min_dist_ix' - min_dist_ix == 0,2);
        repeat_ixs = unique_min_dist_ix(neach > 1);
        for ii_rep = repeat_ixs
            temp = min_dist_ix;
            temp(min_dist_ix ~= ii_rep) = inf;
            [~,ix2keep] = min(temp);
            temp(ix2keep) = inf;
            ix2trash = ~isinf(temp);
            
            min_dist_ix(ix2trash) = NaN;
            valid_ix(ix2trash) = false;
        end
    end
    
    tempix = (min_dist_ix(~isnan(min_dist_ix)));
    align_ixs = false(size(stripe_cenx(ii,:)));
    align_ixs(tempix) = true;
    
    stripe_cenx(ii,align_ixs) = stripe_cenx(ii,valid_ix);
    stripe_cenx(ii,~align_ixs) = NaN;
    
    stripe_ceny(ii,align_ixs) = stripe_ceny(ii,valid_ix);
    stripe_cenx(ii,~align_ixs) = NaN;
    
    stripe_width_left(ii,align_ixs) = stripe_width_left(ii,valid_ix);
    stripe_width_left(ii,~align_ixs) = NaN;
    
    stripe_width_right(ii,align_ixs) = stripe_width_right(ii,valid_ix);
    stripe_width_right(ii,~align_ixs) = NaN;
end

end
```
