## Supplementary material for "Enhancer architecture and chromatin accessibility constrain phenotypic space during development": Script-MLP: realign_stripes_pool.m

```
function [stripe_cenx,stripe_ceny,stripe_width_left,stripe_width_right] = ...
    realign_stripes_pool(expectednstripes, ...
    stripe_cenx,stripe_ceny,stripe_width_left,stripe_width_right,pruneflag)

if nargin < 6
    pruneflag = false;
end

if isempty(expectednstripes)
    warning('no expected number of stripes provided; defaulting to 7');
    expectednstripes = 7;
end

stripe_cenx_pool = nan([size(stripe_cenx),2]);
stripe_ceny_pool = nan([size(stripe_cenx),2]);
stripe_width_left_pool = nan([size(stripe_cenx),2]);
stripe_width_right_pool = nan([size(stripe_cenx),2]);

% sweep one direction
[stripe_cenx_pool(:,:,1),stripe_ceny_pool(:,:,1), ...
    stripe_width_left_pool(:,:,1),stripe_width_right_pool(:,:,1)] = ...
    realign_stripes(expectednstripes, ...
    stripe_cenx,stripe_ceny,stripe_width_left,stripe_width_right,-1,pruneflag);

% sweep other direction
[stripe_cenx_pool(:,:,2),stripe_ceny_pool(:,:,2), ...
    stripe_width_left_pool(:,:,2),stripe_width_right_pool(:,:,2)] = ...
    realign_stripes(expectednstripes, ...
    stripe_cenx,stripe_ceny,stripe_width_left,stripe_width_right,1,pruneflag);

% pick stripe alignments with lowest variance for stripe center
stripe_cenx_var = [var(stripe_cenx_pool(:,:,1),0,1,'omitnan'); ...
    var(stripe_cenx_pool(:,:,2),0,1,'omitnan')];

[~,minixs] = min(stripe_cenx_var,[],1);

for ii = 1:size(stripe_cenx,2)
    stripe_cenx(:,ii) = stripe_cenx_pool(:,ii,minixs(ii));
    stripe_ceny(:,ii) = stripe_ceny_pool(:,ii,minixs(ii));
    stripe_width_left(:,ii) = stripe_width_left_pool(:,ii,minixs(ii));
    stripe_width_right(:,ii) = stripe_width_right_pool(:,ii,minixs(ii));
end

end
```
