## Supplementary material for "Enhancer architecture and chromatin accessibility constrain phenotypic space during development": Script-MLP: remove_stripe_xings.m

```
function [stripe_cenx,stripe_ceny,stripe_width_left,stripe_width_right] = ...
    remove_stripe_xings(stripe_cenx,stripe_ceny,stripe_width_left,stripe_width_right)

maxiter = 9;

stripe_cenx_pool = nan([size(stripe_cenx),2]);
stripe_ceny_pool = nan([size(stripe_cenx),2]);
stripe_width_left_pool = nan([size(stripe_cenx),2]);
stripe_width_right_pool = nan([size(stripe_cenx),2]);

stripe_cenx_pool(:,:,1) = stripe_cenx;
stripe_ceny_pool(:,:,1) = stripe_ceny;
stripe_width_left_pool(:,:,1) = stripe_width_left;
stripe_width_right_pool(:,:,1) = stripe_width_right;

stripe_cenx_pool(:,:,2) = stripe_cenx;
stripe_ceny_pool(:,:,2) = stripe_ceny;
stripe_width_left_pool(:,:,2) = stripe_width_left;
stripe_width_right_pool(:,:,2) = stripe_width_right;

prev_stripe_cenx = nan(size(stripe_cenx_pool(:,:,2)));
for ii = 1:maxiter
    temp_comp = stripe_cenx_pool(:,:,2);
    temp_comp(isnan(temp_comp)) = 0;
    prev_stripe_cenx(isnan(prev_stripe_cenx)) = 0;
    if isequal(temp_comp,prev_stripe_cenx)
        break;
    end
    
    prev_stripe_cenx = stripe_cenx_pool(:,:,2);
    
    [stripe_cenx_pool(:,:,2),stripe_ceny_pool(:,:,2), ...
        stripe_width_left_pool(:,:,2),stripe_width_right_pool(:,:,2)] = ...
        remove_stripe_xings_iterate( ...
        stripe_cenx_pool(:,:,2),stripe_ceny_pool(:,:,2),stripe_width_left_pool(:,:,2),stripe_width_right_pool(:,:,2));
    
    % sort A->P
    [~,sort_ix] = sort(nanmean(stripe_cenx_pool(:,:,2),1),'ascend','MissingPlacement','last');
    stripe_cenx_pool(:,:,2) = stripe_cenx_pool(:,sort_ix,2);
    stripe_ceny_pool(:,:,2) = stripe_ceny_pool(:,sort_ix,2);
    stripe_width_left_pool(:,:,2) = stripe_width_left_pool(:,sort_ix,2);
    stripe_width_right_pool(:,:,2) = stripe_width_right_pool(:,sort_ix,2);
end

ix = 1;
if ii > 2
    fig = figure;
    labels = {'Original','Corrected'};
    for ii_plot = 1:2
        subplot(1,2,ii_plot)
        show_stripes_xy(stripe_cenx_pool(:,:,ii_plot),stripe_ceny_pool(:,:,ii_plot), ...
            stripe_width_left_pool(:,:,ii_plot),stripe_width_right_pool(:,:,ii_plot));
        title(labels{ii_plot});
    end
    
    % allow user to choose
    opt = [];
    
    while ~strcmpi(opt,'y') && ~strcmpi(opt,'n')
        opt = input('Keep correction? (y/n) ','s');
    end
    
    if strcmpi(opt,'y')
        ix = 2;
    end
    
    close(fig);
end

for ii = 1:size(stripe_cenx,2)
    stripe_cenx(:,ii) = stripe_cenx_pool(:,ii,ix);
    stripe_ceny(:,ii) = stripe_ceny_pool(:,ii,ix);
    stripe_width_left(:,ii) = stripe_width_left_pool(:,ii,ix);
    stripe_width_right(:,ii) = stripe_width_right_pool(:,ii,ix);
end

end
```
