## Supplementary material for "Enhancer architecture and chromatin accessibility constrain phenotypic space during development": Script-MLP: remove_stripe_xings_iterate.m

```
function [stripe_cenx,stripe_ceny,stripe_width_left,stripe_width_right] = ...
    remove_stripe_xings_iterate(stripe_cenx,stripe_ceny,stripe_width_left,stripe_width_right)

nbins = size(stripe_cenx,1);
nstripes = size(stripe_cenx,2);

[xq,yq,xlq,xrq] = interpolate_stripes(stripe_cenx,stripe_ceny,stripe_width_left,stripe_width_right);

for jj_stripe = 1:nstripes-1
    % check if current stripe encroaches on right neighbor
    if sum(isnan(stripe_cenx(:,jj_stripe+1))) ~= nbins
        validix = ~isnan(stripe_cenx(:,jj_stripe));
        validneighborix = ~isnan(stripe_cenx(:,jj_stripe+1));
        
        if (sum(validix) > 0) && (sum(validneighborix) > 0)
            neighborbound_left = xq(:,jj_stripe+1) - xlq(:,jj_stripe+1);
            curbound_right = xq(:,jj_stripe) + xrq(:,jj_stripe);
            
            xing = find(curbound_right > neighborbound_left);
            if ~isempty(xing)
                ycoord_xings = yq([xing(1),xing(end)],jj_stripe);
                
                swapix_bin = stripe_ceny(:,jj_stripe) > ycoord_xings(1);
                
                stripe_cenx(swapix_bin,[jj_stripe,jj_stripe+1]) = stripe_cenx(swapix_bin,[jj_stripe+1,jj_stripe]);
                stripe_ceny(swapix_bin,[jj_stripe,jj_stripe+1]) = stripe_ceny(swapix_bin,[jj_stripe+1,jj_stripe]);
                stripe_width_left(swapix_bin,[jj_stripe,jj_stripe+1]) = stripe_width_left(swapix_bin,[jj_stripe+1,jj_stripe]);
                stripe_width_right(swapix_bin,[jj_stripe,jj_stripe+1]) = stripe_width_right(swapix_bin,[jj_stripe+1,jj_stripe]);
                
                ixs = [jj_stripe,jj_stripe+1];
                [xq(:,ixs),yq(:,ixs),xlq(:,ixs),xrq(:,ixs)] = ...
                    interpolate_stripes(stripe_cenx(:,ixs),stripe_ceny(:,ixs), ...
                    stripe_width_left(:,ixs),stripe_width_right(:,ixs));
            end
        end
    end
end

end
```
