## Supplementary material for "Enhancer architecture and chromatin accessibility constrain phenotypic space during development": Script-MLP: show_stripes_xy.m

```
function [xq,yq,xlq,xrq] = show_stripes_xy(stripe_cenx,stripe_ceny, ...
    stripe_width_left,stripe_width_right,ninterpts)

if nargin < 5
    ninterpts = [];
end

nstripes = size(stripe_cenx,2);

[xq,yq,xlq,xrq] = interpolate_stripes(stripe_cenx,stripe_ceny, ...
    stripe_width_left,stripe_width_right,ninterpts);

color_order = get(gca,'colororder');
hold on;
for kk = 1:nstripes
    plot(xq(:,kk),yq(:,kk),'LineWidth',3,'Color',color_order(mod(kk-1,7)+1,:));
    plot(xq(:,kk) - xlq(:,kk),yq(:,kk),'g','LineWidth',1,'Color',color_order(mod(kk-1,7)+1,:));
    plot(xq(:,kk) + xrq(:,kk),yq(:,kk),'g','LineWidth',1,'Color',color_order(mod(kk-1,7)+1,:));
end
hold off;

end
```
