## Supplementary material for "Enhancer architecture and chromatin accessibility constrain phenotypic space during development": Script-MLP: smooth_stripes.m

```
function varargout = ...
    smooth_stripes(stripe_cenx,stripe_ceny,stripe_width_left,stripe_width_right,slope_cutoff)

if nargin < 5 || isempty(slope_cutoff)
    slope_cutoff = 1;
    if nargin < 4
        stripe_width_right = [];
        if nargin < 3
            stripe_width_left = [];
        end
    end
end

[nbins,maxnstripes] = size(stripe_cenx);

for ii_stripe = 1:maxnstripes
    valix = ~isnan(stripe_cenx(:,ii_stripe));
    if sum(valix) > 2
        cur_stripe_cenx = stripe_cenx(valix,ii_stripe);
        cur_stripe_ceny = stripe_ceny(valix,ii_stripe);
        
        slope = abs(diff(cur_stripe_cenx)./diff(cur_stripe_ceny));
        
        if ~isempty(slope)
            over_cutoff = slope > slope_cutoff;
            ix2trash = false(nbins,1);
            
            valix_n = find(valix);
            npts = sum(valix)-1;
            for ii_pt = 1:npts
                if over_cutoff(ii_pt)
                    if ii_pt == 1
                        if ~over_cutoff(ii_pt+1)    % if endpoint
                            ix2trash(valix_n(ii_pt)) = true;
                        end
                    else
                        if over_cutoff(ii_pt-1)
                            ix2trash(valix_n(ii_pt)) = true;
                        elseif ii_pt == npts        % if endpoint
                            ix2trash(valix_n(ii_pt+1)) = true;
                        end
                    end
                end
            end
        end
        
        if any(ix2trash)
            disp(['Scrubbing stripe ' num2str(ii_stripe)]);
        end
        
        stripe_cenx(ix2trash,ii_stripe) = NaN;
        stripe_ceny(ix2trash,ii_stripe) = NaN;
        
        if ~isempty(stripe_width_left)
            stripe_width_left(ix2trash,ii_stripe) = NaN;
        end
        
        if ~isempty(stripe_width_right)
            stripe_width_right(ix2trash,ii_stripe) = NaN;
        end
    end
    
    if nargout == 4
        varargout = {stripe_cenx,stripe_ceny,stripe_width_left,stripe_width_right};
    else
        varargout = {stripe_cenx,stripe_ceny};
    end
end
```
