## Supplementary figures and images for "Enhancer architecture and chromatin accessibility constrain phenotypic space during development"

### TS_UnitTestImage.tif

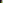
