## Supplemental Figures for "Enhancer architecture and chromatin accessibility constrain phenotypic space during development"

**SUPPLEMENTARY FIGURES:**

https://www.dropbox.com/sh/ayw581rxvzpu4p9/AAAMFvIo7IUPK-bdDXEyMIlya?dl=0

**SUPPLEMENTARY FIGURE LEGENDS**

**Figure S1.** (**a**) Sequence alignment of E3N sequences from 10 Drosophila species (top), and of E3N mutant sequences from the library (bottom). (**b**) Schematic of reporter gene construct used for integration into the *D. melanogaster* genome. (**c-f**) Protein staining of reporter gene expression driven by *E3N* (**c**), *eveS2* (**d**), *rhoNEE* (**e**) and *tinB* (**f**).

**Figure S2.** Assessment of fluorescence intensity across three different regions of a late-stage embryo, each region associated to a different germ-layer: A2 segment/ectoderm (**a** and **a'**, with DAPI overlay), mesoderm-derived tissue beneath A2 segment (**b** and **b'**, with DAPI overlay) and midgut/endoderm (**b** and **b'**). The same embryo is depicted in (**a**) and (**b**), but at different z-plans.

**Figure S3.** Number of mutations per enhancer variant *versus* changes in levels of expression for *twiPE* (**a**), *rhoNEE* (**b**) and *tinB* (**c**) enhancer lines.

**Figure S4.** *rhoNEE* enhancer variants show no evidence for ectopic expression during development; each line is represented at a mid- (left) and late- (right) embryonic stage.

**Figure S5.** *twiPE* enhancer variants show no evidence for ectopic expression during late-stage development.

**Figure S6.** (**a**) Examples of stained embryos from different *tinB* mutant variants.

**Figure S7.** Extensive activation is required to drive expression outside of native zones of expression. (**a-c**) Stage 5 embryos bearing the UAS::TALE-VP64 (TALEA) construct driven by the ubiquitous *nos*::*GAL4* driver and different *rhoNEE-lacZ* constructs, stained for *lacZ* RNA, including wildtype *rhoNEE* (**a**) and *rhoNEE* with either two (**b**) or three (**c**) TALEA binding sites. (**d**) Plots showing measurements of the indicated bounding boxes in panel (**a**). Centre line, mean; upper and lower limits, s.d.; whiskers, 95% confidence intervals (CIs).

**Figure S8.** DNA libraries used to explore the regulatory capacity of random DNA. Random DNA containing core motifs (Zelda and UAS are shown for reference) are cloned into the pLacZattB reporter construct (middle panel). The sequence lengths and nucleotide compositions are shown along the bottom panel.

**Figure S9.** Random DNA sequences display diverse tissue and cell-type-specific activity patterns. Shown are representative embryos across developmental stages. The DNA fragments tested are indicated in the left panels, with the associated stages and embryos across the right panels.

**Figure S10.** Protein staining showing that Zelda is expressed throughout *Drosophila melanogaster* development.

**Figure S11.** Late-embryo knockout of Zelda shows phenotypical consequences, namely extensive misregulation of ectodermal derived cell- and tissue-types.

**Figure S12.** Random DNA sequences biased with a Zelda motif display diverse tissue and cell-type-specific activity patterns. Shown are representative embryos across developmental stages. The DNA fragments tested are indicated in the left panels, with the associated stages and embryos across the right panels.

**Figure S13.** Adding Zelda motifs to endogenous enhancer sequences. (**a**) Examples of early stage embryos harboring a *rhoNEE* (top), *twiPE* (middle) or *eveS2* (bottom) enhancer with Zelda motif(s) on the left flank (left), on the right flank (centre) or on both flanks (right). (**b**) Examples of late stage embryos harboring a *rhoNEE* (first line), *twiPE* (second line), *eveS2* (third line) or *tinB* (fourth line) enhancer with Zelda motif(s) on the left flank (left), on the right flank (centre) or on both flanks (right). (**c**) Quantification of stripe width for stage5-embryos carrying *rhoNEE* enhancers containing different numbers of ectopic Zelda motifs (****$p<0.0001$, compared to wildtype; two-tailed t-test). (**d**) Quantification of nuclear intensities along the stripes for stage5-embryos carrying *rhoNEE* enhancers containing different numbers of ectopic Zelda motifs (****$p<0.0001$, each compared to wildtype; two-tailed t-test). (**e**) Normalised fluorescence intensities in the anterior region of stage5-embryos carrying *twiPE* enhancers containing different numbers of ectopic Zelda motifs (no statistical significance; two-tailed t-test). (**f-h**) Normalised fluorescence intensities across different regions along the anterior-posterior axis of stage5-embryos carrying *eveS2* enhancers containing different numbers of ectopic Zelda motifs.

**Figure S14.** Testing and mutating genomic sequences harboring a Ubx/Hth motif. Four out of seven regions tested are shown. Left (**a**, **d**, **g**, **j**): schematic of genomic region, location of selected sequence, Ubx binding (ChIP-seq) across the locus and sequence conservation across different *Drosophila* species. Centre (**b**, **e**, **h**, **k**): protein staining of late-stage embryos carrying genomic sequences represented on the left. Right (**c**, **f**, **i**, **l**): protein staining of late-stage embryos carrying genomic sequences represented on the left mutated for the Ubx/Hth motif.
