## Supplementary material for "Enhancer architecture and chromatin accessibility constrain phenotypic space during development": Script-GAC: CurateTable.m

```
%which ones
ManualRot=[2,5]

imwidth=1500;
FileList=dir('*.tif');
counter=1;
rotated=0;
num_col=5;

APint=readtable("AP_Intensities.csv");
APintC=table2cell(APint);
APintA=cell2mat(APintC(:,2:(imwidth+1)));

for j=1:length(FileList)
    j
   if 1==(sum(j==ManualRot))
    rotated=rotated+1
    counter
    subplot(5,num_col,counter);
    scatter(1:imwidth,fliplr(APintA(j,:)))
    APintA(j,:)=fliplr(APintA(j,:))
    text(1,20,string(j),'FontSize',16,'Color', 'r')
   elseif 0==(sum(j==ManualRot))
    subplot(5,num_col,counter);
    scatter(1:imwidth,APintA(j,:))
    text(1,20,string(j),'FontSize',16,'Color', 'r')
   end    
   counter=counter+1;
end

compMat=[{FileList.name}' num2cell(APintA)];
Tcomp=cell2table(compMat);
writetable(Tcomp,"AP_Intensities_Curated.csv")
```
