## Supplementary material for "Enhancer architecture and chromatin accessibility constrain phenotypic space during development": Script-GAC: feretProperties.m

```
function T = feretProperties(T)
% Copyright 2017-2018 The MathWorks, Inc.

maxfd = zeros(height(T),1);
maxfd_endpoints = cell(height(T),1);
maxfd_orientation = zeros(height(T),1);

minfd = zeros(height(T),1);
minfd_triangle_points = cell(height(T),1);
minfd_orientation = zeros(height(T),1);

minod = zeros(height(T),1);
minod_lower_points = cell(height(T),1);
minod_upper_points = cell(height(T),1);

minbb = cell(height(T),1);
minbb_a = zeros(height(T),1);

for k = 1:height(T)
    pixels = T.PixelList{k};
    V = pixelHull(pixels,'diamond');
    pairs = antipodalPairs(V);
    [maxfd(k),maxfd_endpoints{k}] = maxFeretDiameter(V,pairs);
    points = maxfd_endpoints{k};
    e = points(2,:) - points(1,:);
    maxfd_orientation(k) = atan2d(e(2),e(1));
    
    [minfd(k),minfd_triangle_points{k}] = minFeretDiameter(V,pairs);
    points = minfd_triangle_points{k};
    e = points(2,:) - points(1,:);
    thetad = atan2d(e(2),e(1));
    minfd_orientation(k) = mod(thetad + 180 + 90,360) - 180;
    
    [minod(k),minod_lower_points{k},minod_upper_points{k}] = ...
        feretDiameter(V,maxfd_orientation(k)+90);
    
    [minbb{k},minbb_a(k)] = minAreaBoundingBox(V,pairs);
end

T.MaxFeretDiameter = maxfd;
T.MaxFeretDiameterEndpoints = maxfd_endpoints;
T.MaxFeretDiameterOrientation = maxfd_orientation;
T.MinFeretDiameter = minfd;
T.MinFeretDiameterTrianglePoints = minfd_triangle_points;
T.MinFeretDiameterOrientation = minfd_orientation;
T.OrthogonalDiameter = minod;
T.OrthogonalDiameterLowerPoints = minod_lower_points;
T.OrthogonalDiameterUpperPoints = minod_upper_points;
T.MinAreaBoundingBox = minbb;
T.MinAreaBoundingBoxArea = minbb_a;
end

function [bb,A] = minAreaBoundingBox(V,antipodal_pairs)
% Copyright 2017-2018 The MathWorks, Inc.

if nargin < 2
    antipodal_pairs = antipodalPairs(V);
end

n = size(antipodal_pairs,1);
p = antipodal_pairs(:,1);
q = antipodal_pairs(:,2);

A = Inf;
thetad = [];

for k = 1:n
    if k == n
        k1 = 1;
    else
        k1 = k+1;
    end
    
    pt1 = [];
    pt2 = [];
    pt3 = [];
    
    if (p(k) ~= p(k1)) && (q(k) == q(k1))
        pt1 = V(p(k),:);
        pt2 = V(p(k1),:);
        pt3 = V(q(k),:);
        
    elseif (p(k) == p(k1)) && (q(k) ~= q(k1))
        pt1 = V(q(k),:);
        pt2 = V(q(k1),:);
        pt3 = V(p(k),:);
    end
    
    if ~isempty(pt1)
        % Points pt1, pt2, and pt3 are possibly on the minimum-area
        % bounding box, with points pt1 and pt2 forming an edge coincident with
        % the bounding box. Call the height of the triangle with base
        % pt1-pt2 the height of the bounding box, h.
        
        h = triangleHeight(pt1,pt2,pt3);
        
        pt1pt2_direction = atan2d(pt2(2)-pt1(2),pt2(1)-pt1(1));
        
        w = feretDiameter(V,pt1pt2_direction);
        
        A_k = h*w;
        if (A_k < A)
            A = A_k;
            thetad = pt1pt2_direction;
        end
    end
end

% Rotate all the points so that pt1-pt2 for the minimum bounding box points
% straight up.

r = 90 - thetad;
cr = cosd(r);
sr = sind(r);
R = [cr -sr; sr cr];

Vr = V * R';

xr = Vr(:,1);
yr = Vr(:,2);
xmin = min(xr);
xmax = max(xr);
ymin = min(yr);
ymax = max(yr);

bb = [ ...
    xmin ymin
    xmax ymin
    xmax ymax
    xmin ymax
    xmin ymin ];

% Rotate the bounding box points back.
bb = bb * R;
end

function h = triangleHeight(P1,P2,P3)
% Copyright 2017-2018 The MathWorks, Inc.

h = 2 * abs(signedTriangleArea(P1,P2,P3)) / norm(P1 - P2);
end

function area = signedTriangleArea(A,B,C)
% Copyright 2017-2018 The MathWorks, Inc.

area = ( (B(1) - A(1)) * (C(2) - A(2)) - ...
    (B(2) - A(2)) * (C(1) - A(1)) ) / 2;
end

function [d,V1,V2] = feretDiameter(V,theta)
% Copyright 2017-2018 The MathWorks, Inc.

% Rotate points so that the direction of interest is vertical.

alpha = 90 - theta;

ca = cosd(alpha);
sa = sind(alpha);
R = [ca -sa; sa ca];

% Vr = (R * V')';
Vr = V * R';

y = Vr(:,2);
ymin = min(y,[],1);
ymax = max(y,[],1);

d = ymax - ymin;

if nargout > 1
    V1 = V(y == ymin,:);
    V2 = V(y == ymax,:);
end
end

function [d,end_points] = maxFeretDiameter(P,antipodal_pairs)
% Copyright 2017-2018 The MathWorks, Inc.

if nargin < 2
    antipodal_pairs = antipodalPairs(P);
end

v = P(antipodal_pairs(:,1),:) - P(antipodal_pairs(:,2),:);
D = hypot(v(:,1),v(:,2));
[d,idx] = max(D,[],1);
P1 = P(antipodal_pairs(idx,1),:);
P2 = P(antipodal_pairs(idx,2),:);

end_points = [P1 ; P2];
end

function [d,triangle_points] = minFeretDiameter(V,antipodal_pairs)
% Copyright 2017-2018 The MathWorks, Inc.

if nargin < 2
    antipodal_pairs = antipodalPairs(V);
end

n = size(antipodal_pairs,1);
p = antipodal_pairs(:,1);
q = antipodal_pairs(:,2);

d = Inf;
triangle_points = [];

for k = 1:n
    if k == n
        k1 = 1;
    else
        k1 = k+1;
    end
    
    pt1 = [];
    pt2 = [];
    pt3 = [];
    
    if (p(k) ~= p(k1)) && (q(k) == q(k1))
        pt1 = V(p(k),:);
        pt2 = V(p(k1),:);
        pt3 = V(q(k),:);
        
    elseif (p(k) == p(k1)) && (q(k) ~= q(k1))
        pt1 = V(q(k),:);
        pt2 = V(q(k1),:);
        pt3 = V(p(k),:);
    end
    
    if ~isempty(pt1)
        % Points pt1, pt2, and pt3 form a possible minimum Feret diameter.
        % Points pt1 and pt2 form an edge parallel to caliper direction.
        % The Feret diameter orthogonal to the pt1-pt2 edge is the height
        % of the triangle with base pt1-pt2.
        
        d_k = triangleHeight(pt1,pt2,pt3);
        if d_k < d
            d = d_k;
            triangle_points = [pt1; pt2; pt3];
        end
    end
end
end      


function V = pixelHull(P,type)

if nargin < 2
    type = 24;
end

if islogical(P)
    P = bwperim(P);
    [i,j] = find(P);
    P = [j i];
end

if strcmp(type,'square')
    offsets = [ ...
         0.5  -0.5
         0.5   0.5
        -0.5   0.5
        -0.5  -0.5 ];

elseif strcmp(type,'diamond')
    offsets = [ ...
         0.5  0
         0    0.5
        -0.5  0
         0   -0.5 ];

else
    % type is number of angles for sampling a circle of diameter 1.
    thetad = linspace(0,360,type+1)';
    thetad(end) = [];

    offsets = 0.5*[cosd(thetad) sind(thetad)];
end

offsets = offsets';
offsets = reshape(offsets,1,2,[]);

Q = P + offsets;
R = permute(Q,[1 3 2]);
S = reshape(R,[],2);

k = convhull(S,'Simplify',true);
V = S(k,:);
end


function h = drawFullLine(ax,point,angle_degrees,varargin)
%drawFullLine Draw a line that spans the entire plot
%
%    drawFullLine(ax,point,angle_degrees) draws a line in the
%    specified axes that goes through the specified point at the
%    specified angle (in degrees). The line is drawn to span the
%    entire plot.
%
%    drawFullLine(___,Name,Value) passes name-value parameter pairs
%    to the line function.

% Steve Eddins


limits = axis(ax);
width = abs(limits(2) - limits(1));
height = abs(limits(4) - limits(3));
d = 2*hypot(width,height);
x1 = point(1) - d*cosd(angle_degrees);
x2 = point(1) + d*cosd(angle_degrees);
y1 = point(2) - d*sind(angle_degrees);
y2 = point(2) + d*sind(angle_degrees);
h = line(ax,'XData',[x1 x2],'YData',[y1 y2],varargin{:});
end

function pq = antipodalPairs(S)
% antipodalPairs Antipodal vertex pairs of simple, convex polygon.
%
%   pq = antipodalPairs(S) computes the antipodal vertex pairs of a simple,
%   convex polygon. S is a Px2 matrix of (x,y) vertex coordinates for the
%   polygon. S must be simple and convex without repeated vertices. It is
%   not checked for satisfying these conditions. S can either be closed or
%   not. The output, pq, is an Mx2 matrix representing pairs of vertices in
%   S. The coordinates of the k-th antipodal pair are S(pq(k,1),:) and
%   S(pq(k,2),:).
%
%   TERMINOLOGY
%
%   For a convex polygon, an antipodal pair of vertices is one where you
%   can draw distinct lines of support through each vertex such that the
%   lines of support are parallel.
%
%   A line of support is a line that goes through a polygon vertex such
%   that the interior of the polygon lies entirely on one side of the line.
%
%   EXAMPLE
%
%     Compute antipodal vertices of a polygon and plot the corresponding
%     line segments.
%
%       x = [0 0 1 3 5 4 0];
%       y = [0 1 4 5 4 1 0];
%       S = [x' y'];
%       pq = antipodalPairs(S);
%
%       plot(S(:,1),S(:,2))
%       hold on
%       for k = 1:size(pq,1)
%           xk = [S(pq(k,1),1) S(pq(k,2),1)];
%           yk = [S(pq(k,1),2) S(pq(k,2),2)];
%           plot(xk,yk,'LineStyle','--','Marker','o','Color',[0.7 0.7 0.7])
%       end
%       hold off
%       axis equal
%
%   ALGORITHM NOTES
%
%   This function uses the "ANTIPODAL PAIRS" algorithm, Preparata and
%   Shamos, Computational Geometry: An Introduction, Springer-Verlag, 1985,
%   p. 174.

%   Steve Eddins


n = size(S,1);

if isequal(S(1,:),S(n,:))
    % The input polygon is closed. Remove the duplicate vertex from the
    % end.
    S(n,:) = [];
    n = n - 1;
end

% The algorithm assumes the input vertices are in counterclockwise order.
% If the vertices are in clockwise order, reverse the vertices.
clockwise = simplePolygonOrientation(S) < 0;
if clockwise
    S = flipud(S);
end

% The following variables, including the two anonymous functions, are set
% up to follow the notation in the pseudocode on page 174 of Preparata and
% Shamos. p and q are indices (1-based) that identify vertices of S. p0 and
% q0 identify starting vertices for the algorithm. area(i,j,k) is the area
% of the triangle with the corresponding vertices from S: S(i,:), S(j,:),
% and S(k,:). next(p) returns the index of the next vertex of S.
%
% The initialization of p0 is missing from the Preparata and Shamos text.
area = @(i,j,k) signedTriangleArea(S(i,:),S(j,:),S(k,:));
next = @(i) mod(i,n) + 1; % mod((i-1) + 1,n) + 1
p = n;
p0 = next(p);
q = next(p);

% The list of antipodal vertices will be built up in the vectors pp and qq.
pp = zeros(0,1);
qq = zeros(0,1);

% ANTIPODAL PAIRS step 3.
while (area(p,next(p),next(q)) > area(p,next(p),q))
    q = next(q);
end
q0 = q;    % Step 4.

while (q ~= p0)    % Step 5.
    p = next(p);   % Step 6.
    % Step 7. (p,q) is an antipodal pair.
    pp = [pp ; p];
    qq = [qq ; q];

    % Step 8.
    while (area(p,next(p),next(q)) > area(p,next(p),q))
        q = next(q);    % Step 9.
        if ~isequal([p q],[q0,p0])
            % Step 10.
            pp = [pp ; p];
            qq = [qq ; q];
        else
            % This loop break is omitted from the Preparata and Shamos
            % text.
            break
        end
    end

    % Step 11. Check for parallel edges.
    if (area(p,next(p),next(q)) == area(p,next(p),q))
        if ~isequal([p q],[q0 n])
            % Step 12. (p,next(q)) is an antipodal pair.
            pp = [pp ; p];
            qq = [qq ; next(q)];
        else
            % This loop break is omitted from the Preparata and Shamos
            % text.
            break
        end
    end
end

if clockwise
    % Compensate for the flipping of the polygon vertices.
    pp = n + 1 - pp;
    qq = n + 1 - qq;
end

pq = [pp qq];
end

function s = vertexOrientation(P0,P1,P2)
% vertexOrientation  Orientation of a vertex with respect to line segment.
%
%   s = vertexOrientation(P0,P1,P2) returns a positive number if P2 is to
%   the left of the line through P0 to P1. It returns 0 if P2 is on the
%   line. It returns a negative number if P2 is to the right of the line.
%
%   Stating it another way, a positive output corresponds to a
%   counterclockwise traversal from P0 to P1 to P2.
%
%   P0, P1, and P2 are two-element vectors containing (x,y) coordinates.
%
%   Reference: http://geomalgorithms.com/a01-_area.html, function isLeft()

% Steve Eddins


s = (P1(1) - P0(1)) * (P2(2) - P0(2)) - ...
    (P2(1) - P0(1)) * (P1(2) - P0(2));
end

function s = simplePolygonOrientation(V)
% simplePolygonOrientation  Determine vertex order for simple polygon.
%
%   s = simplePolygonOrientation(V) returns a positive number if the simple
%   polygon V is counterclockwise. It returns a negative number of the
%   polygon is clockwise. It returns 0 for degenerate cases. V is a Px2
%   matrix of (x,y) vertex coordinates.
%
%   Reference: http://geomalgorithms.com/a01-_area.html, function
%   orientation2D_Polygon()

% Steve Eddins


n = size(V,1);

if n < 3
    s = 0;
    return
end

% Find rightmost lowest vertext of the polygon.

x = V(:,1);
y = V(:,2);
ymin = min(y,[],1);
y_idx = find(y == ymin);
if isscalar(y_idx)
    idx = y_idx;
else
    [~,x_idx] = max(x(y_idx),[],1);
    idx = y_idx(x_idx(1));
end

% The polygon is counterclockwise if the edge leaving V(idx,:) is left of
% the entering edge.

if idx == 1
    s = vertexOrientation(V(n,:), V(1,:), V(2,:));
elseif idx == n
    s = vertexOrientation(V(n-1,:), V(n,:), V(1,:));
else
    s = vertexOrientation(V(idx-1,:), V(idx,:), V(idx+1,:));
end
end
```
