## Supplementary material for "Enhancer architecture and chromatin accessibility constrain phenotypic space during development": Script-GAC: GetIntensities.m

```
function [Intensities,stdim,ymidpoint,stdimOrig,stdimhori,stT,x1feret,x2feret,x1feretMin,x2feretMin]=GetIntensities(Image,valthresh)
    
    iptsetpref('ImshowInitialMagnification','fit');
    filename = Image;
     
    im = imread(filename);
   
    %% max of intensities
    RGBmax = max(im,[],3);

    %% Generate mask: assumes brightness inside embryo above the selected threshold in the GUI
    
    mask = im2bw(RGBmax,valthresh/255);
    
    %% smooth boundaries
    mask = medfilt2(mask,[3,3]); %median filtering to remove noise
    mask = imclose(mask,strel('disk',5,0)); %closure operation to smoothen the boundary

    %% properties for rotation
    L = bwlabel(mask);
    
    statsAr=regionprops(L,'Area');
    statsArea=cell2mat(struct2cell(statsAr));
    [Ma,IndM]=max(statsArea);
    
    %stats before rotation
    statsTBR = regionprops('table',L,'PixelList');
    stTBR = feretProperties(statsTBR);

    %% Rotate the original image
    stdim = imrotate(L,stTBR(IndM,:).MaxFeretDiameterOrientation+180);

    %imshow(stdim)
    statsT = regionprops('table',stdim,'PixelList');
    stT = feretProperties(statsT(IndM,:));

    ymidpoint=(stT.MaxFeretDiameterEndpoints{1}(4));

    x1feret=[(stT.MaxFeretDiameterEndpoints{1}(1)) (stT.MaxFeretDiameterEndpoints{1}(3))];
    x2feret=[(stT.MaxFeretDiameterEndpoints{1}(2)) (stT.MaxFeretDiameterEndpoints{1}(4))];
    x1feretMin=[(stT.OrthogonalDiameterLowerPoints{1}(1)) (stT.OrthogonalDiameterLowerPoints{1}(2))];
    x2feretMin=[(stT.OrthogonalDiameterUpperPoints{1}(1)) (stT.OrthogonalDiameterUpperPoints{1}(2))];
    
    %putative horizontal start and end for the embryo
    imst=round(stT.MaxFeretDiameterEndpoints{1}(1));
    imen=round(stT.MaxFeretDiameterEndpoints{1}(2));
    
    yhalfheight= round(stT.MinFeretDiameter/7);
    
    stdimOrig=imrotate(im,stTBR(IndM,:).MaxFeretDiameterOrientation+180);

    %crop embryo based on the putative start: imst and the putative end : imen
    if imst<imen
        stdimhori = stdimOrig(ymidpoint-yhalfheight:ymidpoint+yhalfheight,imst:imen,:);
    else 
        stdimhori = stdimOrig(ymidpoint-yhalfheight:ymidpoint+yhalfheight,imen:imst,:);
    end

    %% get average intensity per column
    expression = stdimhori; %in case more channels %squeeze(stdimhori);
    %get intensities levels
    Intensities=mean(expression);
    
end


%%
```
