## Supplementary material for "Enhancer architecture and chromatin accessibility constrain phenotypic space during development": Script-GAC: IntensitiesMatrix.m

```
%open all tif images in a directory and call the function that generates intensities
function IntensitiesMatrix(chosenvalthresh,brightval)

imwidth=1500;

%open list of tif files in the current directory
FileList=dir('*.tif');

matrixInt=zeros(length(FileList),imwidth);

%number of columns for subplot
num_col=5;

if num_col>length(FileList)
    num_col=length(FileList)
end

FileList=dir('*.tif');
counter=1;
rotated=0


for j=1:length(FileList)
    ImEmb=FileList(j).name;
    %get intensities for the j embryo and its Feret properties for rotation
    [Intens,StDim,ym,StDimOrigi,StDimHori,stTab,x1fe,x2fe,x1fm,x2fm]=GetIntensities(ImEmb,chosenvalthresh);
    
    [rows, columns]=size(Intens);
    matrixInt(j,1:columns)=Intens;

    subplot(3*ceil(length(FileList)/num_col),num_col,counter);
    
    binaryImage = im2bw(StDim);
    imshow(binaryImage)
    axis on
    hold on;
    boundaries = bwboundaries(binaryImage);
    %
    x = boundaries{1}(:, 2);
    y = boundaries{1}(:, 1);
    
    
    plot(x1fm(1), x1fm(2), 'y+', 'MarkerSize', 10, 'LineWidth',2);
    plot(x2fm(1), x2fm(2), 'y+', 'MarkerSize', 10, 'LineWidth',2);
    plot(x2fe(1), x2fe(2), 'g+', 'MarkerSize', 20, 'LineWidth',2);
    plot(x1fe(1), x1fe(2), 'b+', 'MarkerSize', 20, 'LineWidth',2);
    text(x1fe(1), x1fe(2)-250,string(j),'FontSize',16,'Color', 'r' )
    %   plot(100,ym,'r+','MarkerSize',30,'LineWidth',2);
    hold off;
    subplot(3*ceil(length(FileList)/num_col),num_col,counter+1);
    %embryo number
    j
    
    avx=(x1fm(1)+x2fm(1)/2);
    %increase brightness overall useful for some batches 
    StDimOrigi=StDimOrigi+brightval;
    
    if abs(x2fm(1)-max(x1fe(1),x2fe(1)))>=abs(x2fm(1)-min(x1fe(1),x2fe(1)))
        %imshow(imrotate(StDimHori,-180))
        StDimHori=imrotate(StDimHori,-180);
        
        StDimOrigi=imrotate(StDimOrigi,-180);
        imagesc(StDimOrigi);
        rotated=rotated+1
        subplot(3*ceil(length(FileList)/num_col),num_col,counter+2);
        scatter(1:imwidth,fliplr(matrixInt(j,:)))
        matrixInt(j,:)=fliplr(matrixInt(j,:));
        
    elseif abs(x2fm(1)-max(x1fe(1),x2fe(1)))<abs(x2fm(1)-min(x1fe(1),x2fe(1)))
        
        %imshow(StDimHori)
        imagesc(StDimOrigi);
        subplot(3*ceil(length(FileList)/num_col),num_col,counter+2);
        scatter(1:imwidth,matrixInt(j,:))
    end
    
    counter=counter+3;
end

compMat=[{FileList.name}' num2cell(matrixInt)];
Tcomp=cell2table(compMat);
writetable(Tcomp,"AP_Intensities.csv");
end
```
