## Supplementary material for "Enhancer architecture and chromatin accessibility constrain phenotypic space during development": Script-GAC: Quant_EveZelda.R

```
#library("ggplot2")
library(ggplot2)
library(reshape2)
library(grid)
library(gridExtra)
library(boot)
library(smoother)
library(scales) 
library(gsubfn)


#directory
dirQu<-"/Users/Documents/EveZelda/"
files_list<-list.files(dirQu)

#look for the curated files
files<-files_list[grep("Curated",files_list)]


Addpseudo <- function(data) {
  
  if (is.vector(data)) {
    noise <- runif(length(data), 0, 0.00001)
    addednoise <- data + noise
  } else {
    length <- dim(data)[1] * dim(data)[2]
    noise <- matrix(runif(length, 0, 0.00001), dim(data)[1])
    addednoise <- data + noise
  }
  return(addednoise)
}


Nbins=600
StNm=1
valth=0

MeanStd<-function(FileName){
  newTab<-NULL
  newTabBcd<-NULL
  #smoothing parameter
  wstmh=35
  File<-read.csv(paste0(dirQu,FileName))
  
  valth=0
  indval=5
  
  #extract name from file
  qtitle<-unlist(strsplit(FileName,"_"))
  finame<-qtitle[grep("[0-9]+\\w",qtitle,perl = TRUE)]
  
  #initialize plotting panel for individual embryos
  png(paste0(dirQu,"orig_all_",finame,".png"), width = 1000, height = 1000)
  par(mfrow=c(5,5))
  
  for (i in 1:dim(File)[1]){
    
    ys = smth(as.numeric(as.character(File[i,])),window = wstmh,method = "gaussian") #SMOOTHING
    
    ys[is.na(ys)]<-0
    
    #remove image background
    valth=max(ys)/indval
    ys<-((ys[ys>valth]))
    
    xsl=1:length(ys)
    #linear interpolation for N bins
    appy=approx(xsl, ys, method = "linear",n=Nbins)

    ys=appy$y
    
    last20<-(round(length(ys)*0.8)):(round(length(ys)*0.99))
    
    #embryo background
    thresh=quantile(ys[last20],0.1)

    #from here for changing Ys to different 
    ys1t=ys-thresh
    ys1t[ys1t<=0]<-0

    #normalize by 50% of the signal in background area
    normdif<-quantile(ys1t[round(Nbins*0.75):round(Nbins*0.99)],0.50)*1

    if (normdif>0){
      norm_Vec<-(ys1t)/normdif

      newTab<-rbind(newTab,norm_Vec)
    }  else{
      print("Negative, discard or explore outlier")
      print(finame)
      print(i)
      next;
    }
    plot(1:length(appy$y),appy$y)

  }
  dev.off()
  #bootstrapping function
  boot.mean<-function(x,i){boot.mean<-mean(x[i])}
  #add very small noise for data that is equal zero to avoid bootstrapping issues
  newTab[which(is.nan(newTab))]<-runif(1,0,0.001)
  newTab=Addpseudo(newTab)
  dfnewTab<-as.data.frame(newTab)
  write.csv(dfnewTab,paste0(dirQu,"Expr_Embryo",finame),row.names = F, col.names = T, sep="\t",quote = F)
  b95<-apply(newTab,2,function(y){b<-boot(y,boot.mean,1000); c(mean(b$t),boot.ci(b,type="perc",conf=0.95)$percent[4:5])})
  #centralize to 1? test
  b95=b95-1
  b95[b95<=0]<-0
  
  #statistics
  nTmean<-b95[1,]
  nTsde<-b95[2,]
  nTsde2<-b95[3,]
  
  tabdim=0*Nbins
  print(finame)
  nTT<-cbind(0:length(ys1t)/Nbins*100,nTmean,nTsde,nTsde2)
  colnames(nTT)<-c("indx","mean_nT","sd_nT","sd_nT2")
  
  nTT<-as.data.frame(nTT)
  ExpressionList<-list("Table_Expression"=nTT)#"TableExpressionwBcd"=nBcdnewTab)
  return(list(ExpressionList,dim(File)[1]))
}  


titles<-unlist(strsplit(files,"Curated_"))
titleCode<-titles[grep("[0-9]+\\w",titles,perl = TRUE)]
titleCode=titles[seq(2, length(titles), 2)]
plot_list<- list()
plot_num=1

nfiles=length(files)

allexpn<-NULL

#generate data frames for plots 

for (j in 1:nfiles){


  Q423<-files[j]
  AllOutList<-MeanStd(Q423)
  OutList<-AllOutList[[1]]

  b<-AllOutList[[2]]
  ExpnT<-OutList$Table_Expression
  sav_val<-c(titleCode[j],ExpnT[1:60,2])
  tabsav<-rbind(tabsav,sav_val)
  #plot from 
  cutExpnT<-ExpnT

  cutExpnT<-cbind(cutExpnT,cutExpnT$mean_nT/max(cutExpnT$mean_nT))
  colnames(cutExpnT)[5]<-"normMean"

  cutExpnT<-cbind(cutExpnT,rep(titleCode[j],dim(cutExpnT)[1]))
  colnames(cutExpnT)[6]<-c("Embryo")
  allexpn<-rbind(allexpn,cutExpnT)
  
  
  plot_list[[plot_num]]<-ggplot(cutExpnT)+
    geom_line(aes(indx,mean_nT),group=1)+
    geom_ribbon(aes(x=indx,ymax=sd_nT2,ymin=sd_nT),alpha=0.6,fill="skyblue")+
    labs(title=paste0(titleCode[j],"  N_embryos:",b))+
    xlab("AP %")+ylab("Normalized Intensity")

  #plot counter for list object
  plot_num=plot_num+1
}

ml <- marrangeGrob(plot_list, nrow=1, ncol=1)
ggsave(paste0(dirQu,"BootstrappedExpresionProfiles.pdf"), ml,limitsize = FALSE)
```
